# A global view of human centromere variation and evolution

**DOI:** 10.64898/2025.12.09.693231

**Authors:** Shenghan Gao, Keisuke K. Oshima, Shu-Cheng Chuang, Mark Loftus, Annalaura Montanari, David S. Gordon, Human Genome Structural Variation Consortium, Human Pangenome Reference Consortium, PingHsun Hsieh, Miriam K. Konkel, Mario Ventura, Glennis A. Logsdon

## Abstract

Centromeres are essential for accurate chromosome segregation during cell division, yet their highly repetitive sequence has historically hindered their complete assembly and characterization. Consequently, the full spectrum of centromere diversity across individuals, populations, and evolutionary contexts remains largely unexplored. Here, we address this gap in knowledge by assembling and characterizing 2,110 complete human centromeres from a diverse cohort of individuals representing 5 continental and 28 population groups. By developing a novel suite of bioinformatic tools tailored for centromeric regions, we uncover previously unknown variation within centromeres, including 226 novel centromere haplotypes and 1,870 new α-satellite higher-order repeat (HOR) variants. We find that mobile element insertions are present in 30% of centromeres, with chromosome 16 harboring *Alu* elements within the kinetochore site at an 11-fold higher frequency than expected. While most centromeres have a single kinetochore site, 6% of them have di-kinetochores, and <<1% have tri-kinetochores, which we confirm with long-read CENP-A CUT&RUN, DiMeLo-seq, and multi-generational inheritance. We further show that the position of the kinetochore is not random and is, instead, closely associated with the underlying sequence and structure of the centromere. To understand the nature of evolutionary change, we compared 2,110 complete human centromeres to 5,747 complete centromeres recently assembled from the Human Pangenome Reference Consortium. We show that centromeres have a >50-fold variation in mutation rate, with the most rapidly mutating centromeres on chromosome 1 and the slowest mutating centromeres on chromosome Y. Additionally, a subset of centromeres show evidence of introgression from archaic hominins, shaping their sequence, structure, and evolutionary history. We validate these centromere mutation rates in a four-generation family, spanning 28 family members and 483 accurately assembled centromeres, and show that the kinetochore site is the most rapidly mutating region in the centromere, with twofold more single-nucleotide variants than the rest of the centromeric α-satellite HOR array on average. We propose a model that reveals an ‘arms race’ between centromeric sequence and proteins, with frequent mutations within the site of the kinetochore that lead to changes in genetic and epigenetic landscapes and, ultimately, rapid evolution of these critically important regions.

## INTRODUCTION

Centromeres are essential chromosomal loci that ensure the accurate segregation of chromosomes during mitosis and meiosis. Despite their fundamental importance, their sequence, structure, and variation among the human population are not yet fully known. Centromeres have been largely excluded from variation analyses because they are composed of near-identical tandem repeats known as α-satellite^1–4^, which challenge efforts to assess their variation using standard methods and approaches. Recent advances in long-read sequencing and telomere-to-telomere (T2T) genome assembly algorithms, however, are now making it possible to resolve entire centromeric regions^5–11^. Building on these technological developments, population-scale genome projects have begun to reveal that centromeres harbor far more genetic, epigenetic, and structural diversity than previously appreciated^12–14^. Variation in α-satellite higher-order repeat (HOR) array length, composition, structural organization, and kinetochore position has been uncovered in limited cohorts^15^, but a comprehensive understanding of the full scope of human centromere variation among global populations remains unresolved. Moreover, fundamental questions about how centromeres evolve, how quickly they mutate, and how sequence and chromatin features interact to specify kinetochore sites remain largely unknown.

Here, we address this knowledge gap by assembling and characterizing 2,110 complete centromeres from 65 individuals representing 5 continental and 28 population groups and subsequently validating our findings with a broader set of 6,230 centromeres from 227 diverse individuals and a 28-member pedigree (8,340 centromeres total). By developing new computational tools and pipelines optimized for centromeric regions, we define the landscape of centromere sequence and structural variation, uncover previously unknown haplotypes and α-satellite HOR variants, and map kinetochore positions across all chromosomes. We further integrate long-read methylation data, CENP-A chromatin profiling, multi-generational inheritance, and archaic sequencing data to characterize the functional and evolutionary events shaping centromeres. Together, these analyses provide a comprehensive foundation for understanding how human centromeres vary and evolve and how their underlying architecture supports faithful chromosome segregation during cell division.

## RESULTS

### Complete sequence of 2,110 diverse human centromeres

The recent release of 65 near-complete human genome assemblies spanning 5 continental and 28 population groups by the Human Genome Structural Variation Consortium^12^ (HGSVC; **Supplementary Table 1**) provides a unique opportunity to investigate the diversity in human centromere sequence and structure, variation in epigenetic landscape, and differences in evolutionary trajectories on a global scale. Therefore, we first sought to resolve the sequence of each centromere in these 65 genomes. To do this, we identified centromeres that were completely assembled from p to q arm and free of large sequence assembly errors (**Methods**), identifying 1,246 such centromeres, as previously reported^12^.

Recognizing that centromeres generated by different genome assemblers (e.g. Verkko^16^ and hifiasm^17^) have complementary error profiles that could be leveraged to repair one assembly with the other^12^ (**Supplementary Fig. 1**), we then developed a new tool, AssemblyRepairer, to repair the remaining errors in the Verkko centromeres with the corresponding centromeres assembled with hifiasm. AssemblyRepairer utilizes large, unique *k*-mers shared between the Verkko and hifiasm assemblies to identify corresponding contigs and then patches erroneous regions in the Verkko contigs with the correctly assembled, equivalent sequences in the hifiasm contigs (**Supplementary Fig. 2a**, **Methods**). Using this tool, we repaired an additional 864 centromeres, for a total combined set of 2,110 completely and accurately assembled centromeres (**Supplementary Fig. 2b-d**), a 67% increase over the previously published dataset^12^. We confirmed the accurate assembly of each centromere with a suite of computational tools and approaches, including native long-read mapping as well as validation with HMM-Flagger^18^, NucFlag^19^, and GAVISUNK^20^ (**Supplementary Note 1, Supplementary Table 2**). Additionally, we estimated the quality value (QV) of the centromeres using short-read Illumina sequencing data generated from the same source genome with Merqury^21^ and found them to have an average QV of 60.99, indicating that the centromeres were >99.9999% accurate (**Supplementary Table 3**).

The complete sequence of 2,110 diverse human centromeres allowed us to investigate their genetic and epigenetic variation and determine their haplotype frequency across global populations for the first time. To do this, we developed a new pipeline called CenMAP (<u>cen</u>tromere <u>m</u>apping and <u>a</u>nnotation <u>p</u>ipeline), which automatically annotates centromeric regions at the sequence level to define the active and inactive centromeric α-satellite HOR arrays as well as divergent/monomeric α-satellite, determines the location of hypomethylated regions in the centromere known as “Centromere Dip Regions” (CDRs^22^), which are associated with the site of the kinetochore^6,8^, and generates one-dimensional (1D) local sequence identity heat maps to show the evolutionary relationship of the underlying sequences (**Fig. 1, Supplementary Figs. 3-6**). Applying this pipeline to all 2,110 centromeres revealed 226 new major centromere haplotypes (**Fig. 1, Supplementary Figs. 3-6**) as well as 1,870 new α-satellite HOR variants compared to CHM13 and CHM1 (**Supplementary Table 4**), 97% of which occur in at least one other genome in our dataset or in the broader set of genome assemblies recently released by the Human Pangenome Reference Consortium (HPRC; **Supplementary Table 5; Methods**). Additionally, comparison of these newly assembled centromeres to those in the CHM1 and CHM13 genomes revealed that for 7 out of 24 chromosomes (chromosomes 1, 3, 4, 6, 10, 11 and 13), the most common centromere haplotype was not represented by CHM13 or CHM1 and was instead represented by another haplotype (**Fig. 1, Supplementary Figs. 3-6**), underscoring the benefit of additional genomes for a more comprehensive understanding of centromere variation.

**Figure 1.**
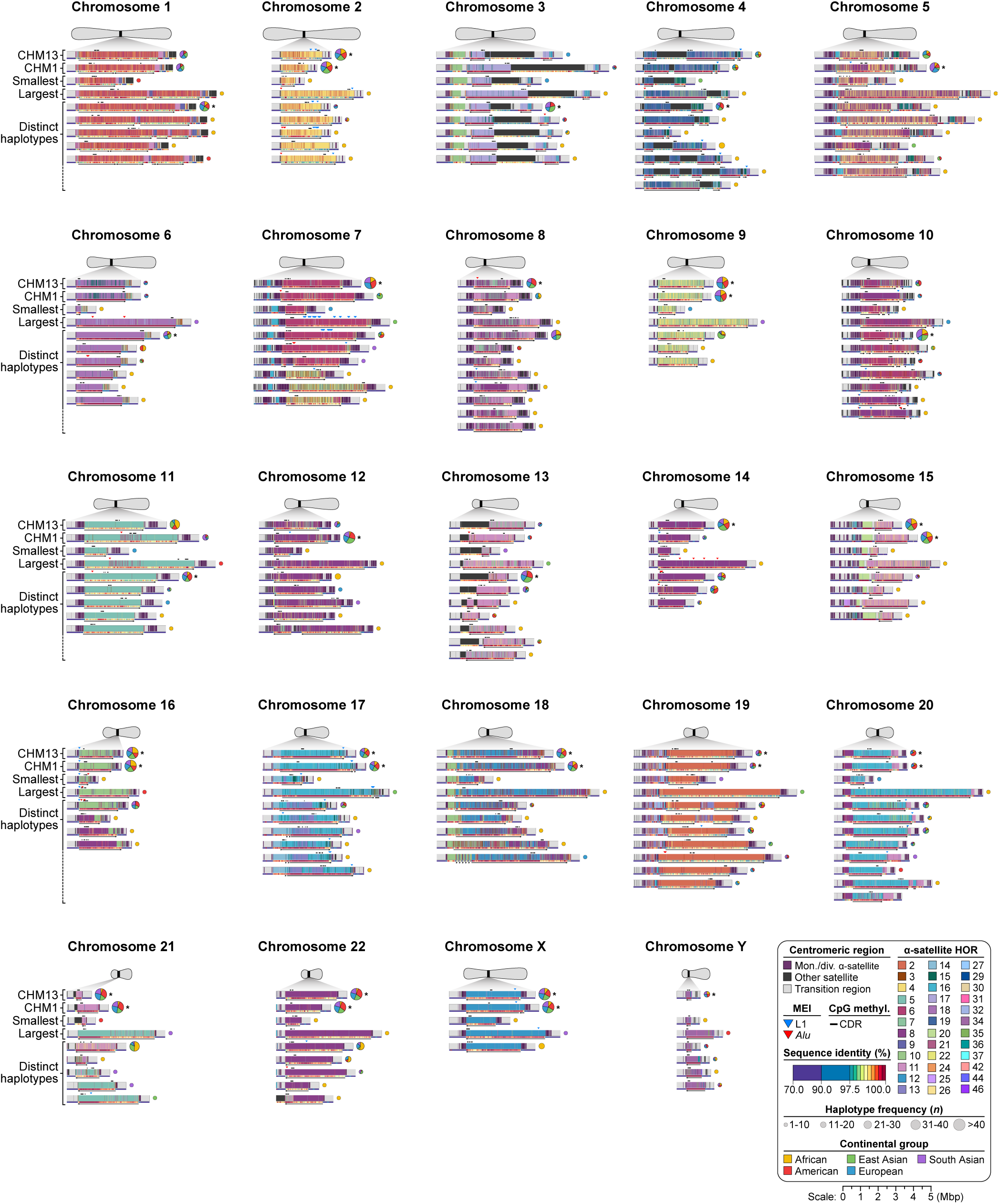
Complete maps and haplotype frequencies of diverse human centromeres. Sequence, structure, methylation pattern, local sequence identity, and haplotype frequency of 2,156 completely assembled centromeres from CHM13^8^, CHM1^9^, and 65 diverse human genomes^12^. The α-satellite HORs are colored by the number of α-satellite monomers within them, and the orientation of the active α-satellite HOR arrays are indicated by an arrow. The site of the putative kinetochore, marked by the centromere dip region (CDR), is shown with a black bar. The local sequence identity across each centromeric region is indicated with a 1D heat map, and mobile element insertions (MEIs) within α-satellite HOR arrays are indicated with colored triangles. Haplotype frequencies are shown as pie charts, with the continental group indicated. The most common haplotype for each chromosome is marked with an asterisk. All 226 new major haplotypes are shown in **Supplementary Fig. 3**. Zoomed-in views of each haplotype and their pie chart are also shown in **Supplementary Figs. 4-6**.

### Global diversity among human centromeres

Using these complete centromere assemblies, we then sought to determine the extent of variation among human centromeres. We first investigated the variation in the length of α-satellite HOR arrays. Human centromeres are typically composed of one or more arrays of tandemly repeated α-satellite HORs, with one array serving as the site of the kinetochore. We, therefore, distinguished between kinetochore-forming and non-kinetochore-forming arrays, marked by the presence or absence of the CDR, respectively, and determined the length of each of these in our dataset (**Supplementary Figs. 7,8**, **Supplementary Table 6, Supplementary Note 2**). We found that kinetochore-forming arrays are, on average, 2.5 Mbp long, with the smallest array only 239 kbp in length (on chromosome 21) and the largest 7.5 Mbp in length (on chromosome 18; **Fig. 2a**, **Supplementary Fig. 7a**, **Supplementary Table 6**). Two chromosomes (chromosomes 1 and 18) have significantly larger kinetochore-forming arrays than the other chromosomes, with a mean length of 4.3 and 4.2 Mbp, respectively, while seven chromosomes (chromosomes 3, 4, 13, 15, 16, 21, and Y) have significantly smaller kinetochore-forming arrays, with a mean length of 1.7, 1.5, 1.6, 1.5, 1.8, 1.4, and 0.7 Mbp, respectively (*p* < 0.05, two-tailed Wilcoxson rank-sum test). The greatest variation in length occurs on chromosome 6, with a 21.5-fold difference between the smallest and largest, while the greatest absolute difference in length occurs on chromosome 18, with a 6.3-Mbp difference (**Fig. 2a**, **Supplementary Fig. 7a**, **Supplementary Table 6**). Comparison of the lengths of kinetochore-forming arrays between haplotypes of the same individual reveals that they are, on average, only 1.5-fold different in length, with <1% of arrays more than 4-fold larger than their counterpart (**Supplementary Fig. 9**, **Supplementary Table 6**).

**Figure 2.**
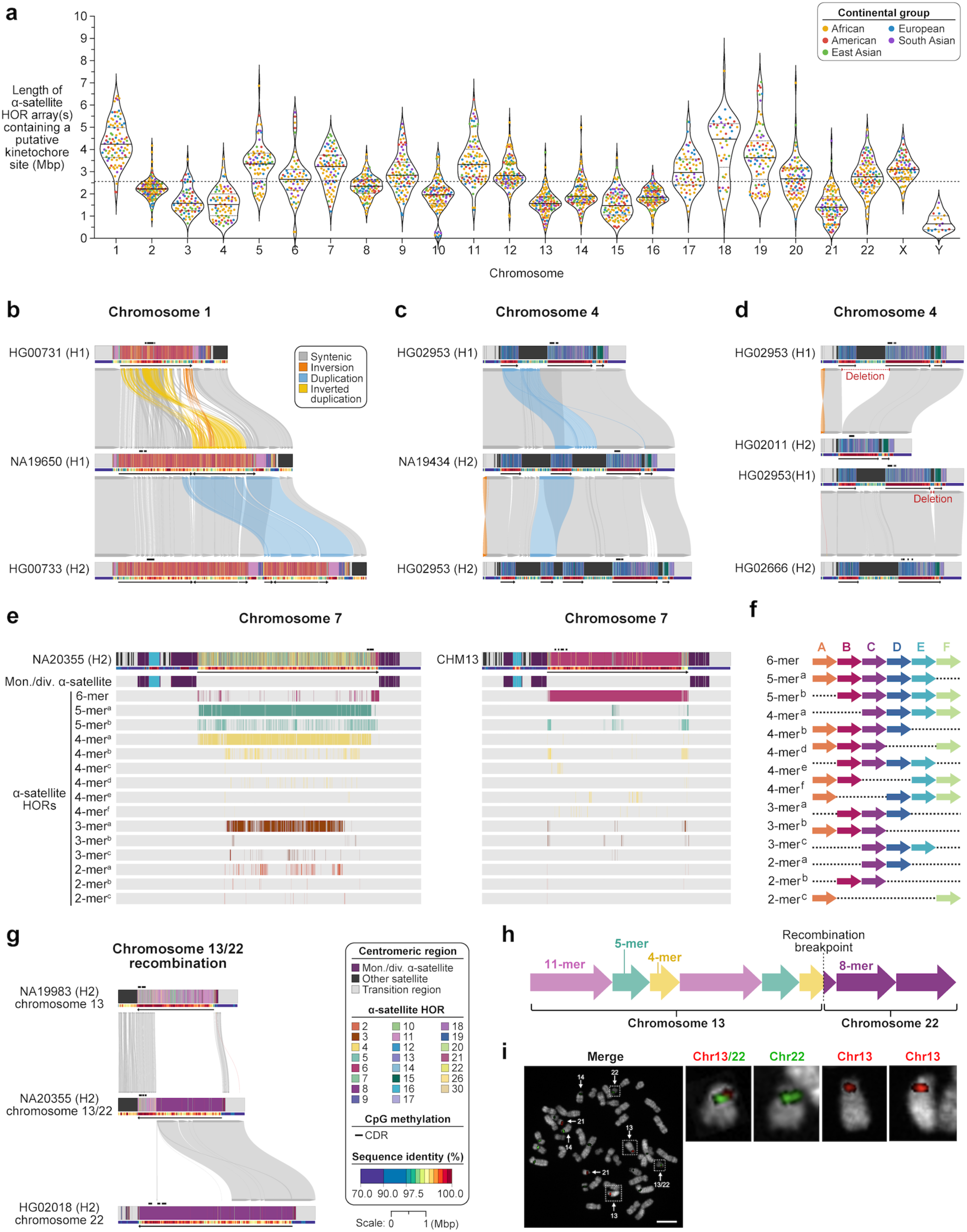
Complex sequence and structural variation among human centromeres. **a)** Variation in the length of α-satellite HOR arrays containing a putative kinetochore site, defined by the presence of a CDR, among 2,110 diverse human centromeres. For each chromosome, the mean is shown as a solid line, and first and third quartiles are shown as dotted lines. The mean across all chromosomes is shown as a dashed line. **b-d)** Examples of large-scale (hundreds of kbp) structural variation among centromeres, including inversions, duplications, inverted duplications, and deletions, within the **b)** chromosome 1 *D1Z7* α-satellite HOR array and **c,d)** chromosome 4 *D4Z1* α-satellite HOR arrays. **e)** Differences in α-satellite HOR array organization within the chromosome 7 *D7Z1* array between the NA19238 (left) and CHM13 (right) genomes. **f)** Structure of α-satellite HOR variants present in the NA19238 (H2) and CHM13 chromosome 7 centromeres. All of them derive from an ancestral 6-mer α-satellite HOR. **g)** A recombination event between the chromosome 13 *D13Z1* α-satellite HOR array and chromosome 22 *D22Z1* α-satellite HOR arrays gives rise to a chromosome 13/22 hybrid α-satellite HOR array. We note that the top and bottom centromeres are not genetically related to the hybrid centromere. **h)** Schematic showing the breakpoint between the *D13Z1* and *D22Z1* α-satellite HOR arrays. The recombination event creates a new α-satellite HOR variant 5 monomers in length. **i)** Fluorescent *in situ* hybridization (FISH) on an NA20355 metaphase chromosome spread showing the recombined chromosome 13/22 chromosome. Fluorescent probes specific to chromosome 13/21 and 14/22 α-satellite DNA colocalize on the recombined chromosome 13/22 chromosome in 100% of cells (*n*=20). Scale bar, 10 μm. Insets are magnified 6.5×.

Across continental groups, we find that nine chromosomes (chromosomes 3, 7, 9, 16, 17, 19, 21, 22, and X) have significantly different kinetochore-forming α-satellite HOR array lengths (**Supplementary Figs. 7b,8**, **Supplementary Table 6**). For chromosome 21, for example, we find that individuals with African ancestry have α-satellite HOR arrays that are 353 kbp larger, on average, than those with non-African ancestry. However, the opposite is true for chromosomes 3, 7, 9, 16, 17, 19, 22, and X, wherein individuals with African ancestry have arrays that are 281, 522, 348, 180, 476, 654, 491, and 594 kbp smaller, on average, than those with non-African ancestry, respectively (**Supplementary Figs. 7b,8**, **Supplementary Table 6**). For the remaining chromosomes, the kinetochore-forming α-satellite HOR array lengths are similar across continental groups, showing no significant correlation with genetic ancestry (**Supplementary Figs. 7b,8**, **Supplementary Table 6**).

In addition to variation in α-satellite HOR array length, we observed both large- and small-scale variation impacting the sequence, structure, and organization of α-satellite HOR arrays (**Fig. 2b-h**, **Supplementary Fig. 10**). We identified ten chromosomes with large duplications, deletions, or inversions (hundreds of kilobase pairs long) that alter the number of α-satellite HOR arrays and/or the organization within them (chromosomes 1, 4, 5, 7, 10, 12, 13, 17, 19, and 20; **Fig. 2b-d**, **Supplementary Fig. 10**). This is perhaps best exemplified by chromosome 1, where we found that a quarter of the individuals in our dataset have a large-scale inversion in the core of the *D1Z7* α-satellite HOR array spanning 637 kbp to 2.47 Mbp (like CHM13^8^ and CHM1^9^); however, the majority (68%) have a single, directly-oriented *D1Z7* α-satellite HOR array, and another 7% have a duplication or triplication of a portion of the *D1Z7* α-satellite HOR array, resulting in two or three distinct arrays spanning 631 kbp to 2.81 Mbp each (**Fig. 2b**, **Supplementary Fig. 10**). We also find that chromosomes 3 and 4 have multiple α-satellite HOR arrays that are interrupted by stretches of human satellite 1A (HSat1A), as previously reported^7,8^. Whereas 97% of chromosome 3 centromeres in our dataset have only two *D3Z1* α-satellite HOR arrays, we find that a much smaller proportion of individuals (56%) have two *D4Z1* α-satellite HOR arrays on chromosome 4, while the rest have three, four, or five arrays (23%, 18%, and 3%, respectively), mainly among those with African ancestry (**Fig. 2c,d**). For chromosomes 5, 7, 12, 13, 17, 19, and 20, almost all individuals in our dataset have a single active α-satellite HOR array; however, at least one individual has a duplication of the active α-satellite HOR array and flanking divergent α-satellite HORs, resulting in two or more distinct α-satellite HOR arrays (**Supplementary Fig. 10**). Finally, chromosomes 10 and 17 often have small active α-satellite HOR arrays that flank the larger α-satellite HOR array, occurring in 100% and 64% of individuals, respectively (**Supplementary Fig. 10**).

Finally, we uncovered variation in the sequence composition of the α-satellite HORs themselves. Comparison of the α-satellite HORs in these 2,110 centromeres to those in the CHM13 and CHM1 genomes revealed 1,870 new α-satellite HOR variants never seen before (**Supplementary Table 4**). We confirmed that 97% of these new α-satellite HOR variants exist in at least one other genome in our dataset or in the broader dataset of 5,747 complete centromeres assembled by the HPRC (**Supplementary Table 5**), underscoring their biological relevance. Many of these new α-satellite HOR variants resulted in completely new α-satellite HOR array organizations relative to CHM13 and CHM1. For 12 chromosomes (chromosomes 4, 6-8, 10, 13, 14, 16-18, 20 and 21), we discovered 26 distinct haplotypes that had markedly different α-satellite HOR compositions compared to CHM13 and CHM1 (**Supplementary Figs. 11-22**). The most pronounced differences occurred on chromosomes 7, 16, and 21. For chromosome 7, we identified three individuals with African ancestry who harbor a centromere with rare α-satellite HOR variants that are up to 251 times more abundant than in the CHM13 or CHM1 centromeres, completely changing the composition and organization of the *D7Z1* α-satellite HOR array (**Fig. 2e,f**, **Supplementary Fig. 13**, **Supplementary Tables 7,8**). Similarly, for chromosome 16, we identified seven individuals with African ancestry who had three or four novel α-satellite HOR variants as well as one 8-mer α-satellite HOR variant that was 18 to 1,378 times more abundant than in the CHM13 and CHM1 genomes (**Supplementary Fig. 18**, **Supplementary Tables 7,8**). In contrast, we identified five individuals with diverse ancestry (one African, one American, one East Asian, and two South Asian) that harbor distinctly different chromosome 21 centromeres (**Supplementary Fig. 22**). These individuals contain up to seven new α-satellite HOR variants as well as a high abundance of 5-mer or 7-mer α-satellite HORs that are nearly absent from the CHM13 and CHM1 centromeres (**Supplementary Tables 7,8**). The presence of these variants within individuals of diverse ancestries suggest they are likely ancient α-satellite HORs, deriving from an ancestral 11-mer α-satellite HOR that resides at the periphery of these new haplotypes and is typically the most common α-satellite HOR variant among chromosome 21 centromeres (**Supplementary Fig. 22b**).

Although rare, we note the discovery of a recombination event between two centromeres from acrocentric chromosomes (chromosomes 13 and 22), creating a hybrid centromere in an individual (NA20355). Previous studies have shown that acrocentric chromosomes 13, 14, 15, 21, and 22 are prone to homologous recombination, resulting in an exchange of p arms or a fusion of the q arms, as with Robertsonian translocations^23,24^. While these recombination events have been reported to occur at the pseudo-homologous regions (PHRs) on the p arms^23–26^, they have never been shown to occur within the centromeric α-satellite itself. Here, we identified an individual that carries a recombined chromosome 13/22 centromere, with the breakpoint within the α-satellite HOR array, the p arm from chromosome 13, and the q arm from chromosome 22 (**Fig. 2g,h**). We validated the breakpoint with both PacBio HiFi and ONT long-read sequencing data (**Supplementary Fig. 23a-c**) as well as with fluorescent *in situ* hybridization (FISH), which showed colocalization of chromosomes 13 and 22 alphoid DNA on the same chromosome in NA20355 (**Fig. 2i**). Close examination of the breakpoint sequence reveals 6 base pairs of homology (AAAAGA) between the α-satellite monomers on chromosome 13 and 22 (**Supplementary Fig. 23d**), indicating that the recombination event may have been mediated by microhomology, as previously suggested^27^. To estimate the frequency of this event in the greater population, we assessed 633 additional chromosome 13 and 22 centromeres assembled from 351 individuals (702 haplotypes) by the HPRC and found 2 such examples of recombined chromosome 13/22 centromeres (from HG00235 and HG00639; **Supplementary Fig. 24**), bringing the estimated rate of occurrence to 1 in 283 in our dataset (*n*=415). The two other individuals (HG00235 and HG00639) have a recombination breakpoint at the exact same 6-bp sequence as in NA20355, indicating a conserved mechanism for the formation of a hybrid chromosome 13/22 centromere. Although all three individuals have different genetic ancestries (NA20355, African; HG00235, European; HG00639, American), they share the same centromere haplotype, suggesting that this structure evolved relatively recently and may predispose individuals to recombination between chromosomes 13 and 22. Biochemical and cell-based experiments that test the molecular basis for the formation of a hybrid chromosome 13/22 centromere and its functional consequences will need to be performed to confirm this hypothesis.

### Epigenetic variation among diverse human centromeres

Functional centromeres are epigenetically defined by two key features: nucleosomes containing the histone H3 variant CENP-A and underlying domains of hypomethylated DNA, known as CDRs^8,22^. Together, these marks establish a unique chromatin foundation for the assembly of the kinetochore, a multi-protein complex that binds spindle microtubules to ensure faithful chromosome segregation during cell division. While all functional centromeres recruit a kinetochore, we lack a clear understanding of the variation in kinetochore location, distribution, and organization among human centromeres, particularly across diverse haplotype structures and genetic ancestries. Using the complete centromere assemblies we generated, we sought to uncover how the underlying sequence and structure within centromeres affects the position and distribution of the kinetochore and how this stratifies by continental groups. To do this, we first determined the methylation profile of each centromere by aligning native ONT long-read sequencing data from the same source genome to each assembly and identifying the location of the CDR along the centromere^28^. Then, we performed CENP-A DiMeLo-seq followed by ONT long-read sequencing on 10% of the cell lines and mapped them back to the respective genome assembly. By normalizing the CENP-A DiMeLo-seq signal with IgG signal, we found that every CDR was enriched with CENP-A (**Supplementary Fig. 25**). Finally, we adapted and optimized CENP-A CUT&RUN for ONT long-read sequencing^29^ and mapped the resulting reads to the relevant genome assembly, confirming the location of CENP-A within the CDR (**Supplementary Fig. 25**; **Methods**).

Using these validated kinetochore sites, we characterized their distribution, length, and position along each centromere (**Fig. 3**). We found that 94% of centromeres contain one kinetochore site, while 6% (119/2110) contain two sites, and <<1% (1/2110) contain three sites, separated by >150 kbp of sequence (**Fig. 3a**, **Supplementary Fig. 26**, **Supplementary Table 9**). Centromeres with multiple kinetochore sites tend to have them clustered within 390 kbp of each other (**Fig. 3b**, **Supplementary Table 9**); however, rarely, they are separated by multiple megabases of sequence, as with the chromosome 11 centromere in HG00513, where the two kinetochores differ in position by 2.9 Mbp (**Supplementary Fig. 27a,b**). Centromeres with multiple kinetochore sites are also meiotically stable, as the chromosome 11 centromere with two distant kinetochores is also observed in the child, HG00514 (**Supplementary Fig. 27c,d**), indicating that this epigenetic architecture is transmissible to the next generation.

**Figure 3.**
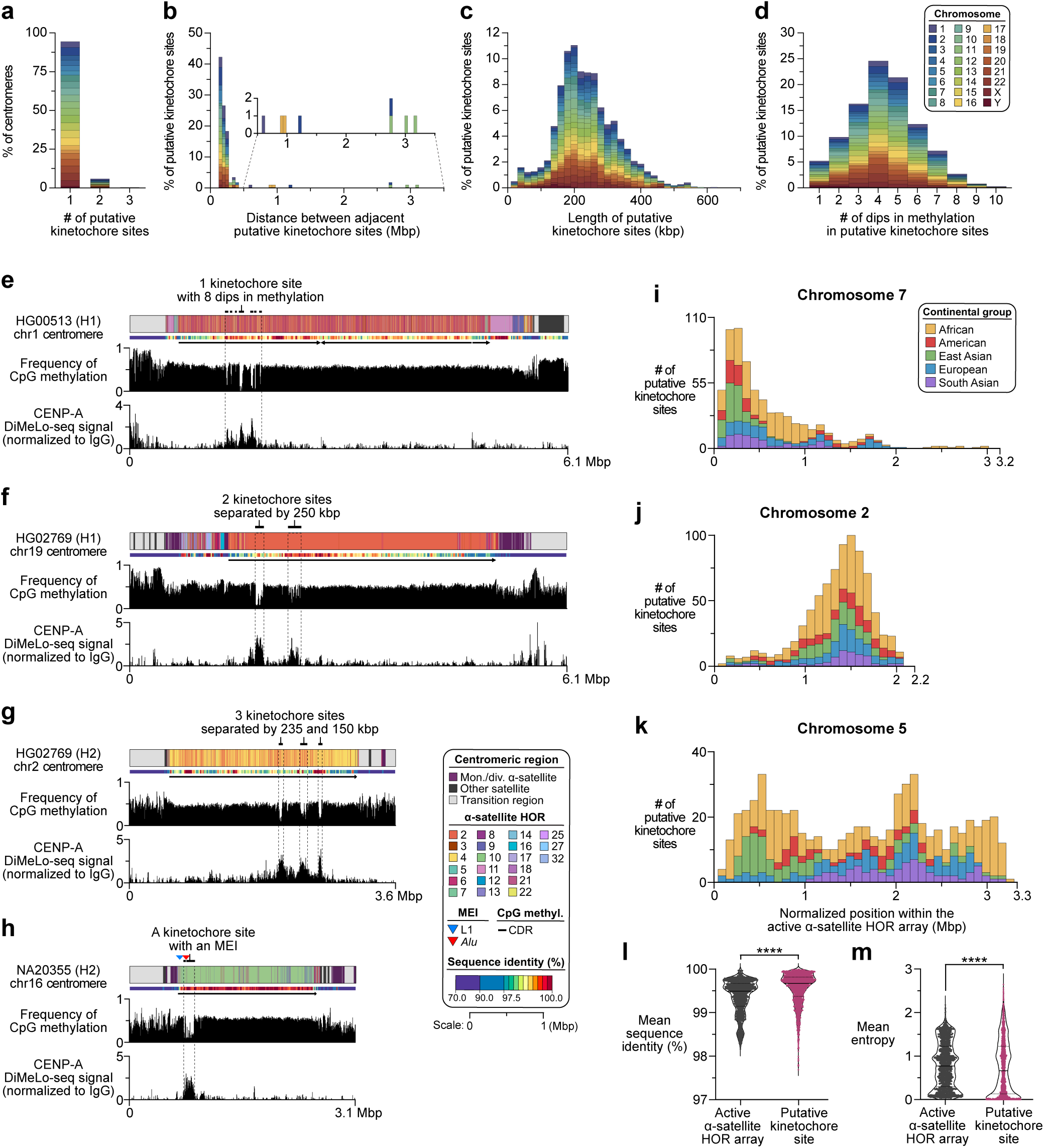
Variation in the length, distribution, and position of putative kinetochore sites among human centromeres. **a)** Number of putative kinetochore sites within each chromosome, marked by hypomethylated DNA and separated by >150 kbp of sequence. **b)** Distance between putative kinetochore sites within a single centromere. Approximately 7% of centromeres have more than one putative kinetochore site separated by >150 kbp of sequence. **c)** Length of putative kinetochore sites, marked by stretches of hypomethylated DNA. **d)** Number of dips in methylation among putative kinetochore sites. **e-g)** Examples of centromeres with **e)** a single kinetochore site, **f)** two kinetochore sites separated by 250 kbp of sequence and residing within different sequence contexts, and **g)** three kinetochore sites separated by 235 and 150 kbp of sequence. **h)** Example of an *Alu* insertion within the kinetochore site of the NA20355 chromosome 16 *D16Z2* α-satellite HOR array. The *Alu* insertion changes the methylation profile of the CDR and breaks a 10-mer α-satellite HOR into a 2- and 8-mer α-satellite HOR. CENP-A is still enriched over the *Alu* element relative to IgG, as determined with DiMeLo-seq (**Supplementary Fig. 28**). **i-k)** Position of the CDR across active α-satellite HOR arrays within the centromeres from chromosomes **i)** 7, **j)** 2, and **k)** 5. On chromosome 7, most CDRs are located on the p arm-proximal side of the array; on chromosome 2, most CDRs are located on the q arm-proximal side of the array; for chromosome 5, CDRs are located on different regions of the α-satellite HOR array depending on their haplotype structure and genetic ancestry. Notably, no CDRs from individuals of South Asian ancestry are located on the p arm-proximal side of the array, indicating that the underlying sequence and structure likely influences the position of the CDR. The distribution of CDRs for each chromosome is shown in **Supplementary Fig. 29**. **l,m)** Comparison of the mean **l)** sequence identity and **m)** entropy between sequences comprising the active α-satellite HOR array vs. those within the CDR. CDR-harboring sequences have higher sequence identity and lower entropy. ****, p<0.0001.

Focusing on individual kinetochore sites, we find that they are typically 240 kbp long on average, with the smallest single site being 90 kbp long and the largest 630 kbp long (**Fig. 3c**, **Supplementary Table 9**). Most kinetochore sites contain multiple dips in methylation that alternate with methylated DNA, with an average of 4.3 dips and a maximum of 10 dips (**Fig. 3d,e**). While the underlying DNA sequence does not often correlate with DNA methylation patterns (**Fig. 3f,g**), we find that, in a subset of centromeres, a change in methylation frequency is due to the insertion of a mobile element within the kinetochore site (**Fig. 3h**). In total, we identify 636 centromeres (30%) in which a mobile element (L1 or *Alu*) is present in the α-satellite HOR array (**Fig. 1**, **Supplementary Fig. 28a**, **Supplementary Table 10**). In 33 of these centromeres (∼5%), the mobile element is present in the putative kinetochore site, altering the DNA methylation profile of the CDR and breaking it into multiple dips in methylation (**Fig. 3h**, **Supplementary Fig. 28b-d**). While MEIs within kinetochore sites are rare, they occur frequently within the chromosome 16 centromere, with 14% (16/112) of centromeres harboring an MEI within the putative kinetochore site, which is 17 times more frequent than in the other chromosomes (**Supplementary Fig. 28a-d**, **Supplementary Table 10**). This is notable, as chromosome 16 is known to have the highest rate of nondisjunction during meiosis and is the most frequently gained or lost chromosome in aneuploid embryos^30,31^. Previous studies suggest that errors in kinetochore-microtubule attachments can lead to increased rates of chromosome missegregation^32^, and MEIs within the kinetochore site may disrupt the 3D chromatin organization required for proper spindle microtubule attachment. Additional studies that test the functional consequences of MEIs within kinetochore sites, such as those found on chromosome 16, will need to be performed to confirm their contribution to increased chromosome missegregation rates.

Perhaps the most striking discovery, however, is that the kinetochore position is not random and is, instead, closely associated with the underlying sequence and structure of the centromere. Indeed, we find that the position of the kinetochore tends to fall into one of three categories: p arm-proximal, q arm-proximal, or distributed in different locations throughout the α-satellite HOR array depending on the haplotype. For 11 out of 24 chromosomes (chromosomes 1, 3, 7, 9, 13-16, 21, 22, and Y), nearly all CDRs are located on the p arm-proximal side of the array (**Fig. 3i, Supplementary Fig. 29**). For one-twelfth of chromosomes (chromosomes 2 and 20), almost all CDRs are located on the q arm-proximal side of the array (**Fig. 3j, Supplementary Fig. 29**). For the remaining chromosomes (chromosomes 4-6, 8, 10-12, 17-19, and X), CDRs are located on different regions of the α-satellite HOR array depending on their haplotype structure (**Fig. 3k, Supplementary Fig. 29**). Importantly, centromeres from individuals with different genetic ancestries have different CDR distributions for these chromosomes. This is perhaps best demonstrated with chromosome 5 (**Fig. 3k**), where individuals with African, American, and East Asian ancestry have CDRs distributed along the length of the α-satellite HOR array, but those with European and South Asian ancestry rarely have a CDR located toward the p arm.

This bias in CDR position suggests that there may be particular sequences or structures that are more likely to serve as the site of the kinetochore than others within the centromeric α-satellite HOR array. To test this hypothesis, we calculated the mean sequence identity within the CDR and across the α-satellite HOR array and found that the CDR resides on sequences that are more highly identical to each other than the rest of the array (mean sequence identity = 99.55% vs. 99.36%, respectively; p<0.0001, two-tailed Mann-Whitney test; **Fig. 3l**, **Supplementary Fig. 30a, Supplementary Table 11**). Additionally, the CDR resides within regions that are more homogenous in sequence and structure and, therefore, have lower entropy (mean entropy = 0.74 vs. 0.80, respectively; p<0.0001, two-tailed Mann-Whitney test; **Fig. 3m**, **Supplementary Fig. 30b, Supplementary Table 11**). This is the case for all centromeres except for those from chromosomes 8 and 13, which have higher entropy among sequences underlying the kinetochore, as previously reported^6^ (**Fig. 3m**, **Supplementary Fig. 30b, Supplementary Table 11**). The finding that the kinetochore typically resides on highly identical, homogenous stretches of α-satellite HORs is consistent with a model first put forth by Alexandrov and colleagues that suggests that centromeric proteins and DNA act in concert, recombining the sequences underneath the kinetochore via a kinetochore-associated recombination machine (KARM) and ultimately leading to increased homogeneity among these sequences^33,34^.

### Ancient and modern evolutionary events shape centromere structure

Our findings reveal that centromeres vary considerably in size, sequence, structure, and epigenetic landscapes among diverse humans, with 226 major centromere haplotypes, 1,870 α-satellite HOR variants, and distinct epigenetic landscapes that correlate with centromere structure (**Figs. 1-3**). However, when these haplotypes first emerged and how rapidly they are evolving remains unclear. To investigate the nature of evolutionary change among centromeres, we leveraged the suppressed meiotic recombination around these regions and identified sequences in the pericentromeres that are in linkage disequilibrium (LD) and associated with particular centromere haplogroups^9^ (**Supplementary Fig. 31**). We then built maximum-likelihood phylogenetic trees for each centromere, using chimpanzee sequences as outgroups and assuming a chimpanzee-human divergence time of 6.2 million years^35^ (**Supplementary Fig. 31**). The phylogenetic trees we built for both of the p and the q arms are remarkably similar, supporting the notion of limited-to-no recombination across centromeres during meiosis (**Supplementary Figs. 32-59**). Additionally, they reveal that centromeres with similar structures and epigenetic landscapes cluster within monophyletic clades, indicating they have a shared evolutionary history.

Phylogenetic analysis reveals evidence that both ancient and modern evolutionary events have shaped human centromere architecture. First, we identify three chromosomes that have centromere haplotypes separated by >1 million years of evolution (chromosomes 10, 12, and 21; **Fig. 4a**, **Supplementary Figs. 41, 43, 52**), as confirmed with the HPRC centromere assemblies (**Supplementary Figs. 57-59**). For chromosome 12, the minor haplotype is present only within a subset of individuals with African ancestry (**Supplementary Figs. 43**, **58**), suggesting that this haplotype arose in Africa and was likely lost during population bottlenecks following the out-of-Africa migration. In contrast, for chromosomes 10 and 21, we find minor haplotypes present within individuals of diverse genetic ancestry [chromosome 10: American, European, East Asian, and African (**Supplementary Fig. 41**, **57**); chromosome 21: East Asian and South Asian (**Fig. 4a, Supplementary Fig. 52**, **59**)], suggesting that these haplotypes possibly emerged as a result of introgression with archaic hominins. To determine if the minor haplotypes from chromosomes 10 and 21 have evidence of introgression with archaic hominins, we applied an alignment-free, *k*-mer-based approach^36^ that leverages high-coverage Neanderthal and Denisovan whole-genome sequencing datasets^37–40^ to identify archaic hominin (ARC)-specific *k*-mers that are not present in modern-day Africans (**Methods**). Using this method, we found that the minor haplotypes from chromosomes 10 and 21 have a significant enrichment of ARC-specific *k*-mers, suggesting the presence of introgressed DNA from Neanderthals and Denisovans in these sets of centromeres (**Fig. 4b**, **Supplementary Figs. 60, 61**). In contrast, more evolutionarily young centromeres from chromosome 21 (**Fig. 4c**) and the minor haplotypes from chromosome 12 (**Supplementary Fig. 62**) do not show this enrichment, indicating that introgression is specific to only a subset of centromere haplotypes.

**Figure 4.**
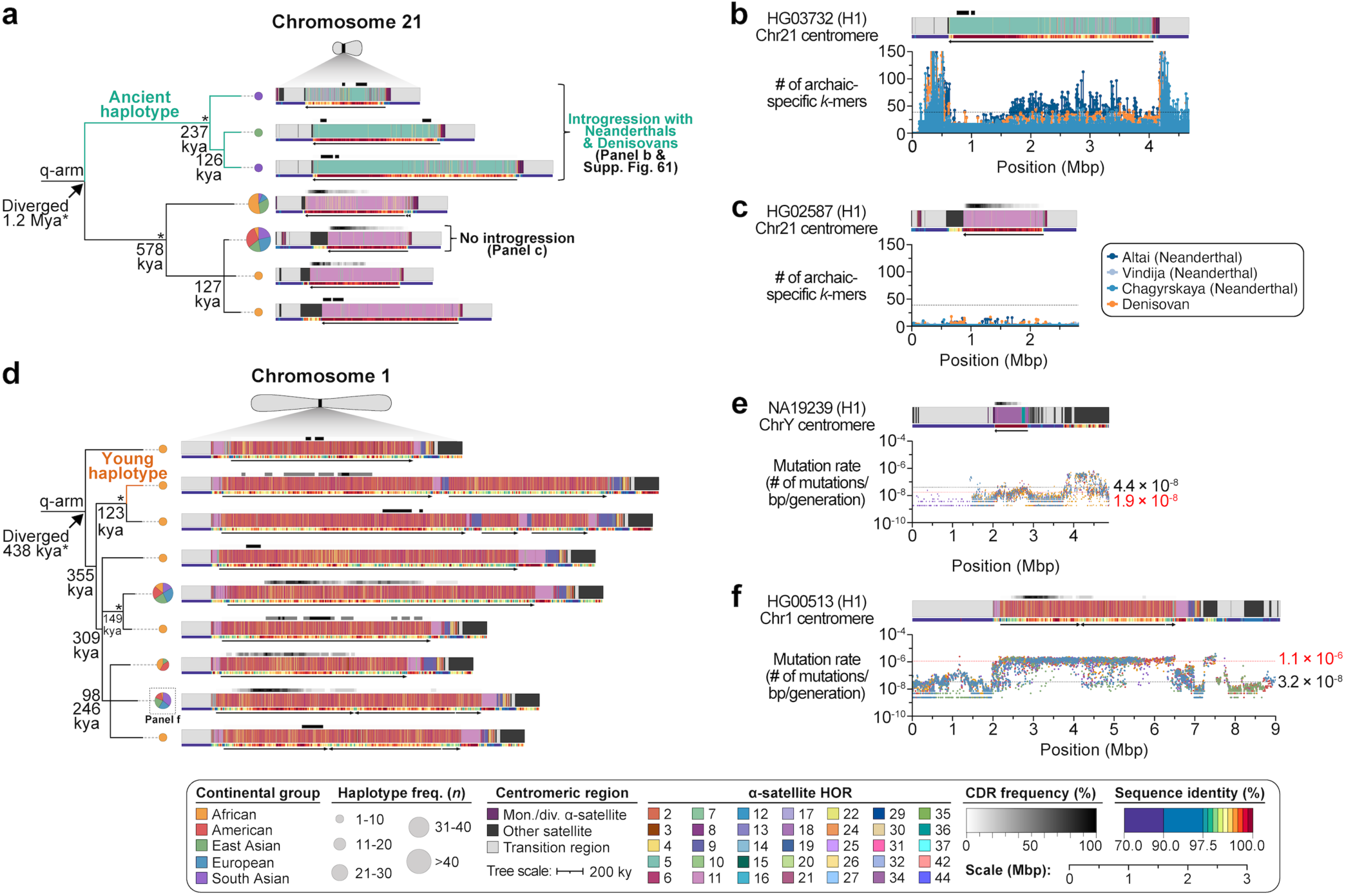
Evidence of introgression from archaic hominins within centromeres. **a)** Maximum-likelihood phylogenetic tree depicting the q-arm topology for chromosome 21 centromere haplogroups along with estimated divergence times and haplotype frequencies. Asterisks indicate nodes with 100% bootstrap support, and nodes with 90-99% bootstrap support are indicated numerically. Nodes without an asterisk or number have bootstrap support <90%. An ancient haplotype with a completely different *D21Z1* α-satellite HOR array organization (composed mainly of 5-mer α-satellite HORs) is observed, with an estimated divergence time of 1.2 Mya. This haplotype is only found within non-African individuals, suggesting it may have arisen due to introgression with archaic hominins. **b,c)** Distribution of archaic-specific DNA *k*-mers across 20 kbp windows of the **b)** HG03732 (H1) or **c)** HG03248 (H1) chromosome 21 centromeric regions using high-quality archaic short-read data from four genomes: Altai (Neanderthal), Vindija (Neanderthal), Chagryskaya (Neanderthal), and Denisovan. Black dashed lines indicate the genome-wide *k*-mer count cutoff, 42, which corresponds to the genome-wide archaic introgression fraction^36^. The ancient haplotype enriched with 5-mer α-satellite HORs has an abundance of archaic DNA k-mers, suggesting it emerged due to introgression with Neanderthals and Denisovans, while the most frequent haplotype enriched with 11-mer α-satellite HORs does not. **d)** Maximum-likelihood phylogenetic tree depicting the q-arm topology for chromosome 1 centromere haplogroups along with estimated divergence times and haplotype frequencies. A young haplotype with a large, multi-megabase duplication of the α-satellite HOR array is observed and is estimated to have emerged in the last 10,000 years (or 500 generations). **e,f)** Plots showing the estimated mutation rate across the chromosome **e)** Y and **f)** 1 centromeric regions. Individual data points from 10-kbp pairwise sequence alignments are shown. Mean mutation rates across the active α-satellite HOR array and flanking non-satellite sequences are shown as dotted red and black lines, respectively. There is a 58-fold difference in mutation rate of the α-satellite HOR arrays between the chromosome Y and 1 centromeres.

In addition to ancient evolutionary events, we also find that modern evolutionary changes have shaped the structure of human centromeres. In fact, we find that nearly all chromosomes have centromere haplotypes that have emerged in the last 10,000 years, such as on chromosome 1, where a 2-4 Mbp duplication of the *D1Z7* active α-satellite HOR array is estimated to have occurred in a subset of African individuals over the course of the last ∼500 generations (**Fig. 4d**, **Supplementary Figs. 10a, 32**), as confirmed with HPRC centromere assemblies (**Supplementary Fig. 56**). Even more recently, a ∼1-Mbp duplication of the *D4Z1* active α-satellite HOR array on chromosome 4 is estimated to have occurred within the last 1,000 years (or 50 generations) among a subset of African individuals (**Fig. 2c**, **Supplementary Fig. 35**), indicating that centromeres are continually mutating and evolving.

### Elevated mutation rates within the centromere and its kinetochore site

The variable evolutionary history among centromeres (described above) suggests that centromeres from different chromosomes may have different mutation rates that alter their sequence and structure. To test this hypothesis, we refined our previous mutation rate estimation method^9^ to use a pan-centromere alignment framework with clade-specific reference sequences in order to account for lineage-level differences (**Supplementary Fig. 31; Methods**). Using this framework, we estimate the mean mutation rate among centromeric α-satellite HOR arrays to be 4.03×10^-7^ mutations per base pair per generation (Ms/bp/G; 95% CI: 4.01-4.05 ×10^-7^ Ms/bp/G), which is 16.6-fold higher than the unique portions flanking the centromeres^9,15^ (**Supplementary Table 12**, **Supplementary Figs. 63-73**). The lowest mutation rate occurs in the chromosome Y centromere, with an average of 1.9×10^-8^ Ms/bp/G across the *DYZ3* α-satellite HOR array (**Fig. 4e**, **Supplementary Figs. 73c,74**), while the highest occurs in the chromosome 1 centromere, with an average of 1.1×10^-6^ Ms/bp/G across the *D1Z7* α-satellite HOR array (**Fig. 4f**, **Supplementary Figs. 63c, 74**), a 57.9-fold difference in mutation rate among centromeres. Even within the same chromosome, we find that centromere haplogroups exhibit diverse mutation rates, with the smallest difference occurring on chromosome 18 (1.03-fold; **Supplementary Figs. 70i,j**, **75**, **Supplementary Table 12**) and the greatest difference occurring on chromosome 12 (5.02-fold; **Supplementary Figs. 68e-j**, **75**, **Supplementary Table 12**). Despite these differences, we find that the α-satellite HOR mutation rates are relatively consistent across individuals with different genetic ancestries, ranging from an average of 4.0×10^-7^ Ms/bp/G in Americans to 4.1×10^-7^ Ms/bp/G in South Asians (**Supplementary Fig. 74b**).

To validate the centromere mutation rates across multiple generations, we leveraged 28 near-complete genome assemblies from a four-generation pedigree (CEPH 1463) that had been recently generated^15^ (**Supplementary Fig. 76a**). These genome assemblies already had 288 centromeres completely and accurately resolved^15^; however, we reasoned that we could repair the assembly errors in many of the remaining centromeres using AssemblyRepairer, taking advantage of the complimentary error profiles of genome assemblies generated with Verkko^16^ and hifiasm^17^. Applying AssemblyRepairer to the Verkko genome assemblies for generations 1-4 (G1-G4) allowed us to resolve an additional 195 centromeres, which we subsequently validated (**Supplementary Note 1**, **Supplementary Tables 13,14**), bringing the total number of centromeres to 483, a 67.7% increase (G1: 134 centromeres; G2: 70 centromeres; G3: 215 centromeres, G4: 64 centromeres; **Supplementary Fig. 76b**). Among these 483 centromeres, 244 of them were transmitted from one generation to the next (**Supplementary Fig. 76c, Supplementary Table 15**), allowing us to identify *de novo* SNVs and SVs that occur within the centromeric regions across generations.

Using these assemblies, we identified 206 *de novo* SNVs within centromeric regions across four generations (**Supplementary Table 16**), which we validated with raw PacBio HiFi and ONT reads as well as with transmission to the next generation when possible. We found that the mutation rate varies depending on the underlying sequence (**Fig. 5a**, **Supplementary Table 17**). The lowest mutation rate occurs in the unique sequences in the pericentromere (7.1×10^-8^ Ms/bp/G), while the monomeric/divergent α-satellite HORs have a slightly higher mutation rate (1.1×10^-7^ Ms/bp/G), and the active α-satellite HORs have a mutation rate more than twice that of the monomeric/divergent α-satellite HORs (2.6×10^-7^ Ms/bp/G). Within the active α-satellite HOR array itself, we find that the CDR has the highest mutation rate (5.1×10^-7^ Ms/bp/G), which is approximately twofold greater than the surrounding α-satellite HORs in the rest of the array (**Fig. 5a**, **Supplementary Table 17**). Importantly, these mutation rates are similar to our earlier pan-centromere-based estimates (**Fig. 4c-f**), which averaged 4.03×10^-7^ Ms/bp/G.

**Figure 5.**
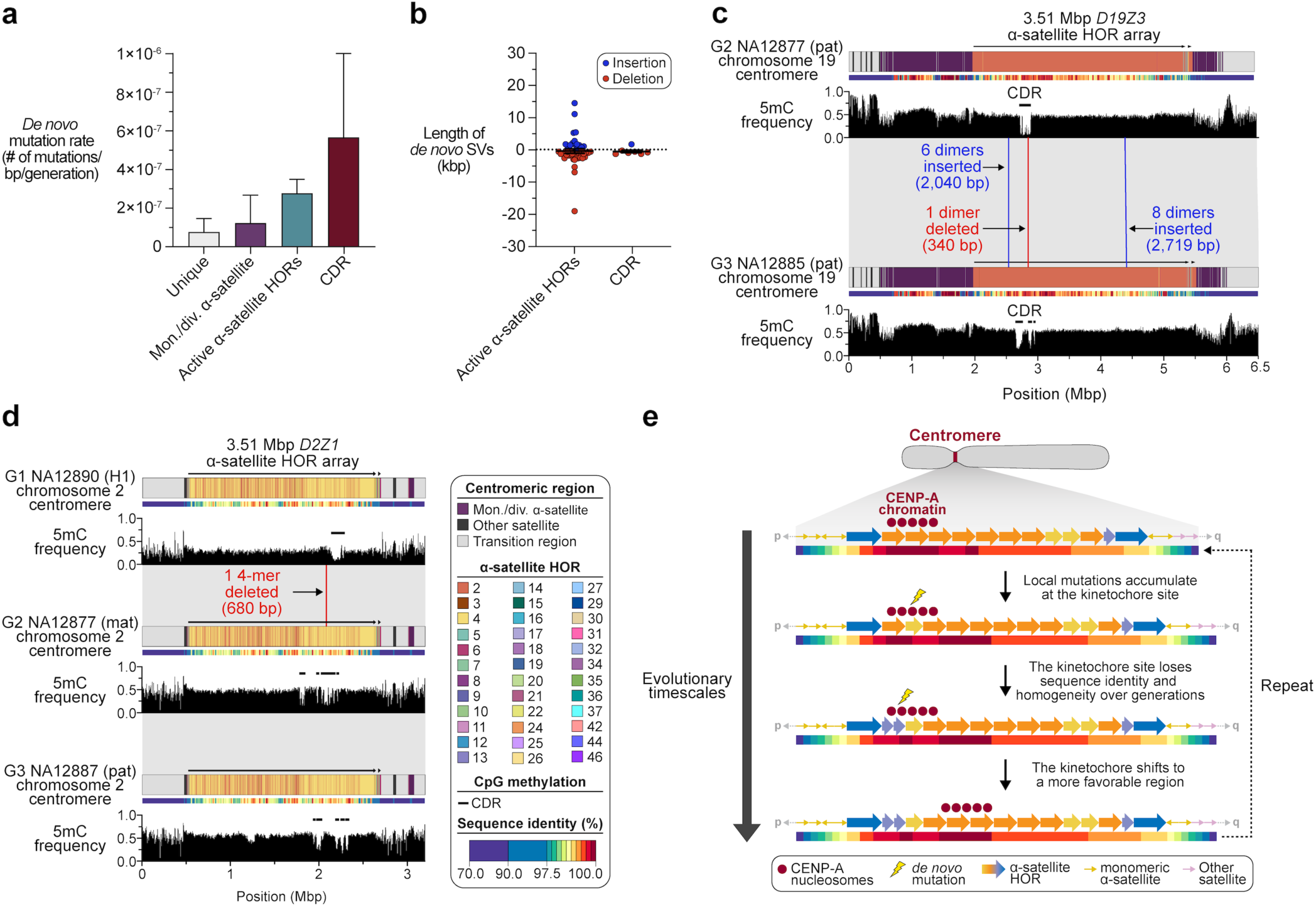
Accelerated mutation rates within the putative kinetochore site results in changes in its position, distribution, and sequence composition across generations. **a)** Estimated *de novo* mutation rates for different sequences within centromeric regions across three generations of a pedigree (CEPH 1463). The active α-satellite HORs and CDR are mutating 3.6- and 7.1-fold faster than the unique sequences in the pericentromere, respectively. **b)** Length of *de novo* SVs within the active α-satellite HORs and CDR. *n*=43 and *n*=7 for the α-satellite HOR array and CDR, respectively. **c,d)** Examples of SVs occurring within the active α-satellite HOR array and CDR in the **c)** chromosome 19 and **d)** chromosome 2 centromeres across generations. Mutations occurring within or near the CDR often result in a change in the position, distribution, and/or length of the CDR from one generation to the next. **e)** Model of centromere mutation and evolution, showing that the CDR is often located within the most identical and homogenous sequences of the centromere and is prone to both SNVs and SVs that alter the genetic and epigenetic landscape of the centromere over time.

In addition to *de novo* SNVs, we also identified 43 *de novo* SVs (defined as an insertion or deletion >50 bp in length) within the α-satellite HOR arrays (15 insertions and 28 deletions) that range in length from 340 bp to 19.4 kbp (**Fig. 5b**, **Supplementary Table 18**). Among these 43 SVs, 7 of them occur within the CDR (1 insertion and 6 deletions). Given that the CDR comprises only 8% of the active α-satellite HOR arrays on average (**Supplementary Table 17**), this is a twofold enrichment of SVs in the CDR than expected by chance. This elevated SV mutation rate in the CDR (**Supplementary Table 19**) may reflect the higher sequence identity of these regions, which are more prone to homologous recombination, leading to expansions and deletions, relative to less identical sequences. Among the 7 SVs that we observe in the CDR across generations, all of them fall within highly identical sequences, such as the deletion of a dimer α-satellite HOR in the chromosome 19 centromere from G2 to G3 (**Fig. 5c**) as well as a deletion of a 4-mer α-satellite HOR in the chromosome 2 centromere from G1 to G2, which is maintained in G3 (**Fig. 5d**). When this occurs, this often alters the epigenetic landscape of the centromere, splitting the CDR into two or more sites and/or shifting its position in the next generation (**Supplementary Table 20**, **Supplementary Fig. 77**).

Together, our findings suggest a model for centromere mutation and evolution that integrates the genetic and epigenetic variation we observe across global populations as well as the mutational landscape among recent generations (**Fig. 5e**). In this model, we find that the putative kinetochore site, marked by the CDR and an enrichment of CENP-A chromatin, typically resides within the most highly identical and homogenous sequences of the centromeric α-satellite HOR array. Due to its high sequence identity and homogeneity, this region is prone to mutations at both the single-nucleotide and structural level, which alter its position, distribution, and length and lead to changes in both genetic and epigenetic landscapes over time. Cell-based assays that identify the mutational processes that shape centromeric diversity across generations will need to be performed in the future to confirm this hypothesis.

## DISCUSSION

Here, we present the first comprehensive, population-scale analysis of human centromere sequence, structure, and epigenetic variation by assembling and characterizing 2,110 complete centromeres from 65 diverse individuals and complementing it with 5,747 centromeres from 227 individuals and 483 centromeres from a four-generation pedigree (8,340 centromeres total). While recent studies have provided valuable insights into centromeric diversity, including comparative analyses of two complete centromere sets (CHM13 and CHM1) that revealed substantial sequence and structural variation between haplotypes^9^, as well as characterization of 1,246 centromeres from diverse genomes focused primarily on structural variants^12^, our work uniquely integrates three critical dimensions at unprecedented scale. First, we expand the number of available complete centromere sequences by more than five-fold compared to previous efforts, revealing 226 new major centromere haplotypes and 1,870 novel α-satellite HOR variants. Second, unlike prior work that either compared limited numbers of complete centromeres or analyzed larger cohorts with incomplete assemblies, we combine deep sequence-level resolution across thousands of centromeres with comprehensive epigenetic profiling and multi-generational mutation tracking. Third, our integrated centromere maps make it possible to systematically link sequence variation, structural rearrangements, MEIs, and epigenetic organization to kinetochore positioning and recent evolutionary dynamics across human populations, establishing the foundation for understanding both the mechanistic basis of centromere evolution and its implications for chromosome segregation fidelity and human disease.

Our population-scale analysis uncovers multiple mechanisms that contribute to centromeric diversity. We identify large duplications, inversions, and deletions on ten chromosomes (chromosomes 1, 4, 5, 7, 10, 12, 13, 17, 19, and 20), as well as a rare recombination event between acrocentric centromeres (e.g., chromosomes 13/22), that alter the composition and organization of the centromeric α-satellite HOR arrays. We find that MEIs are present in ∼30% of centromeres and occur within the kinetochore site in 5% of centromeres, altering local methylation patterns. Chromosome 16 shows a striking 17-fold enrichment of MEIs within the kinetochore site. These structural and insertional changes provide mechanistic routes by which centromeric sequence and epigenetic landscapes can be remodeled on timescales ranging from recent (centuries) to ancient (>100 kya).

A central conclusion from our work is that the position of the kinetochore, marked by the CDR and an enrichment of CENP-A-containing nucleosomes, is not stochastic but is tightly coupled to local centromeric sequence and structure. Across chromosomes, we observe strong positional biases of the kinetochore site (p-arm-proximal, q-arm-proximal, or haplotype-dependent distributions) and show that the sequence underlying the kinetochore is, on average, more homogeneous and of higher sequence identity than the surrounding α-satellite HORs. This association indicates that certain sequence contexts (e.g. highly identical, low-entropy stretches of α-satellite DNA) are preferential substrates for kinetochore establishment and/or maintenance. This finding helps reconcile models in which centromeric proteins and DNA co-evolve (e.g. KARM^33,34^) and suggests that local sequence homogeneity may facilitate the assembly or maintenance of centromeric chromatin.

A particularly novel and biologically important result is the elevated mutation dynamics we observe at centromeres and specifically within kinetochore sites. Using both pan-centromere phylogenetic approaches and direct observation of *de novo* changes in a four-generation pedigree, we show that α-satellite HOR arrays mutate substantially faster than the flanking unique sequence (mean ∼4.0×10^-7^ Ms/bp/G), that centromeres exhibit a 58-fold range of mutation rates (e.g., chromosome Y much slower than chromosome 1), and that the CDR mutates more rapidly than the rest of the active α-satellite HOR array (twofold higher SNV rate and a marked enrichment of structural variants). These observations support a model in which homogeneous, kinetochore-associated regions are intrinsically prone to both nucleotide and structural changes, likely because sequence homogeneity facilitates unequal exchange and recombination events. The accelerated turnover at the functional site of the kinetochore offers a potential mechanistic basis for rapid centromere evolution and for a genetic-epigenetic ‘arms race’ between centromeric DNA and centromere-binding proteins.

Collectively, our findings provide the foundation for a unified model in which (i) kinetochores preferentially form on homogenous stretches of α-satellite HORs; (ii) those stretches are prone to elevated rates of homologous recombination and structural change because their sequence identity promotes misalignment and unequal exchange; and (iii) such genetic changes feed back on chromatin state and kinetochore localization, producing continual shifts in centromere architecture across generations (an evolutionary “treadmill” of DNA-protein interactions). This model integrates our observations of haplotype structure, epigenetic patterns, MEIs, and mutational spectra and is consistent with longstanding hypotheses about centromere drive and centromere-associated recombination machinery.

Implications for human biology and disease are multiple. First, the discovery that MEIs (and other structural changes) can occur within kinetochore sites raises the possibility that sequence perturbations directly influence chromosome segregation fidelity. The elevated incidence of MEIs in the chromosome 16 kinetochore site, a chromosome already known for nondisjunction propensity, is particularly intriguing, and future experiments that test the functional consequence of MEIs within kinetochore sites will need to be performed to confirm their role in increased chromosome missegregation rates. Second, population-stratified haplotypes and ancient introgressed centromeres may influence subtle differences in meiotic behavior or fertility across lineages and could be relevant for interpreting aneuploidy risk and patterns of chromosomal instability in disease. Finally, the rapid evolution of kinetochore-proximal sequence may have consequences for cross-species compatibility of centromere proteins and could inform models of reproductive isolation.

Looking forward, a number of experiments can build on this resource: biochemical reconstitution of kinetochore assembly on defined α-satellite HOR arrays, single-molecule imaging of kinetochore function on engineered centromeres, cell-based assays that measure segregation fidelity after targeted MEI or SV introduction, and population genetic screens for associations between centromere haplotypes and reproductive outcomes. On the computational side, integrating our centromere maps with large-scale clinical and cytogenetic datasets may reveal links between centromere genotype and aneuploidy spectra in embryos or cancer. Finally, extending surveillance to additional pedigrees and to gamete sequencing will clarify the mutation processes (replication errors, recombination-mediated events, retrotransposition) that most strongly shape centromeric evolution.

In summary, complete centromere sequence assemblies and comprehensive analyses presented here reveal centromeres as dynamic genomic regions where sequence homogeneity, structural changes, mobile elements, and epigenetic state interact to shape kinetochore position and evolution. Our results provide a foundation for future mechanistic experiments and for integrating centromere biology into models of human genetic disease, fertility, and genome evolution.

## METHODS

### Centromere mapping and annotation

To identify completely and accurately assembled centromeres in whole-genome assemblies from diverse humans, we developed a pipeline called the Centromere Mapping and Annotation Pipeline (CenMAP; https://github.com/logsdon-lab/CenMAP; v0.4.3.1). Briefly, CenMAP aligns each whole-genome assembly to the centromeric regions of the CHM13 reference genome^7^ (v2.0) using minimap2^41^ (v2.28) with the following parameters: `-x asm20 --secondary=no -s 25000 -K 15G`. The resulting BAM files are then converted into BED files using rustybam (v0.1.34; https://mrvollger.github.io/rustybam), and the BED files are intersected with a BED file containing coordinates of unique regions within the pericentromeres that correspond to each chromosome using BEDTools^42^ (v2.31.1). Contigs spanning the centromeres are subsequently identified and renamed to indicate the chromosome they most accurately align to based on the greatest number of alignments to the pericentromeric region. To confirm that the centromeric contigs contain α-satellite sequences, CenMAP runs dna-brnn^43^ (v0.1-r65), which annotates α-satellite repeats within each centromeric contig. Neighboring α-satellite repeat annotations are iteratively merged if they are within 10 kbp of each other, yielding the coordinates of the centromeric α-satellite HOR array(s). The coordinates of the α-satellite HOR array(s) are extended on either side by approximately 500 kbp, producing the coordinates of the centromeric regions within each contig.

#### Evaluation of contiguity and accuracy within centromeric regions

CenMAP evaluates the contiguity and accuracy of the centromeric regions by first extracting the regions via BEDTools and annotating them with RepeatMasker^44^ (v4.1.0) using the following parameters: `- engine rmblast -species human -dir {output_directory} -pa {threads} {input_contig}`. CenMAP uses these annotations along with a custom tool we developed, CenStats (https://github.com/logsdon-lab/CenStats; v0.0.10), to check the completeness of the contigs and reorient misoriented contigs from p to q arm relative to the CHM13 reference genome (v2.0). This is achieved with the subcommand `status`, which compares the repeat sequence edit distance and Jaccard index relative to the reference genome, selecting the best matching orientation. Completely assembled centromeric contigs are subsequently reoriented to match the orientation of centromeres within the CHM13 reference genome (v2.0) and then evaluated for the presence of assembly errors. To detect assembly errors in the centromeric regions, we developed NucFlag (v0.3.3), an upgraded version of NucFreq^45^ that uses peak detection in the first- and second-highest read pileup nucleotide frequency to detect collapsed and misjoined sequences within the contigs. With NucFlag, PacBio HiFi reads are first aligned to the whole-genome assemblies using pbmm2 (v1.13.1) (https://github.com/PacificBiosciences/pbmm2) with the following parameters: `--log-level DEBUG - -preset SUBREAD --min-length 5000`. Alignments are filtered using samtools^46^ (v1.22) and the filter flag 2308, to remove secondary, supplementary, and low-quality alignments. Only regions annotated as “ALR/Alpha” by RepeatMasker are evaluated, and regions with nucleotide frequencies that are <95% of the mean of the highest nucleotide frequency are considered candidate misjoins, while regions with >225% of the mean highest nucleotide frequency are considered collapses. If the heterozygous site ratio exceeds 20% of the mean second nucleotide frequency, these regions are considered smaller collapses. Only centromeric regions that are free of misjoins, collapses, or other assembly errors are considered accurately assembled and retained for downstream analyses.

#### Annotation of centromeric regions

To determine the sequence composition and organization of all centromeric regions, CenMAP uses a custom tool we developed, Snakemake-HumAS-SD (https://github.com/logsdon-lab/Snakemake-HumAS-SD; v0.1.0), which leverages the α-satellite monomer HMM libraries from HumAS-HMMER (https://github.com/fedorrik/HumAS-HMMER_for_AnVIL) with the rapid annotation of StringDecomposer^47^ to efficiently annotate the centromeric regions. Snakemake-HumAS-SD first constructs a comprehensive α-satellite monomer library for each chromosome based on an existing HumAS-HMMER HMM profile (https://github.com/fedorrik/HumAS-HMMER_for_AnVIL/blob/main/AS-HORs-hmmer3.4-071024.hmm). Chromosomes with shared α-satellite monomers, such as chromosomes 1/5/19 as well as chromosomes 13/14/21/22, had their libraries merged. We developed a new α-satellite HOR variation detection method that generates BED-format annotations of α-satellite HOR variants for each centromere based on the previous StV approach (https://github.com/fedorrik/stv). The method takes α-satellite monomer sequences as input and segments them based on three criteria: (i) adjacent monomers are separated by gaps >160 bp, (ii) adjacent monomers are on opposite strands, or (iii) monomer IDs are not monotonically increasing (on the forward strand) or decreasing (on the reverse strand). Compared to the original StV method, our approach can detect previously unresolvable HOR structures, like the newly identified 8-mer (S1C16H1L.2-9) expansion on chromosome 16, and achieves α-satellite dHOR annotation not annotated by the previous StV method.

#### Estimation of the length of active α-satellite HOR arrays within centromeric regions

To estimate the length of active α-satellite HOR arrays, we ran CenStats (v0.0.10) with the `length` subcommand, which leverages the annotations generated by Snakemake-HumAS-SD (described above) to output length statistics of each active α-satellite HOR array. CenStats first uses Snakemake-HumAS-SD α-satellite HOR StV BED files to identify the coordinates of active α-satellite HOR monomers (those with names containing ‘L’). Adjacent active α-satellite HOR monomers are then merged into one block if they are within ∼1.5 α-satellite monomer lengths (256 bp) of each other, and blocks containing fewer than two α-satellite monomers are removed. To account for interruptions of stretches of α-satellite HORs caused by mobile element insertions (MEIs), adjacent blocks of active α-satellite HORs separated by less than 8 kbp are iteratively merged into a single active α-satellite HOR array. Finally, the coordinates of α-satellite HOR arrays that consist of at least five α-satellite HOR monomers and have at least 90% of their total length composed of active α-satellite HORs is outputted as a BED file. To determine the orientation of active α-satellite HOR arrays, we followed a similar procedure, but incorporated both strand orientation and distance to group active α-satellite HOR monomers into arrays.

#### Detection of CDRs within centromeric regions

In addition to annotating centromeric sequences, CenMAP uses methylation information in native long-read sequencing data to annotate the CpG methylation profile across each centromeric region. CenMAP first aligns native ONT long-read sequencing data from each genome to the relevant whole-genome assembly using minimap2^41^ (v2.28) with the following command: `minimap2 -x lr:hqae -s 4000 - y -I 8G {ref} {ont_reads}`. CenMap filters the alignments using SAMtools^46^ (v1.9) -F 2308 to retain only primary alignments and remove partial and secondary alignments. Finally, CenMAP uses CDR-Finder^28^ (v1.0.4), a tool to detect hypomethylated regions termed “centromere dip regions”, with default parameters, to identify candidate CDRs. We manually curated the CDR calls by checking each region outputted in the BED file using IGV^48^, removing false positives and recovering missed calls.

#### Pairwise sequence identity heat maps

To generate pairwise sequence identity heat maps of each centromeric region, we ran ModDotPlot^49^ (v0.8.4) with the following command: `moddotplot static -f {fasta} -w 5000 -id 70.0`. We normalized the color scale across the ModDotPlots by binning the % sequence identities and recoloring the data points according to the binning.

#### Visualization of centromeric regions

We also developed a Python-based tool, CenPlot (v0.1.4; https://github.com/logsdon-lab/CenPlot), for visualizing the various types of annotations produced by CenMAP. Each data type is formatted into a BED4 or BED9 file and plotted as tracks ordered in a configuration TOML file via the following command: `cenplot draw -t {track_toml} -c {chrom} -d {outdir}`. These tracks include RepeatMasker annotation, α-satellite HOR annotation produced by Snakemake-HumAS-SD (https://github.com/logsdon-lab/Snakemake-HumAS-SD), active α-satellite HOR array strand orientation from CenStats, CpG methylation frequencies produced by CDR-Finder^28^ (v1.0.4) as well as the 1D or 2D sequence identity heat maps produced by ModDotPlot^49^.

### Assembly repair within centromeric regions

With the ability to detect assembly errors (described above), we then developed a new repair tool, AssemblyRepairer (https://github.com/logsdon-lab/AssemblyRepairer; v1.0), to repair erroneous regions within the Verkko centromere assemblies using correctly assembled regions from the hifiasm centromere assemblies. For each Verkko centromere containing errors in α-satellite sequences, we aligned the hifiasm assembly from the same individual using minimap2^41^ (v2.28) with the following parameters: `-x asm5 --eqx --cs --secondary=no -s 25000 -K 8G`. Alignment records with mapping quality (MAPQ) >20 were kept, adjacent alignments within 5 kbp were merged, and alignments with >30% overlap were removed. Resulting alignments <100 kbp were excluded from downstream analysis. This step yielded one or more corresponding regions in the hifiasm assemblies for each erroneous Verkko centromere. NucFlag (v0.3.3) was then run to identify errors in the matched hifiasm regions. Unique *k*-mers of length 5 kbp were generated from each Verkko centromere, and 1,000 were randomly selected as anchors. Anchors mapping to multiple sites in the hifiasm assemblies were discarded to ensure uniqueness. The remaining anchors were used to precisely locate erroneous regions across both assemblies. Erroneous Verkko regions were replaced with the corresponding correct hifiasm sequences, and NucFlag was run on the repaired assemblies to ensure that the regions were accurately repaired. Regions with remaining errors were excluded from downstream analysis. We finally merged the remaining centromeres that were completely and accurately assembled in hifiasm but not in Verkko with repaired Verkko centromeres and generated new whole-genome assemblies for all 65 samples. Each repaired whole-genome assembly was then validated with HMM-Flagger^18^ (v1.1.0), NucFlag^19^ (v0.3.3), GAVISUNK^20^ (v1.0), and Merqury^21^ using PacBio HiFi, ONT, Illumina, or Element data. For HMM-Flagger, we first aligned all PacBio HiFi and ONT reads to the repaired assembly using minimap2^41^ (v2.28) with the following parameters: `-y -a --eqx --cs -x lr:hqae -I8g`. Each BAM was then run through the default HMM-Flagger pipeline using the corresponding data type’s alphaTsv file. We applied GAVISUNK, a singly-unique kmer (SUNK) -based error detection method (with parameter 30-mer), to validate α-satellite regions. Merqury (v1.3) was used to obtain quality value scores for each centromeric region with the following command: `meryl k=21 count {input_reads} output {output_db} && merqury.sh {output_db} {input_asm}`. Finally, we manually checked each centromeric region for uniform PacBio HiFi and ONT coverage, ensuring they were free of large assembly errors.

### Estimation of local sequence identity and entropy across centromeric regions

To determine the local sequence identity across each centromeric region, we averaged the pairwise sequence identity values derived from ModDotPlot^49^ (v0.8.4) BEDPE format files. For each region, genomic intervals were divided into 5-kbp bins, and sequence identity values between bin pairs were used to construct a symmetric identity matrix. A sliding window of 25 kbp (five consecutive bins) was applied along the diagonal of the matrix with a 5-kbp step size. For each window, local sequence identity was calculated as the mean of all off-diagonal identity values, excluding those within a 3-bin neighborhood of the diagonal to reduce self-similarity bias. Average sequence identities of active α-satellite HOR arrays were calculated for arrays >50 kbp. To remove the influence of outliers at array boundaries, we excluded one window from the end of the active α-satellite HOR arrays. To define regions of increased admixture within active α-satellite HOR arrays, we calculated the entropy of the entire α-satellite HOR array using the frequencies of the different types of HOR units in 10-unit windows with 1-unit slide with the following equation:

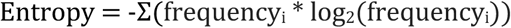

### Structural variation detection within centromeres

To detect large-scale variation between centromeres, we ran SVbyEye^50^ (v0.99.0). We first aligned contigs containing a centromeric region to each other using minimap2^41^ (v2.28) and the following command: `minimap2 -x asm20 --c --eqx --secondary=no {target.fasta} {query.fasta} > {output.paf}`, generating a PAF file. We then visualized these alignments with R (v4.4.2) using the plotting function: `plotMiro(paf.table = paf.table, color.by = “direction”)`.

### Mobile element insertion (MEI) detection, annotation, and curation

To detect mobile element insertions within centromeric α-satellite HOR arrays, we ran RepeatMasker^44^ (v4.1.6) with the Dfam^51^ (v3.8) library and -s parameter to identify unique non-LTR retroposons (*Alu*, LINE-1, and SVAs) within completely and accurately assembled centromeric regions. For each unique non-LTR retroposon, we extracted its sequence along with an additional 30 base pairs of flanking sequence on either side using Pysam^52,53^ (v0.22.0). Then, we reran RepeatMasker^44^ (v4.1.6) with the Dfam^51^ (v3.8) library and -s parameter on these fastas, generating outfile. The RepeatMasker outfiles and fastas were inputted into L1ME-AID (v1.2.0-beta) with default settings for MEI identification, as previously described^12^. MEIs on the same centromeric α-satellite HOR array were aligned to one another using MUSCLE^54^ (v3.8.31), and alignments were manually curated to identify unique insertion events as well as their duplications. Finally, we intersected the MEI calls with coordinates of each centromeric α-satellite HOR array using BEDTools^42^ (v2.31.1) to derive the final callset within centromeric α-satellite HOR arrays.

### Cell culture

Human cell lines HG00512, HG00513, HG00514, HG02769, HG03065, and GM20355 were purchased from the Coriell Institute for Medical Research (Camden, NJ) and cultured in RPMI-1640 medium with 2mM L-glutamine (Corning, MT10040CV) and supplemented with 15% FBS (Gibco, A5669701) and 1% penicillin-streptomycin (Thermo Fisher Scientific, 15140122). All cell lines were cultured in a humidity-controlled environment at 37°C with 95% O_2_ and 5% CO_2_.

### CENP-A DiMeLo-seq

DiMeLo-seq was performed as described in Ref. ^8^ with some modifications for Oxford Nanopore Technologies (ONT) long-read sequencing. Briefly, 7×10^6^ cells were resuspended in 1 ml of Dig-Wash buffer [20 mM HEPES-KOH, pH 7.5, 150 mM NaCl, 0.5 mM Spermidine, 1 tablet of cOmplete, EDTA-free Protease Inhibitor Cocktail (11873580001, Roche) per 50 ml buffer, 0.1% w/v BSA, and 0.02% Digitonin] and incubated on ice for 5 mins to isolate nuclei. Nuclei were pelleted and resuspended in 200 μl of Tween-Wash buffer (20 mM HEPES-KOH, pH 7.5, 150 mM NaCl, 0.5 mM Spermidine, 1 tablet of cOmplete, EDTA-free Protease Inhibitor Cocktail per 50 ml buffer, 0.1% w/v BSA, and 0.1% w/v Tween-20) containing a primary antibody at a 1:50 dilution. Primary antibodies used in this study were mouse anti-CENP-A antibody (ADI-KAM-CC006, Enzo) or rabbit anti-IgG antibody (ab171870, Abcam). Samples were rotated overnight on a nutator at 15 rpm and 4°C to allow for antibody-binding. A mouse nanobody-Hia5 (Nb-Hia5), gifted by Nicolas Altemose (Stanford University), was prepared as a secondary antibody at 200 nM in 200 μl of Tween-Wash buffer. After washing away unbound primary antibody with Tween-Wash buffer, samples were resuspended in secondary antibody buffer and rotated for 2 hours on a nutator at 15 rpm and 4°C. The methyl donor S-Adenosyl Methionine (SAM; B9003S, NEB) was prepared at 800 μM in 100 μl of activation buffer (15 mM Tris, pH 8.0, 15 mM NaCl, 60 mM KCl, 1 mM EDTA, pH 8.0, 0.5 mM EGTA, pH 8.0, 0.05 mM Spermidine, and 0.1% w/v BSA). After washing away unbound Nb-Hia5, samples were resuspended with SAM activation buffer and incubated at 37°C for 2 hours to activate the DNA adenine methyltransferase activity of Hia5. After incubation, samples were collected and resuspended in 400 μl of cold PBS for genomic DNA (gDNA) extraction using the Monarch HMW DNA Extraction Kit for Cells & Blood (T3050S, NEB). For gDNA extraction and ONT library preparation, the FindingNemo protocol^55^ was performed with the following changes: (1) extracted gDNA was kept at RT for at least 48 hours to improve DNA homogeneity, (2) the FRA (transposase fragmentation mix) in the Ultra-Long DNA Sequencing Kit V14 (SQK-ULK114, ONT) was adjusted by adding 8 μl for optimal DNA fragmentation, (3) library DNA was eluted in 100 μl of 10 mM Tris-HCl (pH 9.0) for a single library load. DNA libraries were loaded onto a PromethION 24 R10.4.1 flow cell (FLO-PRO114M, ONT) and sequenced with MinKNOW v25.03.7 (ONT). To enrich centromeric sequences during sequencing, we used the deplete mode of adaptive sampling enabled in MinKNOW to exclude sequencing reads originating from outside centromeric regions. The depletion BED file was made based on centromere annotations of the CHM13 reference genome (v2.0). Libraries were retrieved and reloaded after 20-24 hours, depending on pore availability.

Both anti-CENP-A and anti-IgG DiMeLo-seq ONT data were basecalled with dorado v0.9.6+0949eb8d using the following parameters and model: sup,5mCG_5hmCG,6mA --min-qscore 10; basecall_model=dna_r10.4.1_e8.2_400bps_sup@v5.0.0,modbase_models=dna_r10.4.1_e8.2_400bps _sup@v5.0.0_5mCG_5hmCG@v3,dna_r10.4.1_e8.2_400bps_sup@v5.0.0_6mA@v3. Reads with abnormally high 6mA methylation proportions (# of 6mA sites with a base modification probability greater that 80% / total # of AT sites) were filtered out to reduce noise.

We determined outlier reads by calculating the global mean and standard deviation of the proportion of 6mA over all reads and calculating a standard score for each read. Any read with greater than two z-scores above the mean was filtered using samtools. Then, filtered reads were aligned to their respective assemblies using minimap2^41^ (v2.28) with following parameters: `-y -a --eqx --cs -x lr:hqae - I8g -s 4000`. To quantify 6mA methylation, we applied modkit (v0.3.2, https://github.com/nanoporetech/modkit) using the pileup command with the following parameters: `modkit pileup {input.bam} {output.bed} --ref {input.fa} --motif AT 0 --filter-threshold 0.8`. Finally, using a custom pipeline, Snakemake-DiMeLo-seq-Viz (v0.1.0, https://github.com/logsdon-lab/Snakemake-DiMeLo-seq-Viz), we binned each 6mA window into 5 kbp non-overlapping windows, calculated the average 6mA signal, and subtracted the average IgG signal from the average CENP-A signal to determine the normalized CENP-A signal per sample. The resulting BedGraph file was visualized using CenPlot (v0.1.4).

### CENP-A CUT&RUN and ONT long-read sequencing

CUT&RUN was performed following the CUTANA protocol developed by EpiCypher with some modifications. For each reaction, 550,000 cells were harvested and nuclei were isolated following the nuclei extraction protocol for CUTANA assays (EpiCypher). Nuclei were immobilized with activated Concanavalin A (ConA) beads provided in the CUTANA ChIC/CUT&RUN Kit (14-1048, EpiCypher). After binding to ConA beads, nuclei were incubated overnight at 4°C in 50 μl of Antibody buffer (1X Proteinase Inhibitor, 0.5 mM Spermidine, 0.01% w/v Digitonin, and 2 mM EDTA in Pre-wash buffer) with either mouse anti-CENP-A antibody (ADI-KAM-CC006, Enzo) or a rabbit anti-IgG antibody (ab171870, Abcam) at a 1:50 dilution. An appropriate amount of K-MetSat Panel was added to the reaction before adding the IgG negative control. Following overnight incubation at 4 °C, rotating on a nutator at 15 rpm, samples were washed and incubated in Cell Permeabilization buffer (1X Proteinase Inhibitor, 0.5 mM Spermidine, and 0.01% w/v Digitonin in Pre-wash buffer) with pAG-MNase (15-1016, EpiCypher) at 1:20 dilution for 10 mins at RT. After pAG-MNase bound to the primary antibody, 100 mM CaCl_2_ (EpiCypher) was added to activate chromatin-targeted digestion at 4 °C for 2 hours. Digestion was stopped by adding Stop buffer (EpiCypher), and samples were transferred to low-bind tubes for DNA purification and extraction following the fragment release and cleanup in the protocol of Hainer & Fazzio^56^. Each sample was brought up to 300 μl with 0.1X TE (EpiCypher), followed by addition of 3 ul of 10% SDS and 2.5 ul of 20 mg/ml Proteinase K (P8107S, NEB), then incubated at 70 °C for 10 minutes.

DNA extraction was performed using one phenol:chloroform:isoamyl alcohol (25:24:1 v/v; 15593031, Invitrogen) extraction step, followed by transfer of the aqueous phase to a MaXtract High Density 2ml tube (129056, QIAGEN), then one chloroform (C2432, SIGMA) extraction step, again transfer the aqueous phase to a MaXtract tube. DNA was precipitated with 750 μl of 100% ethanol and 2 μl of Glycogen (AM9510, AMB) and stored overnight at -20 °C. CUT&RUN-enriched DNA was washed with 100% ethanol and eluted in 0.1X TE.

For ONT library preparation and sequencing, 0.5-5 ng of CUT&RUN-enriched DNA was amplified using CUTANA CUT&RUN Library Prep Kit (14-1001 & 14-1002, EpiCypher). Up to the indexing PCR step, the KOD Xtreme Hot Start DNA polymerase (71975-3, SIGMA) was used for fragment enrichment. Each 50ul of PCR reaction contains 2X KOD PCR buffer, 0.4 mM dNTPs, 1.5 μl of i7 primer, 1.5 μl of i5 primer, and 1 μl of KOD polymerase. PCR cycling conditions were: 2-min initial denaturation at 94°C; 20 cycles of 10-second denaturation at 98°C, 30-s primer annealing at 55 °C, and 5-min extension at 68°C; followed by a final 5-min extension at 68°C. The sequencing library was prepared using the Oxford Nanopore Technologies (ONT) SQK-NBD114.24 kit as recommended by Hook et al., unpublished and sequenced on a PromethION 24 using a R10.4.1 flow cell (FLO-PRO114M, ONT).

The resulting CUT&RUN ONT data was processed using a custom pipeline we developed called Snakemake-CutNRun (https://github.com/logsdon-lab/Snakemake-CutNRun). In this pipeline, adapters are first trimmed using cutadapt^57^ (v5.1) with following parameters: `-e 0.1 -q 20 -O 5 -a AGATCGGAAGAGC {reads.fastq}`. Then, reads =100 bp or >10 kbp are removed using seqkit^58^ (v2.10.0) with the following command: `seq -m 100 -M 10000 {reads.fastq}`. The reads are aligned to the relevant whole-genome assembly using BWA-MEM^59^ (v0.7.19) with the following parameters: `-k 50 -c 1000000 {ref.fasta} {reads.fastq}`. Finally, CENP-A alignments are normalized with IgG alignments using deepTools^60^ (v3.5.6) with the following command: `bamCompare - b1 {bam_CENPA} -b2 {bam_IgG} --operation ratio --binSize 5000 --minMappingQuality 0`. The resulting BigWig is visualized with CenPlot (v0.1.4, https://github.com/logsdon-lab/CenPlot).

### Fluorescence *in situ* hybridization (FISH)

To detect a hybrid chromosome 13/22 centromere in human NA20355 cells, we performed FISH on metaphase chromosome spreads as previously described^6,61^ with slight modifications. Briefly, human NA20355 cells were treated with colcemid and swollen in a 0.56% potassium chloride (KCl) hypotonic solution at 37°C. After 30 min, cells were fixed with methanol:acetic acid (3:1) and dropped onto previously cleaned slides. Slides were aged in a drying oven at 90°C for 1.5 h, treated with pepsin (0.005% in 0.01 N HCl) at 37°C for 30 min, hybridized with labelled FISH probes at 70°C for 2 min, and incubated overnight at 37°C.

The FISH probes used to validate the hybrid chromosome 13/22 centromere were either derived from a pZ21A plasmid clone harboring a portion of the *D21Z1* α-satellite HOR array^62^ (Cy3; red) or generated via PCR using primers targeting the p-arm portion of the S2C14/22H1L α-satellite HOR array of chromosomes 14 and 22 (FluorX; green) according to the following procedure. First, a PCR reaction was prepared in a total volume of 25 µL, containing 0.25 µM forward primer (5′-GGTCCTCAAACCGTCCGAAA-3′), 0.25 µM reverse primer (5′-CAGCTTTCAGGTCTATGGTGAG-3′),

5 ng/µL human genomic DNA, 50% DreamTaq PCR Master Mix 2X (DreamTaq DNA Polymerase, 2X DreamTaq buffer, dATP, dCTP, dGTP, and dTTP at 0.4 mM each, and 4 mM MgCl₂), and double-distilled water (ddH₂O). PCR was performed using the following thermal cycling conditions: initial denaturation at 94°C for 30 s, followed by 30 cycles of 94 °C for 30 s (denaturation), 65°C for 1 min (annealing), and 72°C for 1 min (extension). After the last cycle, a final extension was carried out at 72°C for 10 min, and the reaction was held at 4°C. The resulting PCR product was subsequently labeled via PCR in a 25 µL reaction containing 2 uL PCR product, 10% Taq Buffer with (NH_4_)_2_SO_4_ (10X), 2 mM MgCl₂, 0.2 µM forward primer, 0.2 µM reverse primer, 0.2 mM dACG, 0.1 mM fluorescein-dUTP, 0.4% BSA, 0.06 U/µL Taq DNA polymerase, and ddH₂O. PCR amplification was performed as described above.

Slides were washed with high stringency three times in 0.1× SSC at 60°C for 5 min each before mounting in Antifade and DAPI-staining. Slides were imaged on a epifluorescence microscope (Leica DM RXA2) equipped with a charge-coupled device camera (CoolSNAP HQ2) and a 100× 1.6–0.6 NA objective lens. DAPI, Cy3, and fluorescein fluorescence signals, detected through specific filters, were individually collected as grayscale images using the iVision software (v4.5). Pseudocoloring and merging of the images were performed using the Adobe Photoshop software.

### Phylogenetic reconstruction of centromeres

To reconstruct the evolutionary history of centromeres, we first identified orthologous sequences between human and chimpanzee by aligning the CHM13 (v2.0) genome to the mPanTro3 (v2.1) genome^35^ via minimap2^41^ (v2.28) with the following parameters: `-ax asm20 --eqx -Y --secondary=no`. From this set of orthologous sequences, we selected 20 kbp regions in the monomeric α-satellite or unique sequences flanking the α-satellite HOR arrays in both the p and q arms, prioritizing regions closest to the α-satellite HOR arrays. For acrocentric chromosomes (13-15, 21, and 22), only q-arm regions were included due to the high abundance of satellites and segmental duplications on the p-arm as well as the known homologous recombination between acrocentric chromosomes in the p arms that prevent accurate phylogenetic reconstruction^23^. We aligned the 20-kbp CHM13 sequences to the 65 repaired human genome assemblies as well as the mPanTro3 (v2.1) genome^35^ using the same minimap2 parameters described above. We limited our analyses to only those regions with one-to-one unambiguous mapping, and we excluded sequences with alignment lengths >10% of the aligned region (2 kbp). A multiple sequence alignment was performed using MAFFT^63^ (v.7.526) with the --auto setting. Maximum-likelihood phylogenies were inferred using IQ-TREE^64^ (v2.3.6) with automatic model selection (--auto) and 1,000 ultrafast bootstrap replicates, using the chimpanzee sequence as an outgroup. Divergence times were estimated under a molecular clock model by scaling branch lengths, assuming a human–chimpanzee divergence of 6.2 million years^35^. Phylogenetic trees were visualized using iTOL^65^ (v7.2.2; https://itol.embl.de/), and clades were defined by partitioning the phylogenetic tree based on the divergence time of 100 kya.

### Analysis of archaic centromere introgression

We applied ARCkmerFinder^36^ (v1.1, https://github.com/hsiehphLab/ARCkmerFinder), a *k*-mer-based method, to whole-genome assemblies for introgression detection. Briefly, we generated *k*-mer databases for each of the four archaic genomes^37–40^ and a joint *k*-mer database from African genomes in the 1KG short-read collection^66^ using Meryl^21^ (v1.4.1; *k*-mer size = 15 bp). Because low-frequency *k*-mers are more likely to represent sequencing errors, we modeled the observed *k*-mer frequency distribution for each short-read genome using a mixture of negative binomials to determine appropriate cutoff thresholds^36^. Archaic-specific *k*-mers were identified by subtracting the joint African *k*-mers from each archaic database, leveraging the limited archaic introgression in Africans to reduce ancestral polymorphism effects. For each assembly, we scanned contig sequences for the presence of archaic-specific *k*-mers and binned the signal within 2-kbp windows to aggregate local evidence of archaic sequence. Genome-wide cutoffs for detecting archaic introgression were then determined from whole-genome estimates of archaic sequence content.

### Pan-centromeric mutation rate estimation

To estimate the mutation rates of active α-satellite HOR arrays and flanking sequences for each centromere haplogroup, we first identified clades within the phylogenetic tree that contain at least four centromeres, from which one was selected as a reference and the remaining as queries. We then aligned the reference’s active α-satellite HOR array, along with 2 Mb of the adjacent pericentromeric region, to each query in the same clade using minimap2^41^ (v2.28) with the following parameters: `-x asm20 -K 8G --eqx -Y -r 500000,500000`. Unmapped, supplementary, and partial alignments were removed using SAMtools^46^ (v1.22) flag 260. We then generated a BED file of 10-kbp windows located within each reference centromere. We used subseq (https://github.com/EichlerLab/subseq) to extract query sequences. For each window, the corresponding query sequence was required to be between 8,000 and 12,000 bp in length and exhibit a one-to-one, unambiguous mapping. For sequences in each window, a multiple sequence alignment was performed using MAFFT^63^ (v.7.526) with the following parameters: `- -maxiterate 1000 --globalpair`. The mutation rate per segment was estimated based on Kimura’s model of neutral evolution^67^. In brief, we modeled the estimated divergence (D) as a result of between-species substitutions and within-species polymorphisms, i.e.,

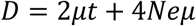

where *μ* is the mutation rate, *t* is the divergence time estimated from our phylogenetic trees, and Ne is the ancestral human effective population size. We assume a generation time of 20 years and a human-chimpanzee divergence time of 6.2 million years. To convert the genetic unit to a physical unit, our computation also assumes Ne=10,000 and uniformly drawn values for the generation and divergence times. The estimated divergence (*D*) is calculated from the Tamura-Nei genetic distance using the libtn93 library (https://github.com/sdwfrost/libtn93?tab=readme-ov-file#api) with a match mode of RESOLVE and a minimum overlap of 100 characters.

### Detection of *de novo* SNVs and SVs among centromeres across generations

To detect SNVs and SVs within centromeric regions across generations (in the CEPH 1463 pedigree), we first aligned each child’s centromere to its corresponding parental centromere using minimap2^41^ (v2.28) with the following parameters: `-ax asm5 --eqx --cs --secondary=no -s 25000 -K 8G`. Only high-confidence, uniquely mapped alignments were retained for downstream analysis. SNVs and SVs were then detected by parsing the CIGAR string. SNVs were identified from ‘X’ operations in the CIGAR string, where each ‘X’ represents a single base mismatch. SVs were identified from insertions (‘I’) or deletions (‘D’) of at least 50 bp in the CIGAR string. To validate the SNVs and SVs, we manually inspected each event using IGV plots generated from both PacBio HiFi and ONT read alignments for the child and parent, allowing us to exclude potential artifacts due to mosaicism or alignment errors.

### Multi-generational centromeric mutation rate estimation

To estimate the mutation rate among centromeric regions (in the CEPH 1463 pedigree), we first annotated the sequence of each centromeric region using RepeatMasker^44^ (v4.1.0) and Snakemake-HumAS-SD (https://github.com/logsdon-lab/Snakemake-HumAS-SD; v0.1.0), Tandem repeats finder^68^ (TRF; v4.09), and CDR-Finder^28^ (v1.0.4). We then intersected the validated SNVs identified from minimap2 alignments (described above) with each of these regions using BEDTools^42^ (v2.31.1). We divided the # of validated SNVs by the total # of base pairs (bps) in the parent for each region per generation to derive the estimated mutation rate, defined as the # of mutations per bp per generation.

## DATA AVAILABILITY

All datasets generated and/or used in this study are listed in **Supplementary Table 21** with their BioProject, accession # (if available), and/or URL. For convenience, we also list their BioProjects and/or URLs here: 65 original human whole-genome assemblies generated by the HGSVC (PRJEB76276 and PRJEB83624); 65 human whole-genome assemblies with repaired centromeric contigs (PRJNA1309774); PacBio HiFi and ONT data from 65 humans generated by the HGSVC (PRJEB58376, PRJEB75216, PRJEB77558, PRJEB75190, PRJNA698480, PRJEB75739, PRJEB36100, PRJNA988114, PRJNA339722, PRJEB41778, ERP159775); Illumina data from 65 humans generated by the HGSVC (ftp://ftp.sra.ebi.ac.uk/vol1/run/ERR323, ftp://ftp.sra.ebi.ac.uk/vol1/run/ERR324, ftp://ftp.sra.ebi.ac.uk/vol1/run/ERR398, ftp://ftp-trace.ncbi.nlm.nih.gov/giab/ftp/data/AshkenazimTrio/HG002_NA24385_son/NIST_Stanford_Illumina_6kb_matepair); CENP-A and IgG DiMeLo-seq data from 6 humans (SRR35424754, SRR35424755, SRR35269316, SRR35269317, SRR35269314, SRR35269315, SRR35144393, SRR35144394, SRR35424752, SRR35424753, SRR35048265, SRR35048266), CENP-A and IgG CUT&RUN ONT data from NA20355 (SRR35048263, SRR35048264); 227 original human whole-genome assemblies generated by the HPRC (https://github.com/human-pangenomics/hprc_intermediate_assembly/blob/main/data_tables/assemblies_pre_release_v0.6.1.index.csv); PacBio HiFi data from 227 humans generated by the HPRC (https://github.com/human-pangenomics/hprc_intermediate_assembly/blob/main/data_tables/sequencing_data/data_hifi_pre_release.index.csv, https://github.com/human-pangenomics/hprc_intermediate_assembly/blob/main/data_tables/sequencing_data/data_deepconsensus_pre_release.index.csv); ONT data from 227 humans generated by the HPRC (https://github.com/human-pangenomics/hprc_intermediate_assembly/blob/main/data_tables/sequencing_data/data_ont_pre_release.index.csv); 28 human whole-genome assemblies, PacBio HiFi data, and ONT data from the CEPH 1463 pedigree (PRJEB86317 or dbGaP accession: phs003793.v1.p1); Element data from the CEPH 1463 pedigree (PRJEB3246); CHM13 whole-genome assembly (PRJNA559484); CHM1 whole-genome assembly (PRJNA975207); Altai Neanderthal (PRJEB1265); Vindija Neanderthal (PRJEB21157); Chagyrskaya Neanderthal (http://ftp.eva.mpg.de/neandertal/Chagyrskaya/); Denisovan (PRJEB3092); and chimpanzee (AG18354) whole-genome assembly (PRJNA916736). In addition, centromere sequences and annotations for the 65 whole-genome assemblies containing repaired centromeres as well as 17 CEPH 1463 whole-genome assemblies containing repaired centromeres consented for public release are available from figshare (https://figshare.com/s/cb839beea6ed06d641db).

## CODE AVAILABILITY

All custom code used in this project are publicly available at the following GitHub repositories: CenMAP (https://github.com/logsdon-lab/CenMAP); CenStats (https://github.com/logsdon-lab/CenStats); NucFlag (https://github.com/logsdon-lab/NucFlag); CenPlot (https://github.com/logsdon-lab/CenPlot); Snakemake-HumAS-SD (https://github.com/logsdon-lab/Snakemake-HumAS-SD); AssemblyRepairer (https://github.com/logsdon-lab/AssemblyRepairer); DiMeLo-seq-Viz (https://github.com/logsdon-lab/Snakemake-DiMeLo-seq-Viz); CUT&RUN-Viz (https://github.com/logsdon-lab/Snakemake-CutNRun); and pan-centromeric mutation rate estimation (https://github.com/logsdon-lab/Snakemake-MutationRate).

## ACKNOWLEDGMENTS

We thank Y. Nechemia-Arbely (Pitt), M. Mahlke (Pitt), G. Hartley (UConn), R.J. O’Neill (UConn), and Z. Liu (UPenn), P. Hook (AJHG), W. Timp (JHU), and N. Altemose (Stanford) for reagents, protocols, and advice. We thank E.E. Eichler (UW) and J.L. Gerton (Stowers) for helpful discussions and suggestions. This research was supported, in part, by funding from the National Institutes of Health (NIH) National Institute for General Medical Sciences (NIGMS) R00GM147352 (to G.A.L.) and 1P20GM139769 (to M.K.K. and M.L.). This work was also supported by funding from the NIH National Human Genome Research Institute (NHGRI) 5R00HG011041 (to. P.H.) and U41HG010972, U01HG010971, U01HG013760, U01HG013755, U01HG013748, U01HG013744, R01HG011274 (to the Human Pangenome Reference Consortium; BioProject ID: PRJNA730823). Additionally, this work was supported by the Searle Scholars Program (to G.A.L.).

## AUTHOR CONTRIBUTIONS

S.G., K.K.O., and G.A.L. conceived the project; S-C.C., A.M., and M.V. performed experiments. S.G., K.K.O., M.L., D.S.G., and G.A.L. analyzed the data; P.H., M.K.K., M.V., and G.A.L. supervised the project; S.G., K.K.O, and G.A.L. wrote, edited, and revised the manuscript. All authors have read and approved the manuscript.

## HUMAN GENOME STRUCTURAL VARIATION CONSORTIUM AUTHORS

Hufsah Ashraf^1,2^, Peter A. Audano^3^, Marcelo Ayllon^4^, Andrey Azov^5^, Parithi Balachandran^3^, Anna O. Basile^6^, Christine R. Beck^3,7^, Marc Jan Bonder^8-10^, Lucy Brooks^5^, Marta Byrska-Bishop^6^, Mark J. P. Chaisson^11^, Zechen Chong^12^, André Corvelo^6^, Jonathan Crabtree^13^, Scott E. Devine^13^, Peter Ebert^14,2^, Jana Ebler^1,2^, Evan E. Eichler^4,15^, Aine Fairbrother-Browne^5^, Chia-Hsuan Fan^12^, Mallory Freeberg^5^, Shenghan Gao^16^, Mark B. Gerstein^17,18^, Bida Gu^11^, Pille Hallast^3^, Patrick Hasenfeld^19^, Mir Henglin^1,2^, Kendra Hoekzema^4^, Kaili Hu^12^, Sarah Hunt^5^, Matthew Jensen^17,18^, Yunzhe Jiang^17,18^, Kwondo Kim^3^, Miriam K. Konkel^20,21^, Jan O. Korbel^19^, Youngjun Kwon^4^, Peter M. Lansdorp^22^, Charles Lee^3^, Tiffany Leung^22^, Jiaqi Li^17,18^, Chong Li^23,24^, Jiadong Lin^4^, Mark Loftus^3^, Glennis A. Logsdon^16^, Tobias Marschall^1,2^, Gianni V. Martino^20^, Ryan E. Mills^25^, Yulia Mostovoy^26^, Katherine M. Munson^4^, Giuseppe Narzisi^6^, Lingbin Ni^4^, Keisuke K. Oshima^16^, Carolyn Paisie^3^, Samarendra Pani^1,2^, Zishan Peng^12^, David Porubsky^4^, Timofey Prodanov^1,2^, Keon Rabbani^11^, Tobias Rausch^19^, Xinghua Shi^23,24^, Yuwei Song^12^, Kaitlyn Sun^4^, Likhitha Surapaneni^5^, Michael E. Talkowski^27-29^, Vasiliki Tsapalou^19^, Andres Veidenberg^5^, Feyza Yilmaz^3^, DongAhn Yoo^4^, Xuefang Zhao^27-29^, Weichen Zhou^25^, Qihui Zhu^3^, Michael C. Zody^6^

1. Institute for Medical Biometry and Bioinformatics, Medical Faculty and University Hospital Düsseldorf, Heinrich Heine University, Düsseldorf, Germany
2. Center for Digital Medicine, Heinrich Heine University, Düsseldorf, Germany
3. The Jackson Laboratory for Genomic Medicine, Farmington, CT, USA
4. University of Washington School of Medicine, Department of Genome Sciences, Seattle, WA, USA
5. European Molecular Biology Laboratory, Wellcome Genome Campus, European Bioinformatics Institute, Cambridge, UK
6. New York Genome Center, New York, NY, USA
7. The University of Connecticut Health Center, Farmington, CT, USA
8. Department of Genetics, University Medical Center Groningen, University of Groningen, Groningen, The Netherlands
9. Oncode Institute, Utrecht, The Netherlands
10. Division of Computational Genomics and Systems Genetics, German Cancer Research Center (DKFZ), Heidelberg, Germany
11. Department of Quantitative and Computational Biology, University of Southern California, Los Angeles, CA, USA
12. Department of Biomedical Informatics and Data Science, Heersink School of Medicine, University of Alabama, Birmingham, AL, USA
13. Institute for Genome Sciences, University of Maryland School of Medicine, Baltimore, MD, USA
14. Core Unit Bioinformatics, Medical Faculty and University Hospital Düsseldorf, Heinrich Heine University, Düsseldorf, Germany
15. Howard Hughes Medical Institute, University of Washington, Seattle, WA, USA
16. Department of Genetics, Epigenetics Institute, Perelman School of Medicine, University of Pennsylvania, Philadelphia, PA, USA
17. Department of Molecular Biophysics and Biochemistry, Yale University, New Haven, CT, USA
18. Program in Computational Biology and Bioinformatics, Yale University, New Haven, CT, USA
19. European Molecular Biology Laboratory (EMBL), Genome Biology Unit, Heidelberg, Germany
20. Clemson University, Department of Genetics & Biochemistry, Clemson, SC, USA
21. Institute for Human Genetics, Clemson University, Greenwood, SC, USA
22. Terry Fox Laboratory, BC Cancer Agency, Vancouver, BC, Canada
23. Department of Computer and Information Sciences, Temple University, Philadelphia, PA, USA
24. Institute for Genomics and Evolutionary Medicine, Temple University, Philadelphia, PA, USA
25. Department of Computational Medicine & Bioinformatics, University of Michigan, MI, USA
26. Cardiovascular Research Institute and Institute for Human Genetics, UCSF School of Medicine, CA, USA
27. Program in Medical and Population Genetics, Broad Institute of MIT and Harvard, Cambridge, MA, USA
28. Center for Genomic Medicine, Massachusetts General Hospital, Boston, MA, USA
29. Department of Neurology, Massachusetts General Hospital and Harvard Medical School, MA, USA

## HUMAN PANGENOME REFERENCE CONSORTIUM AUTHORS

Ahmad Abou Tayoun^1,2^, Derek Albracht^3^, Jamie Allen^4^, Alawi A. Alsheikh-Ali^5^, Casey Andrews^6^, Dmitry Antipov^7^, Lucinda Antonacci-Fulton^3^, Mobin Asri^8^, Marcelo Ayllon^9^, Jennifer R. Balacco^10^, Floris P. Barthel^11^, Edward A. Belter Jr.^3^, Halle D. Bender^8^, Andrew P. Blair^8^, Silvia Buonaiuto^12^, Davide Bolognini^13^, Katherine E. Bonini^14^, Christina Boucher^15^, Guillaume Bourque^16,17,18^, Shuo Cao^12^, Andrew Carroll^19^, Ann M. Mc Cartney^8^, Monika Cechova^8^, Pi-Chuan Chang^19^, Xian Chang^8^, Jitender Cheema^4^, Haoyu Cheng^20^, Claudio Ciofi^21^, Sarah Cody^3^, Vincenza Colonna^12^, Holland C. Conwell^22^, Robert Cook-Deegan^23^, Mark Diekhans^8^, Maria Angela Diroma^21^, Daniel Doerr^24,25,26^, Zheng Dong^6^, Richard Durbin^27,28^, Jana Ebler^24,29^, Evan E Eichler^9,30^, Jordan M. Eizenga^8^, Parsa Eskandar^8^, Eddie Ferro^15^, Anna-Sophie Fiston-Lavier^31,32^, Sarah M. Ford^22^, Willard W. Ford^33^, Giulio Formenti^10^, Adam Frankish^4^, Mallory A. Freeberg^4^, Qichen Fu^6^, Stephanie M. Fullerton^34^, Robert S. Fulton^3^, Yan Gao^35^, Gage H. Garcia^9^, Obed A. Garcia^36^, Joshua M.V. Gardner^8^, Shilpa Garg^37^, Erik Garrison^12^, Nanibaa’ A. Garrison^38,39,40^, John Garza^3^, Margarita Geleta^8^, Mohammadmersad Ghorbani^41^, Tina Graves-Lindsay^3^, Richard E. Green^22^, Cristian Groza^42^, Andrea Guarracino^12^, Melissa Gymrek^33^, Leanne Haggerty^4^, Ira M. Hall^43,44^, Nancy F. Hansen^7^, Mohammad Amiruddin Hashmi^5^, Maximilian Haeussler^8^, Yue Hao^11^, David Haussler^8^, Prajna Hebbar^8^, Peter Heringer^24,25,26^, Glenn Hickey^8^, Todd L. Hillaker^8^, S. Nakib Hossain^4^, Neng Huang^35,45^, Sarah E. Hunt^4^, Toby Hunt^4^, Alexander G. Ioannidis^8^, Nafiseh Jafarzadeh^8^, Nivesh Jain^10^, Erich D. Jarvis^10,30^, Maryam Jehangir^11^, Juan Jiang^6^, Jonathan LoTempio Jr^46^, Eimear E. Kenny^14^, Juhyun Kim^7^, Bonhwang Koo^10^, Sergey Koren^7^, Milinn Kremitzki^3,6^, Ben Langmead^47^, Xiaoyu Zhuo^6^, Heather A. Lawson^6^, Daofeng Li^6^, Heng Li^35,45^, Wen-Wei Liao^43,44^, Jiadong Lin^9^, Tianjie Liu^6^, Glennis A. Logsdon^46^, Ryan Lorig-Roach^8^, Hailey Loucks^8^, Jane E. Loveland^4^, Jianguo Lu^48^, Shuangjia Lu^43,44^, Julian K. Lucas^8^, Juan F. Macias-Velasco^3,6,49^, Maximillian G. Marin^35^, Franco L. Marsico^12^, Kateryna D. Makova^50^, Christopher Markovic^6^, Tobias Marschall^24,29^, Fergal J. Martin^4^, Mira Mastoras^8^, Capucine Mayoud^31^, Brandy McNulty^8^, Jack A. Medico^10^, Julian M. Menendez^8^, Karen H. Miga^8^, Anna Minkina^51^, Matthew W. Mitchell^52^, Saswat K. Mohanty^53^, Younes Mokrab^41,54,55^, Jean Monlong^56^, Shabir Moosa^41^, Avelina Moreno-Ochando^57,58^, Shinichi Morishita^59^, Jonathan M. Mudge^4^, Katherine M. Munson^9^, Njagi Mwaniki^60^, Nasna Nassir^5^, Chiara Natali^21^, Shloka Negi^8^, Lingbin Ni^9^, Adam M. Novak^8^, Chie Owa^59^, Sadye Paez^10^, Benedict Paten^8^, Hiram Clawson^8^, Clelia Peano^13,61^, Adam M. Phillippy^7^, Brandon D. Pickett^7^, Laura Pignata^12^, Nadia Pisanti^60^, David Porubsky^9^, Pjotr Prins^12^, Anandi Radhakrishnan^8^, T. Rhyker Ranallo-Benavidez^11^, Brian J. Raney^8^, Mikko Rautiainen^62^, Alessandro Raveane^13^, Luyao Ren^9,30^, Arang Rhie^7^, Farnaz Salehi^12^, Samuel Sacco^22^, Michael C. Schatz^47,63^, Laura B. Scheinfeldt^52^, Aarushi Sehgal^33^, William E. Seligmann^22^, Mahsa Shabani^64^, Kishwar Shafin^19^, Shadi Shahatit^31^, Ruhollah Shemirani^14^, Vikram S. Shivakumar^47^, Swati Sinha^4^, Jouni Sirén^8^, Linnéa Smeds^53^, Steven J. Solar^7^, Marco Sollitto^10,21^, Nicole Soranzo^13,27,28^, Andrew B. Stergachis^9,51^, Marie-Marthe Suner^4^, Yoshihiko Suzuki^59^, Arda Söylev^24,29^, Jack A.S. Tierney^4^, Chad Tomlinson^3^, Francesca Floriana Tricomi^4^, Mohammed Uddin^5,65^, Matteo Tommaso Ungaro^22,66^, Rahul Varki^15^, Flavia Villani^12^, Mitchell R. Vollger^51^, Brian P. Walenz^7^, Charles Wang^67^, Lisa E. Wang^14^, Ting Wang^3,6,49^, Aaron M. Wenger^68^, Conor V. Whelan^10^, Zilan Xin^6^, Zheng Xu^6^, Kai Ye^69^, DongAhn Yoo^9^, Wenjin Zhang^6^, Ying Zhou^35^, Ivo Violich^8^, Giulia Zunino^13^

1. Center for Genomic Discovery, Mohammed Bin Rashid University, Dubai Health, UAE
2. Dubai Health Genomic Medicine Center, Dubai Health, UAE
3. McDonnell Genome Institute, Washington University School of Medicine, St. Louis, MO 63108, USA
4. European Molecular Biology Laboratory, European Bioinformatics Institute (EMBL-EBI), Wellcome Genome Campus, Hinxton, Cambridge CB10 1SD, UK
5. Center for Applied and Translational Genomics (CATG), Mohammed Bin Rashid University of Medicine and Health Sciences, Dubai, United Arab Emirates
6. Department of Genetics, Washington University School of Medicine, St. Louis, MO 63110, USA
7. Genome Informatics Section, Center for Genomics and Data Science Research, National Human Genome Research Institute, National Institutes of Health, Bethesda, MD 20894, USA
8. UC Santa Cruz Genomics Institute, University of California, Santa Cruz, 2300 Delaware Avenue, Santa Cruz, CA 95060, USA
9. Department of Genome Sciences, University of Washington School of Medicine, Seattle, WA 98195, USA
10. The Vertebrate Genome Laboratory, The Rockefeller University, 1230 York Ave, NY 10065, USA
11. The Translational Genomics Research Institute (TGen), 445 N 5th St 4th Floor, Phoenix, AZ 85004, USA
12. Department of Genetics, Genomics and Informatics, University of Tennessee Health Science Center, 71 S Manassas St, Memphis, TN 38163, USA
13. Human Technopole, Milan, Italy
14. Institute for Genomic Health, Icahn School of Medicine at Mount Sinai, New York, NY 10029, USA
15. Department of Computer and Information Science and Engineering, University of Florida, Gainesville, FL 32611, USA
16. Canadian Center for Computational Genomics, McGill University, Montréal, QC, Canada
17. Department of Human Genetics, McGill University, Montréal, QC, Canada
18. Victor Phillip Dahdaleh Institute of Genomic Medicine, Montréal, QC, Canada
19. Google LLC, 1600 Amphitheatre Pkwy, Mountain View, CA 94043, USA
20. Department of Biomedical Informatics and Data Science, Yale School of Medicine, New Haven, CT 06510, USA
21. University of Florence, Department of Biology, Via Madonna del Piano, 6, Sesto Fiorentino (FI) 50019, Italy
22. Department of Ecology and Evolutionary Biology, University of California, Santa Cruz, Santa Cruz, CA 95064, USA
23. Arizona State University, Consortium for Science, Policy & Outcomes, 1800 I St NW, Washington, DC 20006, USA
24. Center for Digital Medicine, Heinrich Heine University Düsseldorf, Germany
25. Department for Endocrinology and Diabetology, Medical Faculty and University Hospital Düsseldorf, Heinrich Heine University Düsseldorf, Germany
26. German Diabetes Center (DDZ), Leibniz Institute for Diabetes Research, Düsseldorf, Germany
27. University of Cambridge, Cambridge, UK
28. Wellcome Sanger Institute, Genome Campus, Hinxton, CB10 1HH, UK
29. Institute for Medical Biometry and Bioinformatics, Medical Faculty and University Hospital Düsseldorf, Heinrich Heine University, Düsseldorf, Germany
30. Howard Hughes Medical Institute, Chevy Chase, MD 20815, USA
31. ISEM, Univ Montpellier, CNRS, IRD, Montpellier, France
32. Institut Universitaire de France, Paris, France
33. Department of Computer Science and Engineering, University of California, San Diego, La Jolla, CA 92093, USA
34. Department of Bioethics & Humanities, University of Washington School of Medicine, Seattle, WA 98195, USA
35. Department of Data Science, Dana-Farber Cancer Institute, MA 02215, USA
36. Department of Anthropology, University of Kansas, KS 66044 USA
37. University of Manchester, Manchester M13 9PL, UK
38. Traditional, ancestral and unceded territory of the Gabrielino/Tongva peoples, Institute for Society & Genetics, University of California, Los Angeles, Los Angeles, CA 90095, USA
39. Traditional, ancestral and unceded territory of the Gabrielino/Tongva peoples, Institute for Precision Health, David Geffen School of Medicine, University of California, Los Angeles, Los Angeles, CA 90095, USA
40. Traditional, ancestral and unceded territory of the Gabrielino/Tongva peoples, Division of General Internal Medicine & Health Services Research, David Geffen School of Medicine, University of California, Los Angeles, Los Angeles, CA 90095, USA
41. Medical and Population Genomics Lab, Sidra Medicine, Doha, Qatar
42. Montreal Heart Institute, Montreal, Quebec, Canada
43. Center for Genomic Health, Yale University School of Medicine, New Haven, CT 06510, USA
44. Department of Genetics, Yale University School of Medicine, New Haven, CT 06510, USA
45. Department of Biomedical Informatics, Harvard Medical School, MA 02115, USA
46. Department of Genetics, Epigenetics Institute, Perelman School of Medicine, University of Pennsylvania, Philadelphia, PA 19104, USA
47. Department of Computer Science, Johns Hopkins University, Baltimore, MD 21218, USA
48. Sun Yat-sen University, No. 135, Xingang Xi Road, Guangzhou, China
49. Edison Family Center for Genome Sciences & Systems Biology, Washington University School of Medicine, St. Louis, MO 63110, USA
50. Center for Medical Genomics, Penn State University, University Park, PA 16802, USA
51. Division of Medical Genetics, Department of Medicine, University of Washington School of Medicine, Seattle, WA 98195, USA
52. Coriell Institute for Medical Research, Camden, NJ 08103, USA
53. Department of Biology, Penn State University, University Park, PA 16802, USA
54. Department of Biomedical Science, College of Health Sciences, Qatar University, Doha, Qatar
55. Department of Genetic Medicine, Weill Cornell Medicine-Qatar, Doha, Qatar
56. IRSD - Digestive Health Research Institute, University of Toulouse, INSERM, INRAE, ENVT, UPS, Toulouse, France
57. MATCH biosystems, S.L., Spain
58. Universidad Miguel Hernández de Elche, Spain
59. Department of Computational Biology and Medical Sciences, The University of Tokyo, Kashiwa, Chiba 277-8561, Japan
60. University of Pisa, Pisa, Italy
61. Institute of Genetics and Biomedical Research, UoS of Milan, National Research Council, Milan, Italy
62. Institute for Molecular Medicine Finland, Helsinki Institute of Life Science, University
63. Department of Biology, Johns Hopkins University, Baltimore, MD 21218, USA
64. University of Amsterdam, Amsterdam, Netherlands
65. GenomeArc Inc, Mississauga, ON, Canada
66. Department of Biology and Biotechnologies “Charles Darwin”, University of Rome “La Sapienza”, 00185 Rome, Italy
67. Center for Genomics, Loma Linda University School of Medicine, Loma Linda, CA 92350, USA
68. PacBio, 1305 O’Brien Drive, Menlo Park, CA 94025, USA
69. The first affiliated hospital of Xi’an Jiaotong University, Xi’an Jiaotong University, Xi’an, Shaanxi, 710049, China

