## Supplementary Information for "A global view of human centromere variation and evolution"

**This PDF file includes:**

1. **Supplementary Notes 1 and 2**
2. **Supplementary Figures 1-77**
3. **Supplementary Tables 1-21**
4. **Supplementary References**

### SUPPLEMENTARY NOTES

**Supplementary Note 1.** To confirm the accurate assembly of each human centromere, we assessed each of them with a suite of computational tools and approaches. First, we aligned native long-read Pacific Biosciences high-fidelity (PacBio HiFi) and Oxford Nanopore Technologies (ONT) sequencing data generated from each genome to the corresponding genome assembly and assessed for uniform read depth across each centromere. We found that all centromeres had uniform coverage of both PacBio HiFi and ONT data, suggesting that they were free of large structural errors (**Supplementary Table 2**). Then, we applied HMM-Flagger[1], a tool that uses a Hidden Markov Model (HMM) to detect anomalies in coverage based on PacBio HiFi alignments, and found that nearly all α-satellite sequences within the centromeres were free of assembly errors (**Supplementary Table 2**). Next, we ran NucFlag[2], a tool that assesses for assembly errors based on first and second most common bases in aligned PacBio HiFi or ONT reads, and found that 100% of centromeres were free of such errors (**Supplementary Table 2**). Then, we applied GAVISUNK[3], a tool that compares singly unique nucleotide *k*-mers (SUNKs) within the assembly to those found in native ONT reads, and found that 98% of α-satellite sequences were supported with raw ONT data, with the remaining 2% due to validation gaps[3] (**Supplementary Table 2**). Finally, we estimated the quality value (QV) of the centromeres with Merqury[4] using short-read Illumina sequencing data generated from the same source genome and found them to have an average QV of 60.99, indicating that the centromeres were >99.9999% accurate on average (**Supplementary Table 3**).

**Supplementary Note 2.** A third of all chromosomes contain non-kinetochore-forming arrays that flank the larger kinetochore-forming array(s) (chromosomes 3, 4, 10, 17-21). These small α-satellite HOR arrays are present at all centromeres from chromosomes 3, 4, and 10, but are only present at 54%, 2%, 1%, 4%, and 7% of centromeres from chromosomes 17-21, respectively. The mean length of the non-kinetochore-forming arrays is 332 kbp, with the smallest on chromosome 10 (1.7 kbp) and the largest on chromosome 12 due to a large duplication (3.4 Mbp; **Fig. 2a**, **Supplementary Fig. 7a**, **Supplementary Table 6**).

### SUPPLEMENTARY FIGURES

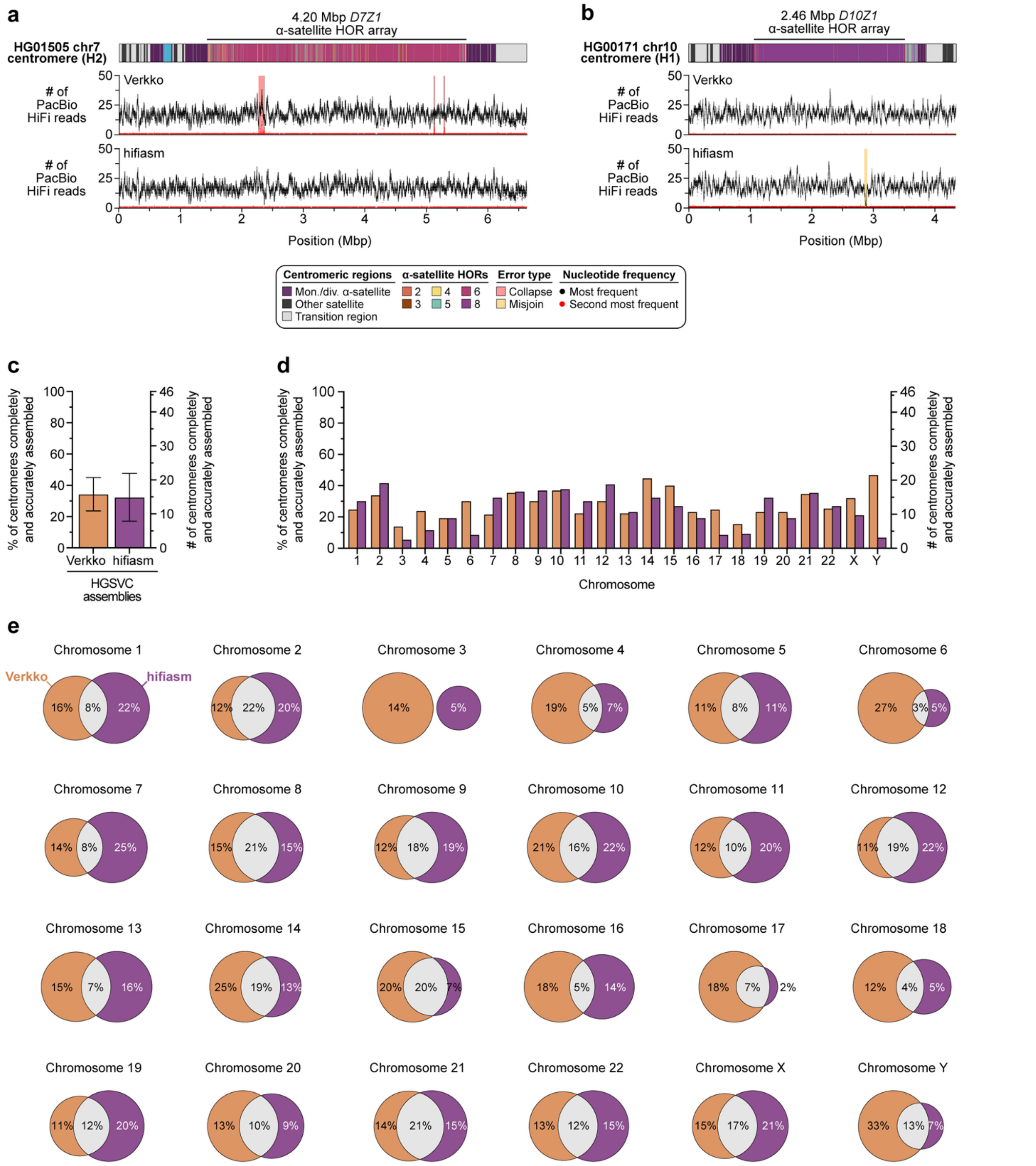


**Supplementary Figure 1. Centromeres assembled with Verkko and hifiasm have complementary error profiles. a,b)** Plots showing **a)** the HG01505 chromosome 7 centromere (haplotype 2; H2) or **b)** the HG00171 chromosome 10 centromere (haplotype 1; H1) assembled with either Verkko or hifiasm and the different assembly errors detected in each. **c,d)** Quantification of the percentage and number of centromeres completely and accurately assembled in Verkko- or hifiasm-assembled genomes for **b)** all or **c)** individual chromosomes. **e)** Venn diagrams showing the percentage of centromeres completely and accurately assembled by Verkko, hifiasm, or both for each chromosome.

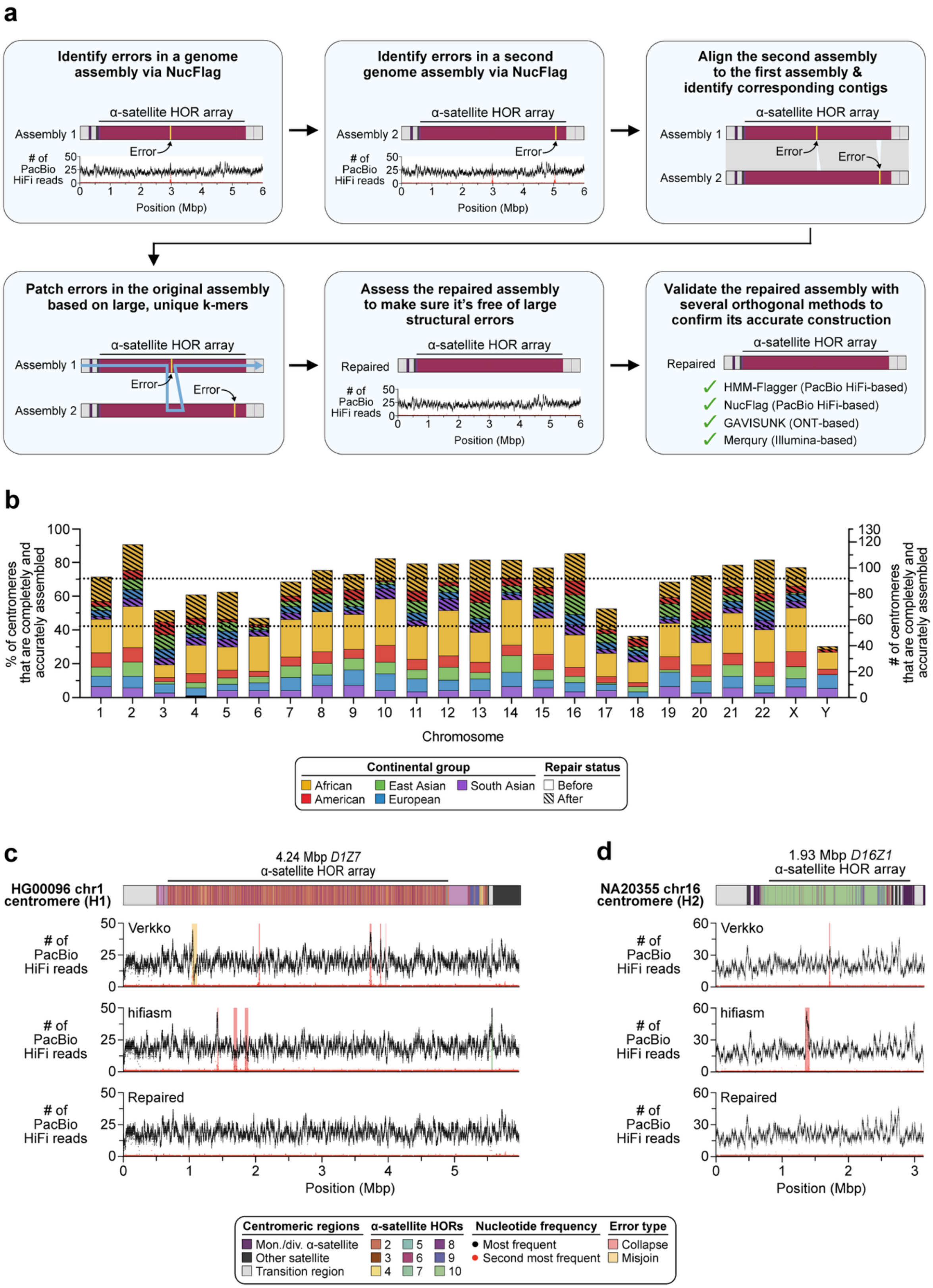


**Supplementary Figure 2. AssemblyRepairer successfully resolves errors in human centromeres, increasing the percentage of accurately resolved centromeres from 42.2% to 70.4%. a)** Method for resolving errors in genome assemblies via AssemblyRepairer, which uses two complementary assemblies generated from the same genome to patch errors in one of them. **b)** Bar plot showing the increase in completely and accurately assembled centromeres after repair. Before repair, 54 centromeres per chromosome (42.2%) were accurately resolved on average; after repair, 92 centromeres per chromosome (70.4%) were accurately resolved on average. Chromosome 3 has the greatest increase in the number of correctly assembled centromeres (172% increase), while chromosome Y has the least (18.8%). Chromosome 2 had the highest percentage of centromeres that were correctly assembled overall (92%), while chromosome 18 had the fewest (34%), likely due to its large size (**Fig. 2a**). Dotted lines, mean before and after repair. **c,d)** Plots showing the assembly errors detected in the **c)** HG00096 chromosome 1 centromere (H1) and **d)** NA20355 chromosome 16 centromere (H2) before and after repair. Before repair, there are complementary errors in the Verkko and hifiasm assemblies, which AssemblyRepairer uses to patch the errors in the Verkko assembly, generating an accurately resolved centromere.

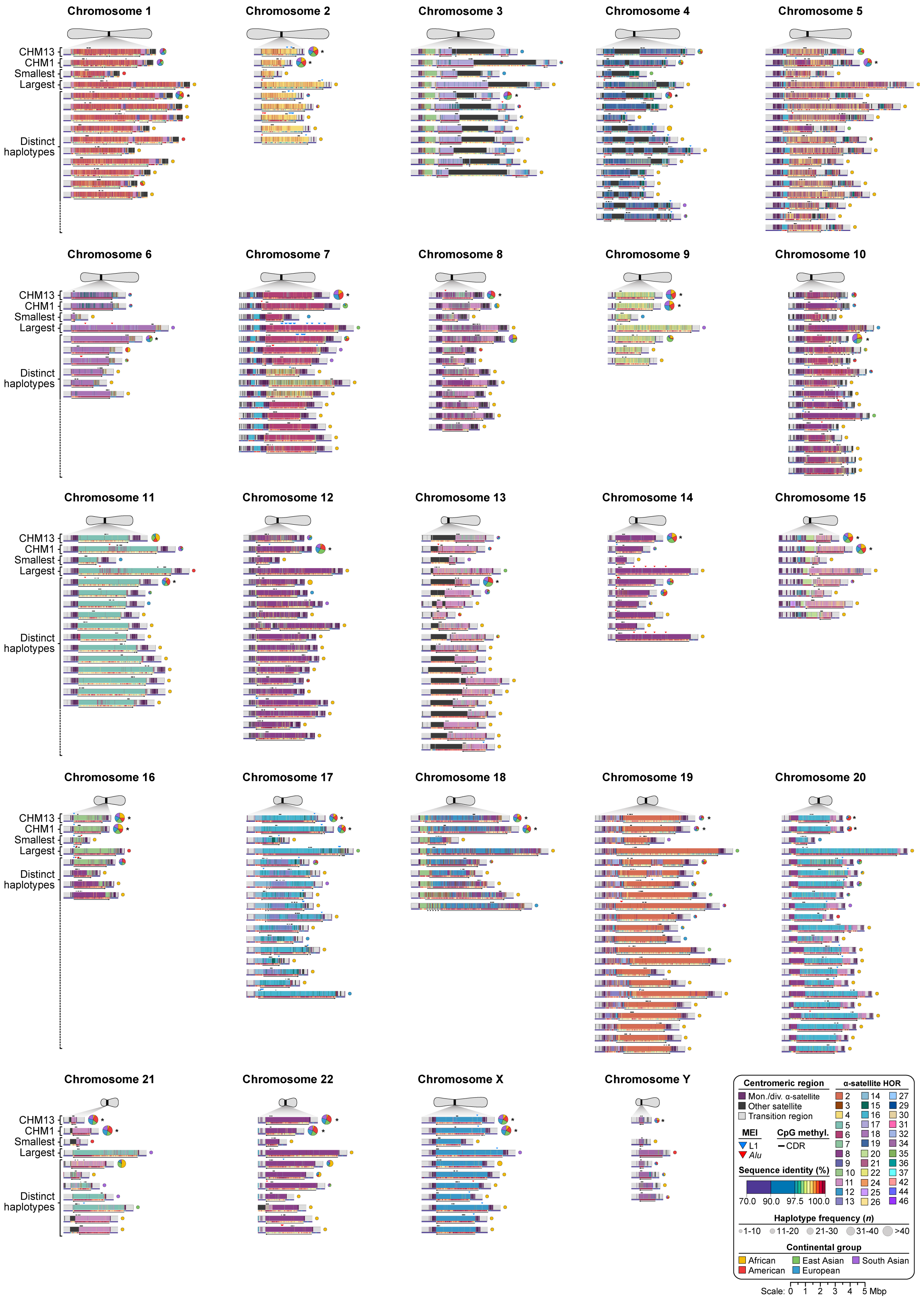


**Supplementary Figure 3. Overview of 226 new major haplotypes and their frequency among 2,156 complete human centromeres.** High-resolution maps showing the sequence, structure, methylation, sequence identity, and mobile element insertions (MEIs) among centromeres from CHM13[5], CHM1[6], and 65 diverse human genomes[7]. The α-satellite HORs are colored by the number of α-satellite monomers within them, and the orientation of the active α-satellite HOR arrays are indicated by an arrow. The site of the putative kinetochore, marked by the centromere dip region (CDR), is shown with a black bar. The local sequence identity across each centromeric region is indicated with a 1D heat map, and mobile element insertions (MEIs) within α-satellite HOR arrays are indicated with colored triangles. Haplotype frequencies are shown as pie charts, with the continental group indicated. The most common haplotype for each chromosome is marked with an asterisk. Zoomed-in views of centromeres from each chromosome are shown in **Supplementary Figs. 4-6**.

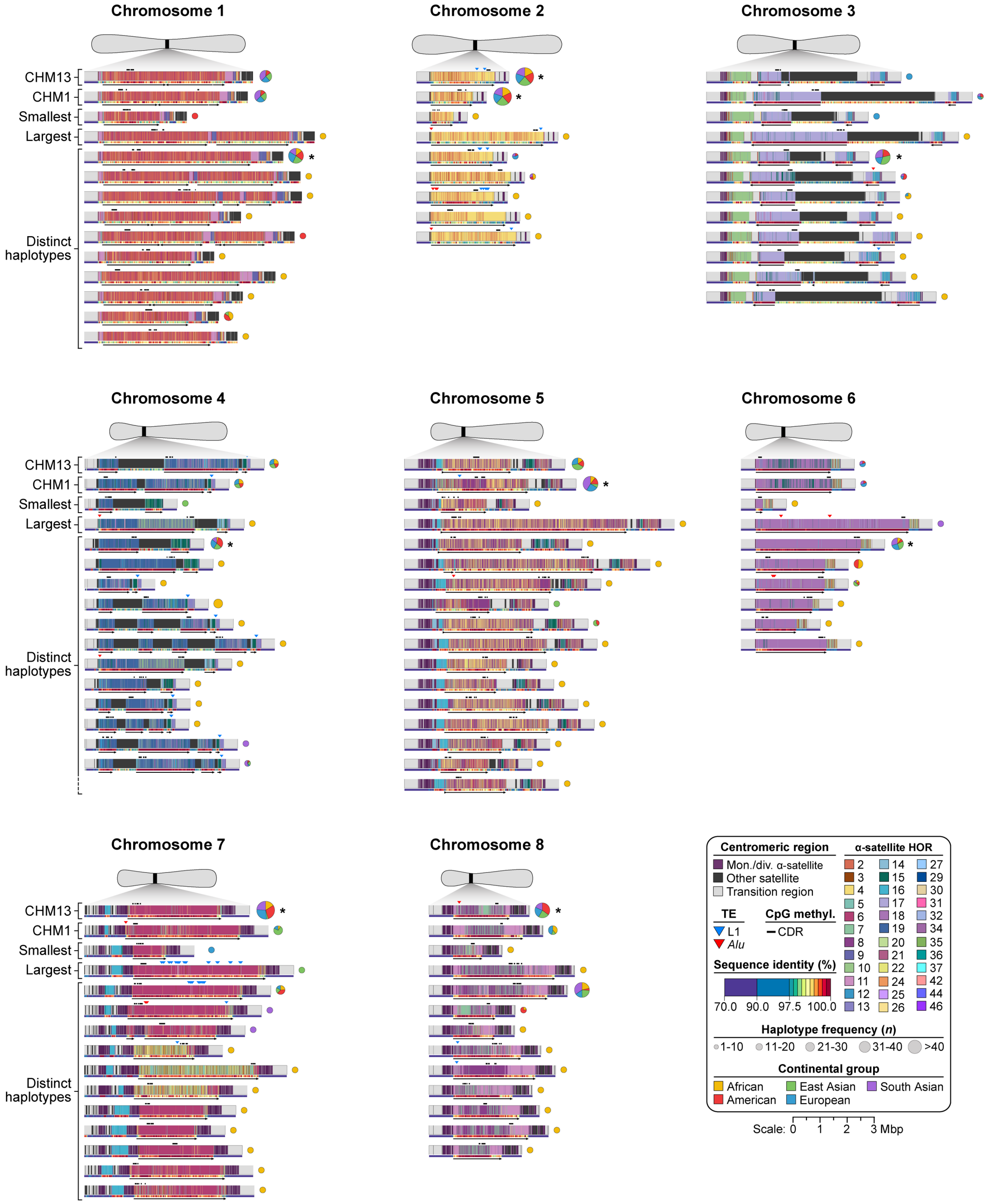


**Supplementary Figure 4. Variation in the sequence, structure, methylation pattern, evolutionary landscape among centromeres from chromosomes 1-8.** High-resolution maps showing the sequence, structure, methylation, sequence identity, and mobile element insertions (MEIs) among centromeres from chromosomes 1-8 in CHM13[5], CHM1[6], and 65 diverse human genomes[7]. The α-satellite HORs are colored by the number of α-satellite monomers within them, and the orientation of the active α-satellite HOR arrays are indicated by an arrow. The site of the putative kinetochore, marked by the CDR, is shown with a black bar. The local sequence identity across each centromeric region is indicated with a 1D heat map, and MEIs within α-satellite HOR arrays are indicated with colored triangles. Haplotype frequencies are shown as pie charts, with the continental group indicated. The most common haplotype for each chromosome is marked with an asterisk.Zoomed-in views of centromeres from chromosomes 9-22, X, and Y are shown in **Supplementary Figs. 5,6**.

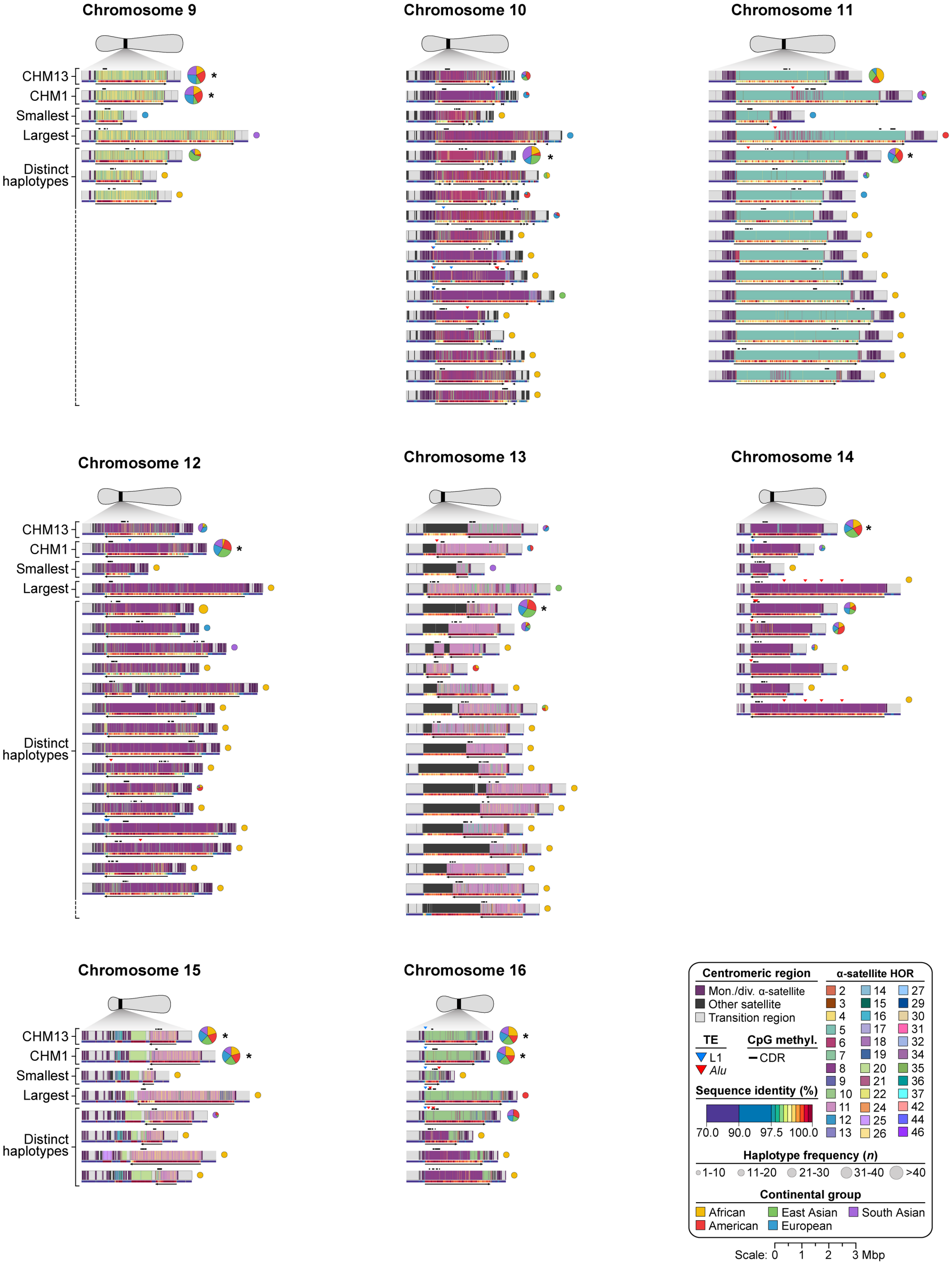


**Supplementary Figure 5. Variation in the sequence, structure, methylation pattern, evolutionary landscape among centromeres from chromosomes 9-16.** High-resolution maps showing the sequence, structure, methylation, sequence identity, and mobile element insertions (MEIs) among centromeres from chromosomes 17-22, X, and Y in CHM13[5], CHM1[6], and 65 diverse human genomes[7]. The α-satellite HORs are colored by the number of α-satellite monomers within them, and the orientation of the active α-satellite HOR arrays are indicated by an arrow. The site of the putative kinetochore, marked by the CDR, is shown with a black bar. The local sequence identity across each centromeric region is indicated with a 1D heat map, and MEIs within α-satellite HOR arrays are indicated with colored triangles. Haplotype frequencies are shown as pie charts, with the continental group indicated. The most common haplotype for each chromosome is marked with an asterisk. Zoomed-in views of centromeres from chromosomes 1-8 and 17-22, X, and Y are shown in **Supplementary Figs. 4,6**, respectively.

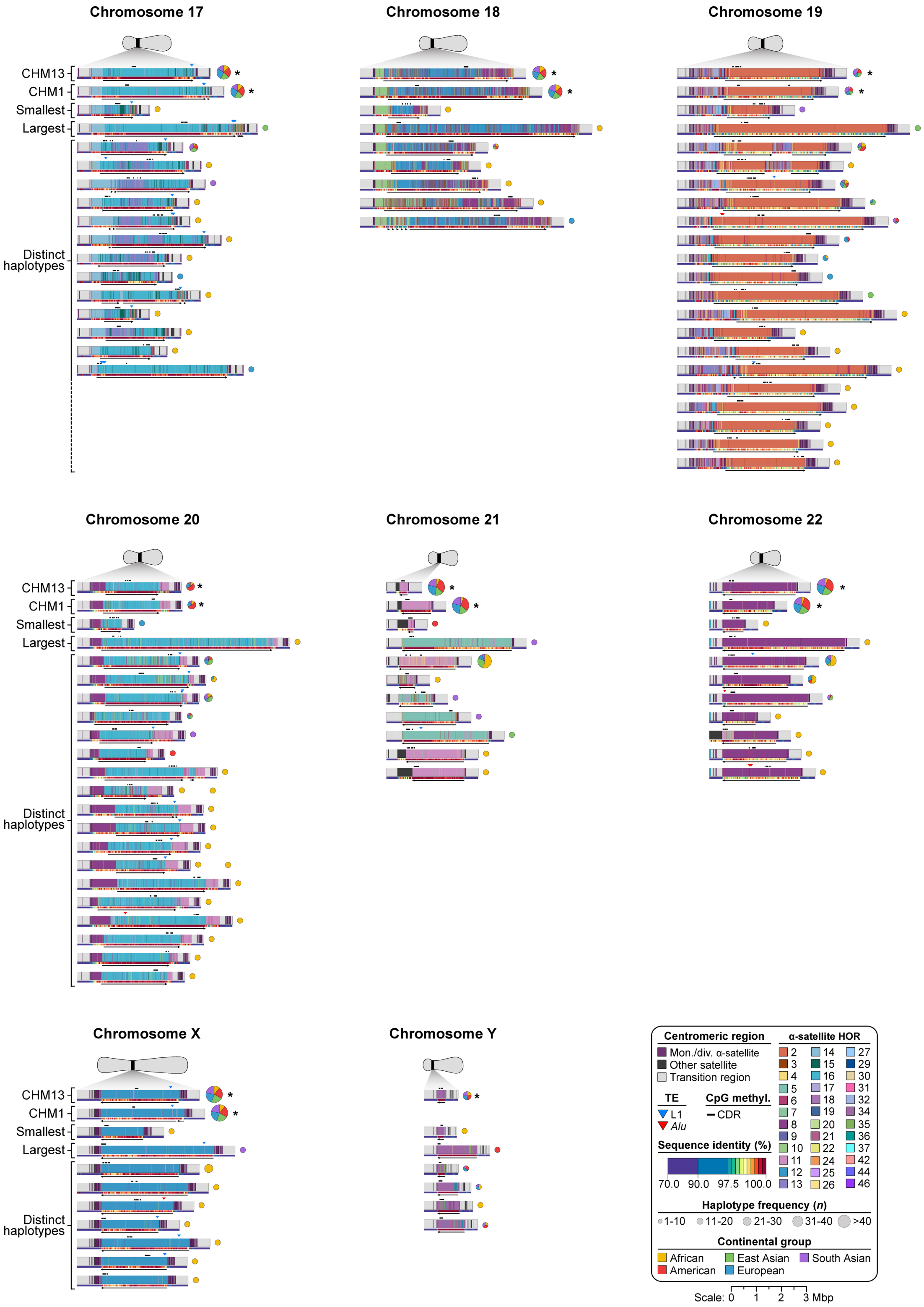


**Supplementary Figure 6. Variation in the sequence, structure, methylation pattern, evolutionary landscape among centromeres from chromosomes 17-22, X, and Y.** High-resolution maps showing the sequence, structure, methylation, sequence identity, and mobile element insertions (MEIs) among centromeres from chromosomes 17-22, X, and Y in CHM13[5], CHM1[6], and 65 diverse human genomes[7]. The α-satellite HORs are colored by the number of α-satellite monomers within them, and the orientation of the active α-satellite HOR arrays are indicated by an arrow. The site of the putative kinetochore, marked by the CDR, is shown with a black bar. The local sequence identity across each centromeric region is indicated with a 1D heat map, and MEIs within α-satellite HOR arrays are indicated with colored triangles. Haplotype frequencies are shown as pie charts, with the continental group indicated. The most common haplotype for each chromosome is marked with an asterisk. Zoomed-in views of centromeres from chromosomes 1-16 are shown in **Supplementary Figs. 4,5**.

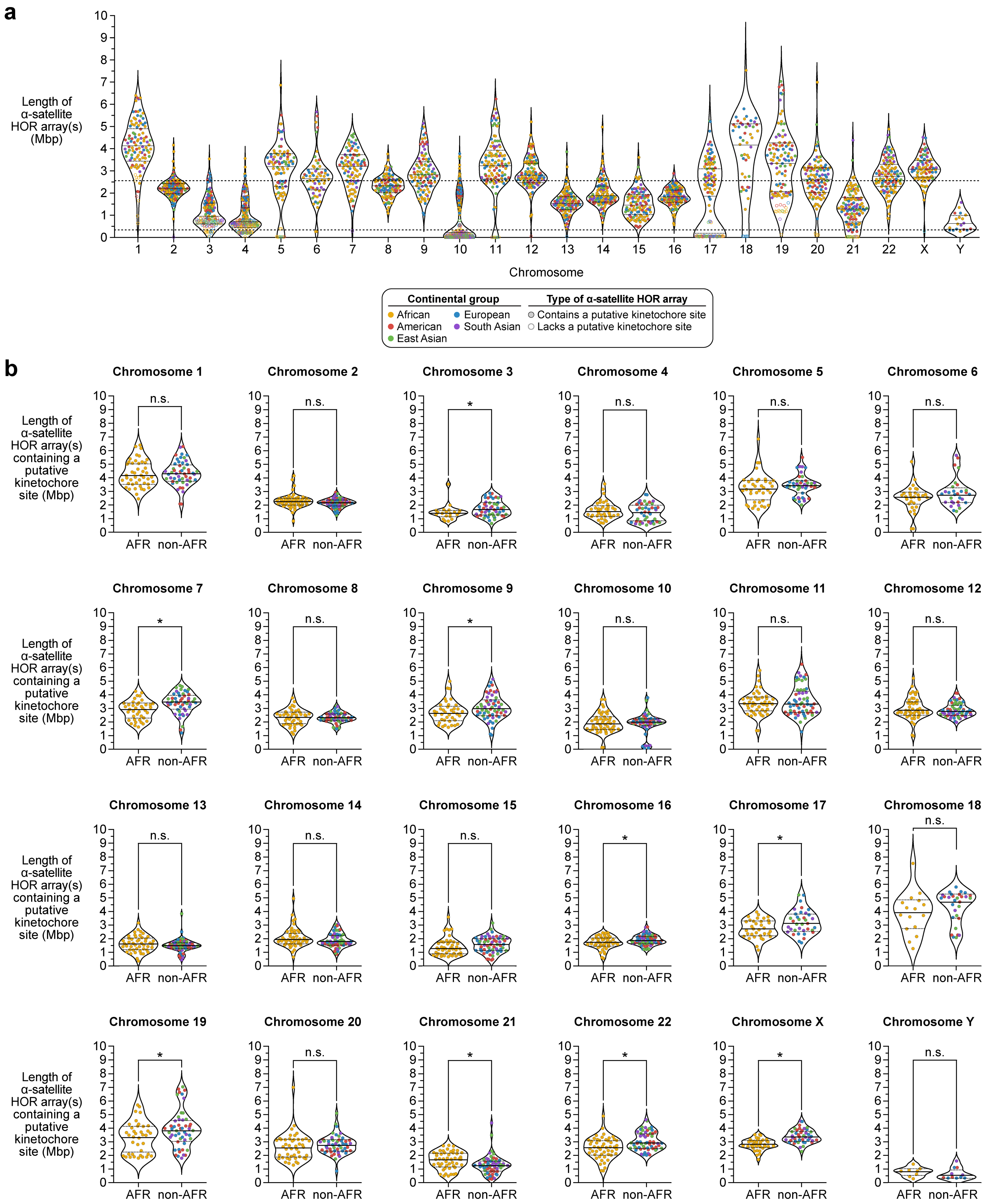


**Supplementary Figure 7. Variation in the length of α-satellite HOR arrays among centromeres. a)** Differences in the length of α-satellite HOR arrays containing or lacking a putative kinetochore site, defined by the presence or absence of the CDR, respectively. For each chromosome, the mean is shown as a solid line, and first and third quartiles are shown as dotted lines. The means across all chromosomes are shown as dashed lines. **b)** Comparison of α-satellite HOR array lengths containing a putative kinetochore site among individuals with African (AFR) or non-African (non-AFR) ancestry. For each ancestry, the mean is shown as a solid line, and first and third quartiles are shown as dotted lines. Nine chromosomes have statistically significant differences in kinetochore-containing α-satellite HOR array lengths between individuals with African (AFR) and non-African (non-AFR) ancestry. For chromosome 21, individuals with African ancestry have α-satellite HOR arrays that are 353 kbp larger, on average, than those with non-African ancestry. In contrast, for chromosomes 3, 7, 9, 16, 17, 19, 22, and X, individuals with non-African ancestry have arrays that are 281, 522, 348, 180, 476, 654, 491, and 594 kbp larger, on average, than those with African ancestry, respectively. For the remaining chromosomes, there is no significant difference, as determined with a two-tailed Wilcoxson rank-sum test. *, $p < 0.05$; n.s., not significant.

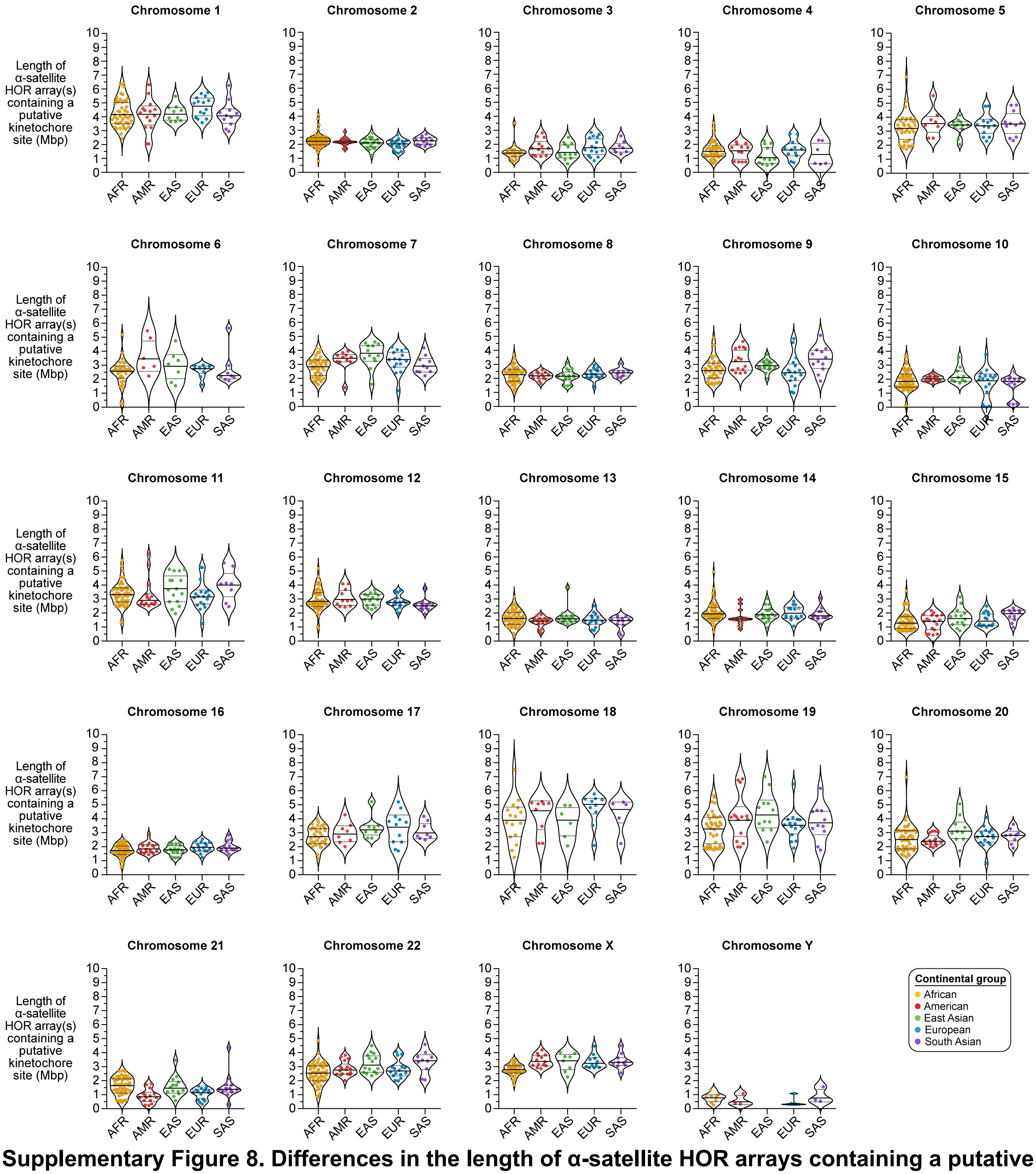


**Supplementary Figure 8. Differences in the length of α-satellite HOR arrays containing a putative kinetochore site among centromeres from individuals with diverse ancestries.** Variation in length of α-satellite HOR arrays containing a putative kinetochore site, defined by the presence of the CDR, among individuals with African (AFR), American (AMR), East Asian (EAS), European (EUR), and South Asian (SAS) ancestry. For each ancestry, the mean is shown as a solid line, and first and third quartiles are shown as dotted lines.

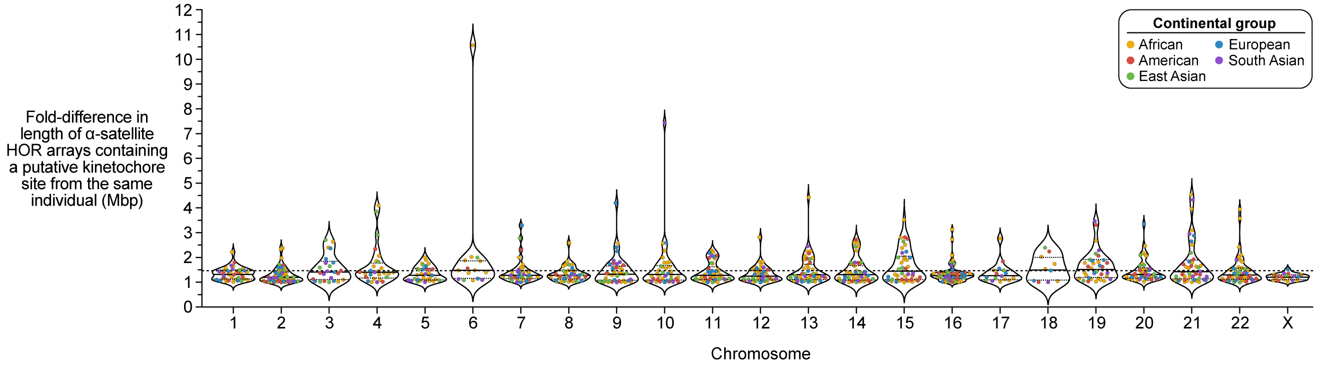


**Supplementary Figure 9. Variation in the length of α-satellite HOR arrays containing a putative kinetochore site between haplotypes of the same individual.** Fold-difference in the length of α-satellite HOR arrays containing a putative kinetochore site, defined by the presence of the CDR, in the same individual. For each chromosome, the mean is shown as a solid line, and first and third quartiles are shown as dotted lines. Mean (1.47-fold) is shown as a dotted line. Only 2.2% of individuals have a >3-fold difference in α-satellite HOR array lengths, and 0.9% have a >4-fold difference.

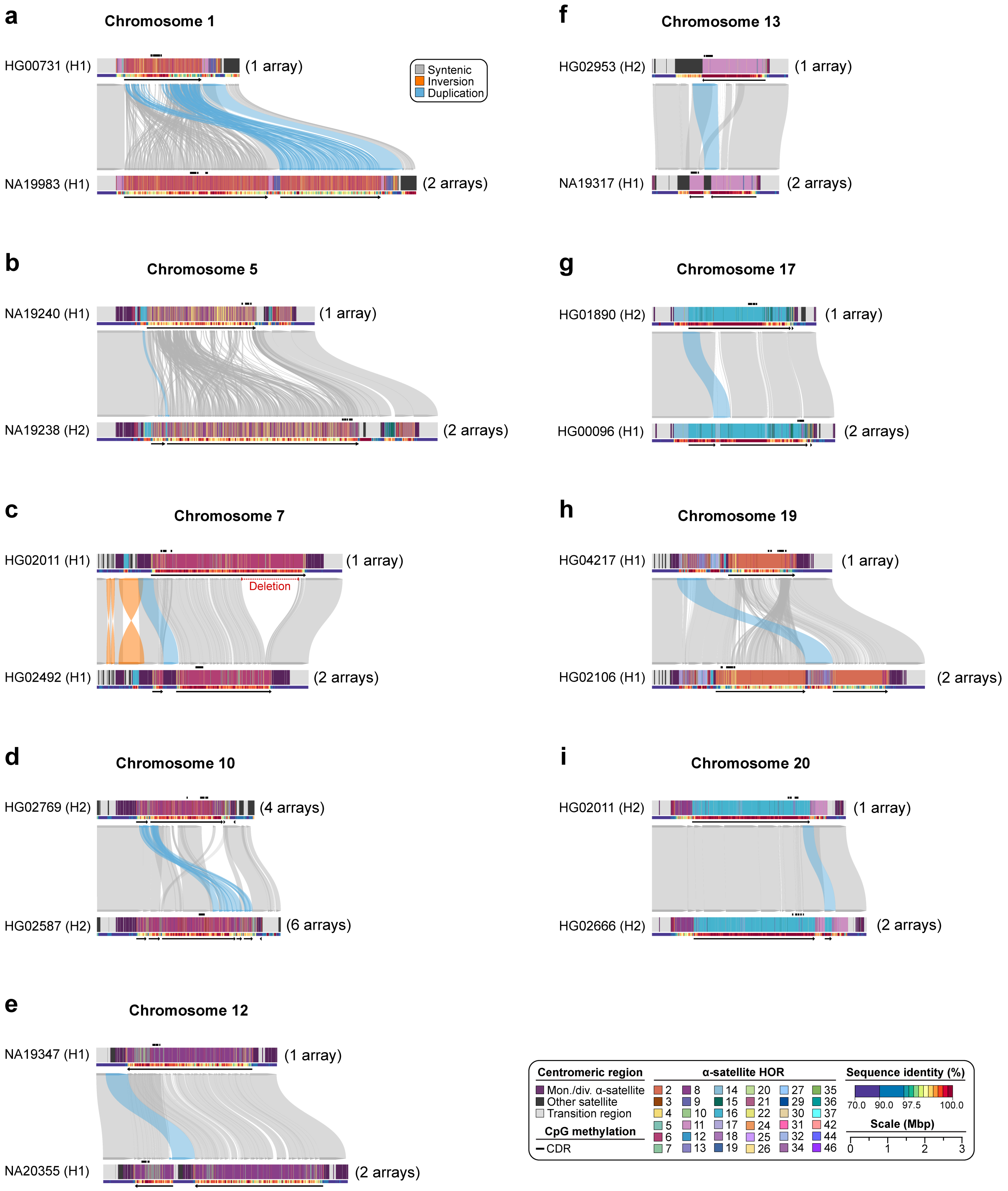


**Supplementary Figure 10. Examples of large-scale structural variation among centromeres from chromosomes 1, 5, 7, 10, 12, 13, 17, 19, and 20. a-i)** Duplications, inversions, and deletions spanning tens to hundreds of kilobase pairs alter the overall structure of centromeres by changing the number and organization of active α-satellite HOR arrays found within chromosomes **a)** 1, **b)** 5, **c)** 7, **d)** 10, **e)** 12, **f)** 13, **g)** 17, **h)** 19, and **i)** 20.

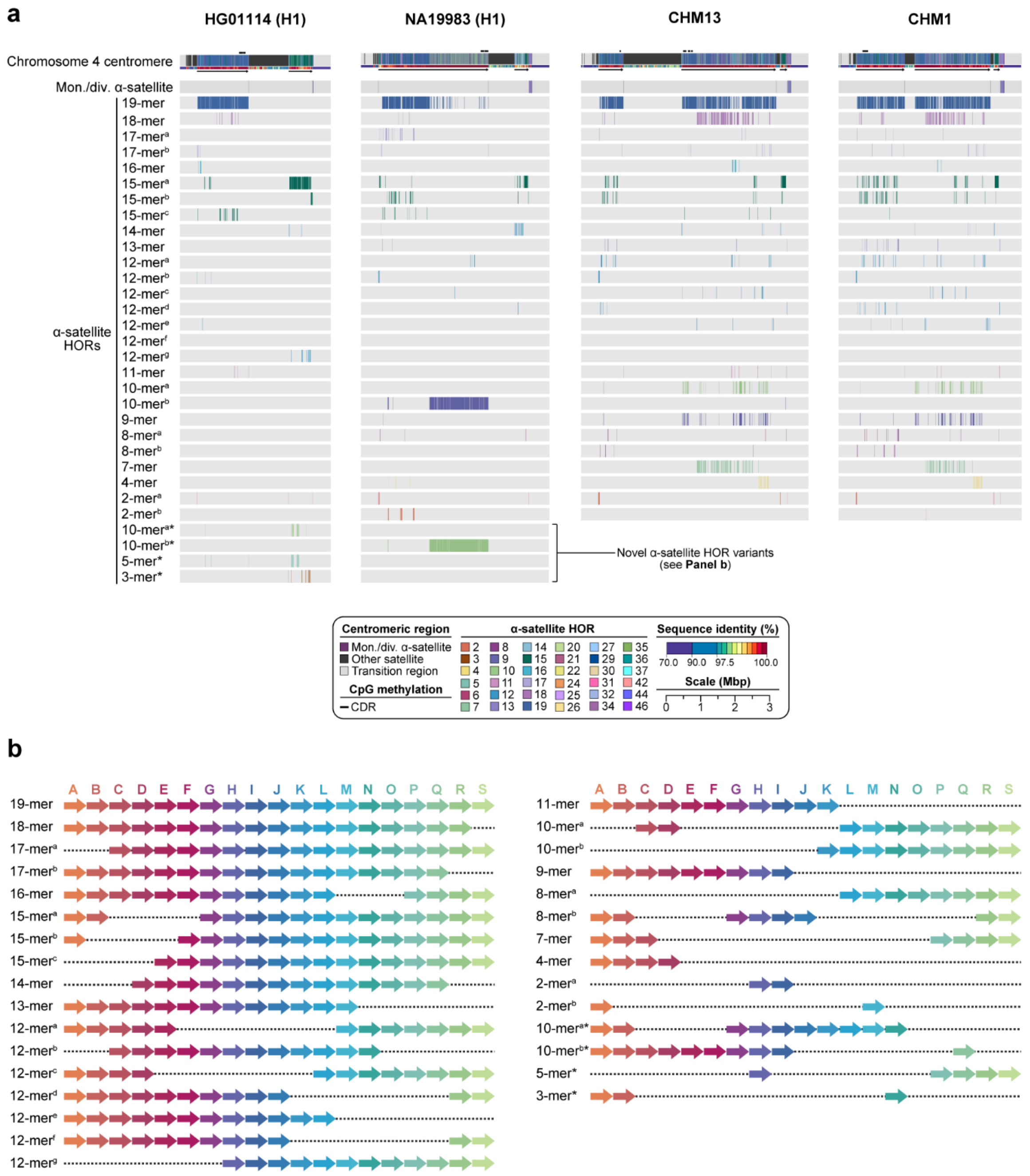


**Supplementary Figure 11. Novel α-satellite HOR variants within chromosome 4 centromeres. a)** Comparison of the HG01114 (H1), NA19983 (H1), CHM13, and CHM1 chromosome 4 *D4Z1* α-satellite HOR arrays, showing that that HG01114 (H1) has three novel α-satellite HOR variants and NA19983 (H1) has one novel α-satellite HOR variant, which alters the overall composition of the α-satellite HOR array. **b)** Structure of the α-satellite HOR variants found in the HG01114 (H1), NA19983 (H1), CHM13, and CHM1 chromosome 4 centromeres. All of them derive from an ancestral 19-mer α-satellite HOR. Novel α-satellite HOR variants are indicated with an asterisk.

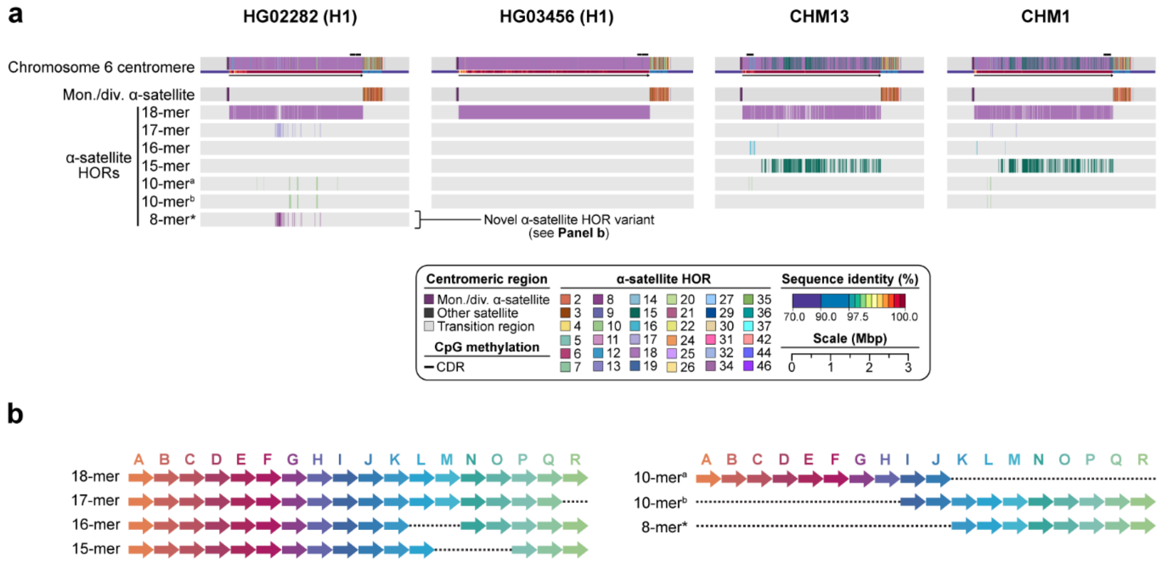


**Supplementary Figure 12. Novel α-satellite HOR variants within chromosome 6 centromeres. a)** Comparison of the HG02282 (H1), HG03456 (H1), CHM13, and CHM1 chromosome 6 *D6Z1* α-satellite HOR arrays, showing that HG02282 (H1) has a novel α-satellite HOR variant and HG03456 (H1) is missing an α-satellite HOR variant, which alters the overall composition of the α-satellite HOR arrays. **b)** Structure of the α-satellite HOR variants found in the HG02282 (H1), HG03456 (H1), CHM13, and CHM1 chromosome 6 centromeres. All of them derive from an ancestral 18-mer α-satellite HOR. The novel 8-mer α-satellite HOR variant is indicated with an asterisk.

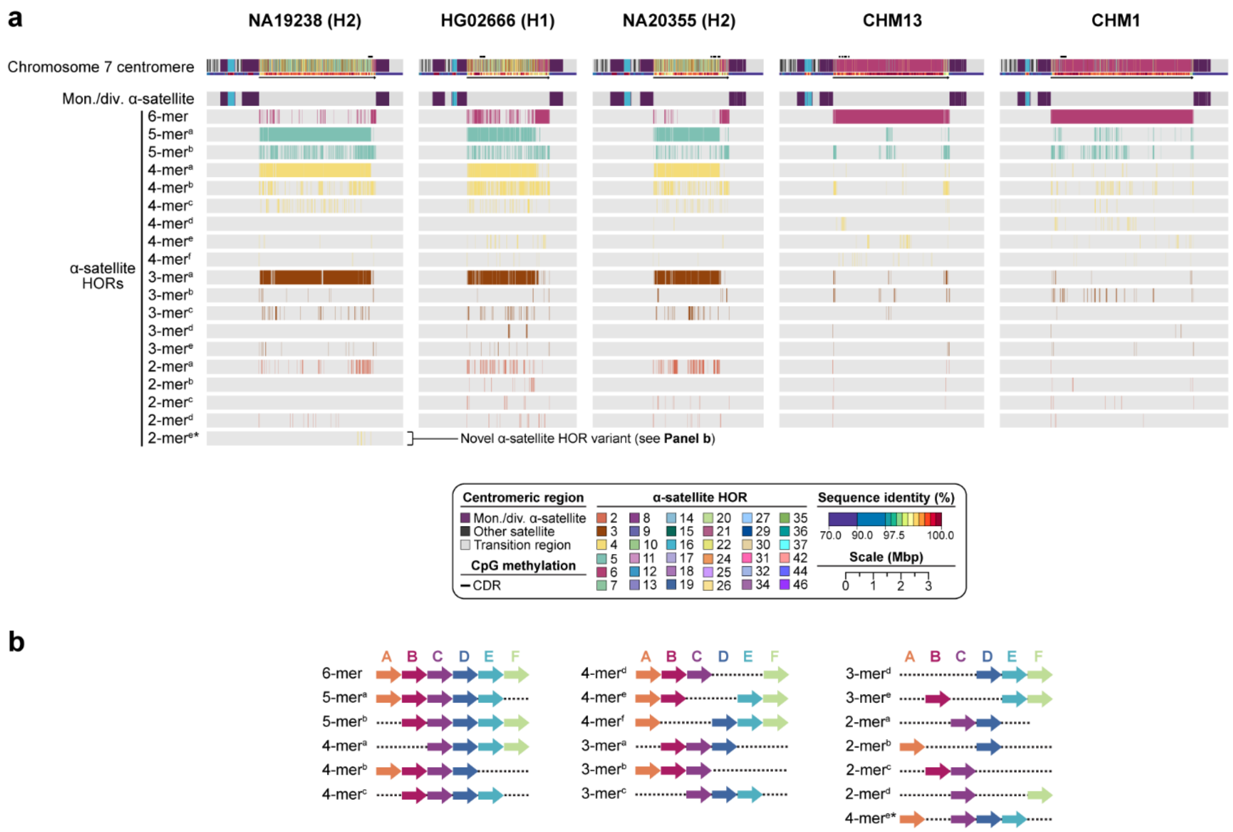


**Supplementary Figure 13. Novel α-satellite HOR variants within chromosome 7 centromeres. a)** Comparison of the NA19238 (H2), HG02666 (H1), NA20355 (H2), CHM13, and CHM1 chromosome 7 *D7Z1* α-satellite HOR arrays, showing that NA19238 (H2) has one novel α-satellite HOR variant and NA19238 (H2), HG02666 (H1), NA20355 (H2) have α-satellite HOR variants in much higher abundance than CHM13 and CHM1, which alters the overall composition of the α-satellite HOR array. **b)** Structure of the α-satellite HOR variants found in the NA20355 (H2), CHM13, and CHM1 chromosome 7 centromeres. All of them derive from an ancestral 6-mer α-satellite HOR. The novel 2-mer α-satellite HOR variant is indicated with an asterisk.

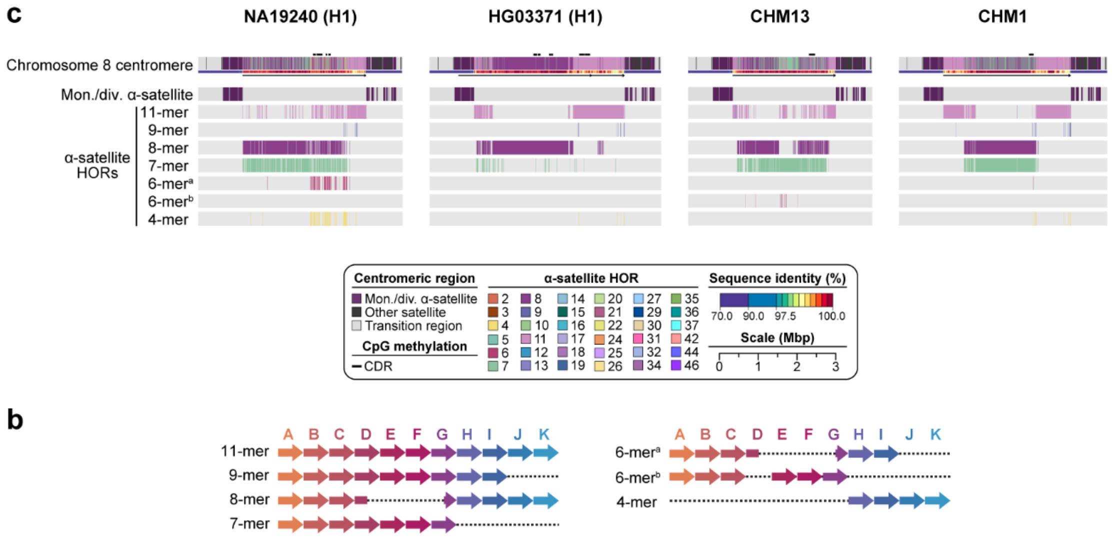


**Supplementary Figure 14. Differences in composition and abundance of α-satellite HOR variants within chromosome 8 centromeres. a)** Comparison of the NA19240 (H1), HG03371 (H1), CHM13, and CHM1 chromosome 8 *D8Z2* α-satellite HOR arrays reveals that NA19240 (H1) has two α-satellite HOR variants in much higher abundance than CHM13 and CHM1 and HG003371 (H1) has an α-satellite HOR variant in lower abundance, which alters the overall composition of the α-satellite HOR arrays. **b)** Structure of the α-satellite HOR variants found in the NA19240 (H1), HG03371 (H1), CHM13, and CHM1 chromosome 8 centromeres. All of them derive from an ancestral 11-mer α-satellite HOR.

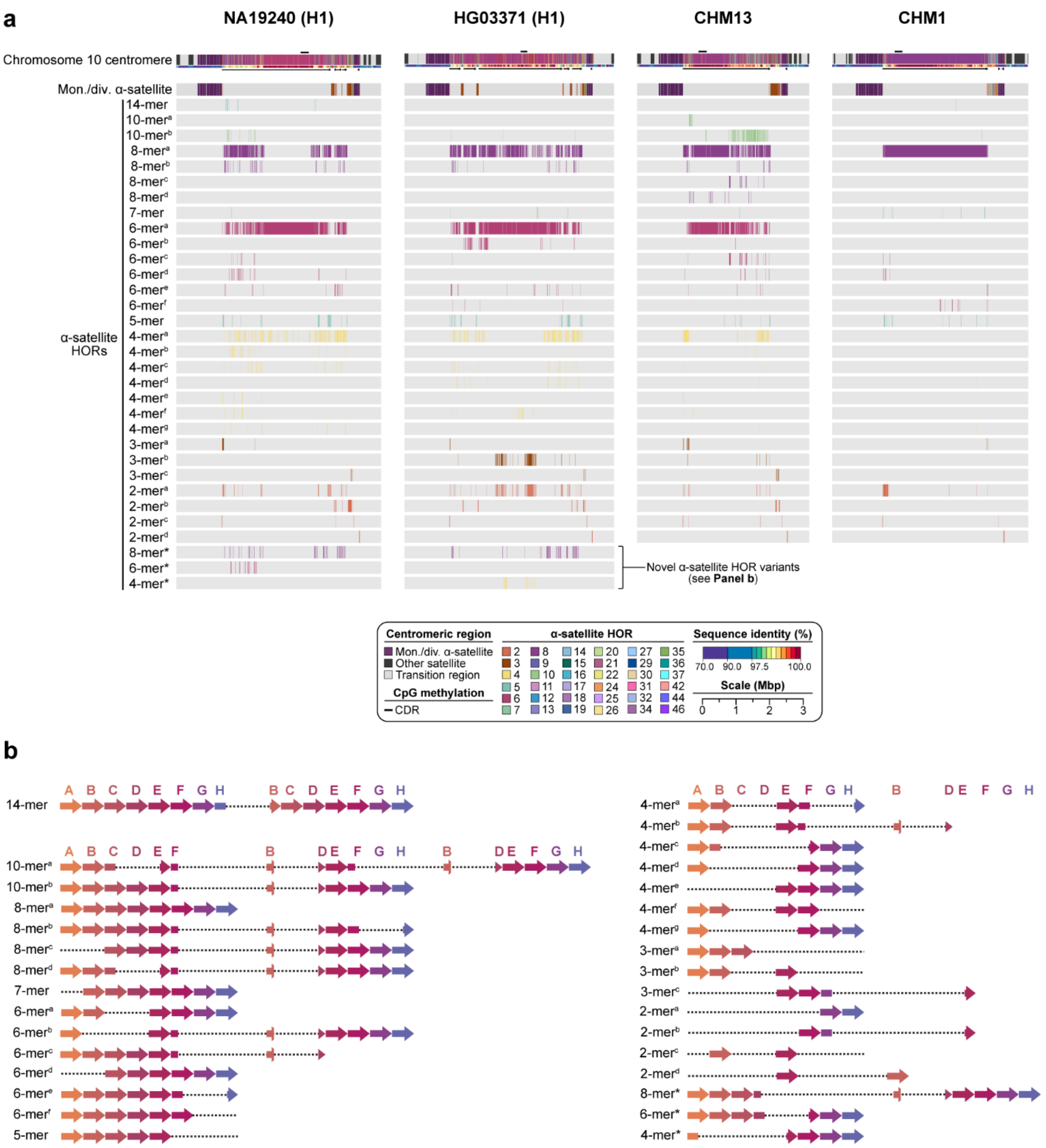


**Supplementary Figure 15. Novel α-satellite HOR variants within chromosome 10 centromeres. a)** Comparison of the NA19240 (H1), HG03371 (H1), CHM13, and CHM1 chromosome 10 *D10Z1* α-satellite HOR arrays, showing that NA19240 (H1) and HG03371 (H1) have two novel α-satellite HOR variants each, which alter the overall composition of the α-satellite HOR arrays. **b)** Structure of the α-satellite HOR variants found in the NA19240 (H1), HG03371 (H1), CHM13, and CHM1 chromosome 10 centromeres. All of them derive from an ancestral 8-mer α-satellite HOR. Novel α-satellite HOR variants are indicated with an asterisk.

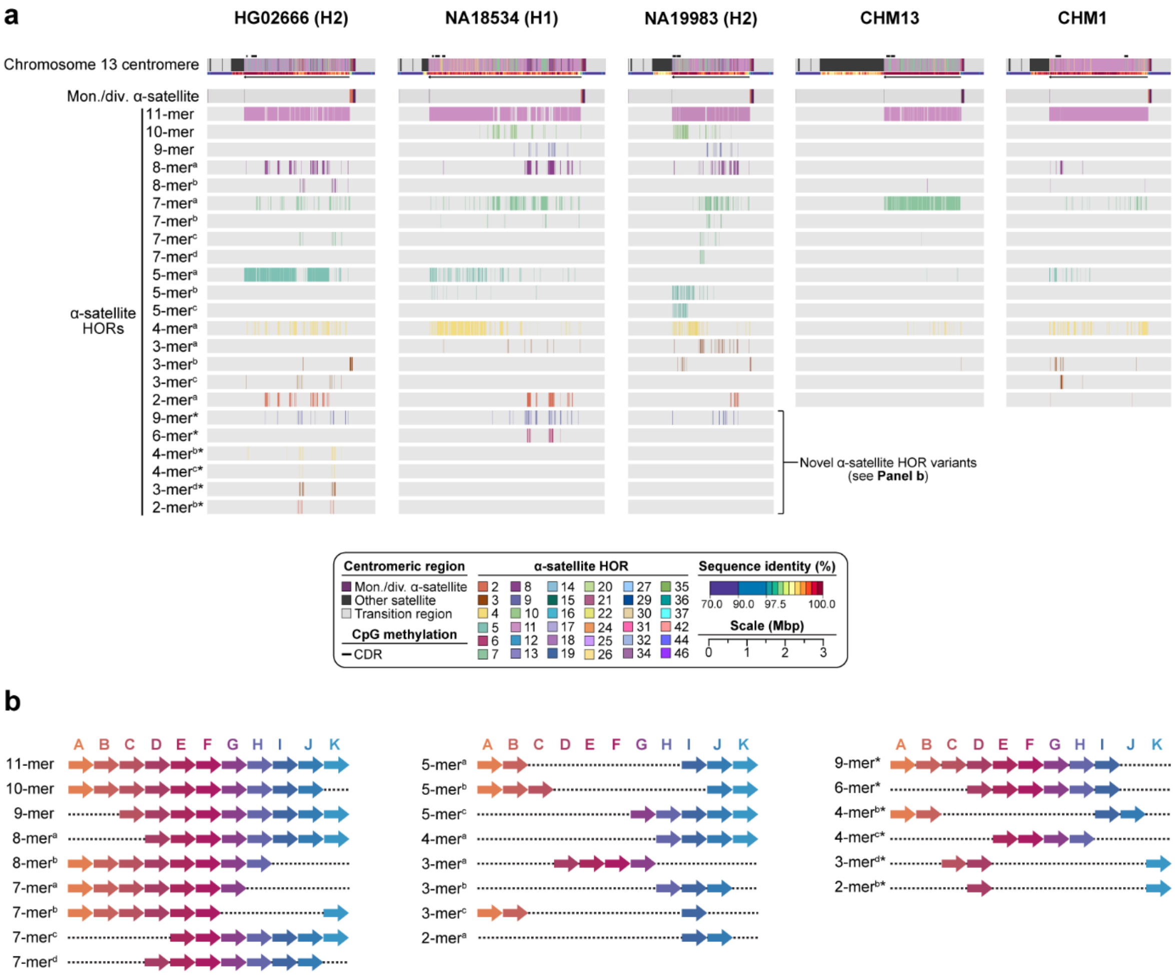


**Supplementary Figure 16. Novel α-satellite HOR variants within chromosome 13 centromeres. a)** Comparison of the HG02666 (H2), NA18534 (H1), NA19983 (H2), CHM13, and CHM1 chromosome 13 *D13Z1* α-satellite HOR arrays, showing that HG02666 (H2), NA18534 (H1), and NA19983 (H2) have five, two, and one novel α-satellite HOR variant, respectively, which alters the overall composition of the α-satellite HOR arrays. **b)** Structure of the α-satellite HOR variants found in the HG02666 (H2), NA18534 (H1), NA19983 (H2), CHM13, and CHM1 chromosome 13 centromeres. All of them derive from an ancestral 11-mer α-satellite HOR. Novel α-satellite HOR variants are indicated with an asterisk.

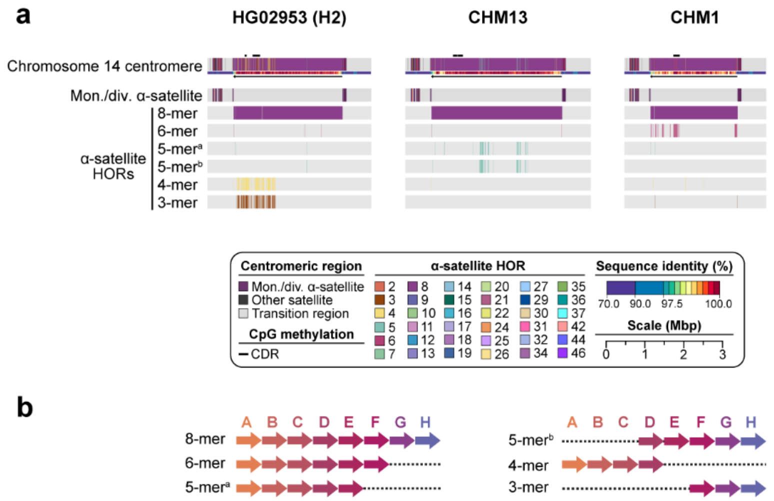


**Supplementary Figure 17. Differences in composition and abundance of α-satellite HOR variants within chromosome 14 centromeres. a)** Comparison of the HG02953 (H2), CHM13, and CHM1 chromosome 14 *D14Z9* α-satellite HOR arrays reveals that HG02953 (H2) has two α-satellite HOR variants in much higher abundance than CHM13 and CHM1, which alters the overall composition of the α-satellite HOR array. **b)** Structure of the α-satellite HOR variants found in the HG02953 (H2), CHM13, and CHM1 chromosome 14 centromeres. All of them derive from an ancestral 8-mer α-satellite HOR.

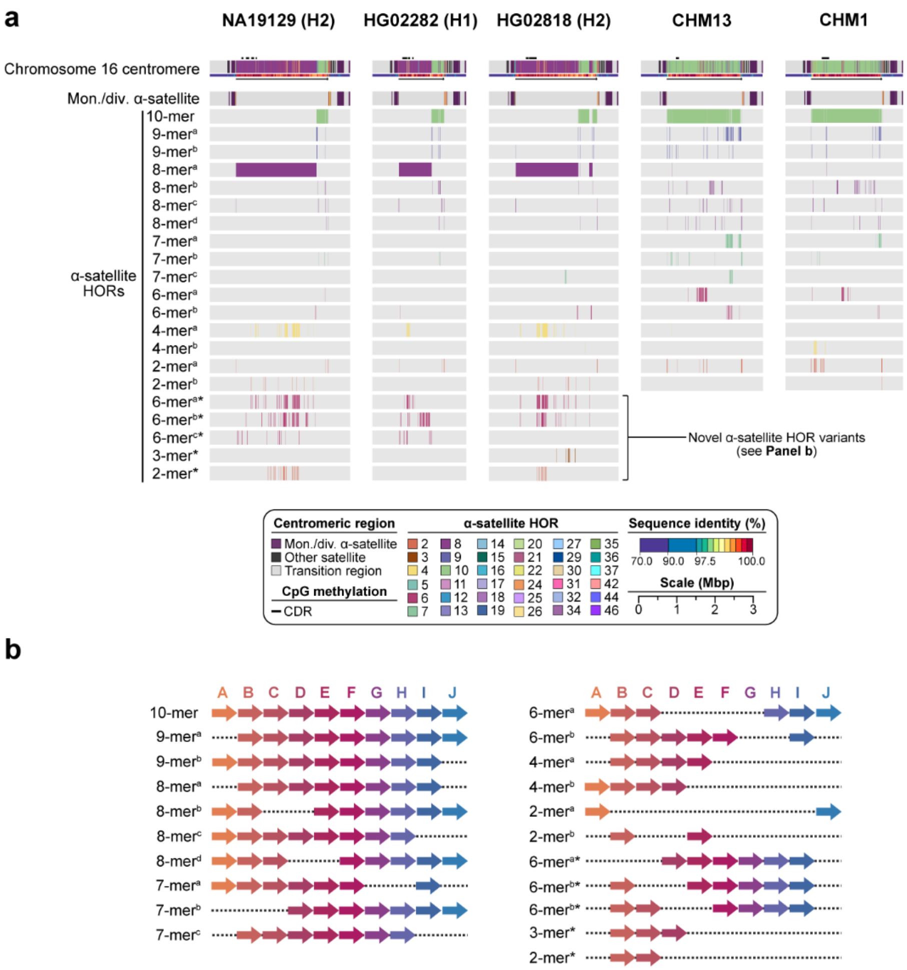

**Supplementary Figure 18. Novel α-satellite HOR variants within chromosome 16 centromeres. a)** Comparison of the NA19129 (H2), HG02282 (H1), HG02818 (H2), CHM13, and CHM1 chromosome 16 *D16Z2* α-satellite HOR arrays, showing that NA19129 (H2), HG02282 (H1), and HG02818 (H2) have three or four novel α-satellite HOR variants, which alter the overall composition of the α-satellite HOR arrays. **b)** Structure of the α-satellite HOR variants found in the NA19129 (H2), HG02282 (H1), HG02818 (H2), CHM13, and CHM1 chromosome 16 centromeres. All of them derive from an ancestral 10-mer α-satellite HOR. Novel α-satellite HOR variants are indicated with an asterisk.

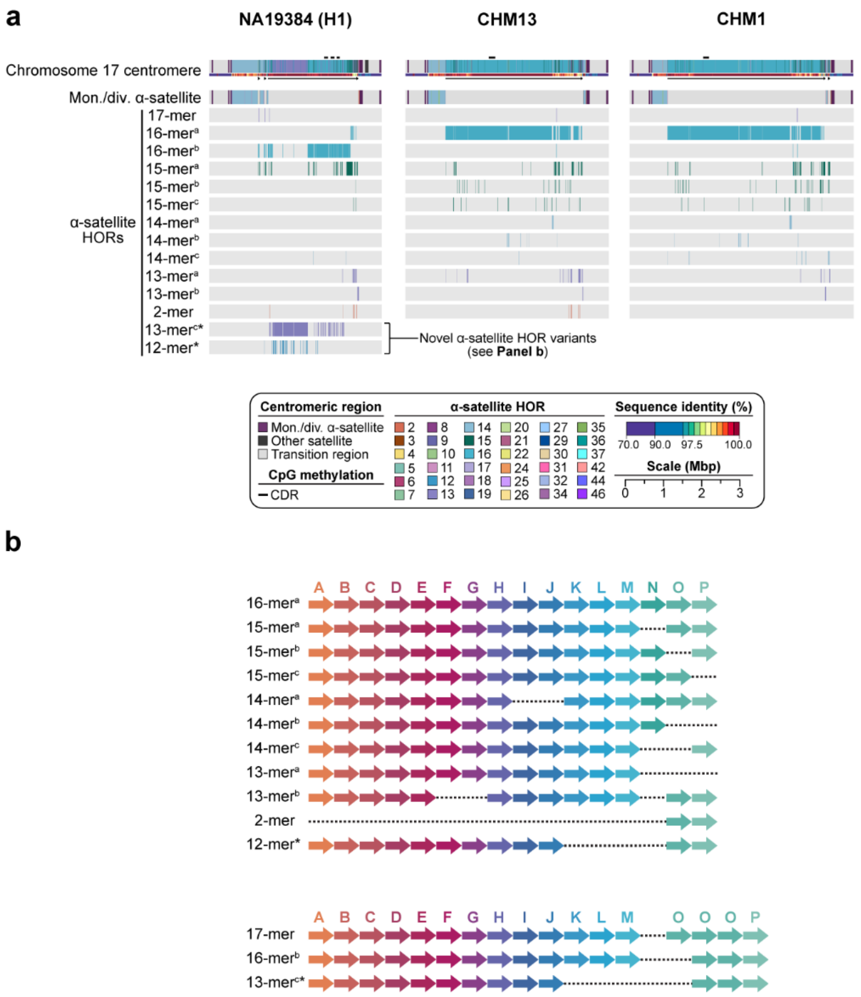


**Supplementary Figure 19. Novel α-satellite HOR variants within chromosome 17 centromeres. a)** Comparison of the NA19384 (H2), CHM13, and CHM1 chromosome 17 *D17Z1* α-satellite HOR arrays, showing that NA19384 (H2) has two novel α-satellite HOR variants, which alter the overall composition of the α-satellite HOR array. **b)** Structure of the α-satellite HOR variants found in the NA19384 (H2), CHM13, and CHM1 chromosome 17 centromeres. All of them derive from an ancestral 16-mer α-satellite HOR. Novel α-satellite HOR variants are indicated with an asterisk.

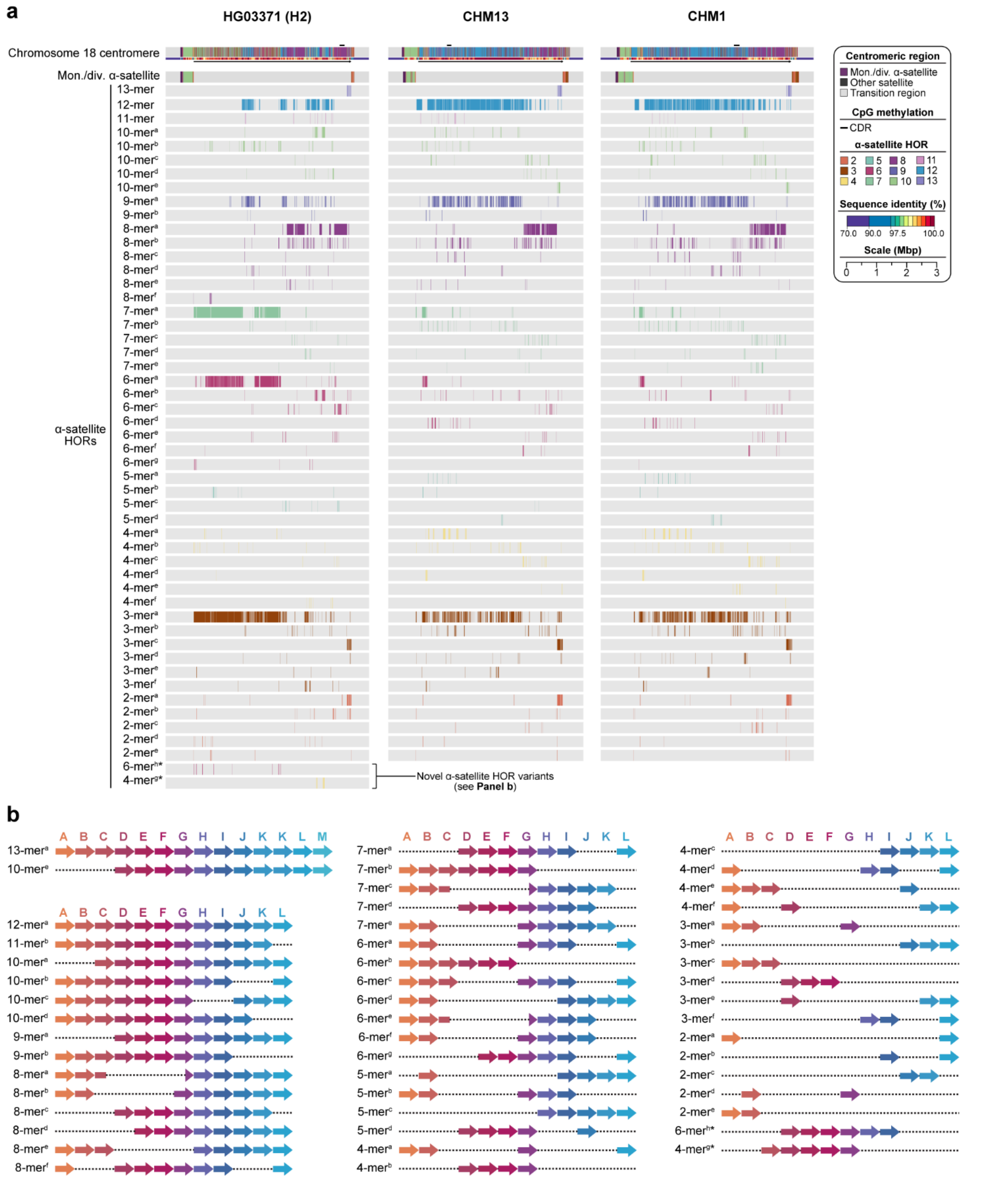


**Supplementary Figure 20. Novel α-satellite HOR variants within chromosome 18 centromeres. a)** Comparison of the HG03371 (H2), CHM13, and CHM1 chromosome 18 *D18Z1* α-satellite HOR arrays, showing that HG03371 (H2) have two novel α-satellite HOR variants, which alter the overall composition of the α-satellite HOR array. **b)** Structure of the α-satellite HOR variants found in the HG03371 (H2), CHM13, and CHM1 chromosome 18 centromeres. All of them derive from an ancestral 12-mer α-satellite HOR. Novel α-satellite HOR variants are indicated with an asterisk.

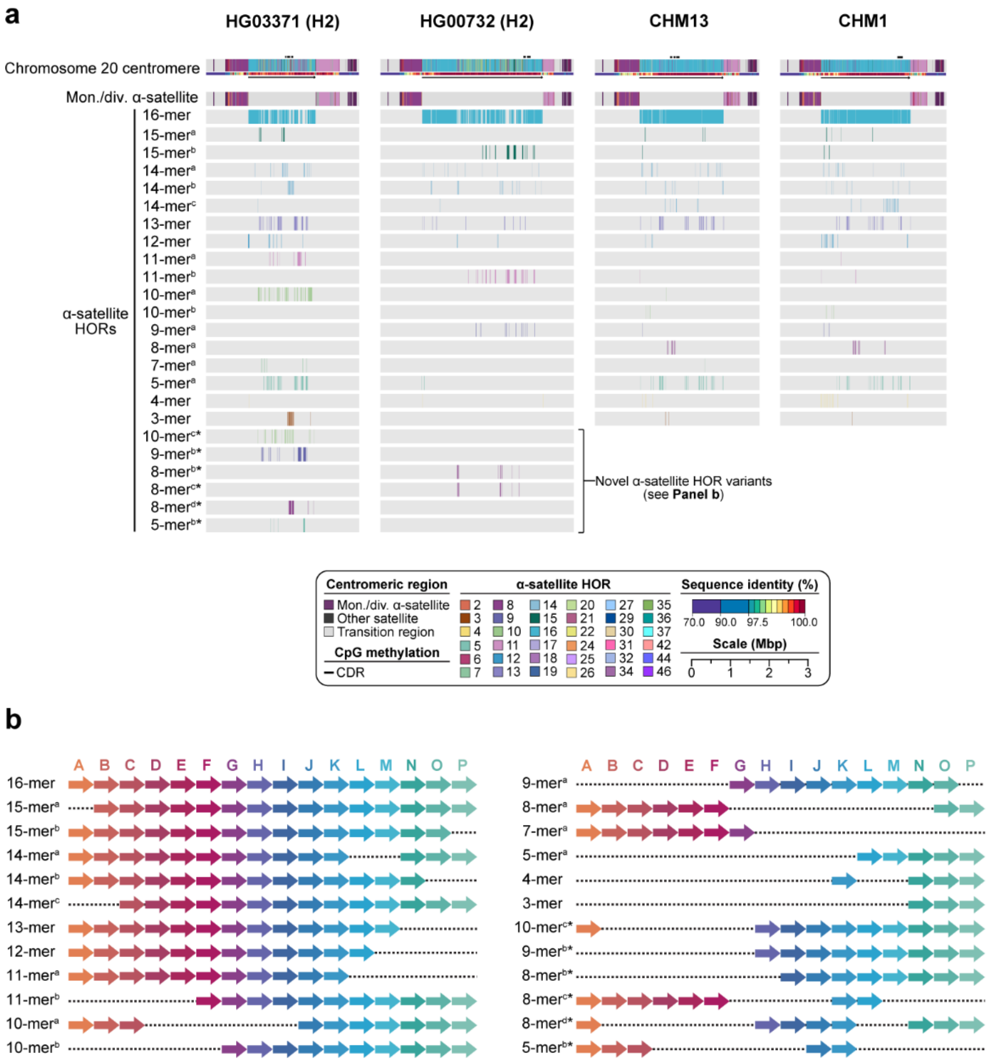


**Supplementary Figure 21. Novel α-satellite HOR variants within chromosome 20 centromeres. a)** Comparison of the HG03371 (H2), HG00732 (H2), CHM13, and CHM1 chromosome 20 *D20Z2* α-satellite HOR arrays, showing that HG03371 (H2) and HG00732 (H2) have four and two novel α-satellite HOR variants, respectively, which alter the overall composition of the α-satellite HOR arrays. **b)** Structure of the α-satellite HOR variants found in the HG03371 (H2), HG00732 (H2), CHM13, and CHM1 chromosome 20 centromeres. All of them derive from an ancestral 16-mer α-satellite HOR. Novel α-satellite HOR variants are indicated with an asterisk.

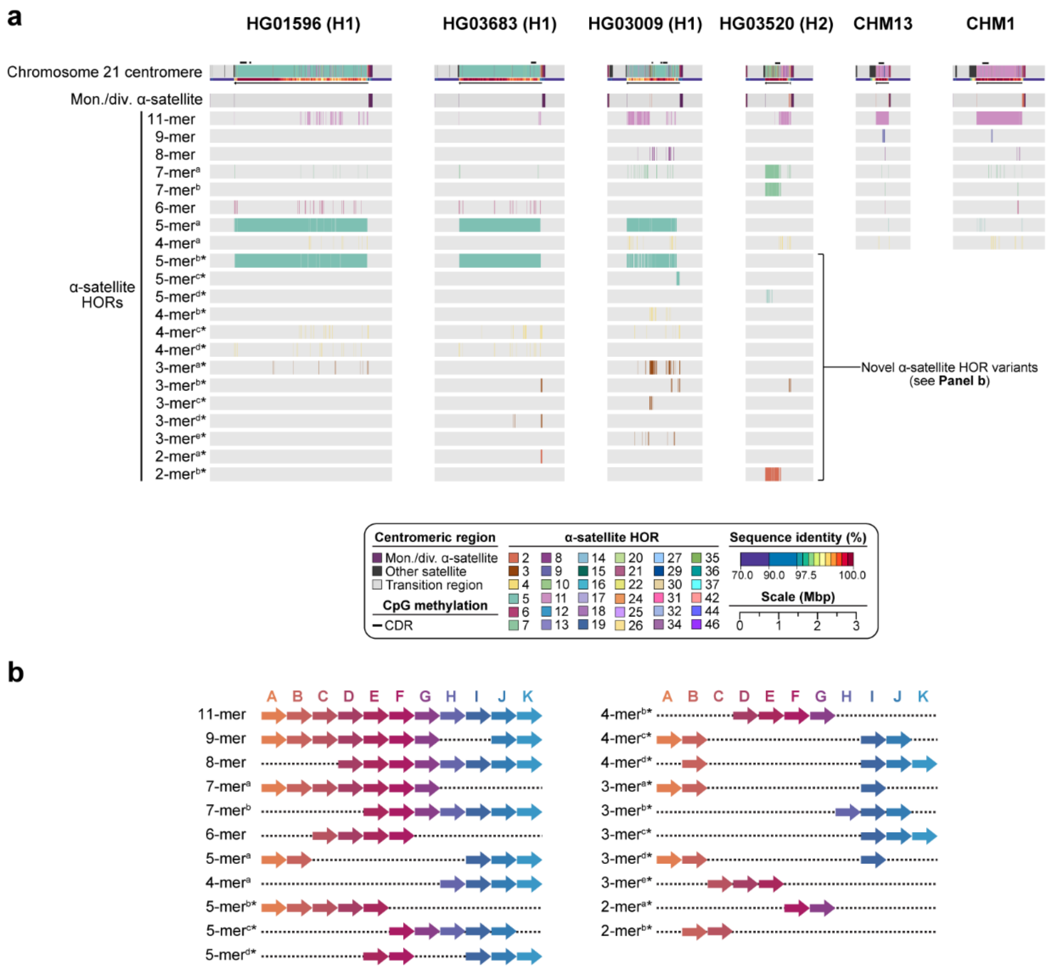


**Supplementary Figure 22. Novel α-satellite HOR variants within chromosome 21 centromeres. a)** Comparison of the HG01596 (H1), HG03683 (H1), HG03009 (H1), HG03520 (H2), CHM13, and CHM1 chromosome 13 *D13Z1* α-satellite HOR arrays, showing that HG01596 (H1), HG02666 (H2), NA18534 (H1), and NA19983 (H2) have up to eight novel α-satellite HOR variants, which alter the overall composition of the α-satellite HOR arrays. **b)** Structure of the α-satellite HOR variants found in the HG01596 (H1), HG03683 (H1), HG03009 (H1), HG03520 (H2), CHM13, and CHM1 chromosome 21 centromeres. All of them derive from an ancestral 11-mer α-satellite HOR. Novel α-satellite HOR variants are indicated with an asterisk.

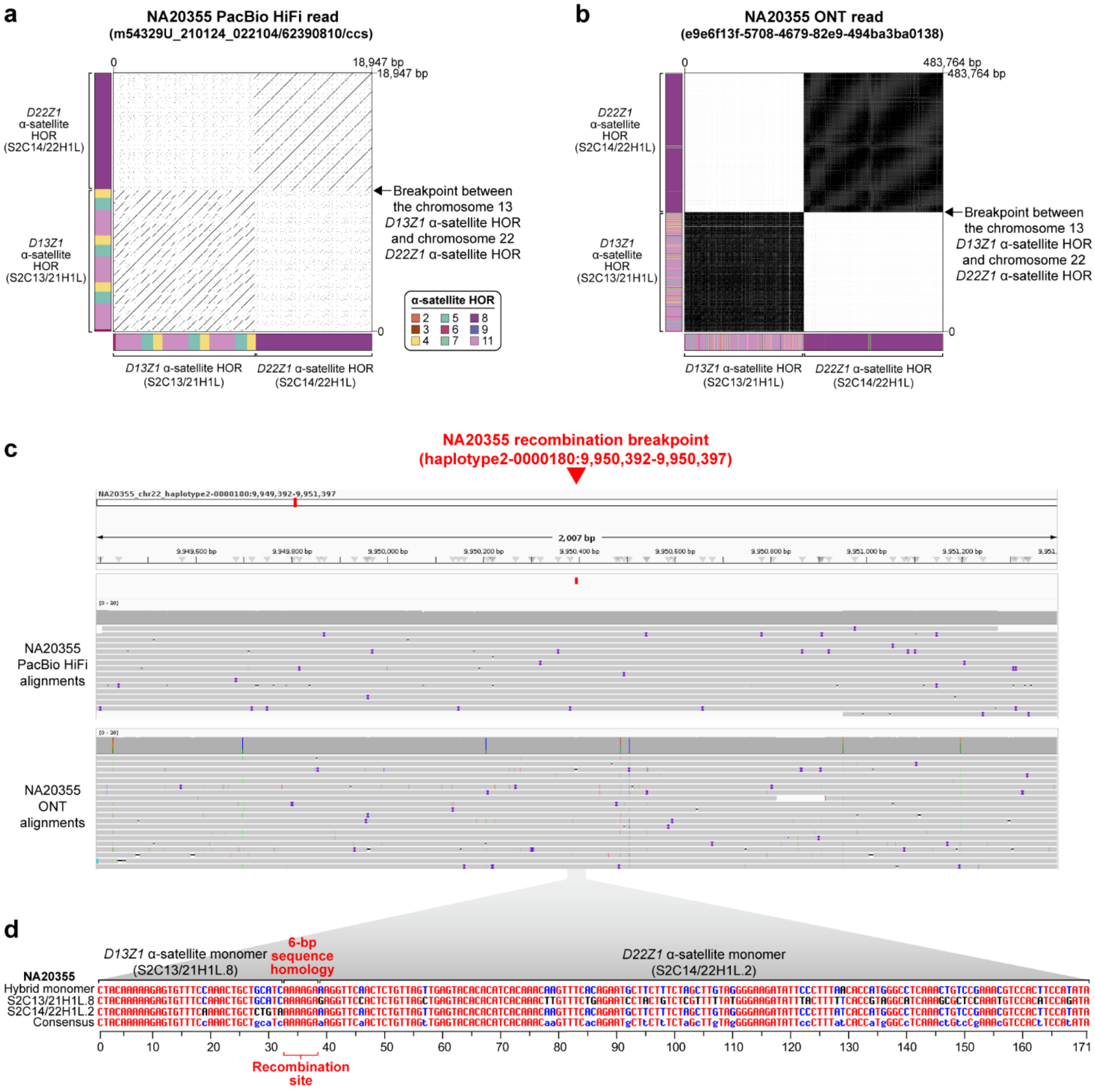


**Supplementary Figure 23. Evidence of a recombination event between chromosome 13 and chromosome 22 centromeres in NA20355, giving rise to a hybrid 13/22 chromosome. a,b)** Dot plots of **a)** a PacBio HiFi read and **b)** an ONT read showing the transition between chromosome 13 *D13Z1* and chromosome 22 *D22Z1* α-satellite HORs in NA20355. Word length, 20 bp for the PacBio HiFi read, 50 bp for the ONT read, generated with Gepard[8]. **c)** Detection of the breakpoint within NA20355 and support for the junction with raw NA20355 PacBio HiFi and ONT reads. **d)** Comparison of the NA20355 hybrid monomer to the S2C13/21H1L.8 and S2C144/22H1L.2 monomers up and downstream of the breakpoint. A 6-bp homology sequence (AAAAGA) is found among all three and likely contributes to microhomology-mediated recombination, leading to the hybrid 13/22 chromosome.

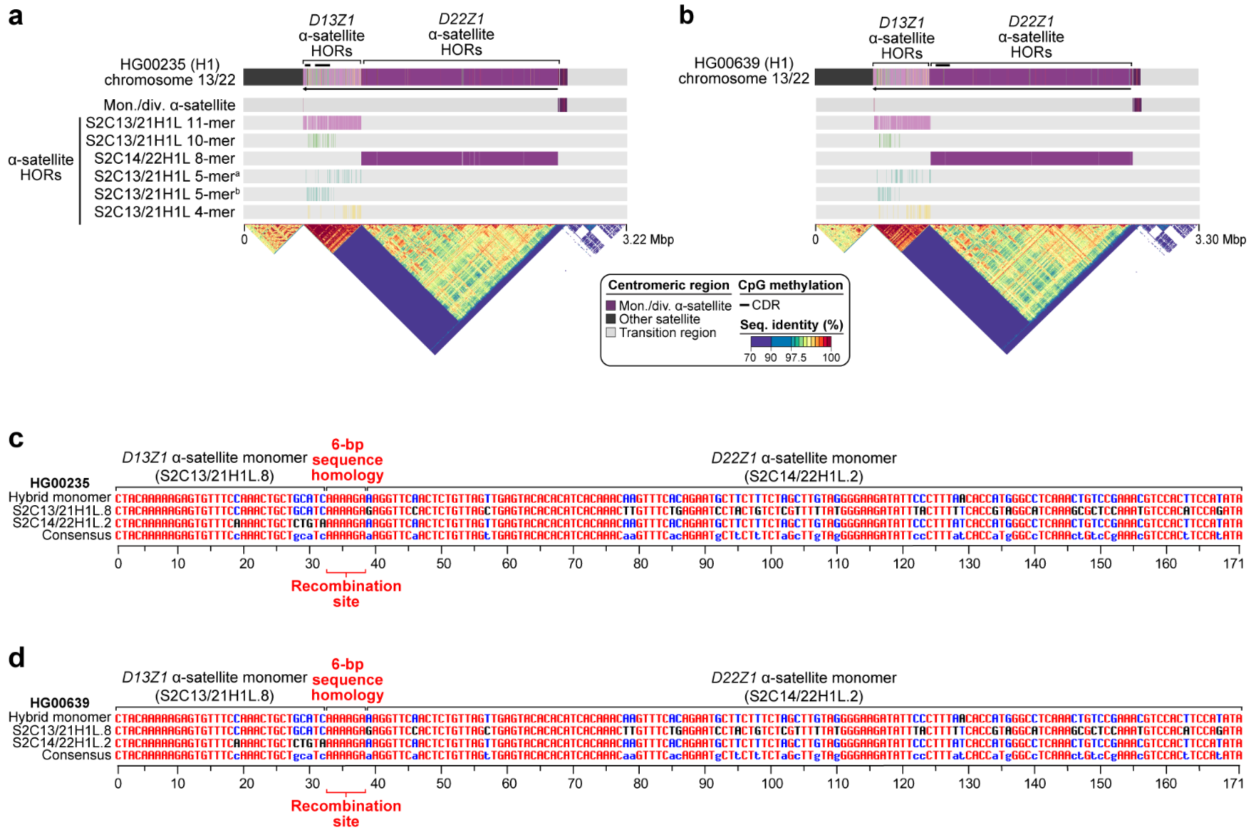


**Supplementary Figure 24. Discovery of a hybrid chromosome 13/22 in two other individuals, with a recombination breakpoint in the centromere. a,b)** Structure and composition of the chromosome 13/22 centromeres in **a)** HG00235 and **b)** HG00639. **c,d)** Comparison of the chromosome 13/22 hybrid monomer to the S2C13/21H1L.8 and S2C144/22H1L.2 monomers up and downstream of the breakpoint in **c)** HG00235 and **d)** HG00639. A 6-bp homology sequence (AAAAGA) is found among all three and likely contributes to microhomology-mediated recombination.

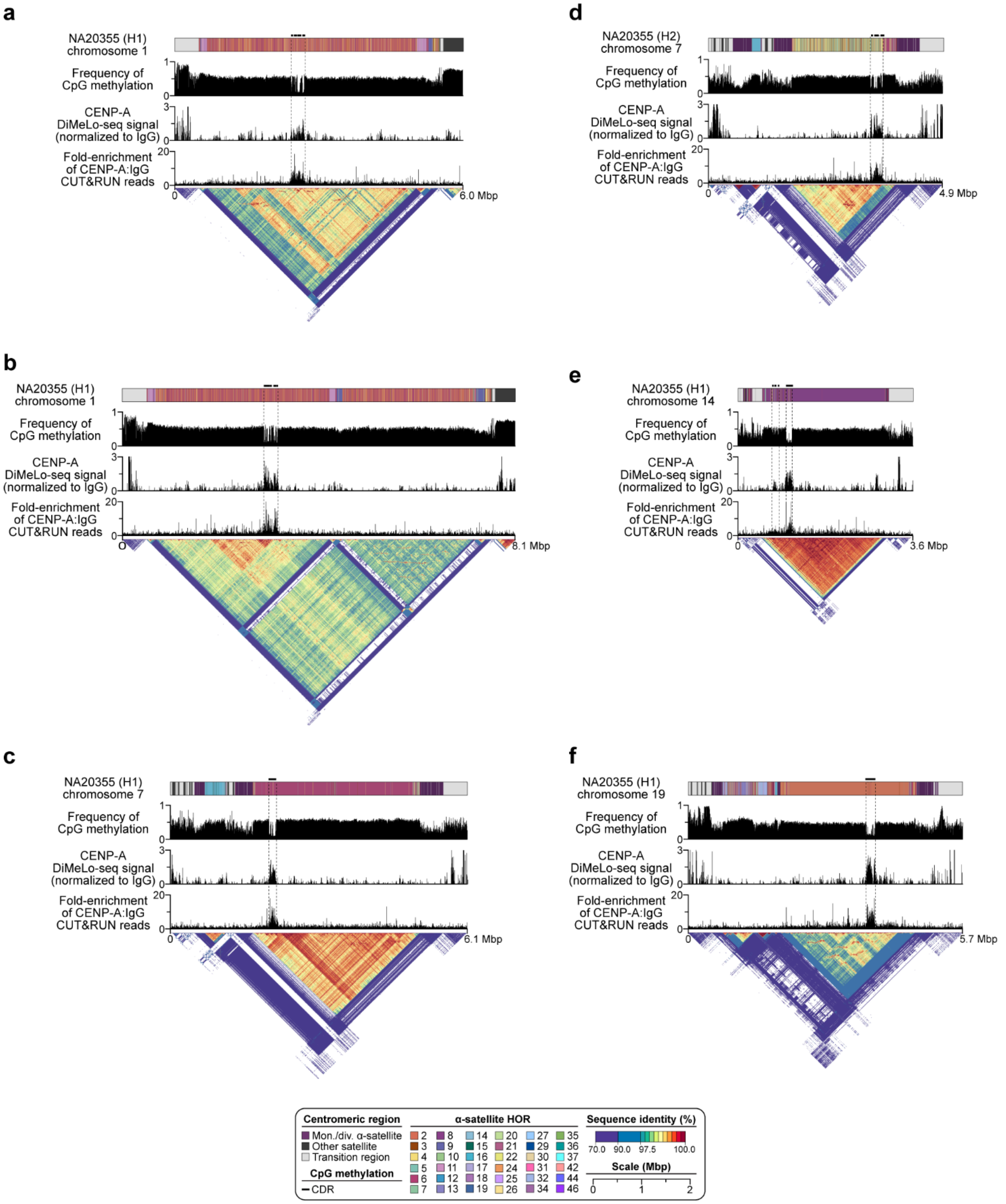


**Supplementary Figure 25. CENP-A DiMeLo-seq and CUT&RUN followed by ONT long-read sequencing shows enrichment of CENP-A within the CDR of NA20355 centromeres. a-f)** Plots showing the presence of a CDR enriched with CENP-A within the centromeres of chromosomes **a,b)** 1, **c,d)** 7, **e)** 14, and **f)** 19. Both CENP-A DiMeLo-seq and long-read CENP-A CUT&RUN are concordant and show an enrichment of CENP-A within the CDRs.

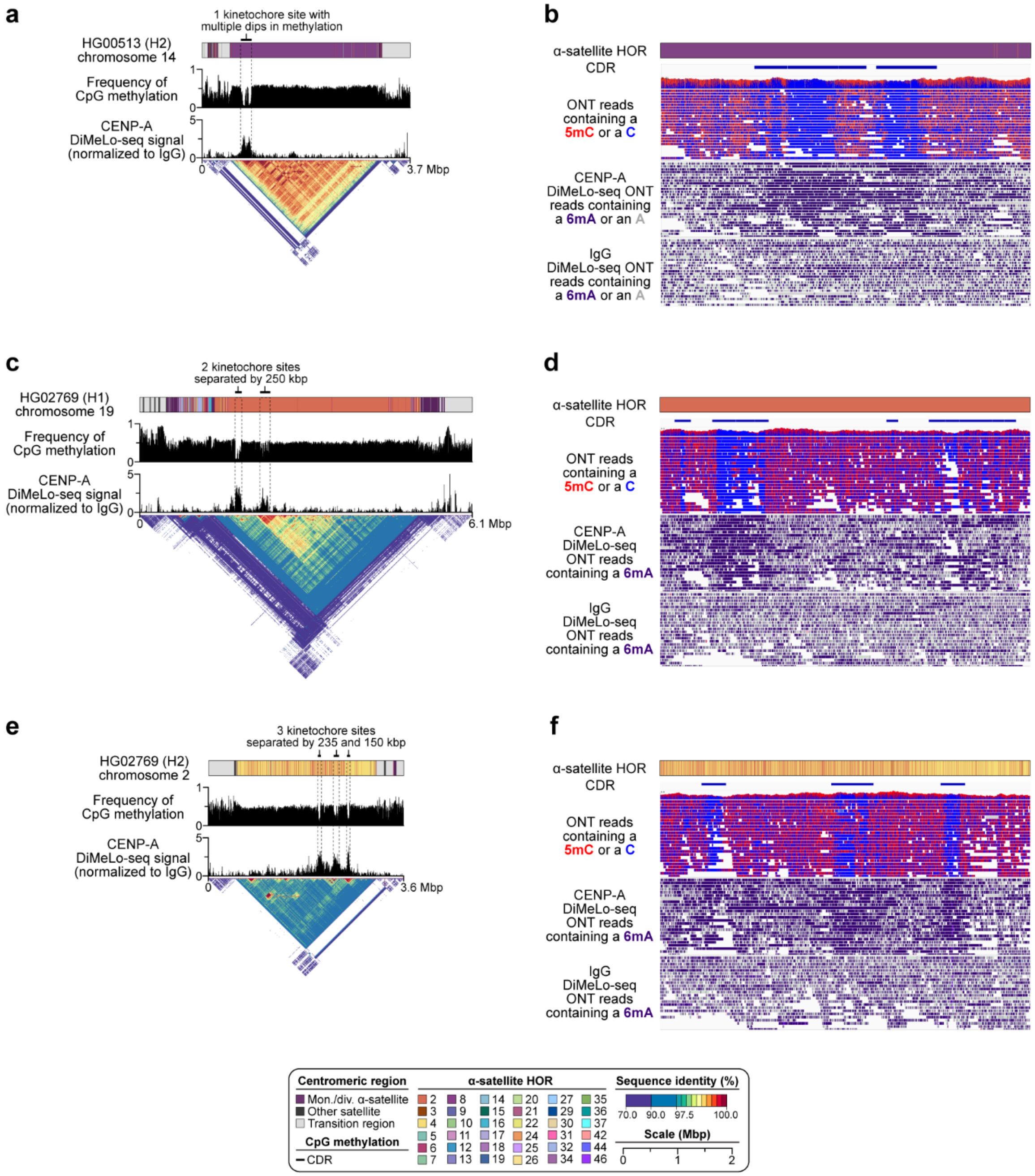


**Supplementary Figure 26. Examples of single and multi-kinetochore sites within human centromeres. a)** Plot showing a single kinetochore site within the HG00513 chromosome 1 centromere (H2) as determined by the presence of a CDR enriched with CENP-A. **b)** Raw ONT sequencing reads showing a hypomethylated region with four dips in methylation enriched with CENP-A, as determined with DiMeLo-seq. **c)** Plot showing two kinetochore sites within the HG02769 chromosome 19 centromere (H1) as determined by the presence of two CDRs enriched with CENP-A, separated by 250 kbp of sequence. **d)** Raw ONT sequencing reads showing two hypomethylated regions enriched with CENP-A, as determined with DiMeLo-seq. **e)** Plot showing three kinetochore sites within the HG02789 chromosome 2 centromere (H2) as determined by the presence of three CDRs enriched with CENP-A, separated by 235 and 150 kbp of sequence. **f)** Raw ONT sequencing reads showing three hypomethylated regions enriched with CENP-A, as determined with DiMeLo-seq.

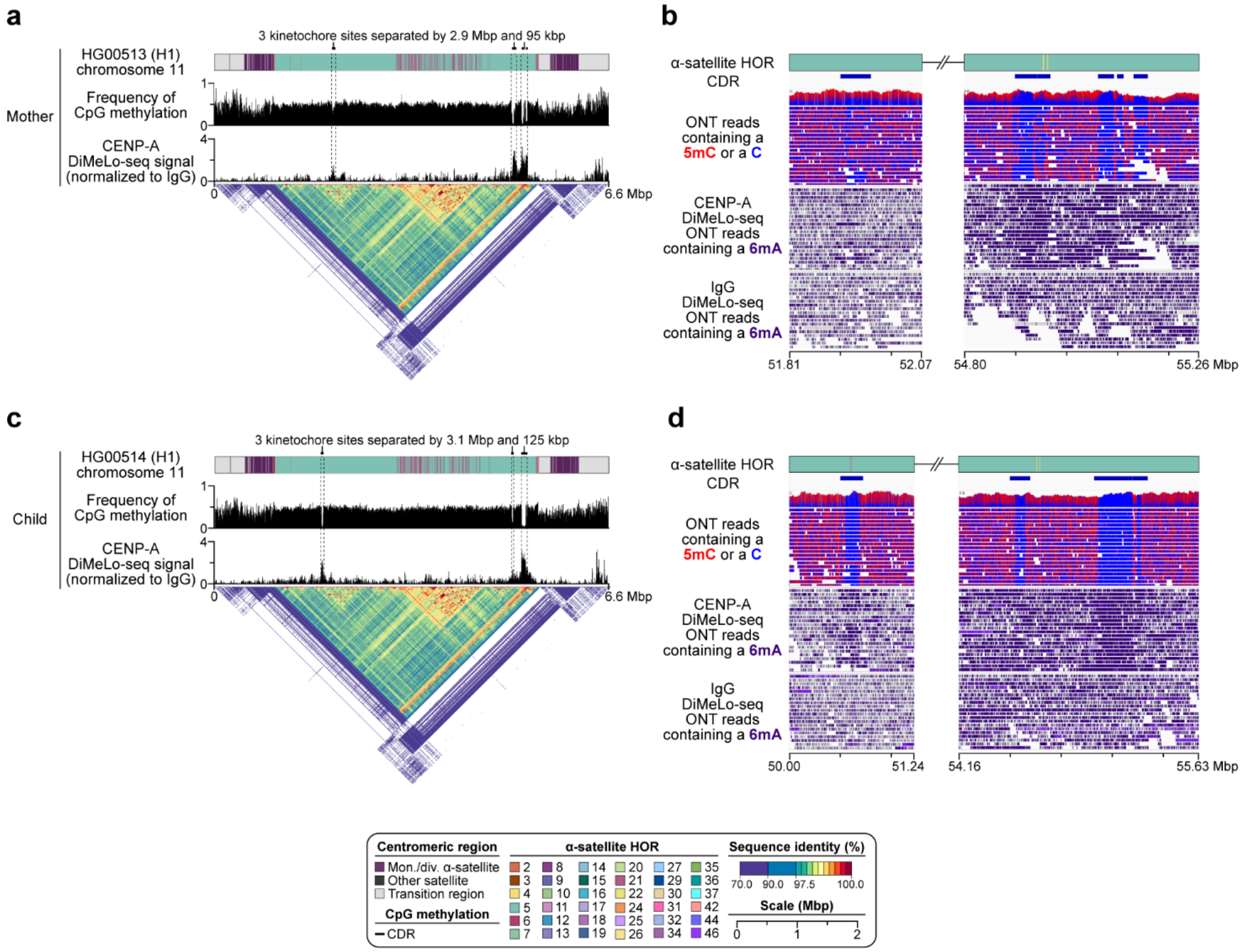


**Supplementary Figure 27. Di-kinetochores separated by 2.9 Mbp of sequence are stably transmitted across generations. a)** Plot showing multiple kinetochore sites within the HG00513 chromosome 11 centromere (H1), with two of them separated by ~2.9 Mbp of sequence and the third separated by 95 kbp of sequence. **b)** Raw ONT sequencing reads showing hypomethylated regions enriched with CENP-A within the HG00513 chromosome 11 centromere (H1), as determined with DiMeLo-seq. **c)** Plot showing that multiple kinetochores within the chromosome 11 centromere are transmitted to the child, HG00514, with slight changes in position. **d)** Raw ONT sequencing reads showing hypomethylated regions enriched with CENP-A within the HG00514 chromosome 11 centromere (H1), as determined with DiMeLo-seq.

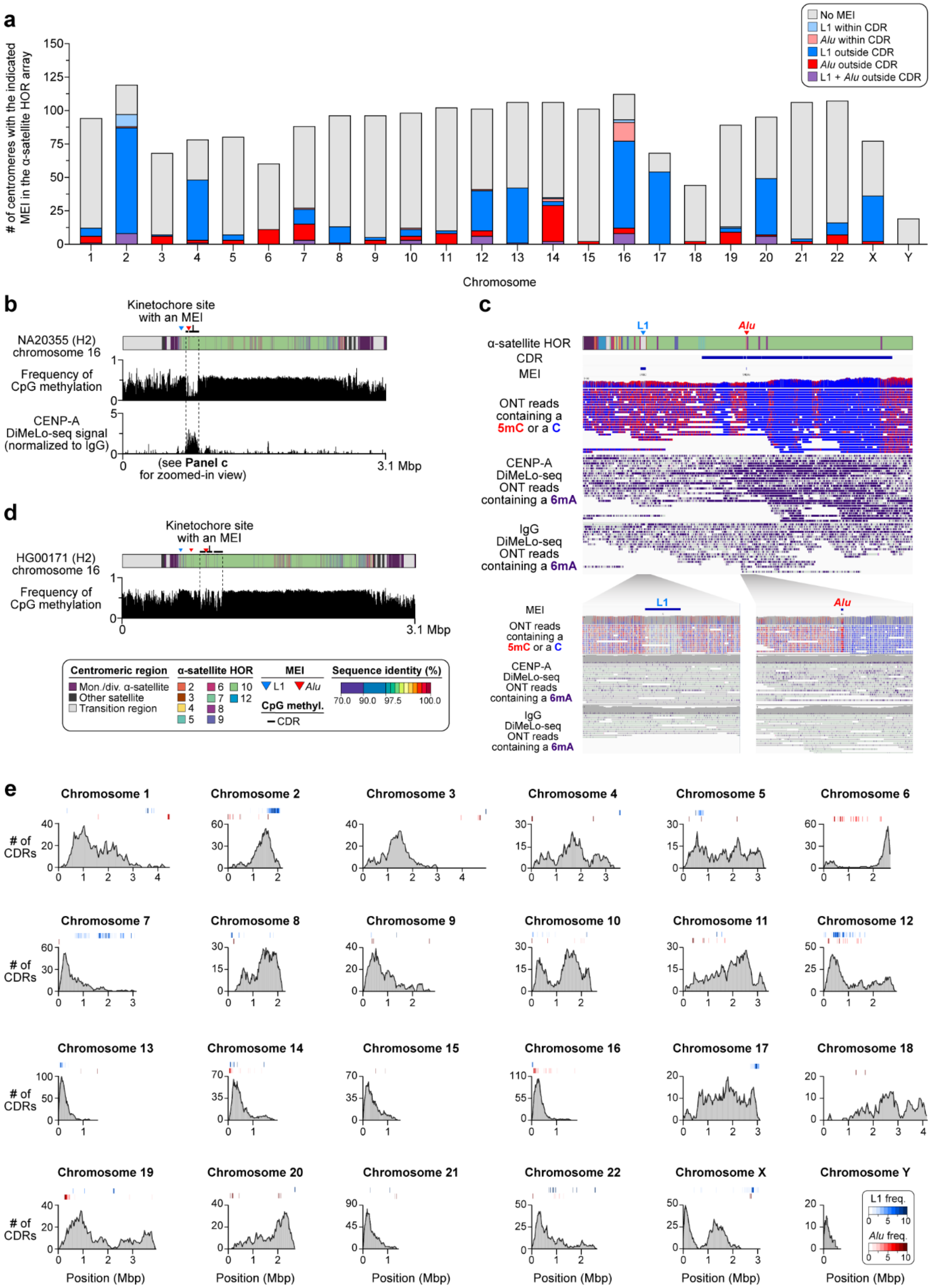


**Supplementary Figure 28. Frequency of MEIs within centromeres and putative kinetochore sites. a)** Number of centromeres with an MEI within the α-satellite HOR array or the putative kinetochore site, defined by the presence of the CDR. **b)** Example of an *Alu* element inserted in a putative kinetochore site of the NA20355 chromosome 16 centromere (H2). An L1 element is also inserted upstream of the kinetochore site. **c)** Raw ONT sequencing reads showing that the *Alu* element changes the methylation profile of the CDR. Despite the presence of methylated cytosines, CENP-A is still enriched over the *Alu* element relative to IgG, as determined with DiMeLo-seq. In contrast, the L1 element upstream of the kinetochore site has similar levels of both CENP-A and IgG DiMeLo-seq signals. **d)** Plot showing an *Alu* element inserted within the putative kinetochore site of the HG00171 chromosome 16 centromere (H2). Another *Alu* element and an L1 element are also inserted upstream of the putative kinetochore site. **e)** Distribution of MEIs across centromeric α-satellite HOR arrays relative to CDR position (shown in gray). L1s are shown in blue, and *Alu*s are shown in red.

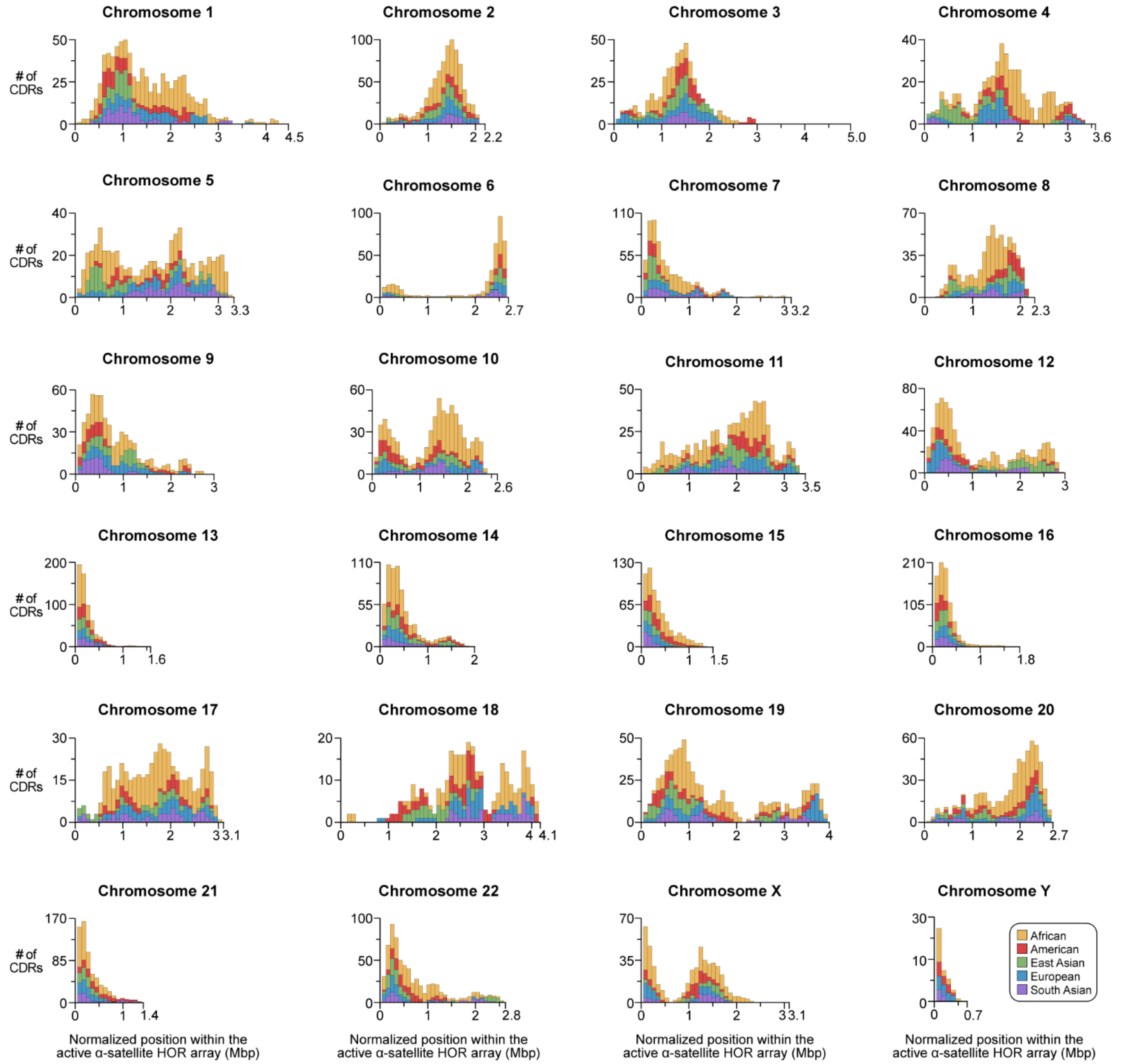


**Supplementary Figure 29. The position of the putative kinetochore is closely associated with the underlying sequence and structure of the centromere.** The position of the CDR across active α-satellite HOR arrays for chromosomes 1-22, X and Y. For 11 out of 24 chromosomes (chromosomes 1, 3, 7, 9, 13-16, 21, 22, and Y), CDRs are mainly located on the p arm-proximal side of the array. For 2 out of 24 chromosomes (chromosomes 2 and 20), CDRs are mainly located on the q arm-proximal side of the array. For the remaining 11 out of 24 chromosomes (chromosomes 4-6, 8, 10-12, 17-19, and X), CDRs are located on different regions of the α-satellite HOR array depending on their haplotype structure.

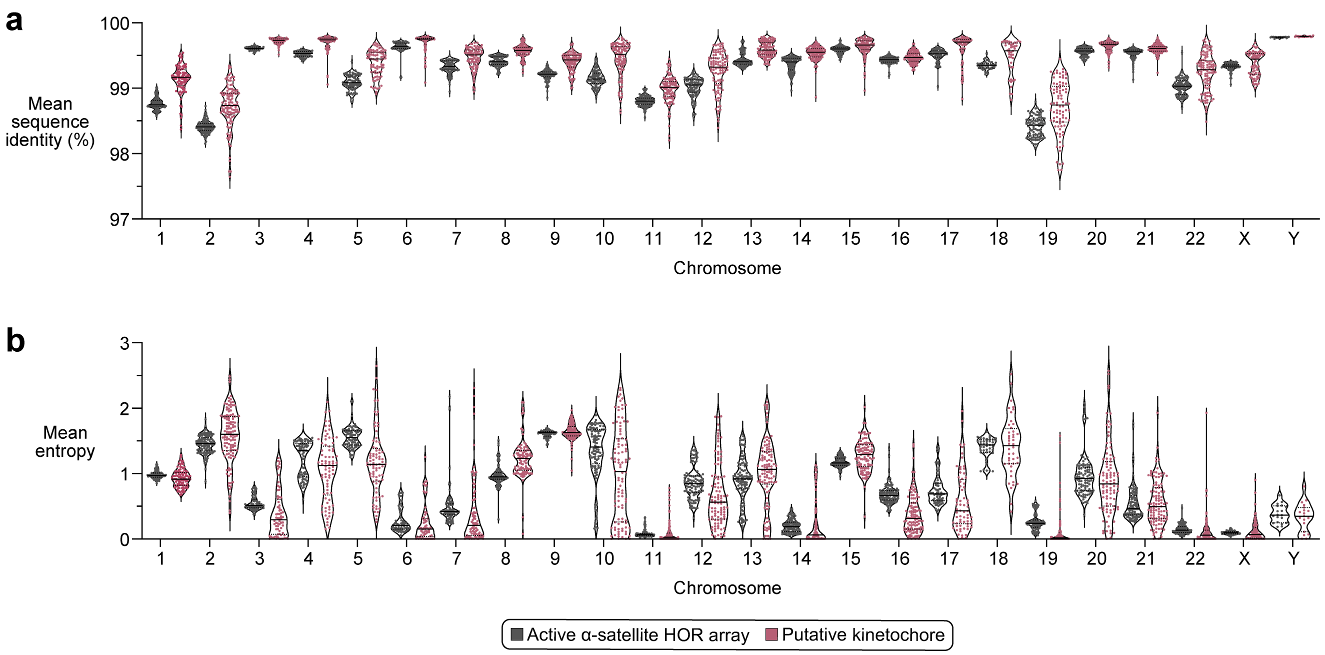


**Supplementary Figure 30. Sequences underlying the putative kinetochore have a higher mean sequence identity and lower entropy than the rest of the active α-satellite HOR array. a,b)** Plot showing the **a)** mean sequence identity or **b)** mean entropy for the active α-satellite HOR array vs. the putative kinetochore site for each centromere in 65 diverse human genomes. All chromosomes have a higher mean sequence identity for the putative kinetochore site compared to the rest of the α-satellite HOR array. Almost all chromosomes, except for chromosomes 8 and 13, have a lower mean entropy for the putative kinetochore site compared to the rest of the α-satellite HOR array.

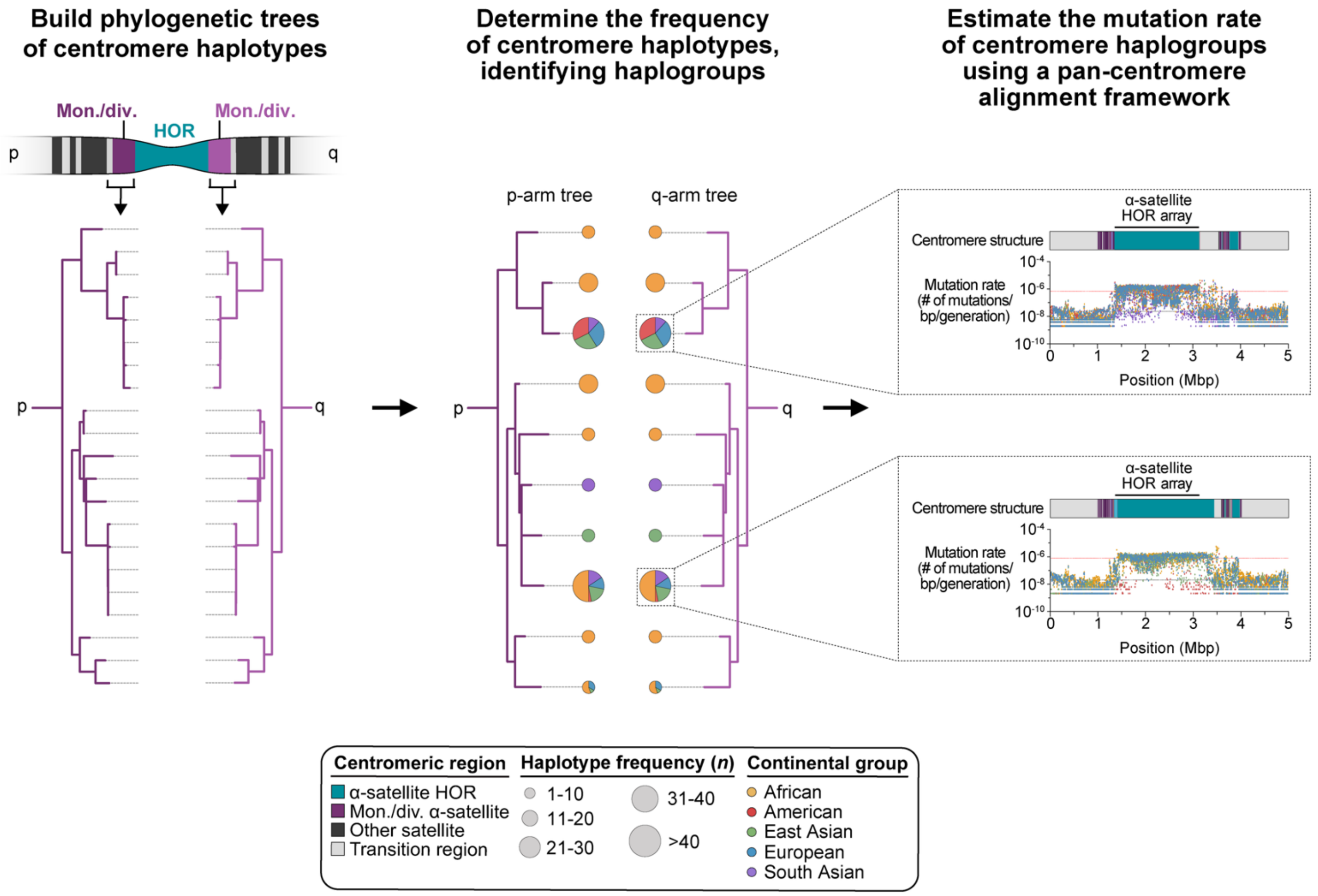


**Supplementary Figure 31. Method to reconstruct the evolutionary history of human centromeres, calculate their haplotype frequency, and estimate their mutation rate among global populations.** Using sequences within the monomeric/divergent α-satellite and/or neighboring sequences in the p and q arms of each centromere, we built maximum-likelihood phylogenetic trees, using chimpanzee as an outgroup and assuming a human-chimpanzee divergence time of 6.2 million years. We identified centromere haplogroups specific to particular populations and/or shared among others and determined their frequency among global populations. Finally, we estimated the mutation rate of each centromere haplogroup using a pan-centromere alignment framework, revealing a 58-fold difference in mutation rate among centromeres from different chromosomes (**Methods**).

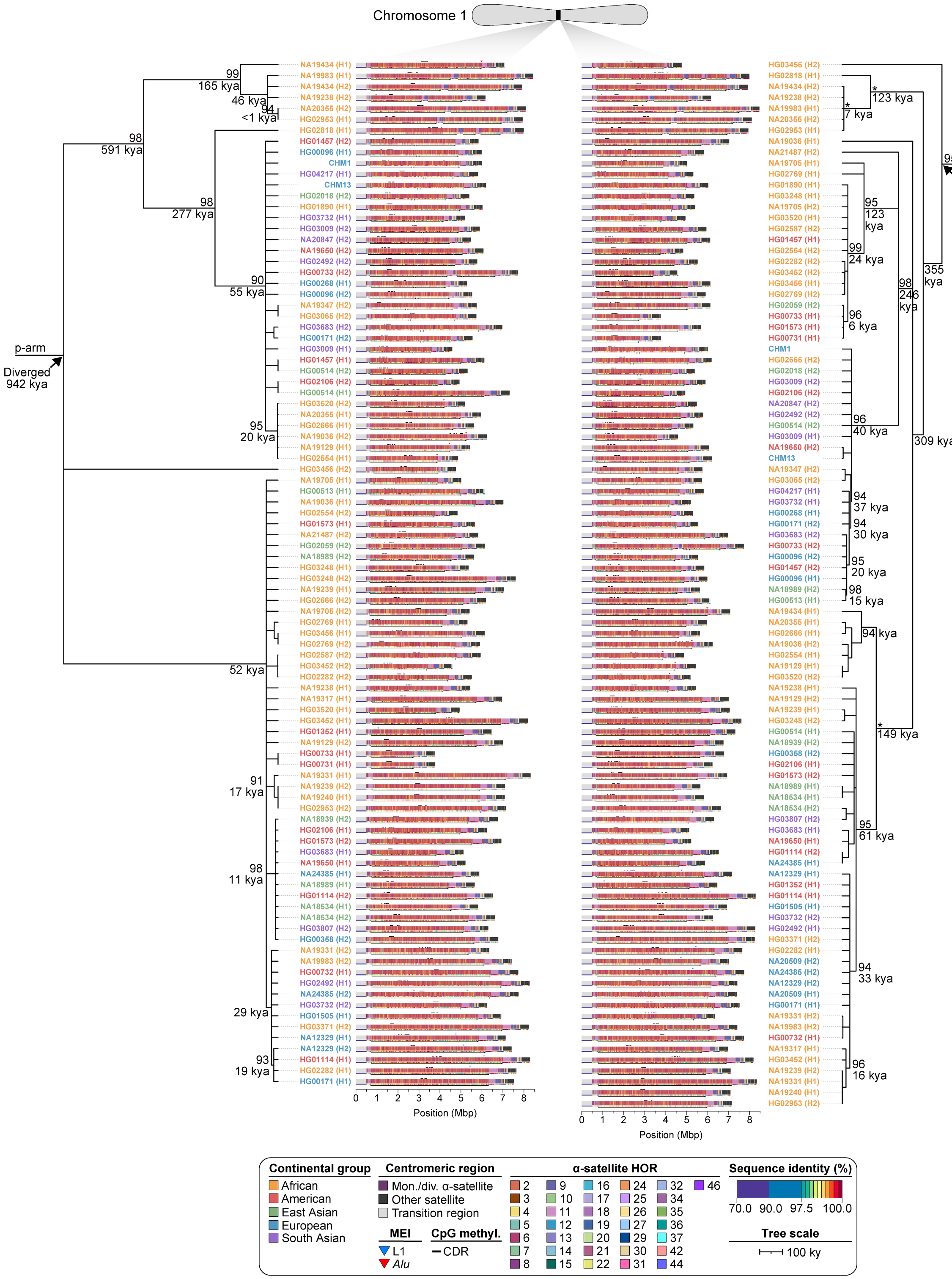


**Supplementary Figure 32. Phylogenetic reconstruction of chromosome 1 centromere haplotypes.** Maximum-likelihood phylogenetic tree depicting the p- and q-arm topologies along with estimated divergence times for the chromosome 1 (*D1Z7*) α-satellite HOR arrays from diverse human genomes. Asterisks indicate nodes with 100% bootstrap support, and nodes with 90-99% bootstrap support are indicated numerically. Nodes without an asterisk or number have bootstrap support <90%. We note that most differences in the order of the haplotypes occur at the terminal branches where the order of sequence taxa can be readily reshuffled to establish near-complete concordance. Thus, there are no significant changes in the overall topologies of the phylogenetic tree.

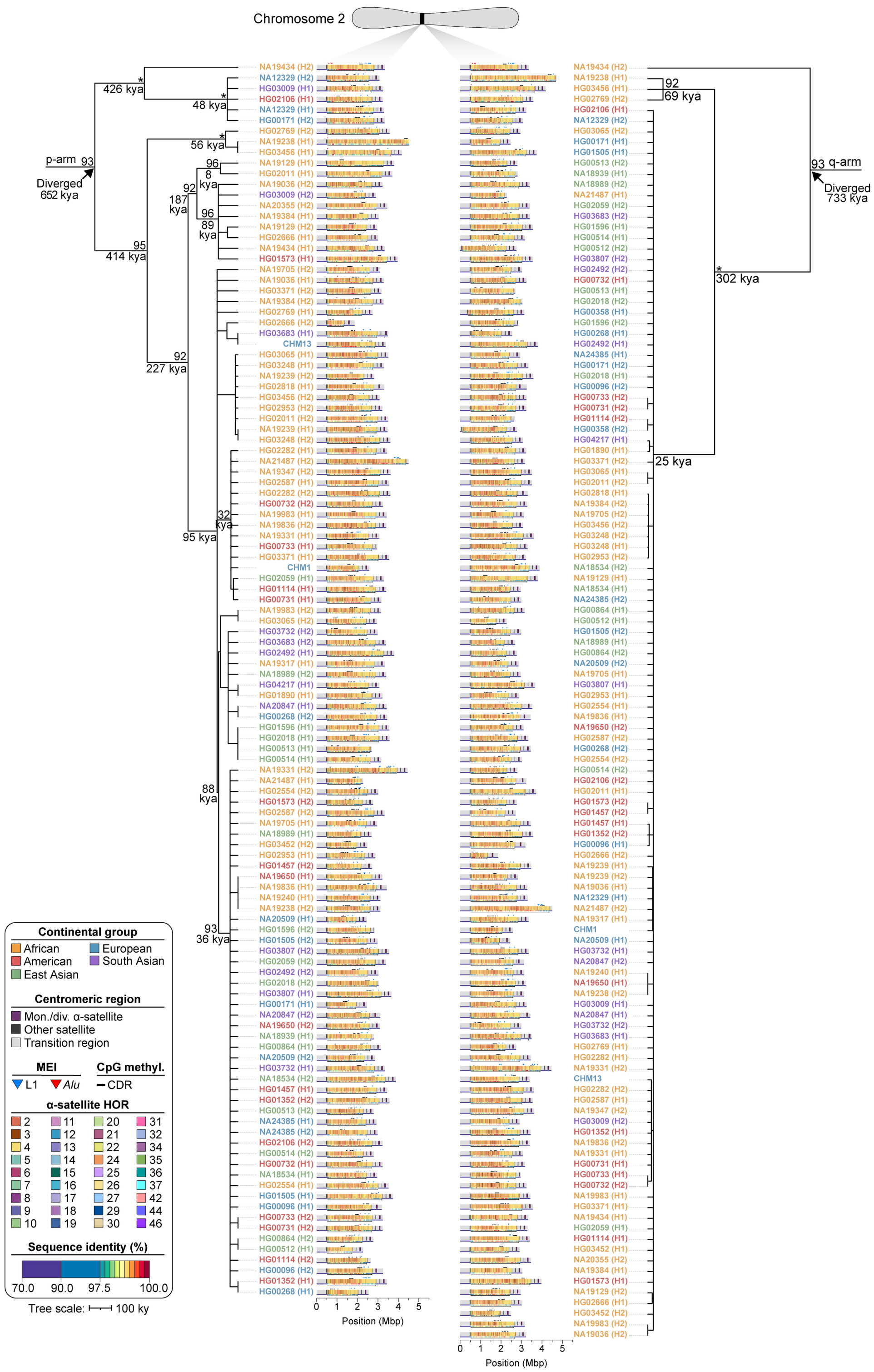


**Supplementary Figure 33. Phylogenetic reconstruction of chromosome 2 centromere haplotypes.** Maximum-likelihood phylogenetic tree depicting the p- and q-arm topologies along with estimated divergence times for the chromosome 2 (*D2Z1*) α-satellite HOR arrays from diverse human genomes. Asterisks indicate nodes with 100% bootstrap support, and nodes with 90-99% bootstrap support are indicated numerically. Nodes without an asterisk or number have bootstrap support <90%. We note that most differences in the order of the haplotypes occur at the terminal branches where the order of sequence taxa can be readily reshuffled to establish near-complete concordance. Thus, there are no significant changes in the overall topologies of the phylogenetic tree.

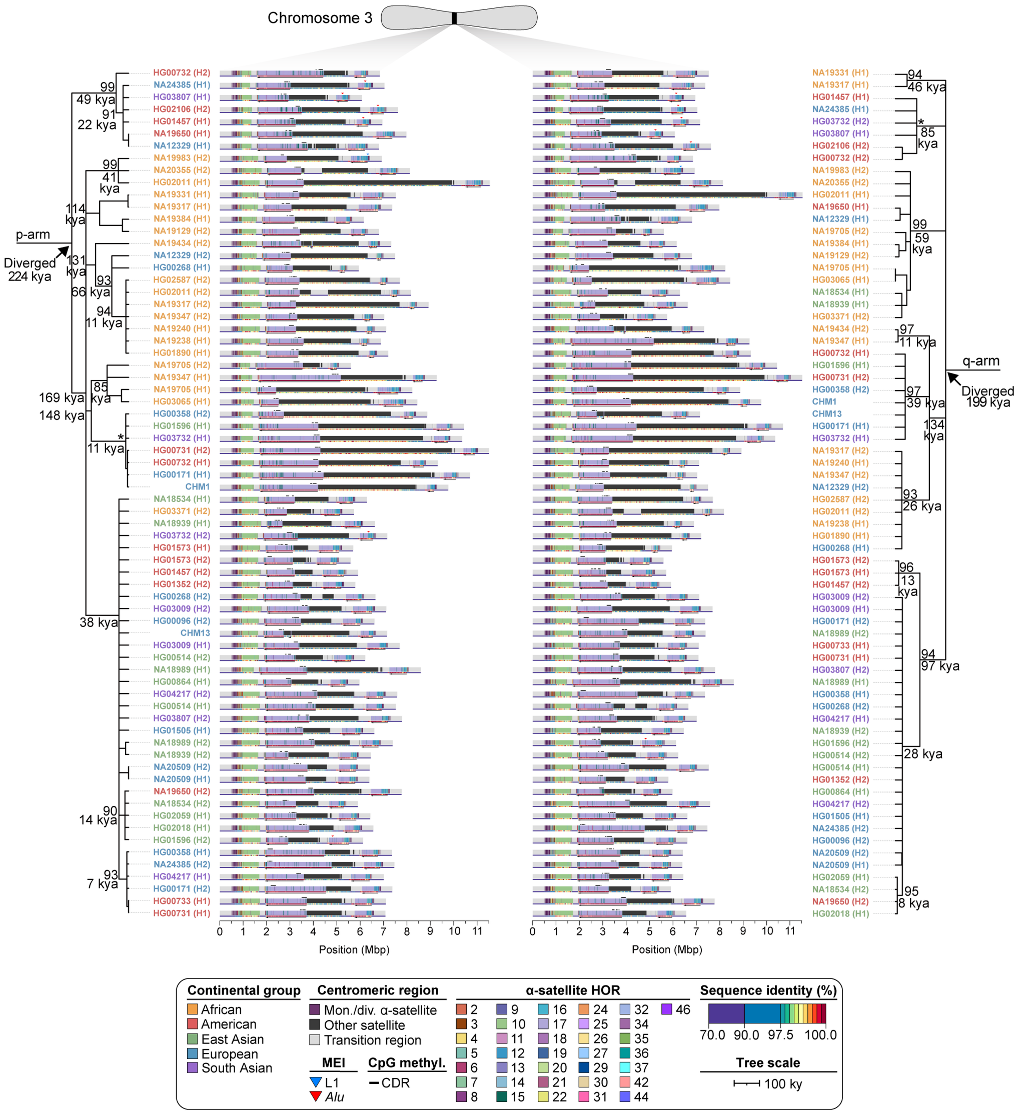


**Supplementary Figure 34. Phylogenetic reconstruction of chromosome 3 centromere haplotypes.** Maximum-likelihood phylogenetic tree depicting the p- and q-arm topologies along with estimated divergence times for the chromosome 3 (*D3Z1*) α-satellite HOR arrays from diverse human genomes. Asterisks indicate nodes with 100% bootstrap support, and nodes with 90-99% bootstrap support are indicated numerically. Nodes without an asterisk or number have bootstrap support <90%. We note that most differences in the order of the haplotypes occur at the terminal branches where the order of sequence taxa can be readily reshuffled to establish near-complete concordance. Thus, there are no significant changes in the overall topologies of the phylogenetic tree.

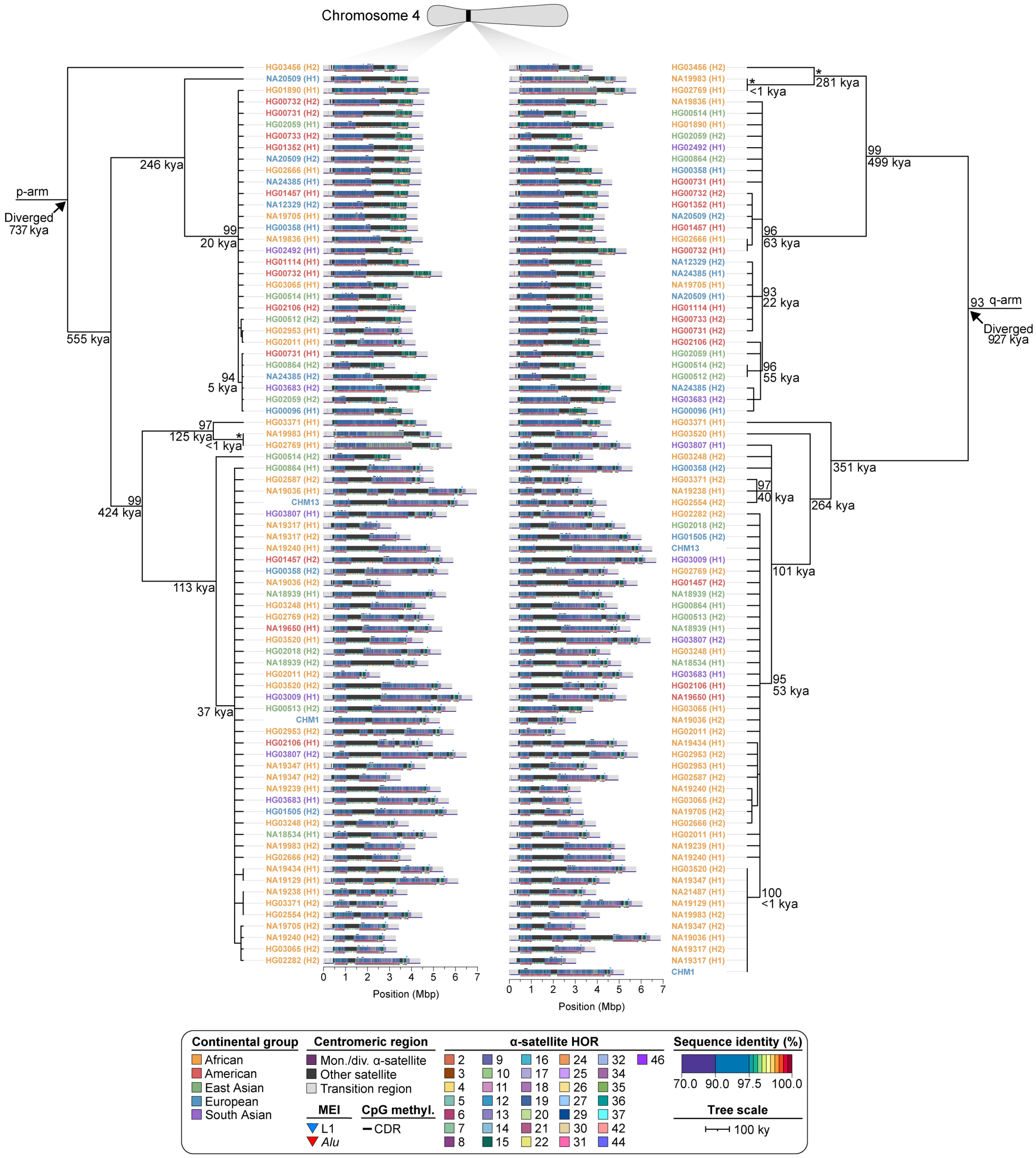


**Supplementary Figure 35. Phylogenetic reconstruction of chromosome 4 centromere haplotypes.** Maximum-likelihood phylogenetic tree depicting the p- and q-arm topologies along with estimated divergence times for the chromosome 4 (*D4Z1*) α-satellite HOR arrays from diverse human genomes. Asterisks indicate nodes with 100% bootstrap support, and nodes with 90-99% bootstrap support are indicated numerically. Nodes without an asterisk or number have bootstrap support <90%. We note that most differences in the order of the haplotypes occur at the terminal branches where the order of sequence taxa can be readily reshuffled to establish near-complete concordance. Thus, there are no significant changes in the overall topologies of the phylogenetic tree.

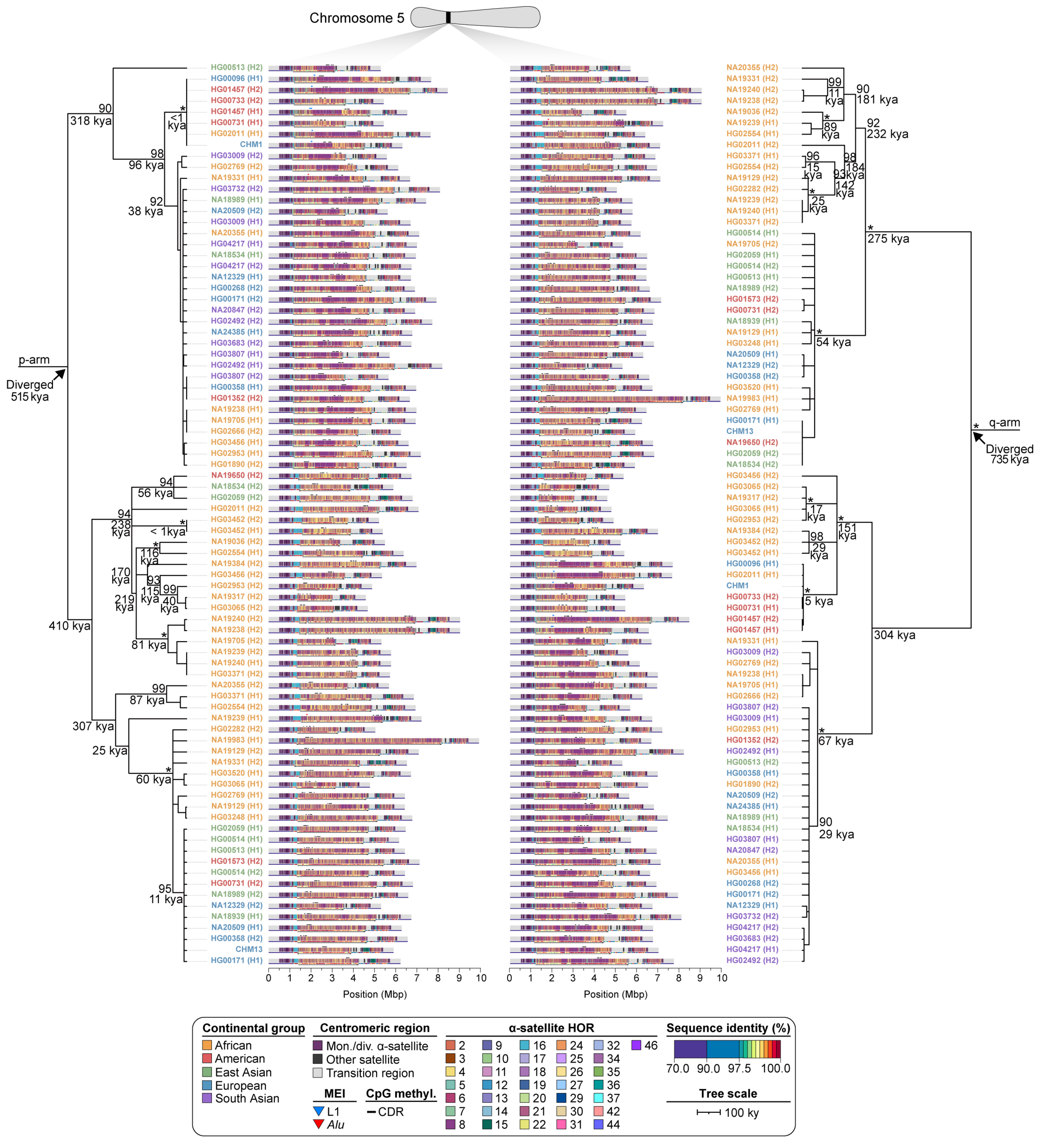


**Supplementary Figure 36. Phylogenetic reconstruction of chromosome 5 centromere haplotypes.** Maximum-likelihood phylogenetic tree depicting the p- and q-arm topologies along with estimated divergence times for the chromosome 5 (*D5Z2*) α-satellite HOR arrays from diverse human genomes. Asterisks indicate nodes with 100% bootstrap support, and nodes with 90-99% bootstrap support are indicated numerically. Nodes without an asterisk or number have bootstrap support <90%. We note that most differences in the order of the haplotypes occur at the terminal branches where the order of sequence taxa can be readily reshuffled to establish near-complete concordance. Thus, there are no significant changes in the overall topologies of the phylogenetic tree.

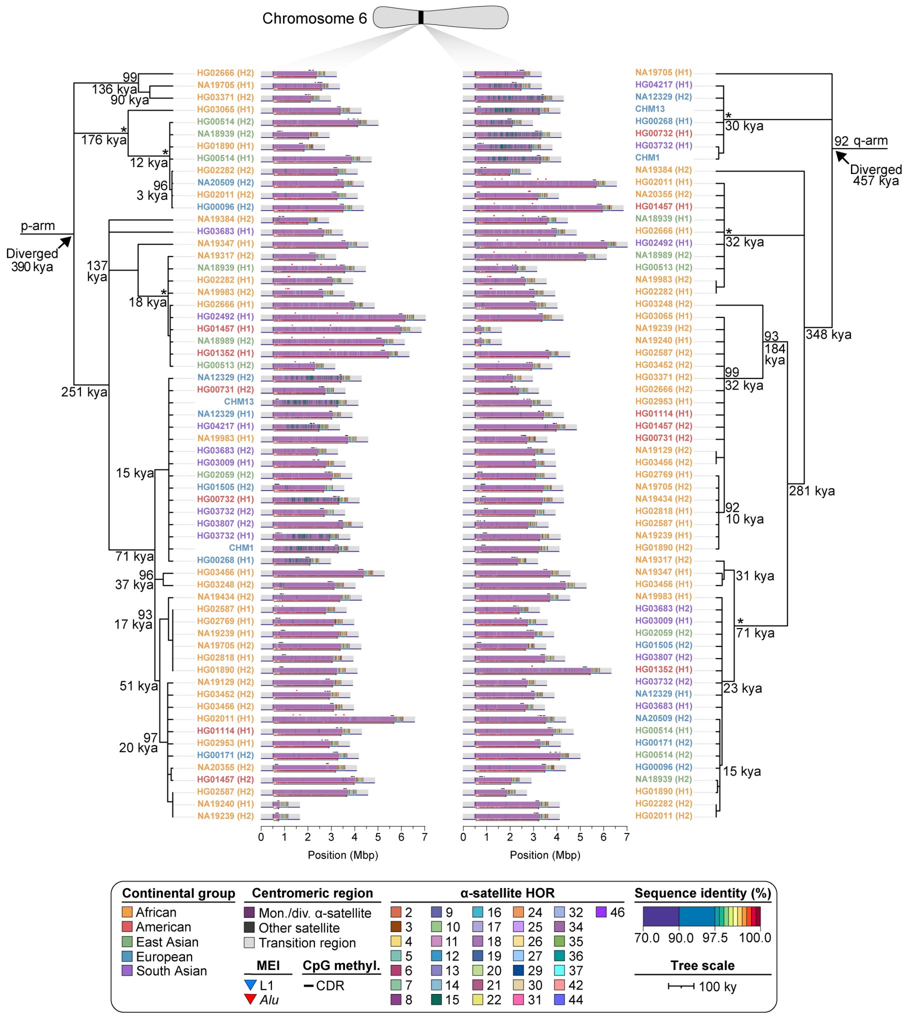


**Supplementary Figure 37. Phylogenetic reconstruction of chromosome 6 centromere haplotypes.** Maximum-likelihood phylogenetic tree depicting the p- and q-arm topologies along with estimated divergence times for the chromosome 6 (*D6Z1*) α-satellite HOR arrays from diverse human genomes. Asterisks indicate nodes with 100% bootstrap support, and nodes with 90-99% bootstrap support are indicated numerically. Nodes without an asterisk or number have bootstrap support <90%. We note that most differences in the order of the haplotypes occur at the terminal branches where the order of sequence taxa can be readily reshuffled to establish near-complete concordance. Thus, there are no significant changes in the overall topologies of the phylogenetic tree.

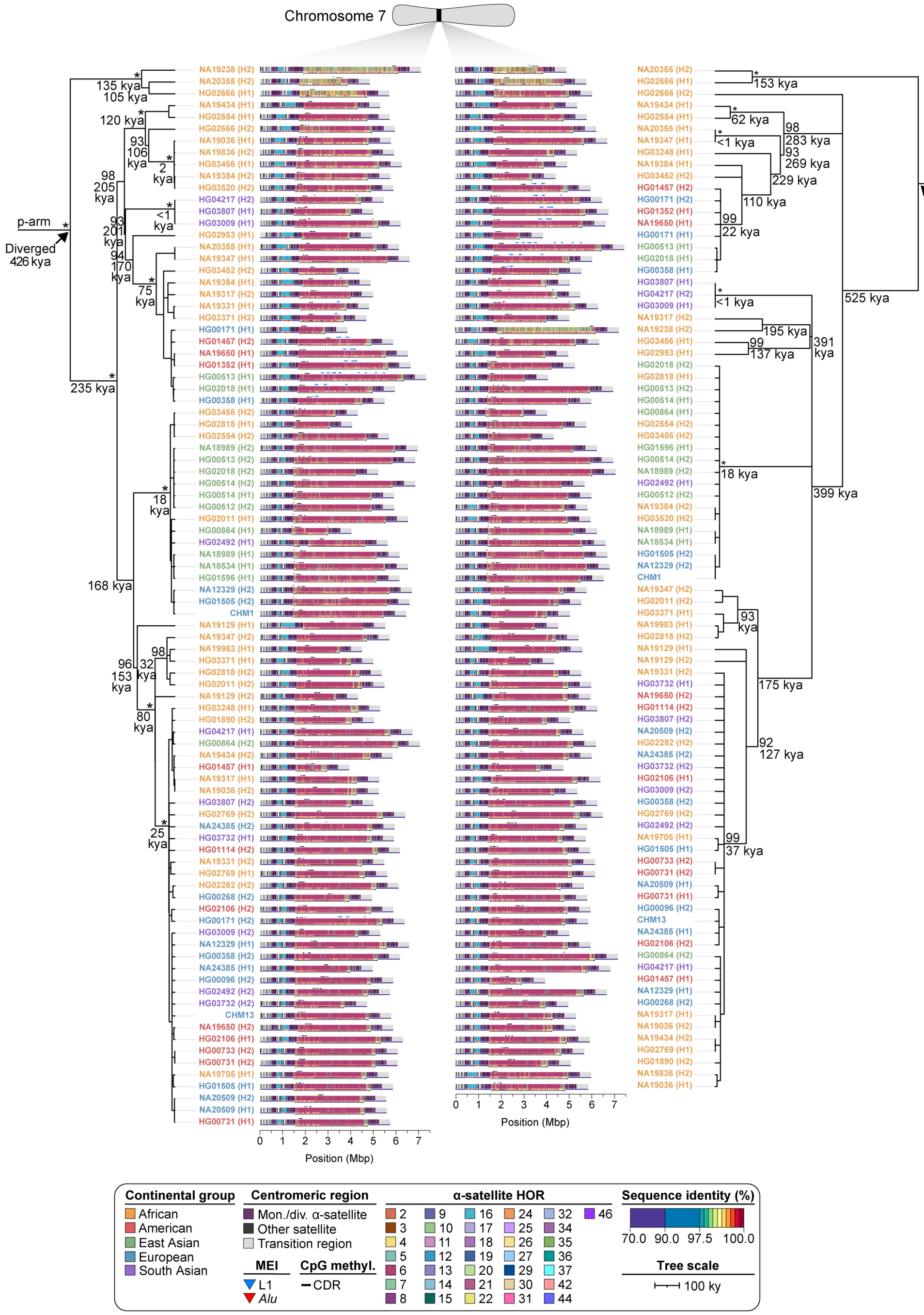

**Supplementary Figure 38. Phylogenetic reconstruction of chromosome 7 centromere haplotypes.** Maximum-likelihood phylogenetic tree depicting the p- and q-arm topologies along with estimated divergence times for the chromosome 7 (*D7Z1*) α-satellite HOR arrays from diverse human genomes. Asterisks indicate nodes with 100% bootstrap support, and nodes with 90-99% bootstrap support are indicated numerically. Nodes without an asterisk or number have bootstrap support <90%. We note that most differences in the order of the haplotypes occur at the terminal branches where the order of sequence taxa can be readily reshuffled to establish near-complete concordance. Thus, there are no significant changes in the overall topologies of the phylogenetic tree.

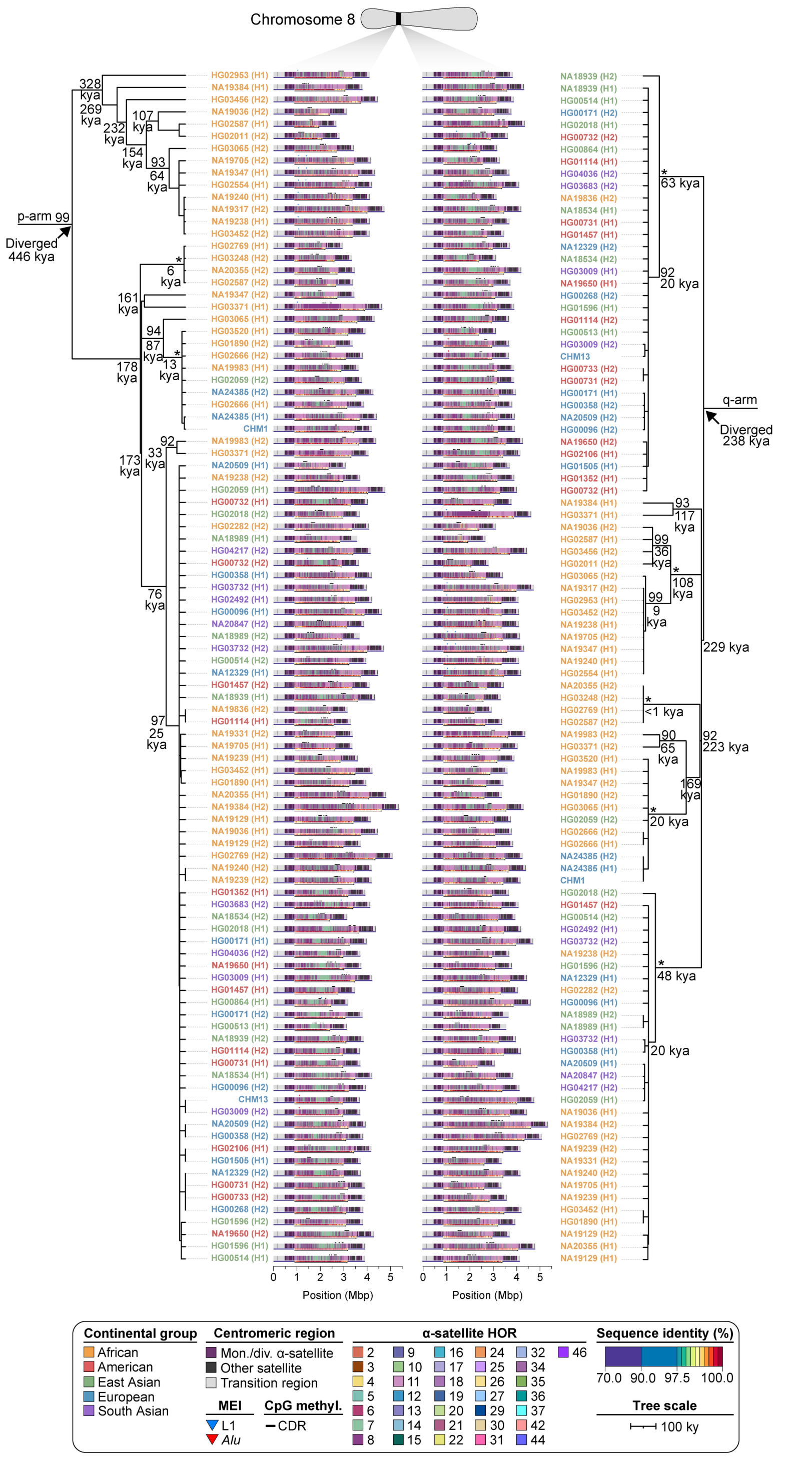


**Supplementary Figure 39. Phylogenetic reconstruction of chromosome 8 centromere haplotypes.** Maximum-likelihood phylogenetic tree depicting the p- and q-arm topologies along with estimated divergence times for the chromosome 8 (*D8Z2*) α-satellite HOR arrays from diverse human genomes. Asterisks indicate nodes with 100% bootstrap support, and nodes with 90-99% bootstrap support are indicated numerically. Nodes without an asterisk or number have bootstrap support <90%. We note that most differences in the order of the haplotypes occur at the terminal branches where the order of sequence taxa can be readily reshuffled to establish near-complete concordance. Thus, there are no significant changes in the overall topologies of the phylogenetic tree.

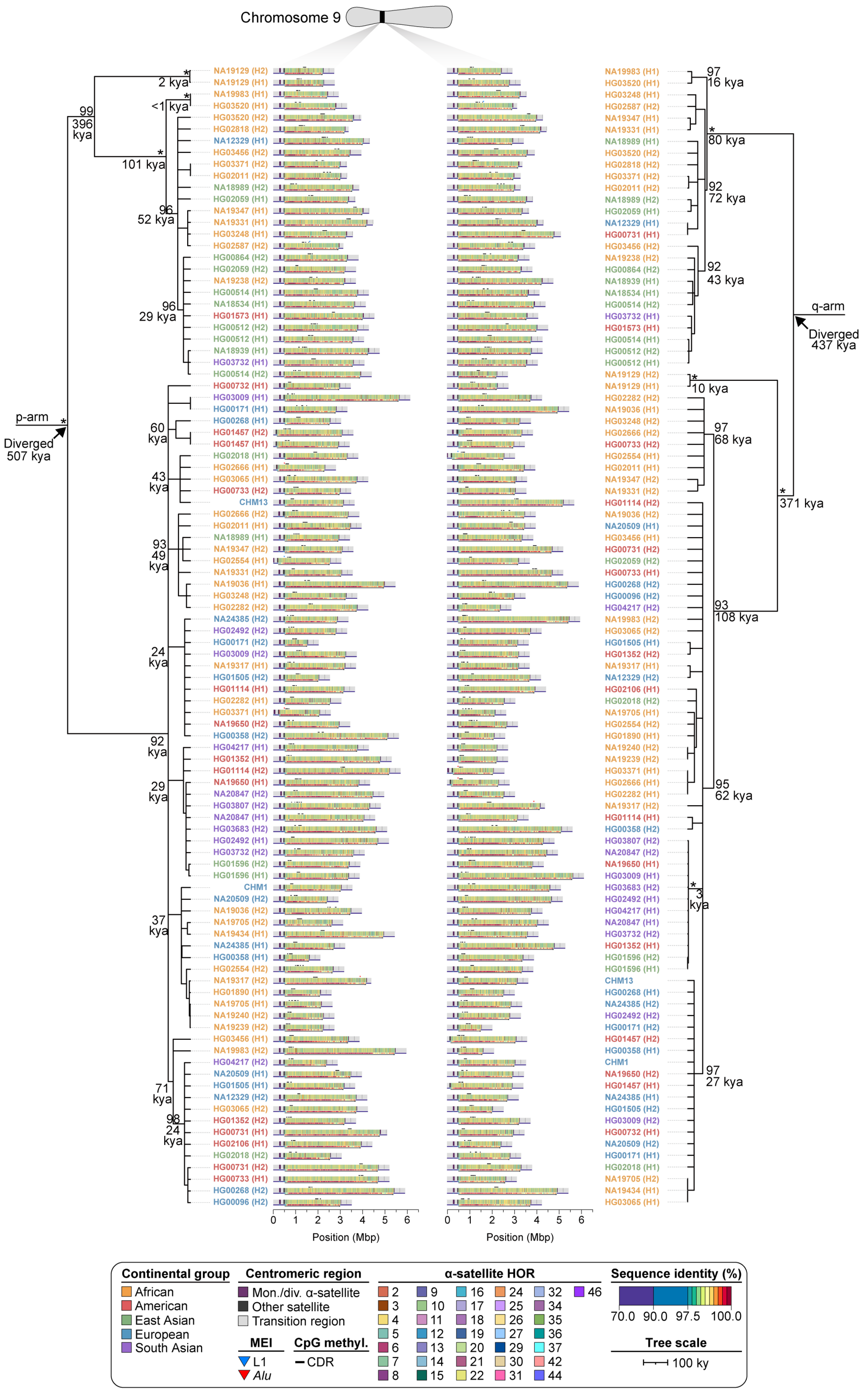


**Supplementary Figure 40. Phylogenetic reconstruction of chromosome 9 centromere haplotypes.** Maximum-likelihood phylogenetic tree depicting the p- and q-arm topologies along with estimated divergence times for the chromosome 9 (*D9Z4*) α-satellite HOR arrays from diverse human genomes. Asterisks indicate nodes with 100% bootstrap support, and nodes with 90-99% bootstrap support are indicated numerically. Nodes without an asterisk or number have bootstrap support <90%. We note that most differences in the order of the haplotypes occur at the terminal branches where the order of sequence taxa can be readily reshuffled to establish near-complete concordance. Thus, there are no significant changes in the overall topologies of the phylogenetic tree.

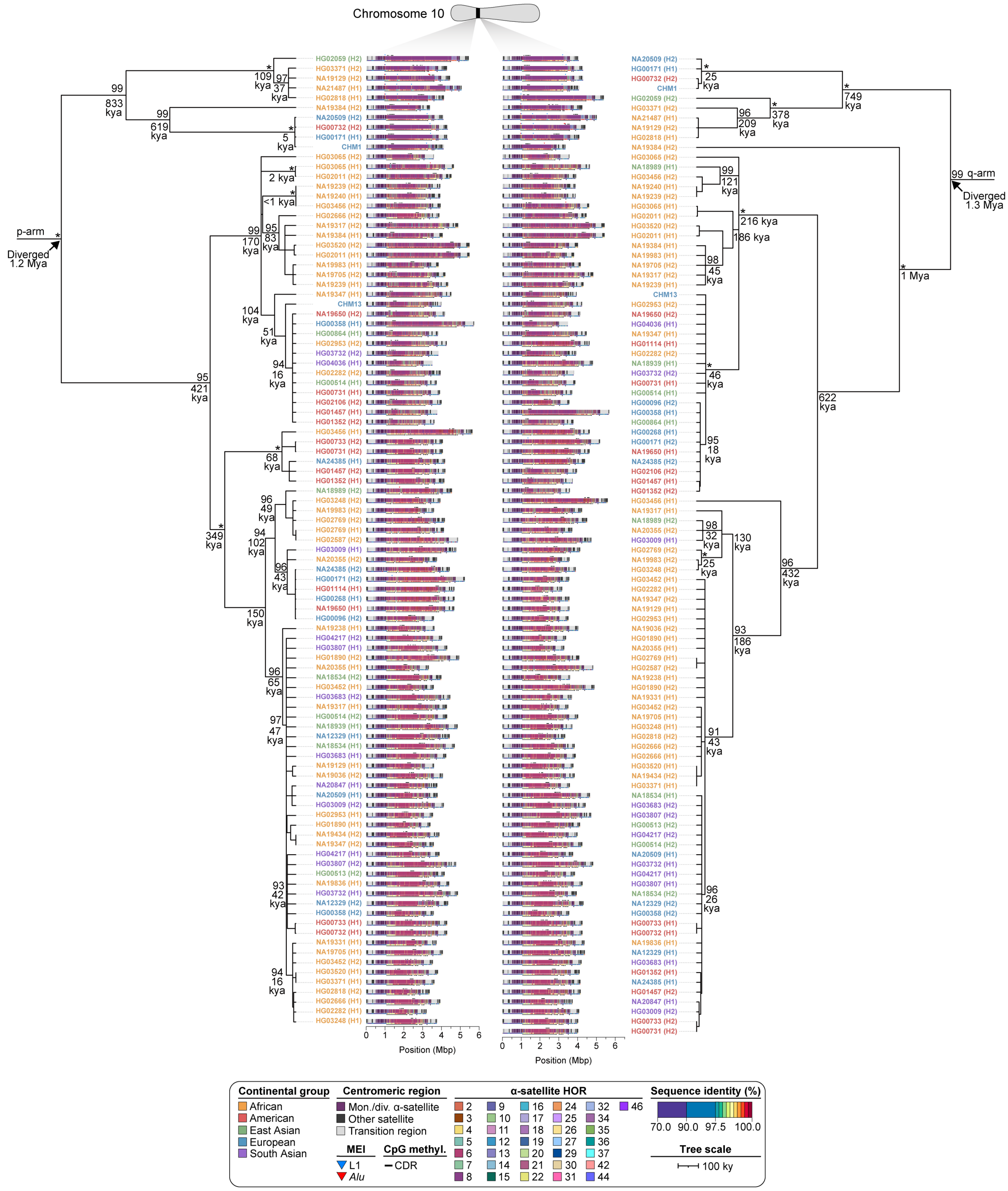

**Supplementary Figure 41. Phylogenetic reconstruction of chromosome 10 centromere haplotypes.** Maximum-likelihood phylogenetic tree depicting the p- and q-arm topologies along with estimated divergence times for the chromosome 10 (*D10Z1*) α-satellite HOR arrays from diverse human genomes. Asterisks indicate nodes with 100% bootstrap support, and nodes with 90-99% bootstrap support are indicated numerically. Nodes without an asterisk or number have bootstrap support <90%. We note that most differences in the order of the haplotypes occur at the terminal branches where the order of sequence taxa can be readily reshuffled to establish near-complete concordance. Thus, there are no significant changes in the overall topologies of the phylogenetic tree.

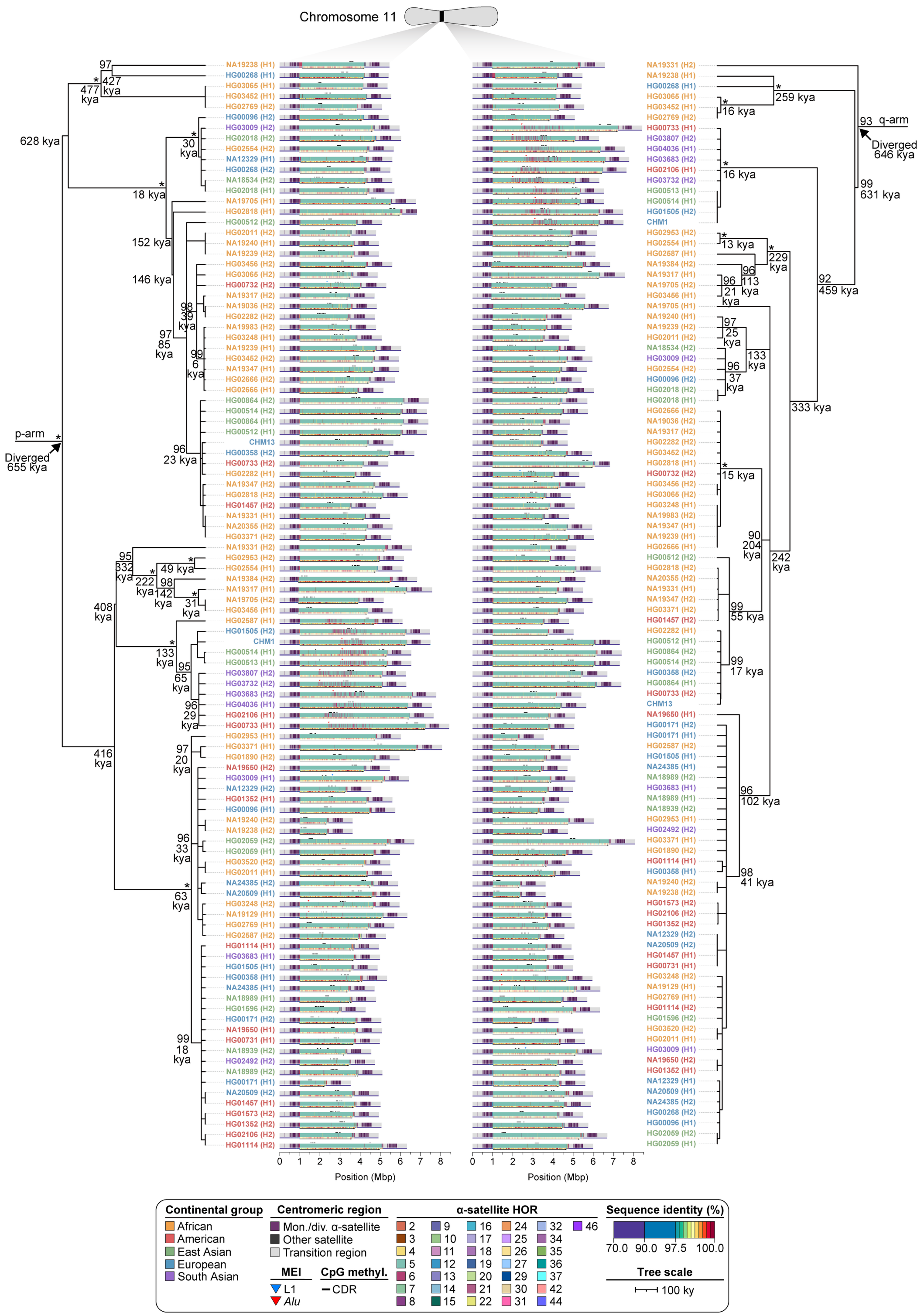


**Supplementary Figure 42. Phylogenetic reconstruction of chromosome 11 centromere haplotypes.** Maximum-likelihood phylogenetic tree depicting the p- and q-arm topologies along with estimated divergence times for the chromosome 11 (*D11Z1*) α-satellite HOR arrays from diverse human genomes. Asterisks indicate nodes with 100% bootstrap support, and nodes with 90-99% bootstrap support are indicated numerically. Nodes without an asterisk or number have bootstrap support <90%. We note that most differences in the order of the haplotypes occur at the terminal branches where the order of sequence taxa can be readily reshuffled to establish near-complete concordance. Thus, there are no significant changes in the overall topologies of the phylogenetic tree.

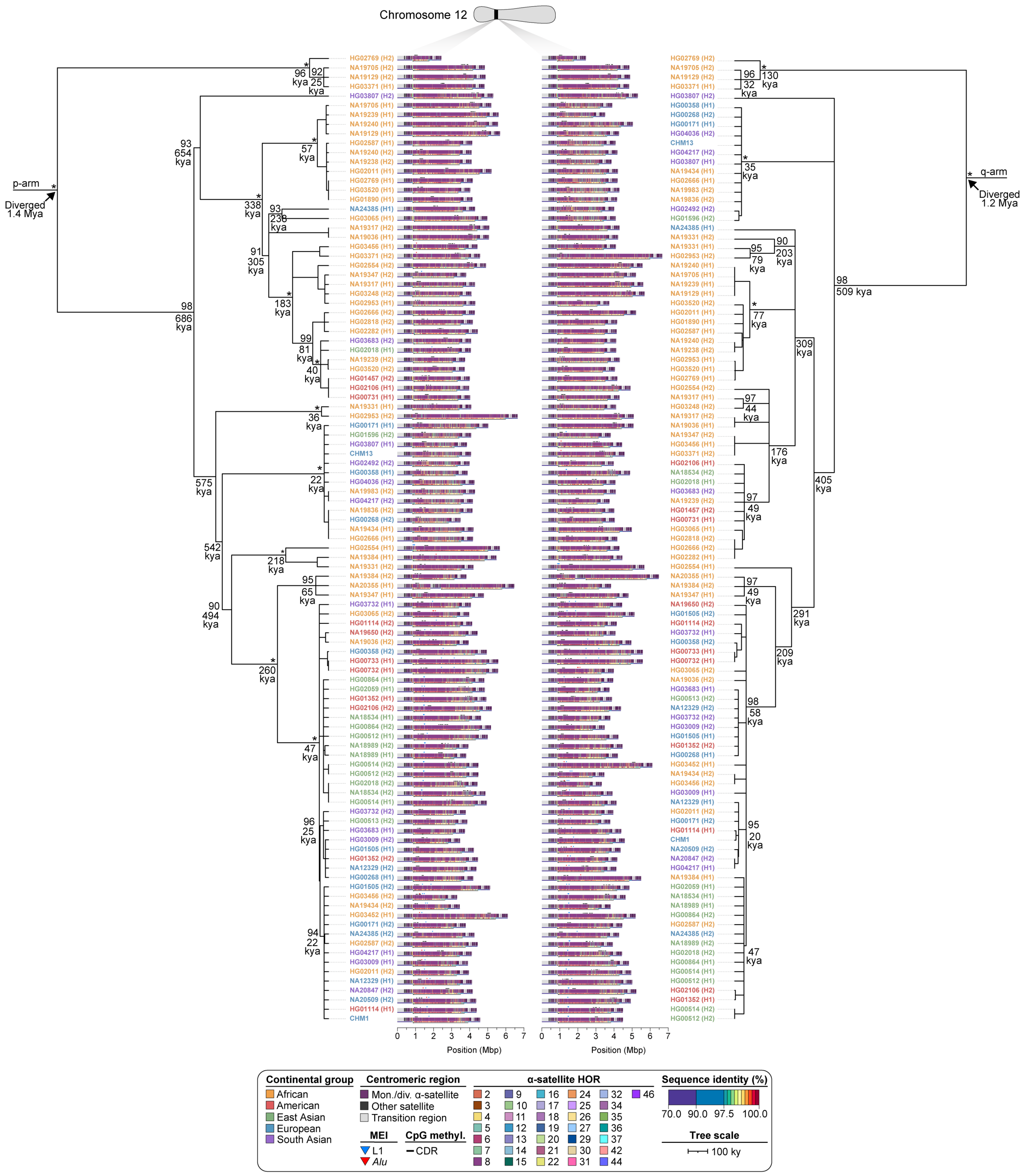


**Supplementary Figure 43. Phylogenetic reconstruction of chromosome 12 centromere haplotypes.** Maximum-likelihood phylogenetic tree depicting the p- and q-arm topologies along with estimated divergence times for the chromosome 12 (*D12Z3*) α-satellite HOR arrays from diverse human genomes. Asterisks indicate nodes with 100% bootstrap support, and nodes with 90-99% bootstrap support are indicated numerically. Nodes without an asterisk or number have bootstrap support <90%. We note that most differences in the order of the haplotypes occur at the terminal branches where the order of sequence taxa can be readily reshuffled to establish near-complete concordance. Thus, there are no significant changes in the overall topologies of the phylogenetic tree.

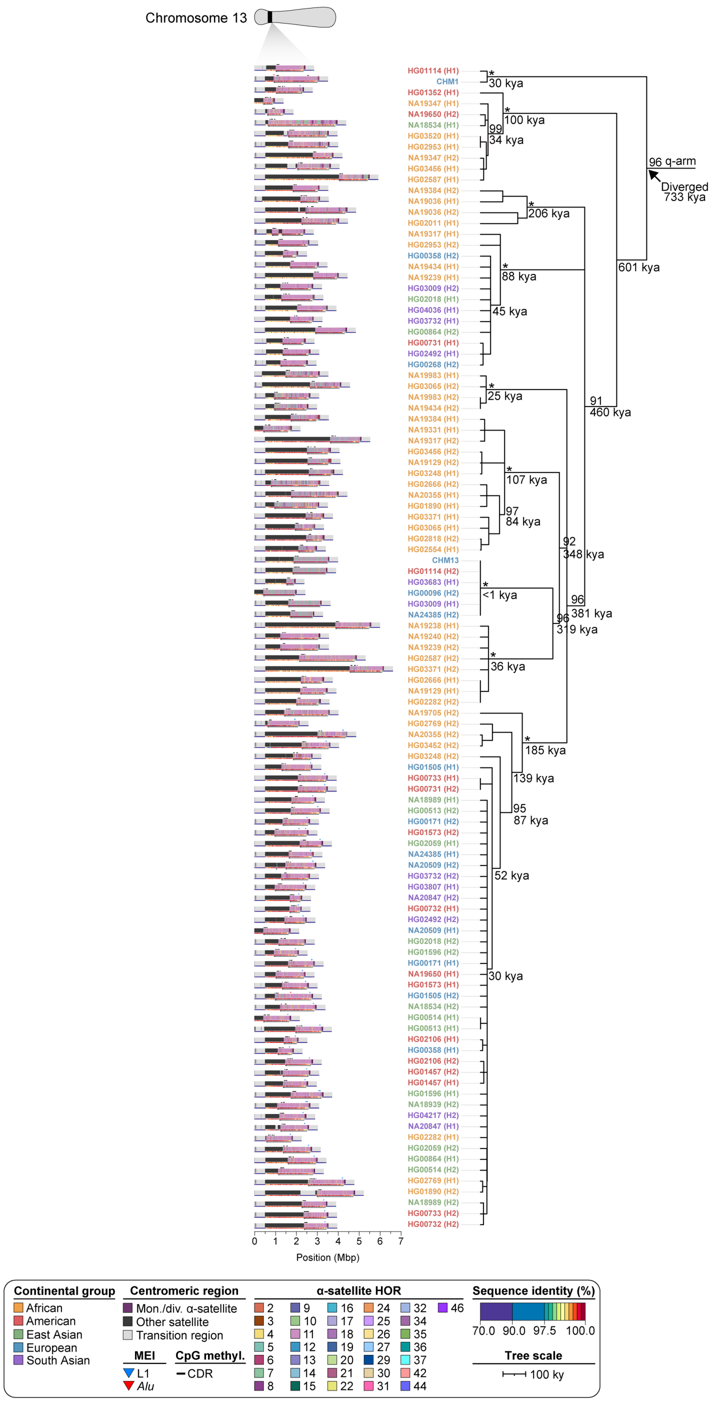


**Supplementary Figure 44. Phylogenetic reconstruction of chromosome 13 centromere haplotypes.** Maximum-likelihood phylogenetic tree depicting the q-arm topology along with estimated divergence times for the chromosome 13 (*D13Z1*) α-satellite HOR arrays from diverse human genomes. The phylogenetic tree from the p arm does not preserve the overall topology of the tree due to the known homologous recombination between acrocentric p arms[9], so only the q-arm phylogenetic tree is shown. Asterisks indicate nodes with 100% bootstrap support, and nodes with 90-99% bootstrap support are indicated numerically. Nodes without an asterisk or number have bootstrap support <90%. We note that most differences in the order of the haplotypes occur at the terminal branches where the order of sequence taxa can be readily reshuffled to establish near-complete concordance. Thus, there are no significant changes in the overall topologies of the phylogenetic tree.

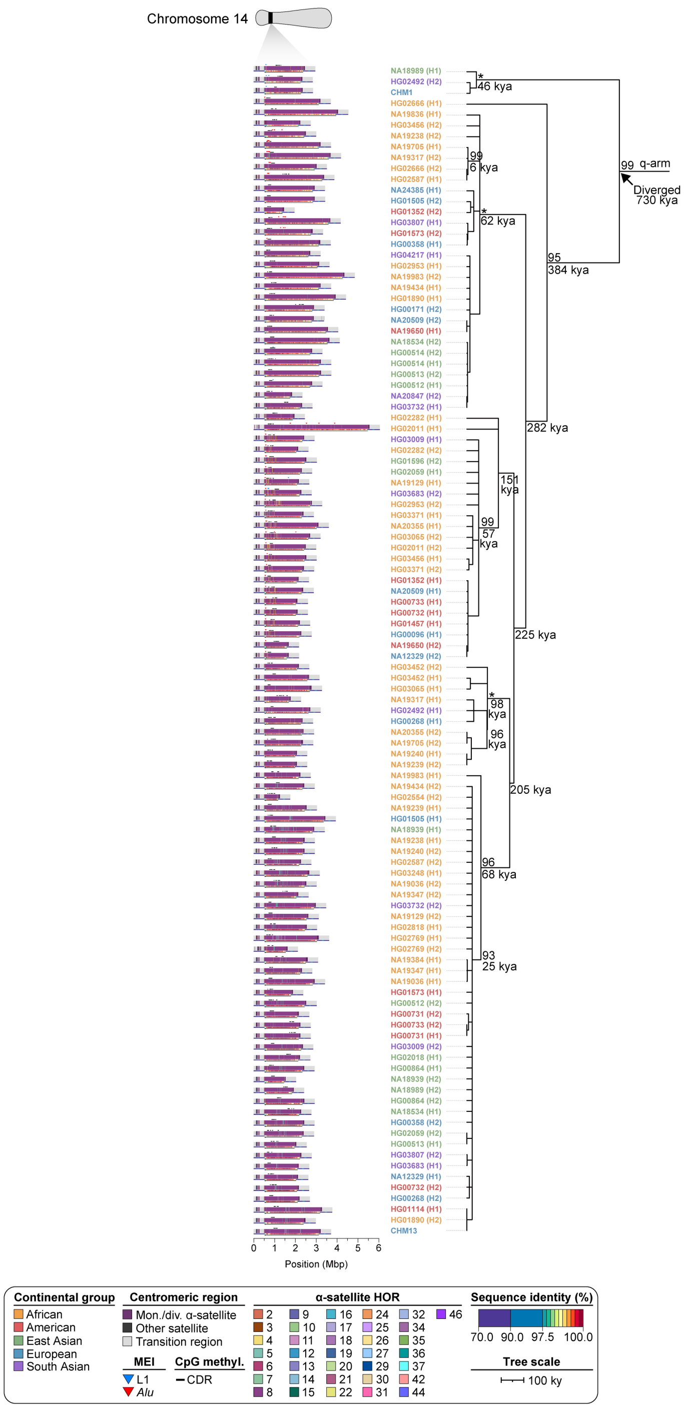


**Supplementary Figure 45. Phylogenetic reconstruction of chromosome 14 centromere haplotypes.** Maximum-likelihood phylogenetic tree depicting the q-arm topology along with estimated divergence times for the chromosome 14 (*D14Z9*) α-satellite HOR arrays from diverse human genomes. The phylogenetic tree from the p arm does not preserve the overall topology of the tree due to the known homologous recombination between acrocentric p arms[9], so only the q-arm phylogenetic tree is shown. Asterisks indicate nodes with 100% bootstrap support, and nodes with 90-99% bootstrap support are indicated numerically. Nodes without an asterisk or number have bootstrap support <90%. We note that most differences in the order of the haplotypes occur at the terminal branches where the order of sequence taxa can be readily reshuffled to establish near-complete concordance. Thus, there are no significant changes in the overall topologies of the phylogenetic tree.

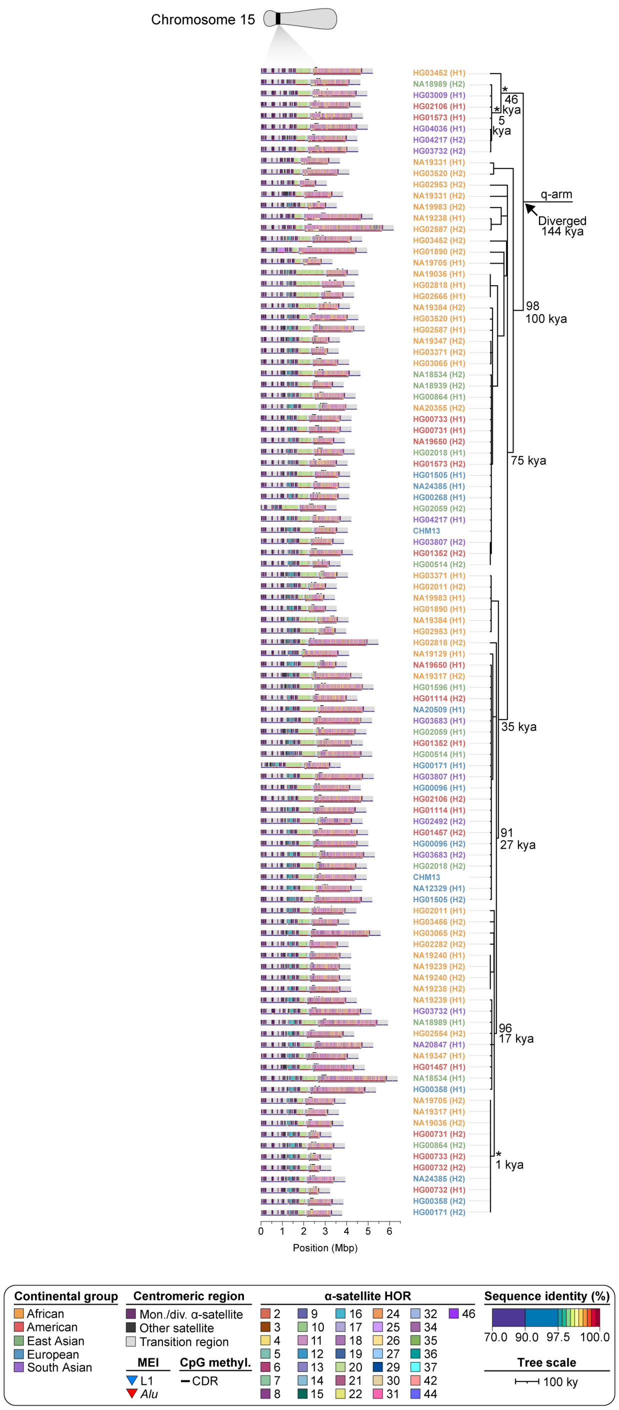


**Supplementary Figure 46. Phylogenetic reconstruction of chromosome 15 centromere haplotypes.** Maximum-likelihood phylogenetic tree depicting the q-arm topology along with estimated divergence times for the chromosome 15 (*D15Z3*) α-satellite HOR arrays from diverse human genomes. The phylogenetic tree from the p arm does not preserve the overall topology of the tree due to the known homologous recombination between acrocentric p arms[9], so only the q-arm phylogenetic tree is shown. Asterisks indicate nodes with 100% bootstrap support, and nodes with 90-99% bootstrap support are indicated numerically. Nodes without an asterisk or number have bootstrap support <90%. We note that most differences in the order of the haplotypes occur at the terminal branches where the order of sequence taxa can be readily reshuffled to establish near-complete concordance. Thus, there are no significant changes in the overall topologies of the phylogenetic tree.

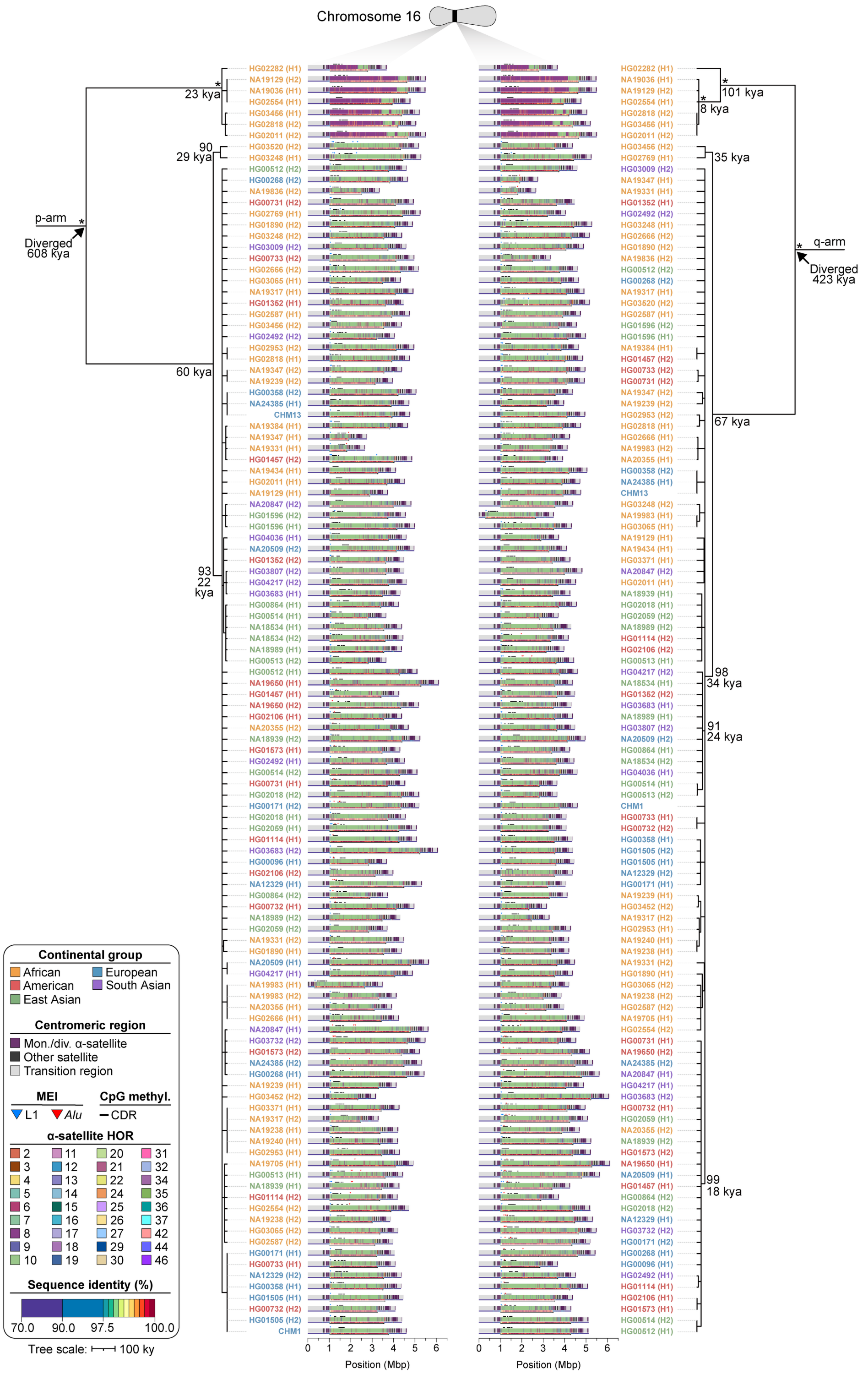


**Supplementary Figure 47. Phylogenetic reconstruction of chromosome 16 centromere haplotypes.** Maximum-likelihood phylogenetic tree depicting the p- and q-arm topologies along with estimated divergence times for the chromosome 16 (*D16Z2*) α-satellite HOR arrays from diverse human genomes. Asterisks indicate nodes with 100% bootstrap support, and nodes with 90-99% bootstrap support are indicated numerically. Nodes without an asterisk or number have bootstrap support <90%. We note that most differences in the order of the haplotypes occur at the terminal branches where the order of sequence taxa can be readily reshuffled to establish near-complete concordance. Thus, there are no significant changes in the overall topologies of the phylogenetic tree.

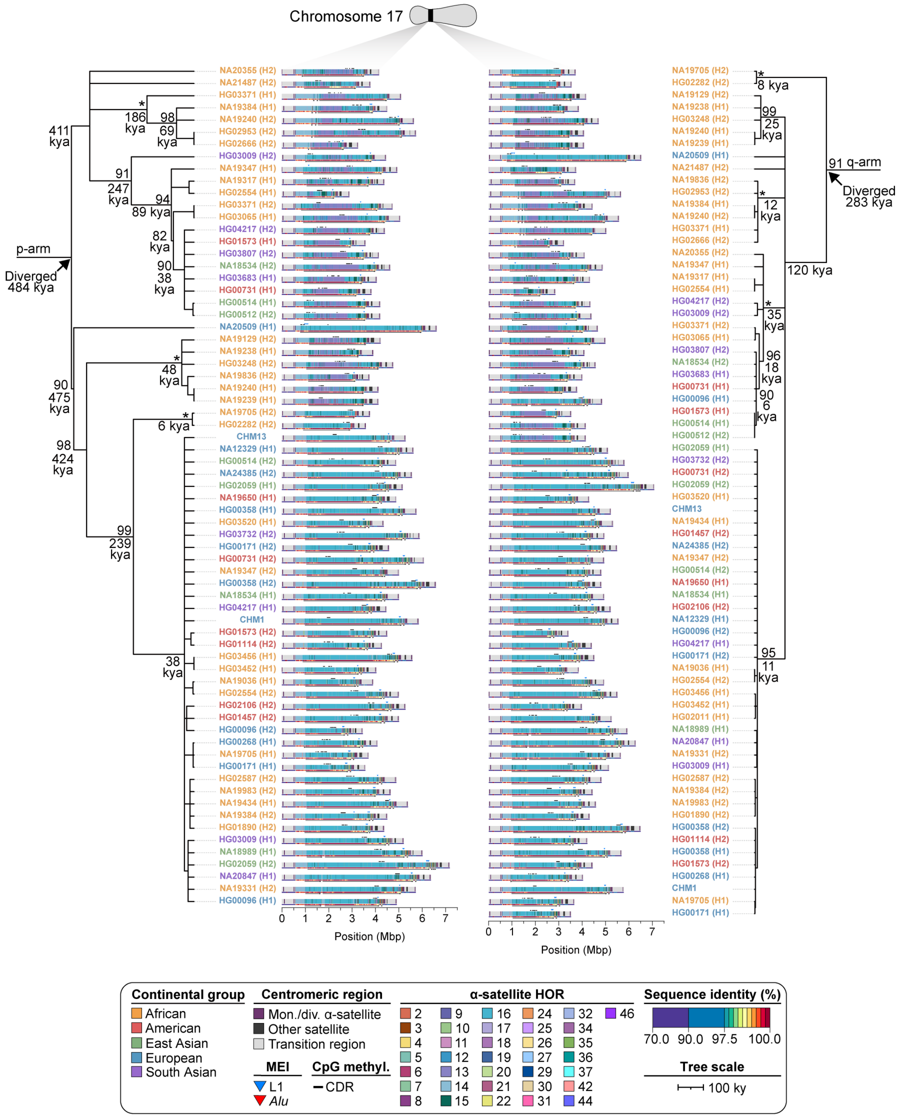


**Supplementary Figure 48. Phylogenetic reconstruction of chromosome 17 centromere haplotypes.** Maximum-likelihood phylogenetic tree depicting the p- and q-arm topologies along with estimated divergence times for the chromosome 17 (*D17Z1*) α-satellite HOR arrays from diverse human genomes. Asterisks indicate nodes with 100% bootstrap support, and nodes with 90-99% bootstrap support are indicated numerically. Nodes without an asterisk or number have bootstrap support <90%. We note that most differences in the order of the haplotypes occur at the terminal branches where the order of sequence taxa can be readily reshuffled to establish near-complete concordance. Thus, there are no significant changes in the overall topologies of the phylogenetic tree.

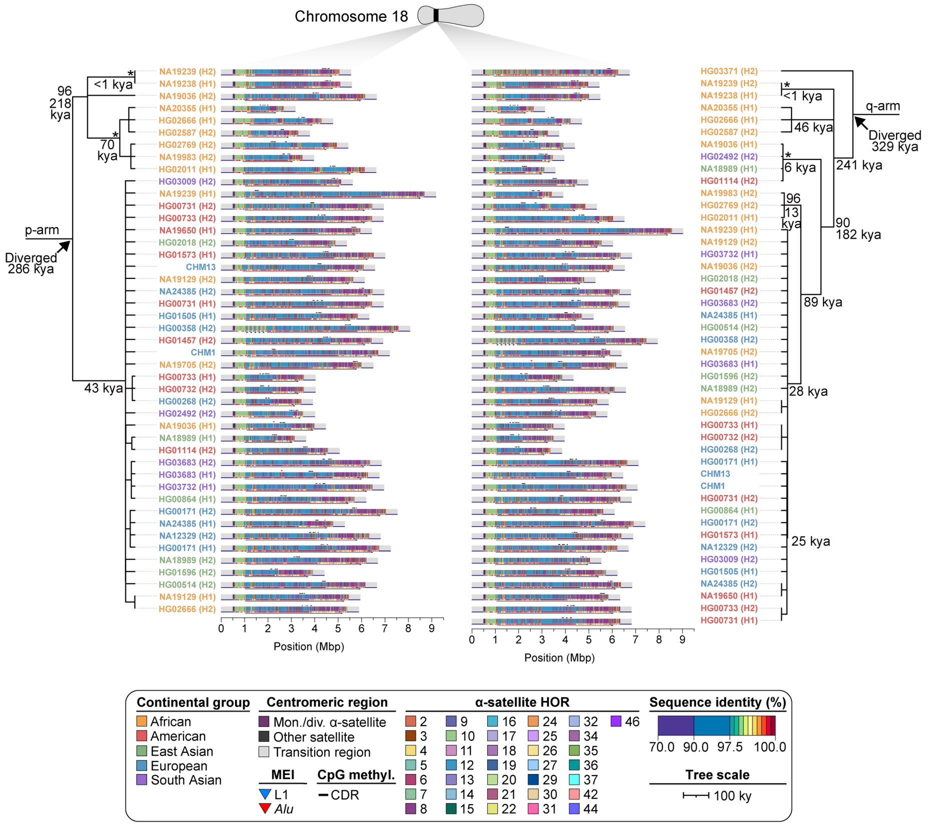


**Supplementary Figure 49. Phylogenetic reconstruction of chromosome 18 centromere haplotypes.** Maximum-likelihood phylogenetic tree depicting the p- and q-arm topologies along with estimated divergence times for the chromosome 18 (*D18Z1*) α-satellite HOR arrays from diverse human genomes. Asterisks indicate nodes with 100% bootstrap support, and nodes with 90-99% bootstrap support are indicated numerically. Nodes without an asterisk or number have bootstrap support <90%. We note that most differences in the order of the haplotypes occur at the terminal branches where the order of sequence taxa can be readily reshuffled to establish near-complete concordance. Thus, there are no significant changes in the overall topologies of the phylogenetic tree.

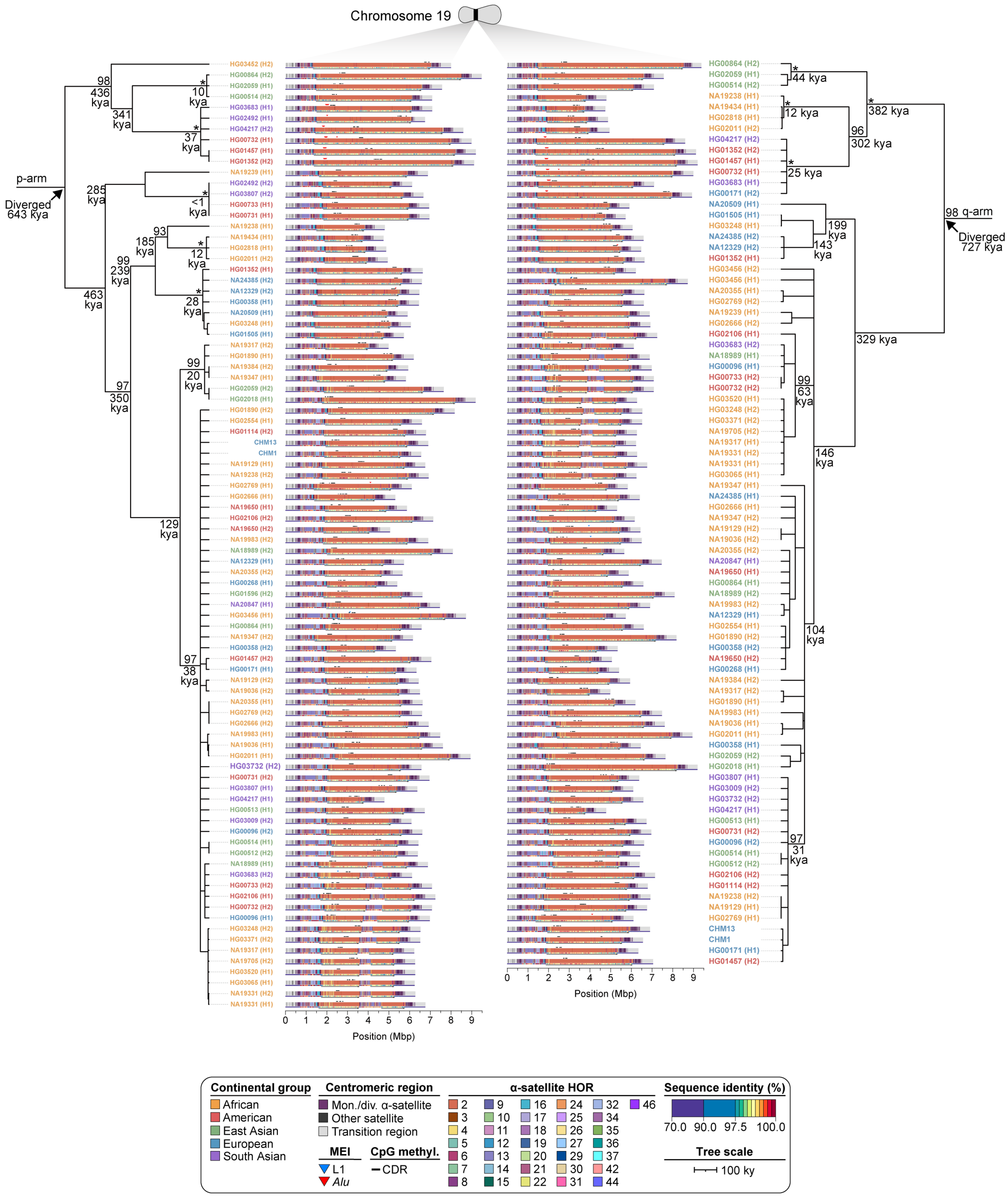


**Supplementary Figure 50. Phylogenetic reconstruction of chromosome 19 centromere haplotypes.** Maximum-likelihood phylogenetic tree depicting the p- and q-arm topologies along with estimated divergence times for the chromosome 19 (*D19Z3*) α-satellite HOR arrays from diverse human genomes. Asterisks indicate nodes with 100% bootstrap support, and nodes with 90-99% bootstrap support are indicated numerically. Nodes without an asterisk or number have bootstrap support <90%. We note that most differences in the order of the haplotypes occur at the terminal branches where the order of sequence taxa can be readily reshuffled to establish near-complete concordance. Thus, there are no significant changes in the overall topologies of the phylogenetic tree.

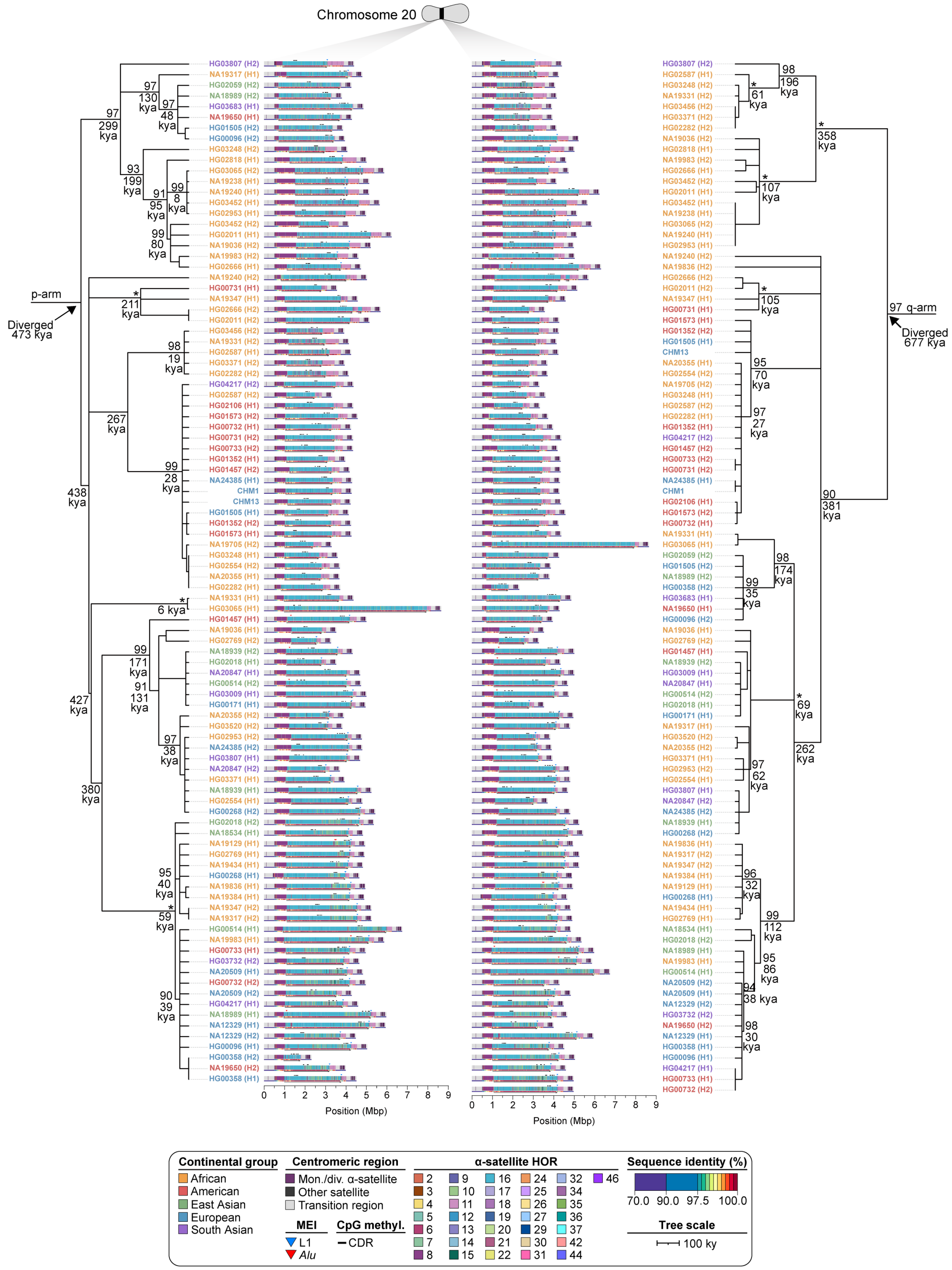


**Supplementary Figure 51. Phylogenetic reconstruction of chromosome 20 centromere haplotypes.** Maximum-likelihood phylogenetic tree depicting the p- and q-arm topologies along with estimated divergence times for the chromosome 20 (*D20Z2*) α-satellite HOR arrays from diverse human genomes. Asterisks indicate nodes with 100% bootstrap support, and nodes with 90-99% bootstrap support are indicated numerically. Nodes without an asterisk or number have bootstrap support <90%. We note that most differences in the order of the haplotypes occur at the terminal branches where the order of sequence taxa can be readily reshuffled to establish near-complete concordance. Thus, there are no significant changes in the overall topologies of the phylogenetic tree.

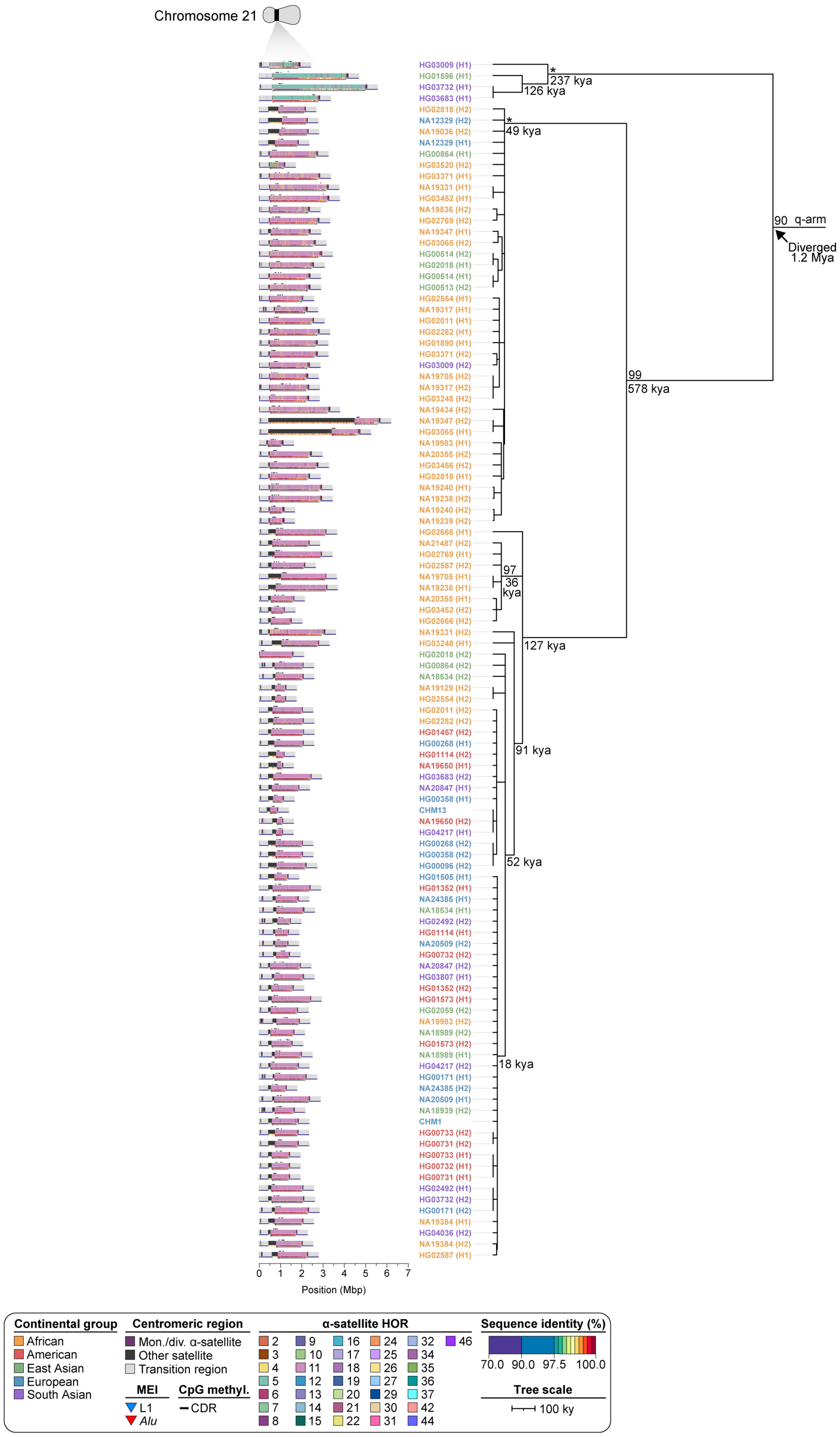


**Supplementary Figure 52. Phylogenetic reconstruction of chromosome 21 centromere haplotypes.** Maximum-likelihood phylogenetic tree depicting the q-arm topology along with estimated divergence times for the chromosome 21 (*D21Z1*) α-satellite HOR arrays from diverse human genomes. The phylogenetic tree from the p arm does not preserve the overall topology of the tree due to the known homologous recombination between acrocentric p arms[9], so only the q-arm phylogenetic tree is shown. Asterisks indicate nodes with 100% bootstrap support, and nodes with 90-99% bootstrap support are indicated numerically. Nodes without an asterisk or number have bootstrap support <90%. We note that most differences in the order of the haplotypes occur at the terminal branches where the order of sequence taxa can be readily reshuffled to establish near-complete concordance. Thus, there are no significant changes in the overall topologies of the phylogenetic tree.

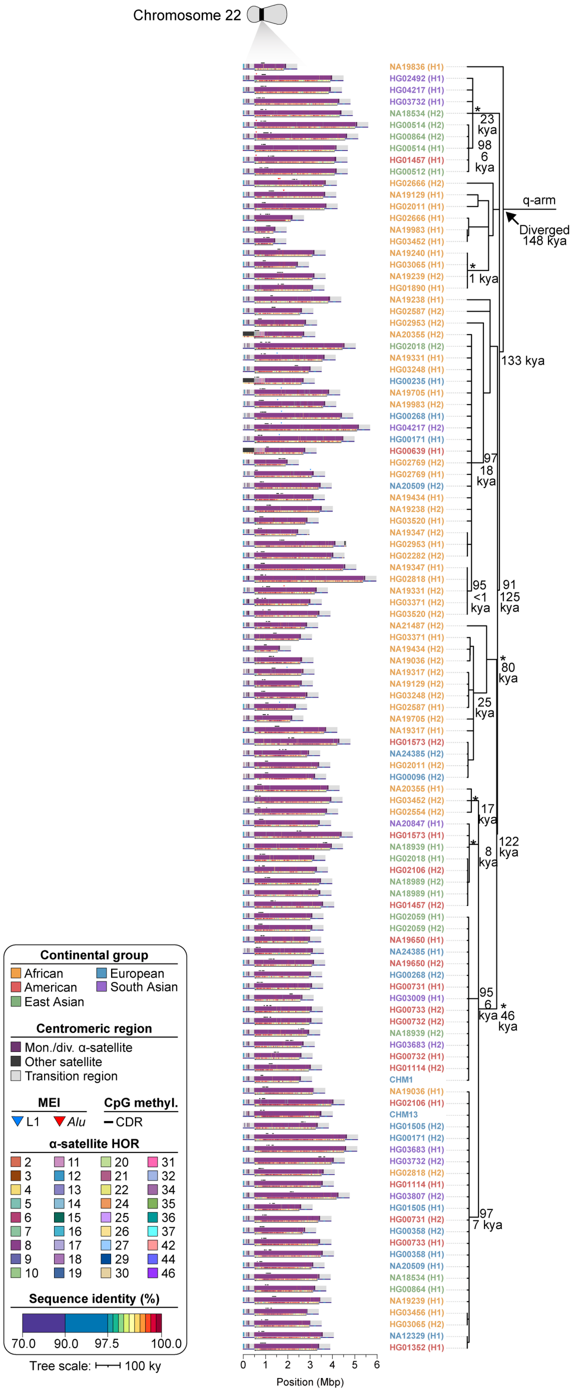


**Supplementary Figure 53. Phylogenetic reconstruction of chromosome 22 centromere haplotypes.** Maximum-likelihood phylogenetic tree depicting the q-arm topology along with estimated divergence times for the chromosome 22 (*D22Z1*) α-satellite HOR arrays from diverse human genomes. The phylogenetic tree from the p arm does not preserve the overall topology of the tree due to the known homologous recombination between acrocentric p arms[9], so only the q-arm phylogenetic tree is shown. Asterisks indicate nodes with 100% bootstrap support, and nodes with 90-99% bootstrap support are indicated numerically. Nodes without an asterisk or number have bootstrap support <90%. We note that most differences in the order of the haplotypes occur at the terminal branches where the order of sequence taxa can be readily reshuffled to establish near-complete concordance. Thus, there are no significant changes in the overall topologies of the phylogenetic tree.

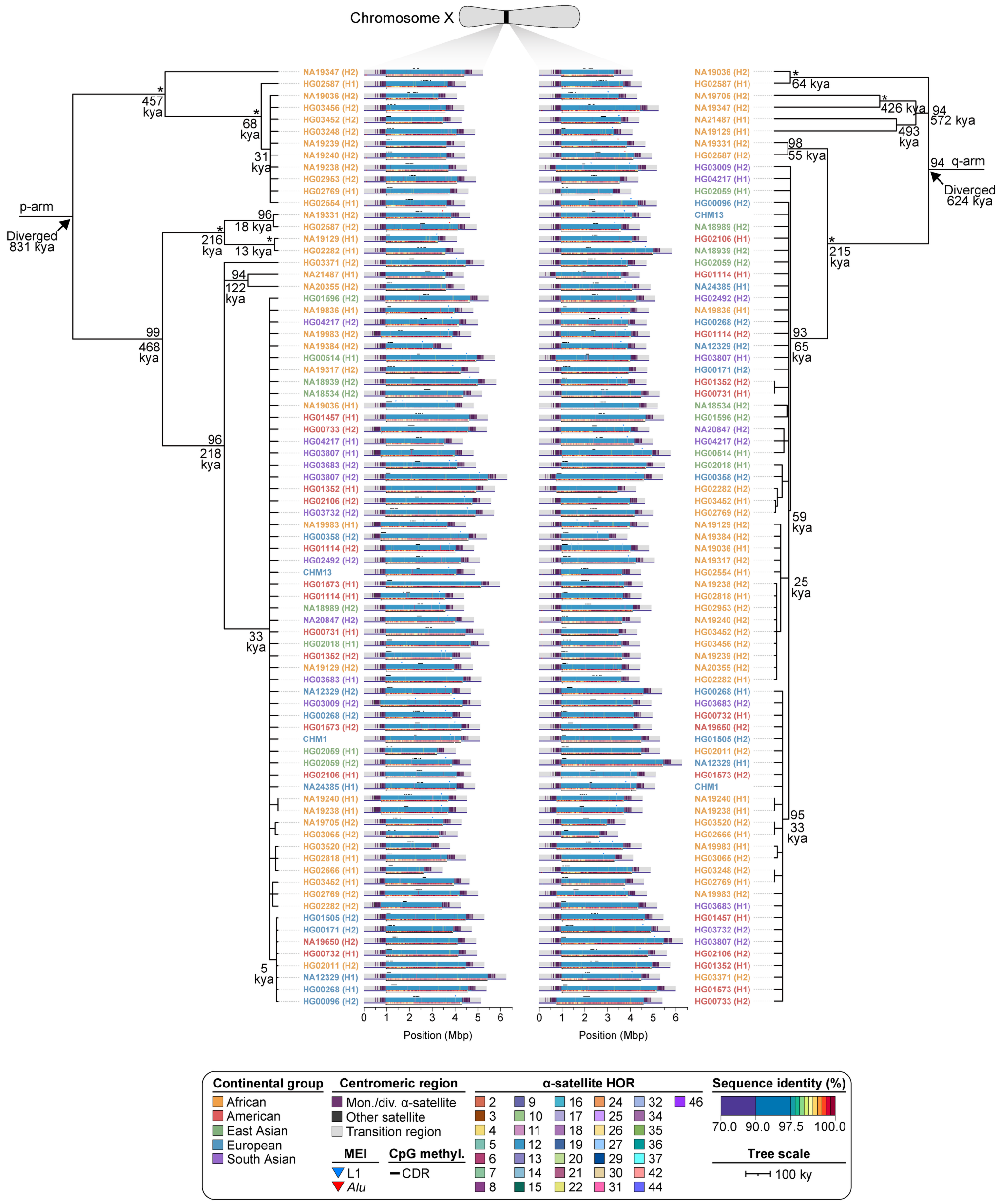


**Supplementary Figure 54. Phylogenetic reconstruction of chromosome X centromere haplotypes.** Maximum-likelihood phylogenetic tree depicting the p- and q-arm topologies along with estimated divergence times for the chromosome X (*DXZ1*) α-satellite HOR arrays from diverse human genomes. Asterisks indicate nodes with 100% bootstrap support, and nodes with 90-99% bootstrap support are indicated numerically. Nodes without an asterisk or number have bootstrap support <90%. We note that most differences in the order of the haplotypes occur at the terminal branches where the order of sequence taxa can be readily reshuffled to establish near-complete concordance. Thus, there are no significant changes in the overall topologies of the phylogenetic tree.

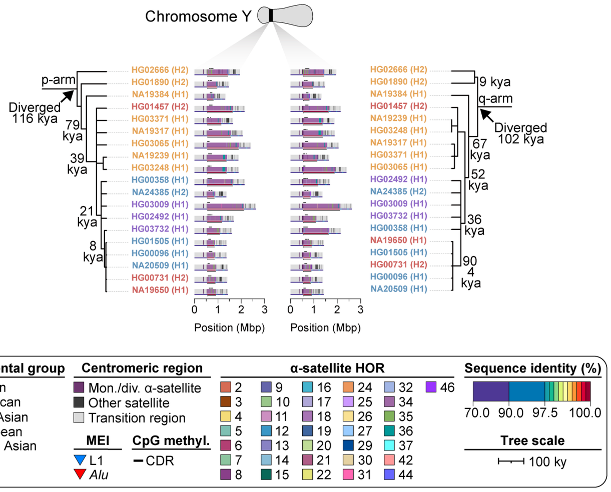


**Supplementary Figure 55. Phylogenetic reconstruction of chromosome Y centromere haplotypes.** Maximum-likelihood phylogenetic tree depicting the p- and q-arm topologies along with estimated divergence times for the chromosome Y (*DYZ3*) α-satellite HOR arrays from diverse human genomes. Asterisks indicate nodes with 100% bootstrap support, and nodes with 90-99% bootstrap support are indicated numerically. Nodes without an asterisk or number have bootstrap support <90%. We note that most differences in the order of the haplotypes occur at the terminal branches where the order of sequence taxa can be readily reshuffled to establish near-complete concordance. Thus, there are no significant changes in the overall topologies of the phylogenetic tree.

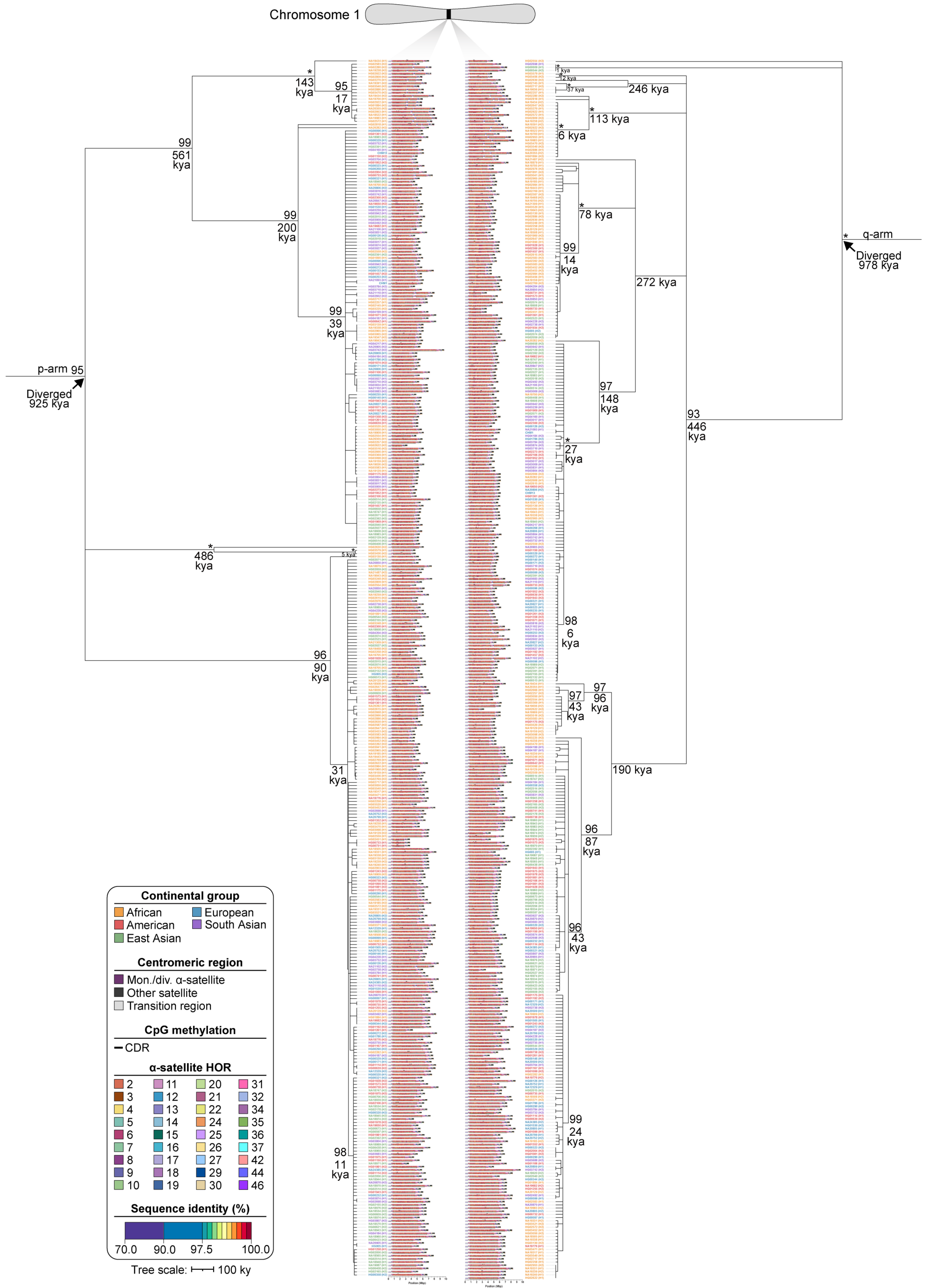


**Supplementary Figure 56. Phylogenetic reconstruction of chromosome 1 centromere haplotypes across the HGSVC and HPRC datasets.** Maximum-likelihood phylogenetic tree depicting the p- and q-arm topologies along with estimated divergence times for the chromosome 1 (*D1Z7*) α-satellite HOR arrays from 296 diverse human genomes. Asterisks indicate nodes with 100% bootstrap support, and nodes with 90-99% bootstrap support are indicated numerically. Nodes without an asterisk or number have bootstrap support <90%. The overall topology of the tree is preserved with the inclusion of complete centromeres recently assembled by the HPRC.

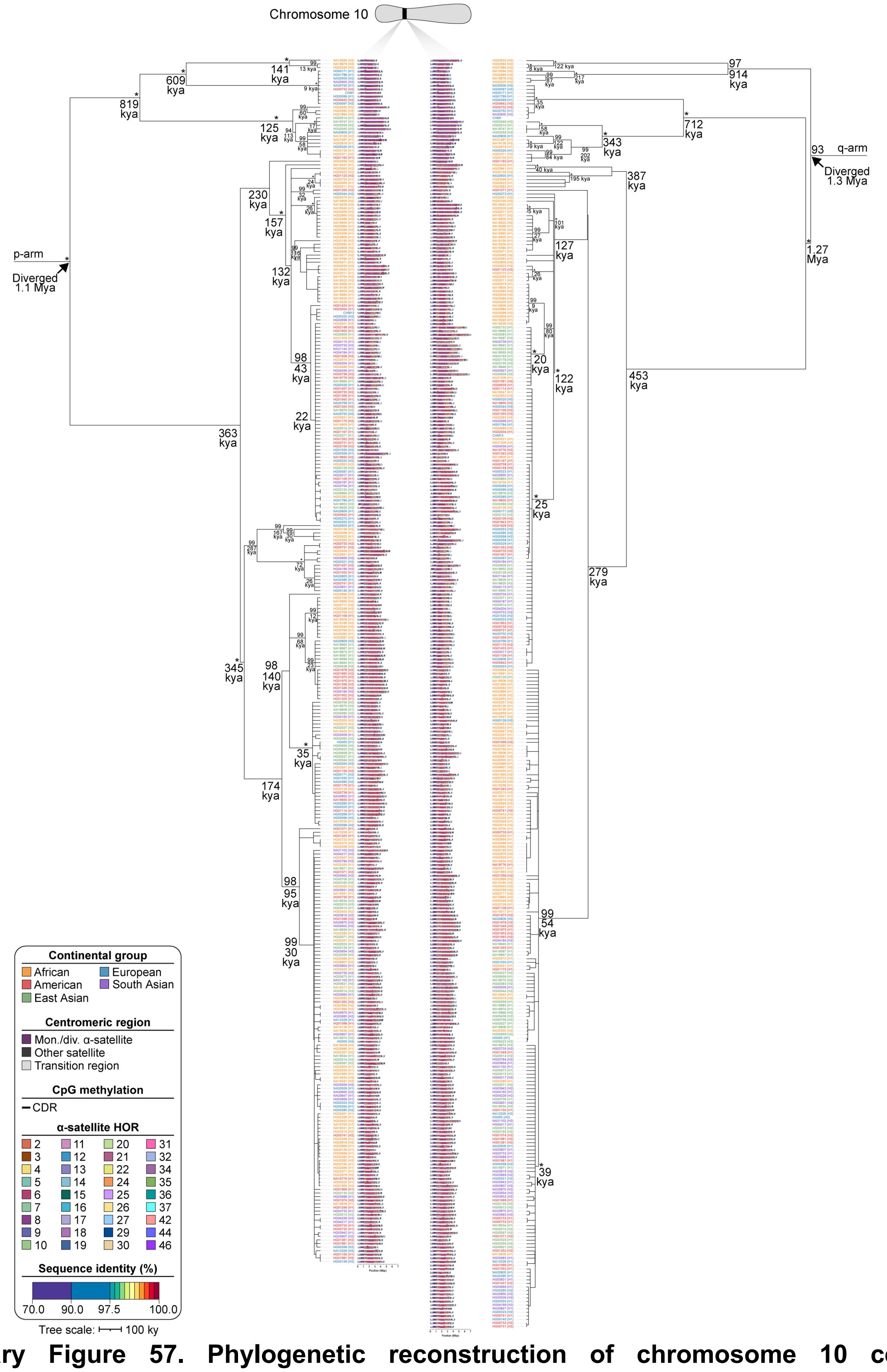


**Supplementary Figure 57. Phylogenetic reconstruction of chromosome 10 centromere haplotypes across the HGSVC and HPRC datasets.** Maximum-likelihood phylogenetic tree depicting the p- and q-arm topologies along with estimated divergence times for the chromosome 10 (*D10Z1*) α-satellite HOR arrays from 296 diverse human genomes. Asterisks indicate nodes with 100% bootstrap support, and nodes with 90-99% bootstrap support are indicated numerically. Nodes without an asterisk or number have bootstrap support <90%. The overall topology of the tree is preserved with the inclusion of complete centromeres recently assembled by the HPRC.

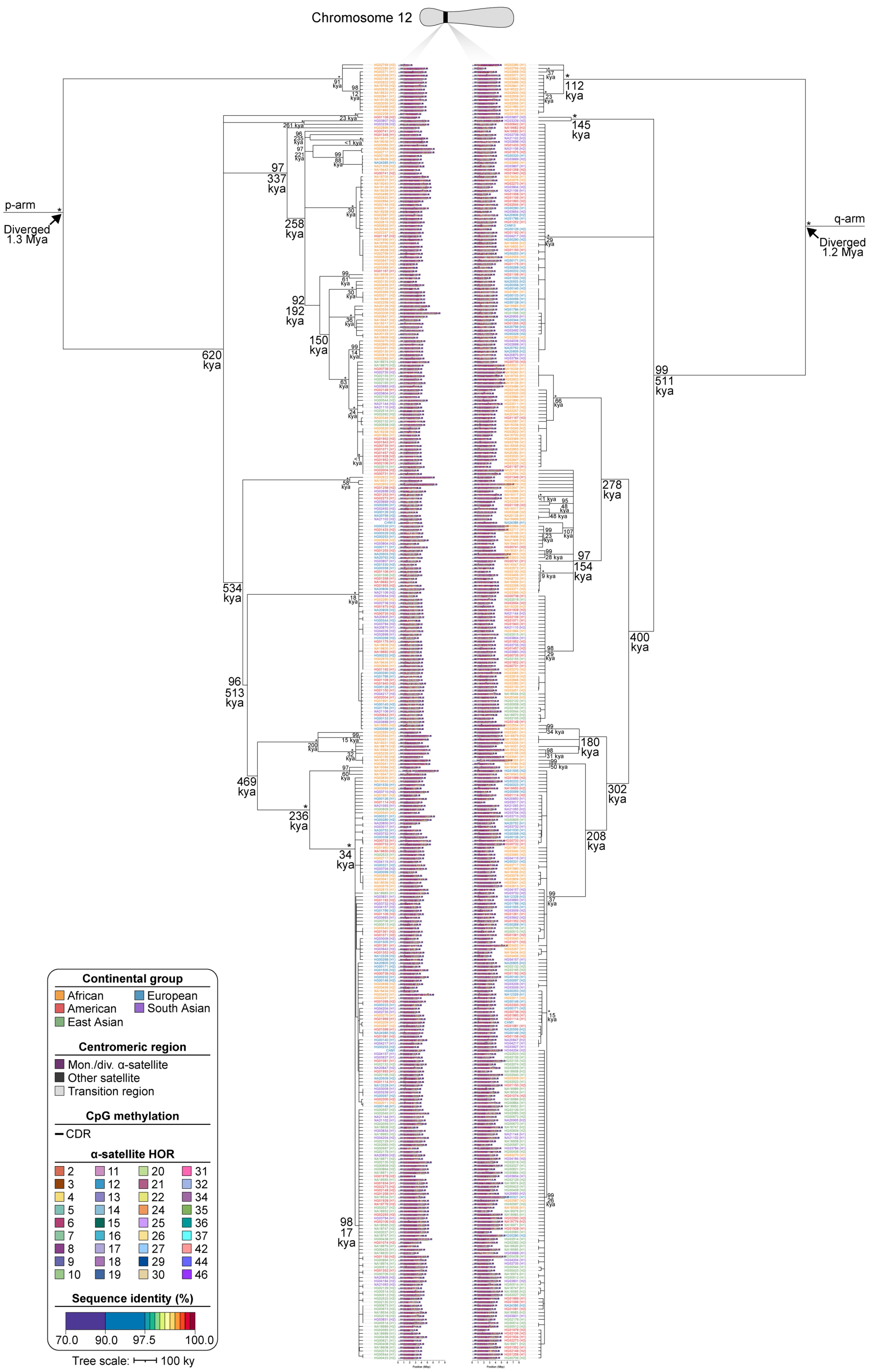


**Supplementary Figure 58. Phylogenetic reconstruction of chromosome 12 centromere haplotypes across the HGSVC and HPRC datasets.** Maximum-likelihood phylogenetic tree depicting the p- and q-arm topologies along with estimated divergence times for the chromosome 12 (*D12Z3*) α-satellite HOR arrays from 296 diverse human genomes. Asterisks indicate nodes with 100% bootstrap support, and nodes with 90-99% bootstrap support are indicated numerically. Nodes without an asterisk or number have bootstrap support <90%. The overall topology of the tree is preserved with the inclusion of complete centromeres recently assembled by the HPRC.

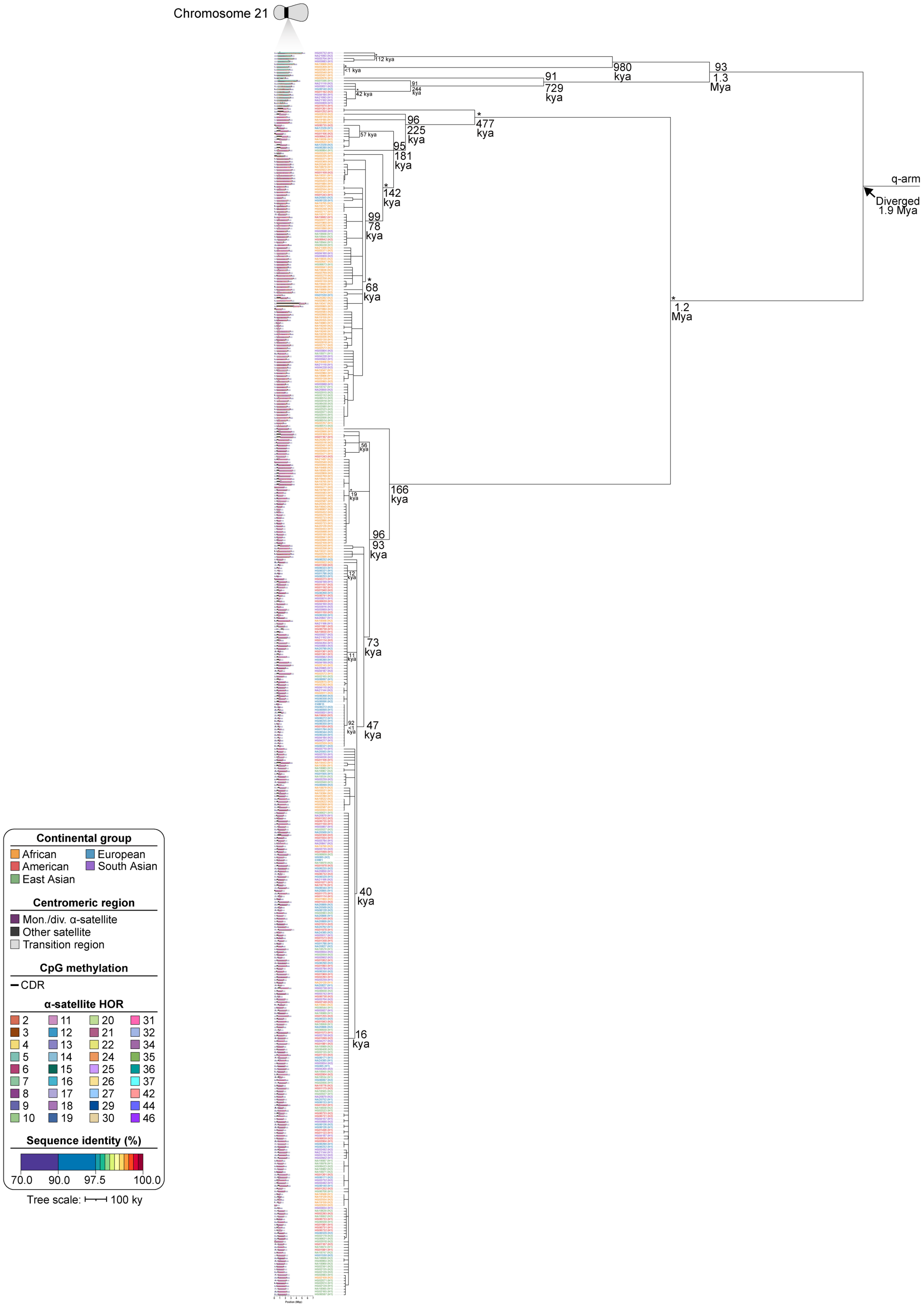


**Supplementary Figure 59. Phylogenetic reconstruction of chromosome 21 centromere haplotypes across the HGSVC and HPRC datasets.** Maximum-likelihood phylogenetic tree depicting the p- and q-arm topologies along with estimated divergence times for the chromosome 21 (*D21Z1*) α-satellite HOR arrays from 296 diverse human genomes. Asterisks indicate nodes with 100% bootstrap support, and nodes with 90-99% bootstrap support are indicated numerically. Nodes without an asterisk or number have bootstrap support <90%. The overall topology of the tree is preserved with the inclusion of complete centromeres recently assembled by the HPRC.

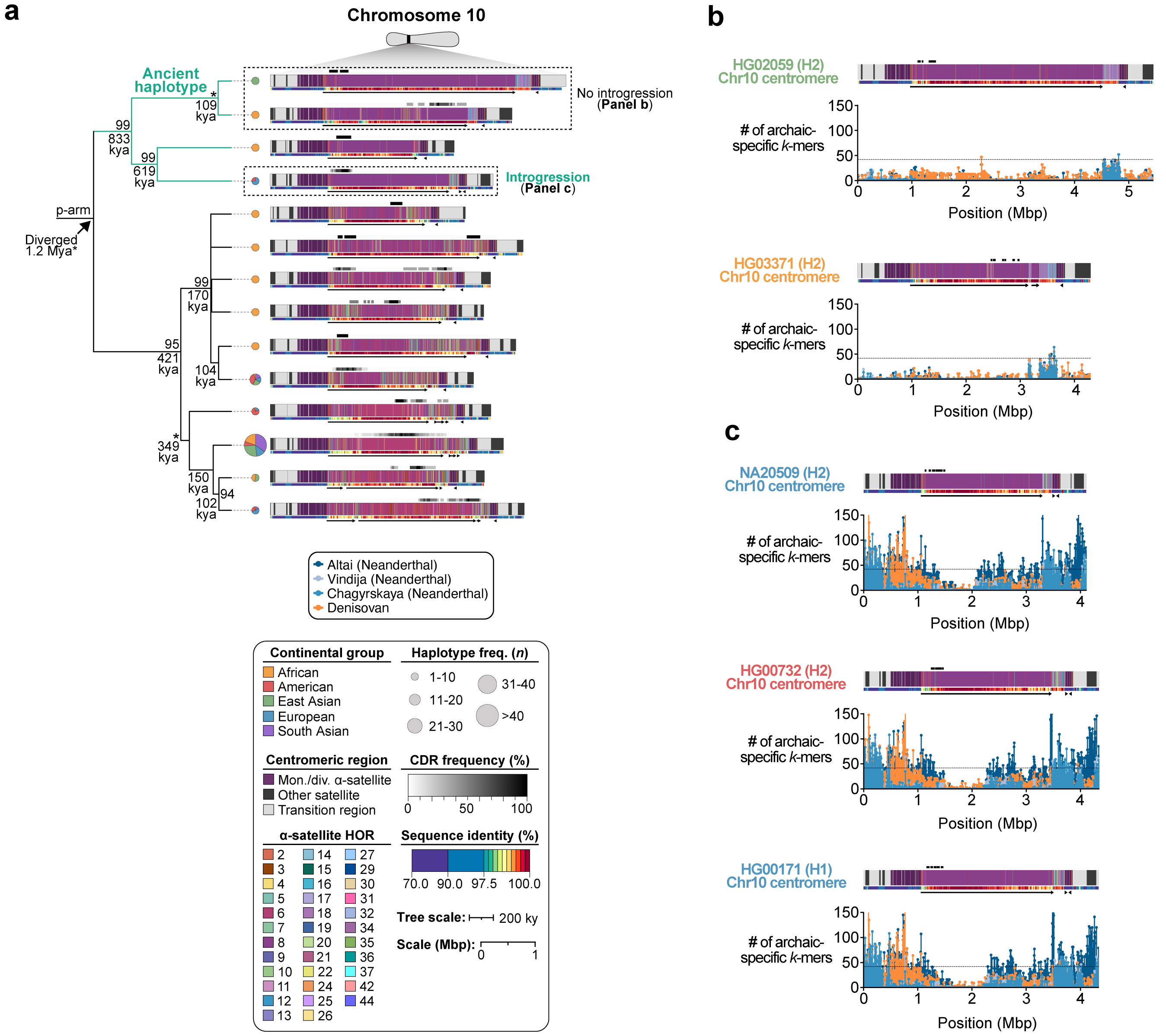


**Supplementary Figure 60. Evidence of archaic introgression in a subset of centromeres from chromosome 10. a)** Maximum-likelihood phylogenetic tree depicting the p-arm topology for chromosome 10 centromere haplogroups along with estimated divergence times and haplotype frequencies. Asterisks indicate nodes with 100% bootstrap support, and nodes with 90-99% bootstrap support are indicated numerically. Nodes without an asterisk or number have bootstrap support <90%. **b,c)** Distribution of archaic-specific DNA *k*-mers across 20 kbp windows of the **b)** HG02059 (H2), HG03371 (H2) or **c)** NA20509 (H2), HG00732 (H2), or HG00171 (H1) chromosome 10 centromeric regions using archaic short-read data from four genomes: Altai (Neanderthal), Vindija (Neanderthal), Chagryskaya (Neanderthal), and Denisovan. The dashed line indicates the genome-wide *k*-mer count cutoff, 42, which corresponds to the genome-wide archaic introgression fraction[10]. While the two ancient haplotypes shown in Panel b do not have an enrichment of archaic DNA *k*-mers, the haplotypes in Panel c do have an enrichment, suggesting this haplotype resulted, at least in part, due to introgression with Neanderthals and Denisovans.

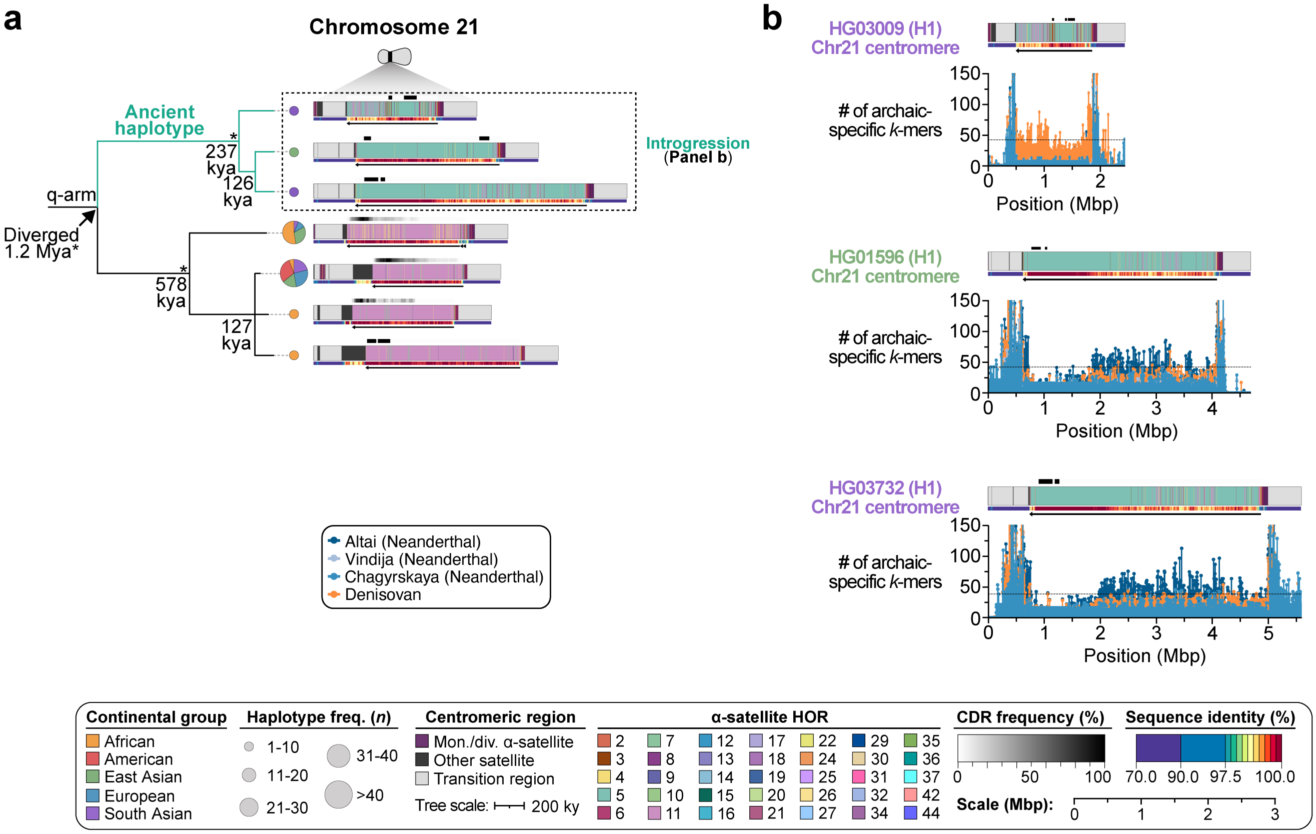


**Supplementary Figure 61. Evidence of archaic introgression in a subset of centromeres from chromosome 21. a)** Maximum-likelihood phylogenetic tree depicting the p-arm topology for chromosome 21 centromere haplogroups along with estimated divergence times and haplotype frequencies. Asterisks indicate nodes with 100% bootstrap support, and nodes with 90-99% bootstrap support are indicated numerically. Nodes without an asterisk or number have bootstrap support <90%. **b)** Distribution of archaic-specific DNA *k*-mers across 20 kbp windows of the HG03009 (H1), HG01596 (H1), HG03732 (H1) chromosome 21 centromeric regions using high-quality archaic short-read data from four genomes: Altai (Neanderthal), Vindija (Neanderthal), Chagryskaya (Neanderthal), and Denisovan. The dashed line indicates the genome-wide *k*-mer count cutoff, 42, which corresponds to the genome-wide archaic introgression fraction[10]. All three ancient haplotypes shown in Panel b have an enrichment of archaic DNA *k*-mers, suggesting these haplotypes emerged, at least in part, due to introgression with Neanderthals and Denisovans.

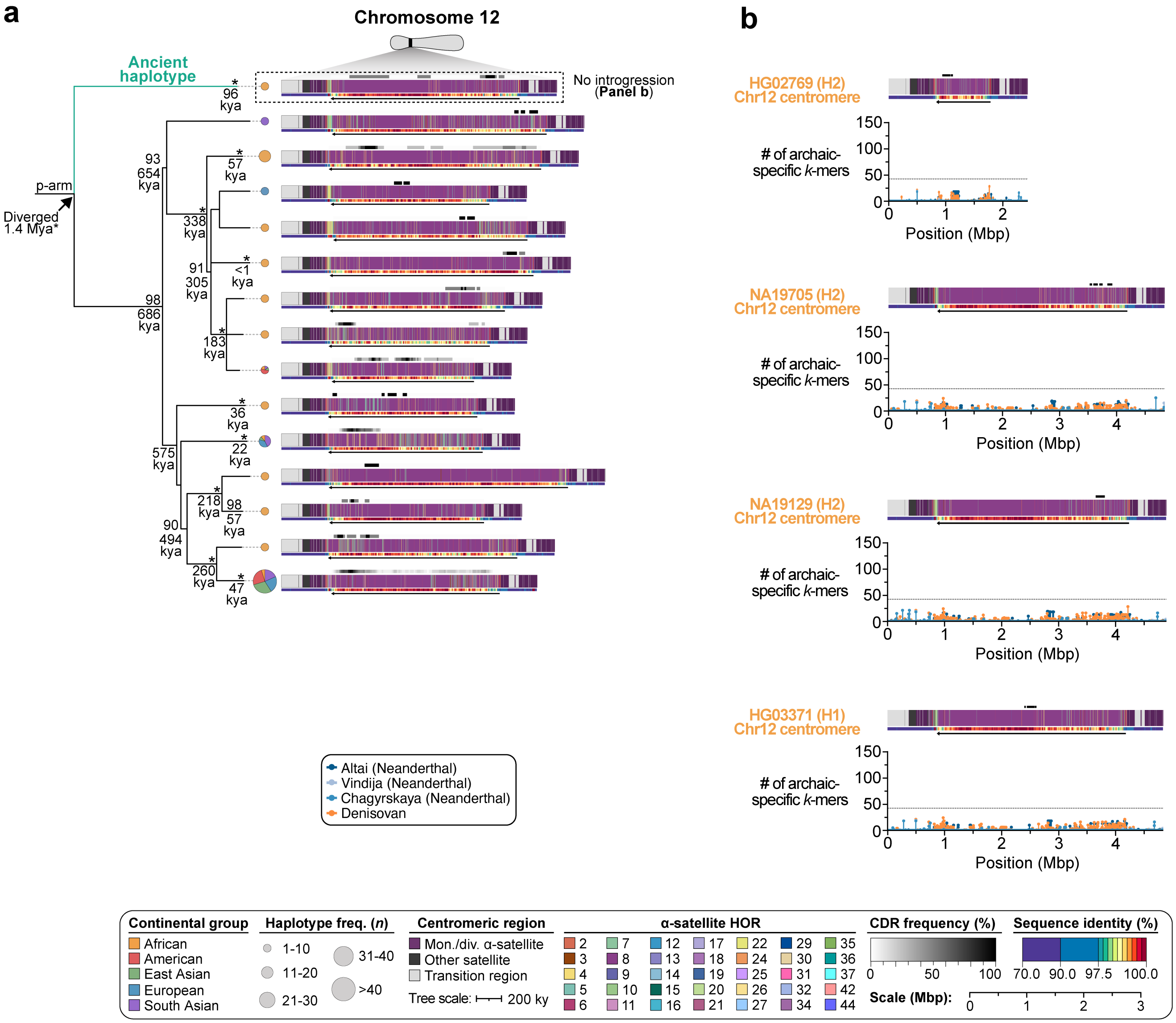


**Supplementary Figure 62. Lack of archaic introgression in ancient centromeres from chromosome 12. a)** Maximum-likelihood phylogenetic tree depicting the p-arm topology for chromosome 12 centromere haplogroups along with estimated divergence times and haplotype frequencies. Asterisks indicate nodes with 100% bootstrap support, and nodes with 90-99% bootstrap support are indicated numerically. Nodes without an asterisk or number have bootstrap support <90%. **b)** Distribution of archaic-specific DNA *k*-mers across 20 kbp windows of the **b)** HG02769 (H2), NA19705 (H2), NA19129 (H2), or HG03371 (H1) chromosome 12 centromeric regions using high-quality archaic short-read data from four genomes: Altai (Neanderthal), Vindija (Neanderthal), Chagryskaya (Neanderthal), and Denisovan. The dashed line indicates the genome-wide *k*-mer count cutoff, 42, which corresponds to the genome-wide archaic introgression fraction[10]. None of the ancient haplotypes shown in Panel b have an enrichment of archaic DNA *k*-mers, suggesting this haplotype likely arose before the out-of-Africa migration and may have been lost due to population bottlenecks.

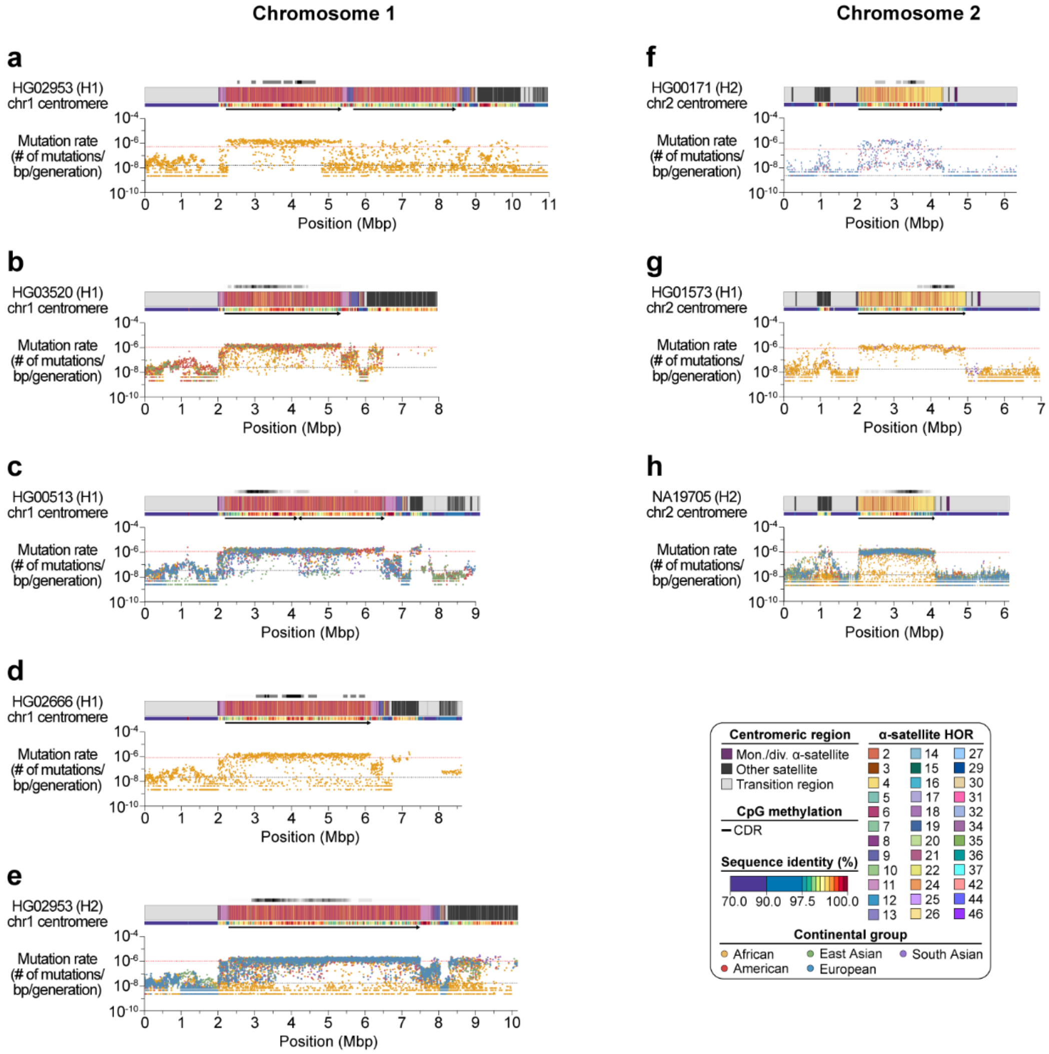


**Supplementary Figure 63. Estimated mutation rates of major centromere haplogroups from chromosomes 1 and 2. a-h)** Plots showing the estimated mutation rate across the **a-e)** chromosome 1 and **f-h)** chromosome 2 centromeric regions, separated by haplogroup. Individual data points from 10-kbp pairwise sequence alignments are shown. Mean mutation rates across the active α-satellite HOR array and flanking non-satellite sequences are shown as dotted red and black lines, respectively.

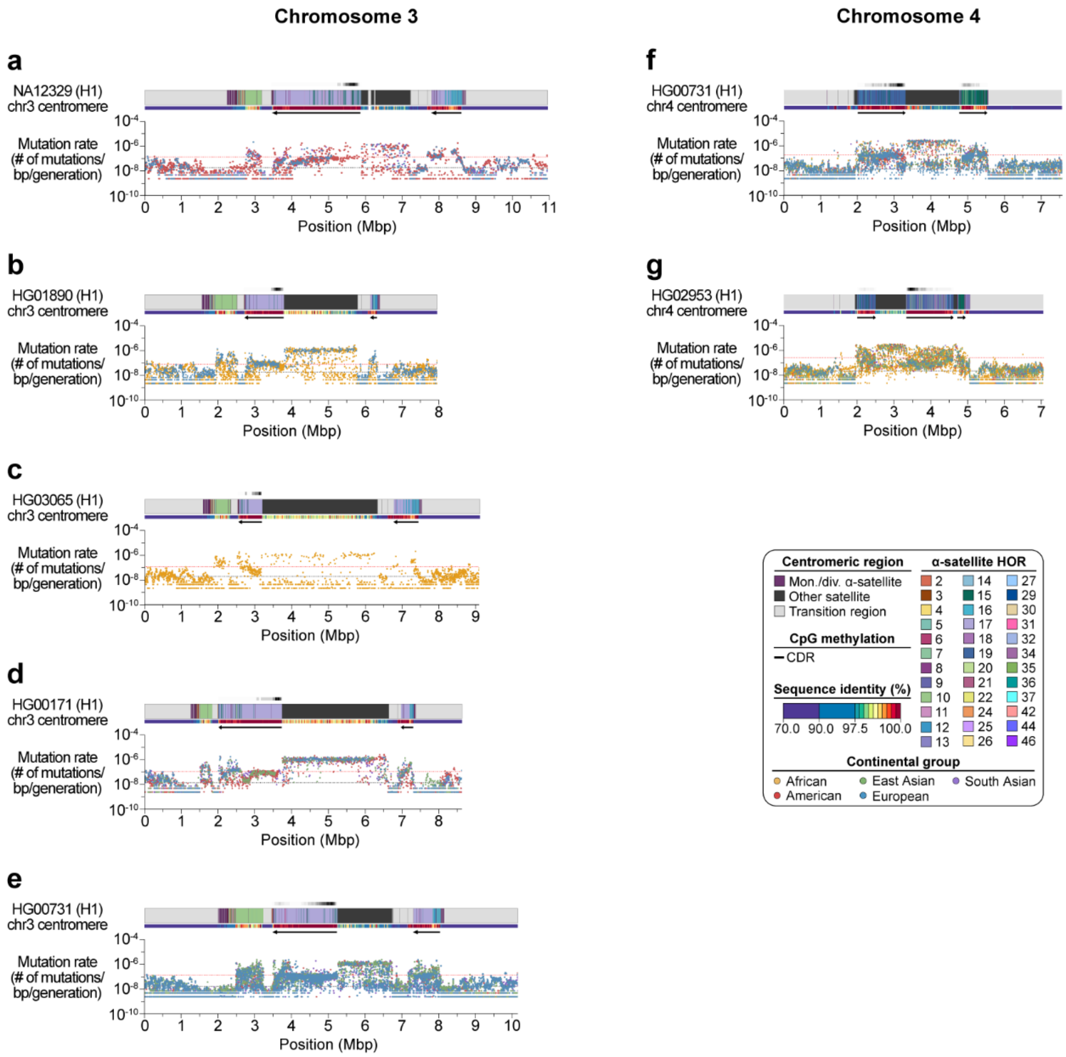


**Supplementary Figure 64. Estimated mutation rates of major centromere haplogroups from chromosomes 3 and 4. a-g)** Plots showing the estimated mutation rate across the **a-e)** chromosome 3 and **f,g)** chromosome 4 centromeric regions, separated by haplogroup. Individual data points from 10-kbp pairwise sequence alignments are shown. Mean mutation rates across the active α-satellite HOR array and flanking non-satellite sequences are shown as dotted red and black lines, respectively.

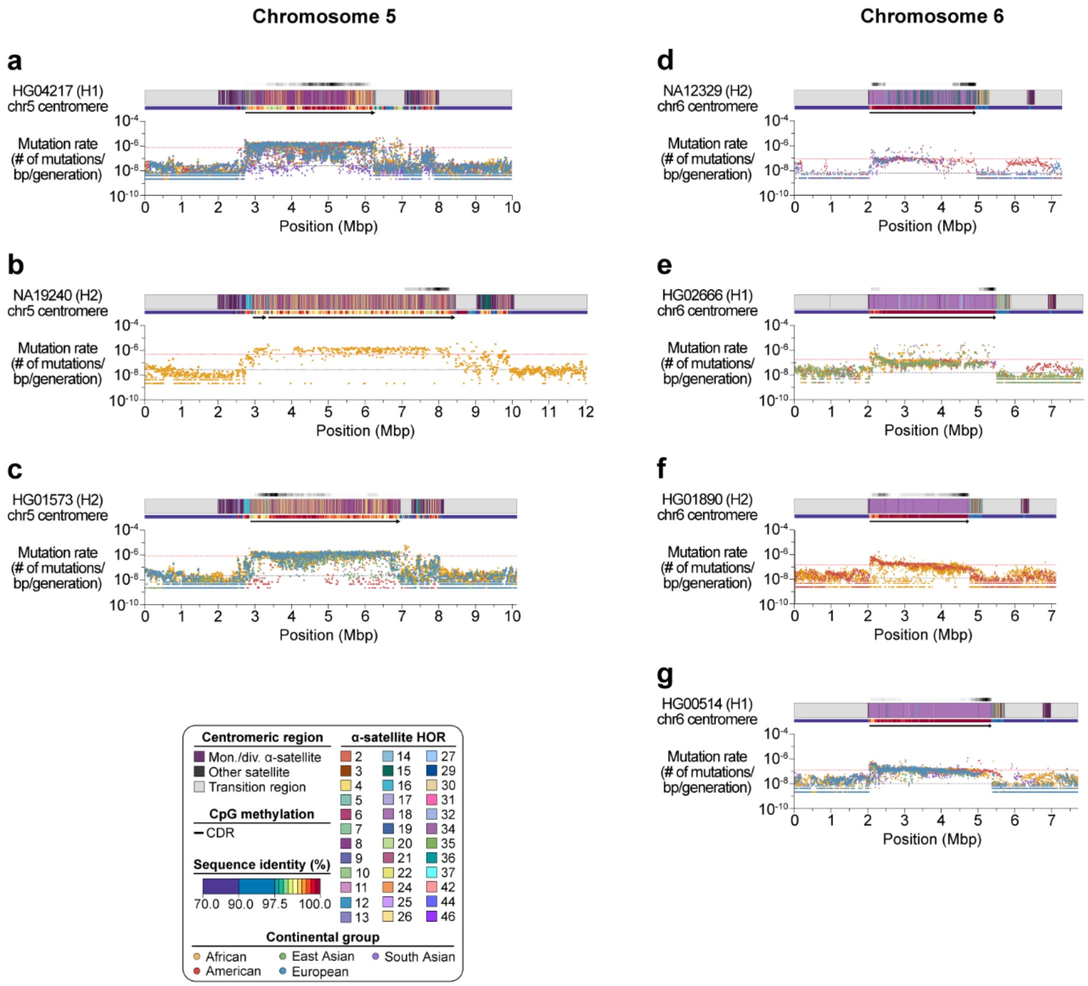


**Supplementary Figure 65. Estimated mutation rates of major centromere haplogroups from chromosomes 5 and 6. a-g)** Plots showing the estimated mutation rate across the **a-c)** chromosome 5 and **d-g)** chromosome 6 centromeric regions, separated by haplogroup. Individual data points from 10-kbp pairwise sequence alignments are shown. Mean mutation rates across the active α-satellite HOR array and flanking non-satellite sequences are shown as dotted red and black lines, respectively.

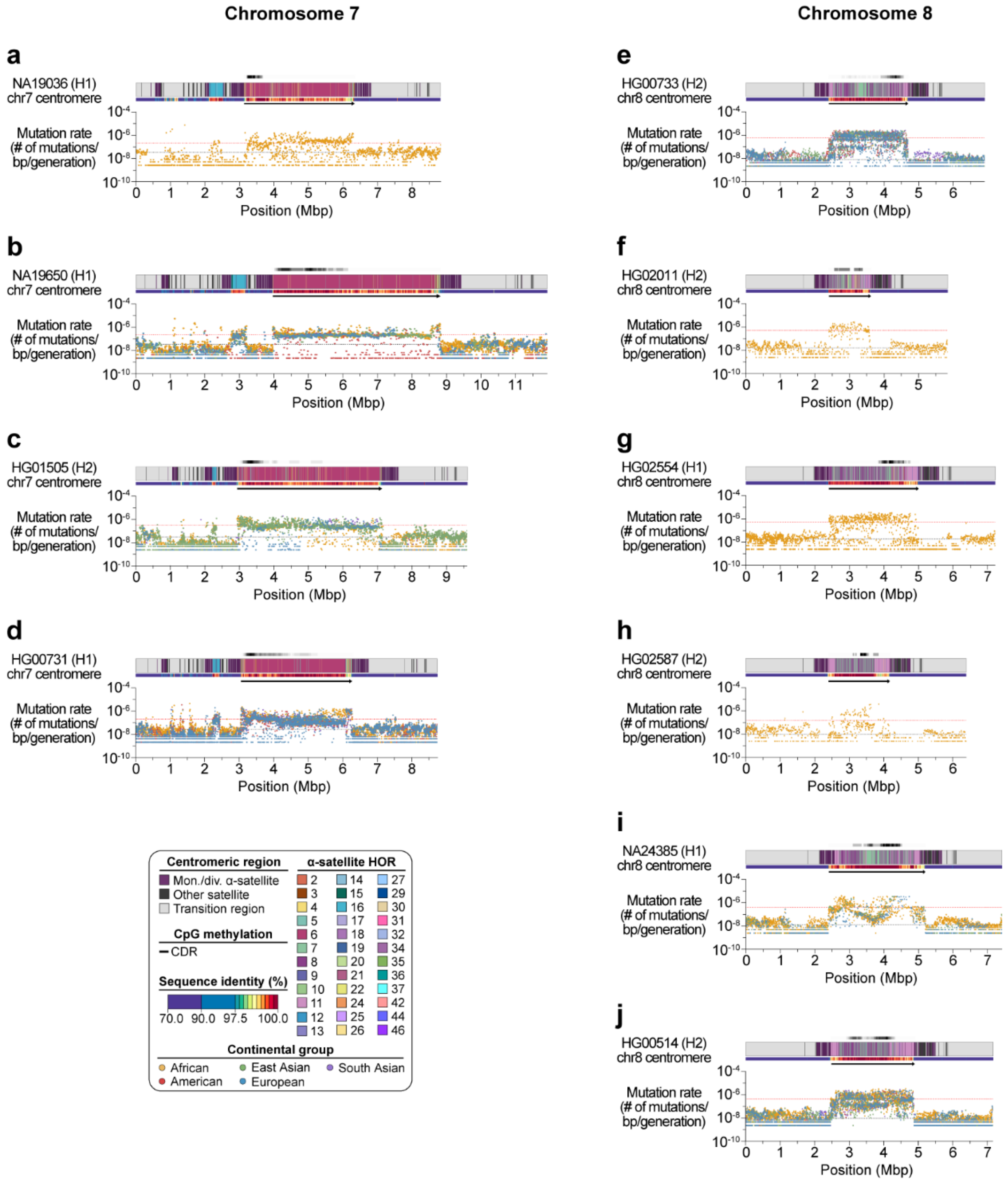


**Supplementary Figure 66. Estimated mutation rates of major centromere haplogroups from chromosomes 7 and 8. a-j)** Plots showing the estimated mutation rate across the **a-d)** chromosome 7 and **e-j)** chromosome 8 centromeric regions, separated by haplogroup. Individual data points from 10-kbp pairwise sequence alignments are shown. Mean mutation rates across the active α-satellite HOR array and flanking non-satellite sequences are shown as dotted red and black lines, respectively.

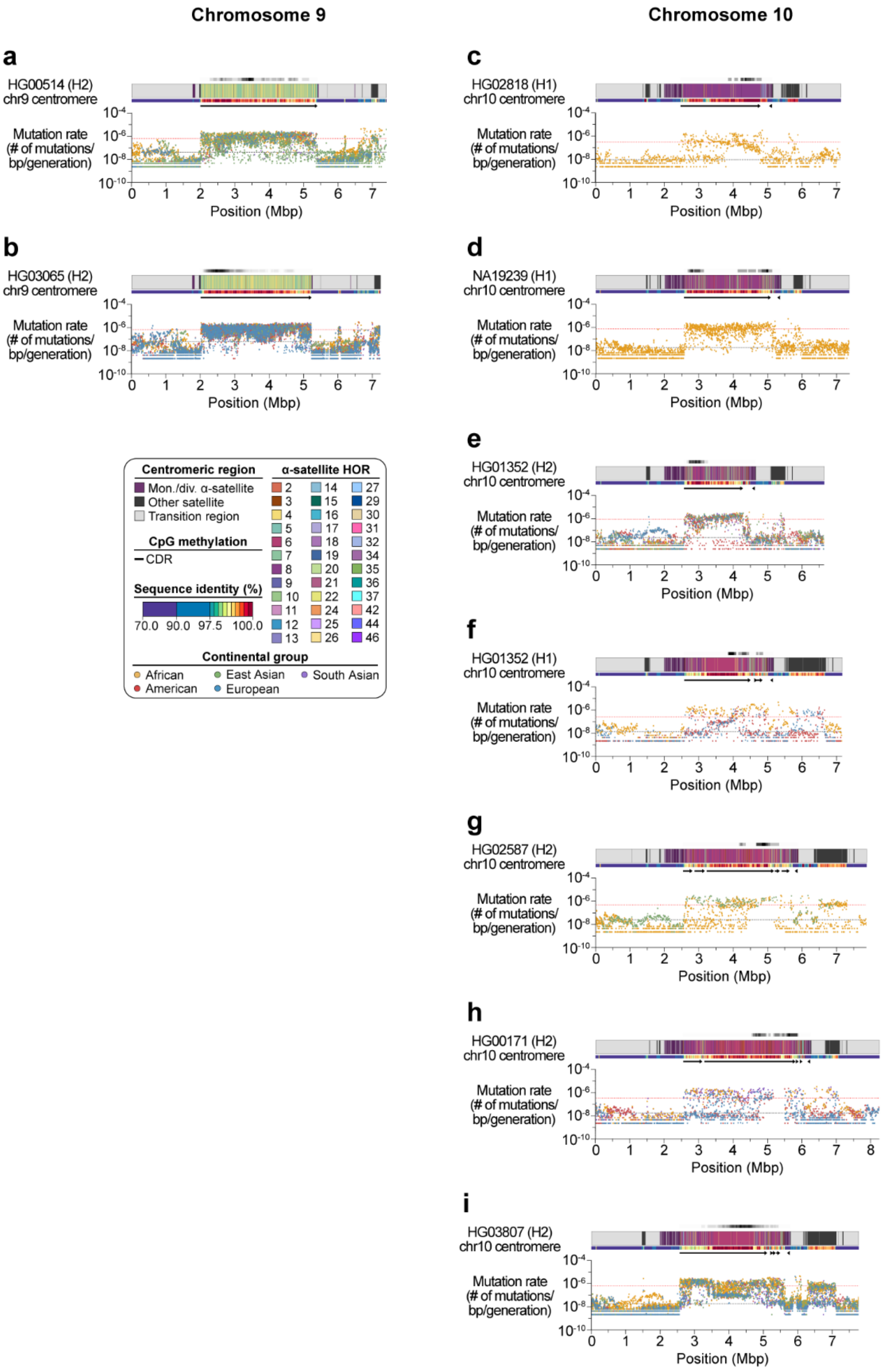


**Supplementary Figure 67. Estimated mutation rates of major centromere haplogroups from chromosomes 9 and 10. a-i)** Plots showing the estimated mutation rate across the **a,b)** chromosome 9 and **c-i)** chromosome 10 centromeric regions, separated by haplogroup. Individual data points from 10-kbp pairwise sequence alignments are shown. Mean mutation rates across the active α-satellite HOR array and flanking non-satellite sequences are shown as dotted red and black lines, respectively.

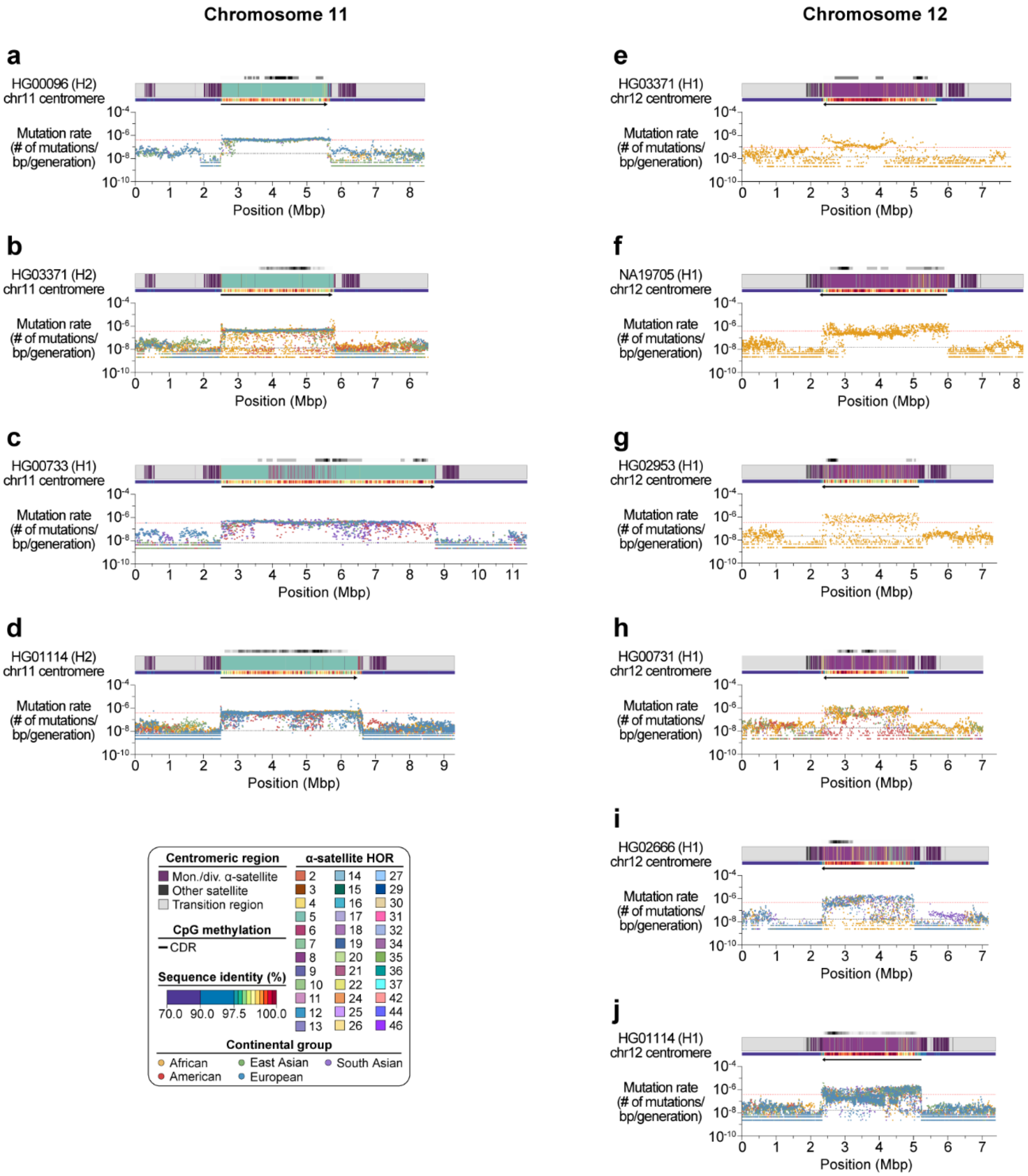


**Supplementary Figure 68. Estimated mutation rates of major centromere haplogroups from chromosomes 11 and 12. a-j)** Plots showing the estimated mutation rate across the **a-d)** chromosome 11 and **e-j)** chromosome 12 centromeric regions, separated by haplogroup. Individual data points from 10-kbp pairwise sequence alignments are shown. Mean mutation rates across the active α-satellite HOR array and flanking non-satellite sequences are shown as dotted red and black lines, respectively.

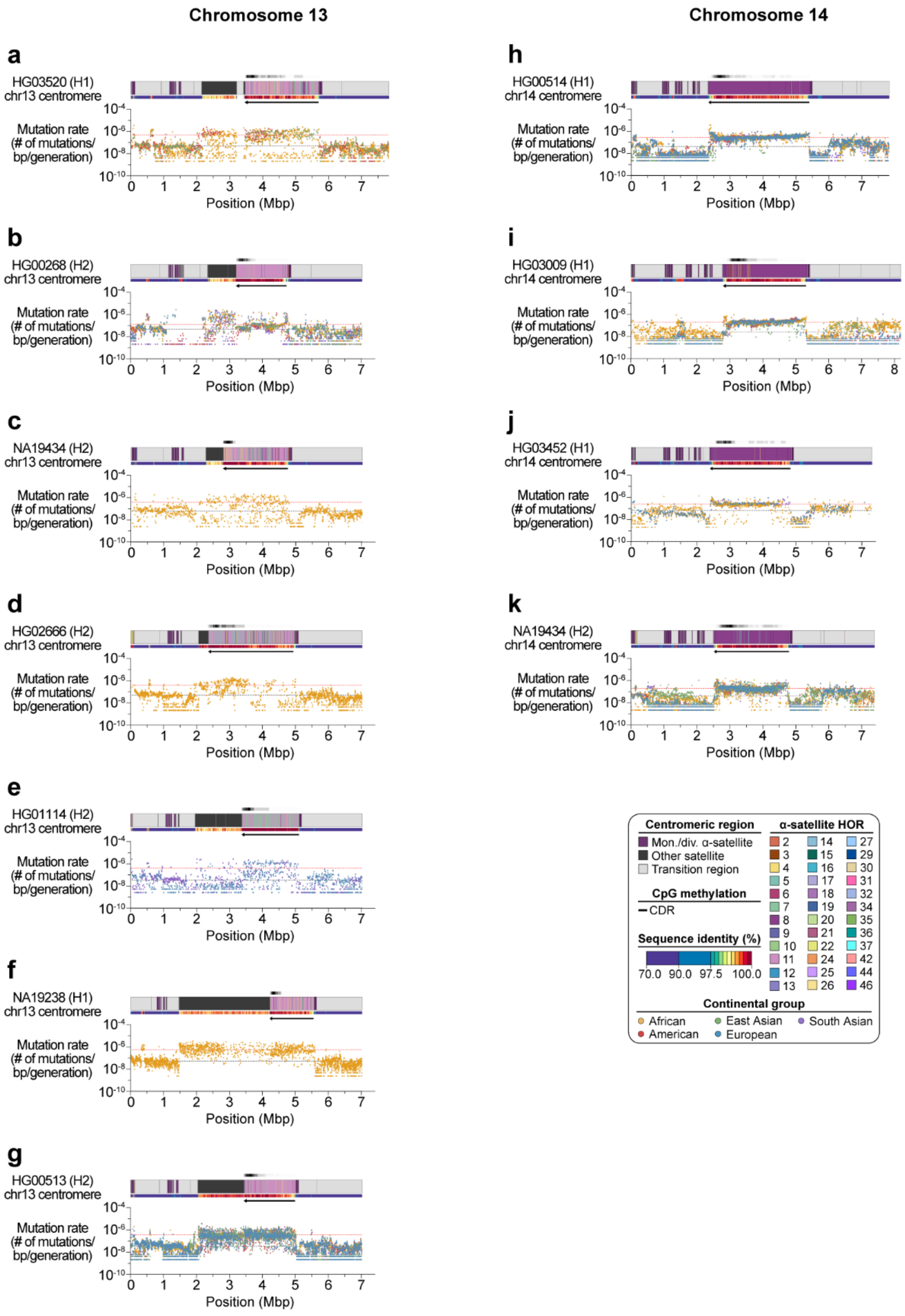


**Supplementary Figure 69. Estimated mutation rates of major centromere haplogroups from chromosomes 13 and 14. a-k)** Plots showing the estimated mutation rate across the **a-g)** chromosome 13 and **h-k)** chromosome 14 centromeric regions, separated by haplogroup. Individual data points from 10-kbp pairwise sequence alignments are shown. Mean mutation rates across the active α-satellite HOR array and flanking non-satellite sequences are shown as dotted red and black lines, respectively.

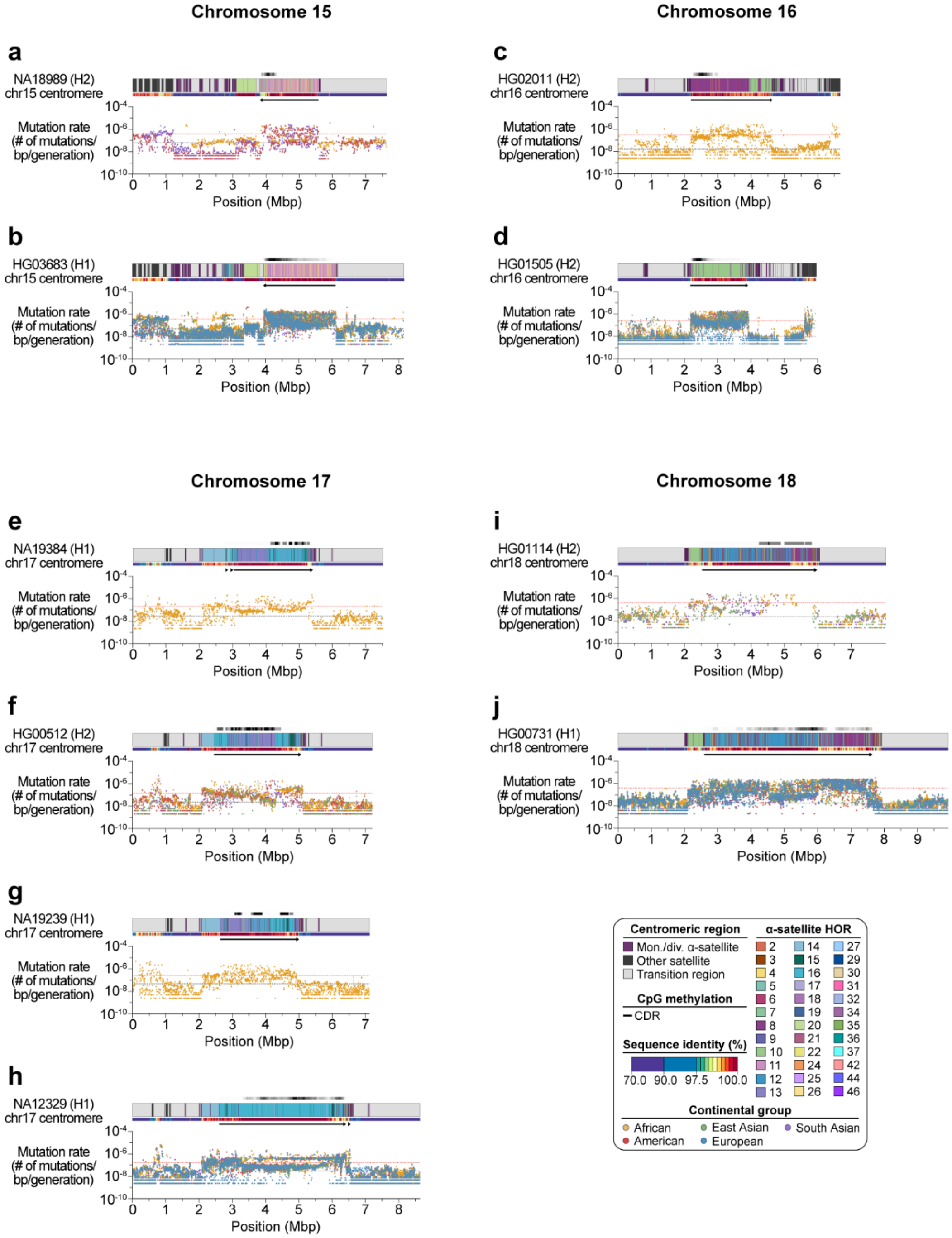


**Supplementary Figure 70. Estimated mutation rates of major centromere haplogroups from chromosomes 15-18. a-j)** Plots showing the estimated mutation rate across the **a,b)** chromosome 15, **c,d)** chromosome 16, **e-h)** chromosome 17, and **i,j)** chromosome 18 centromeric regions, separated by haplogroup. Individual data points from 10-kbp pairwise sequence alignments are shown. Mean mutation rates across the active α-satellite HOR array and flanking non-satellite sequences are shown as dotted red and black lines, respectively.

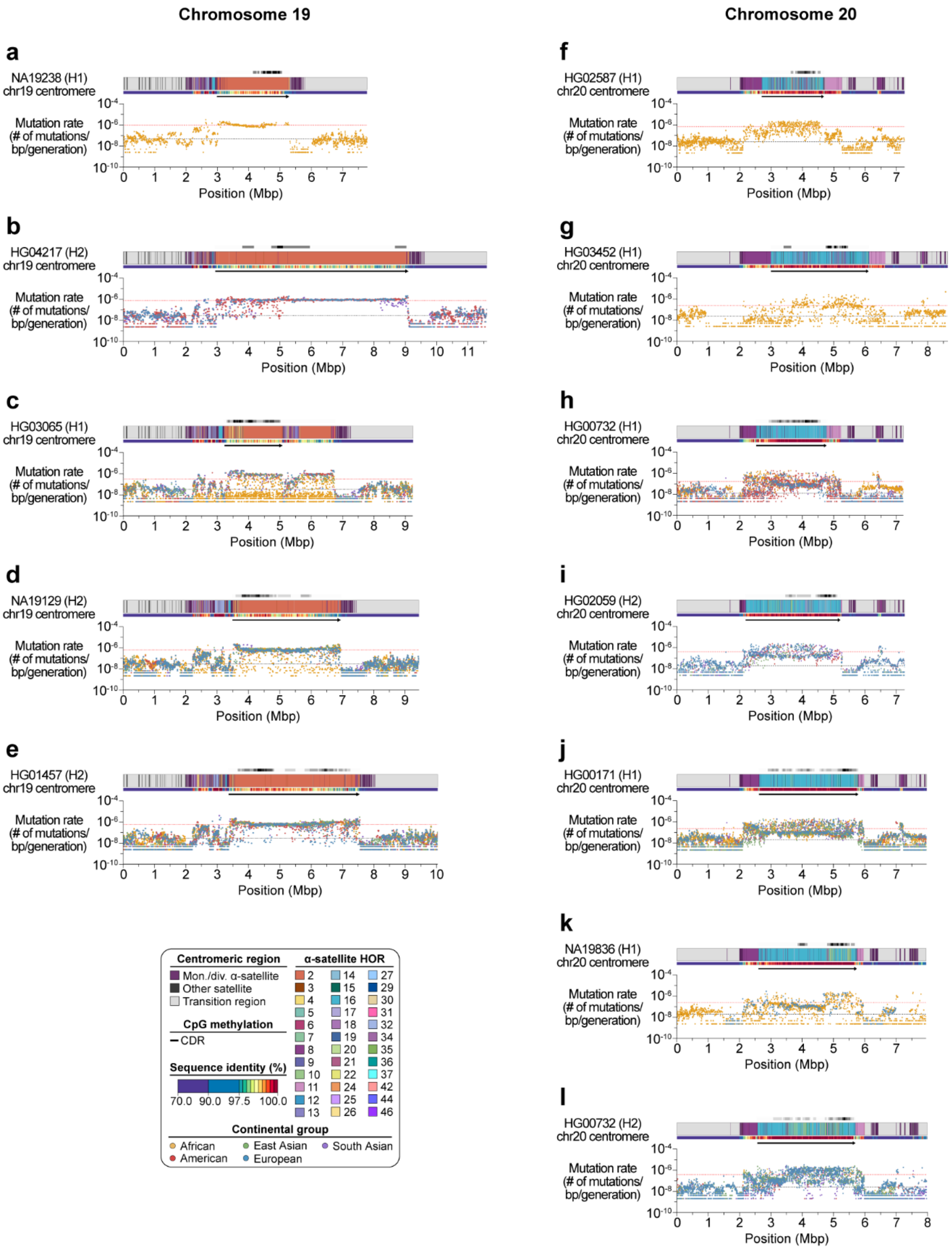


**Supplementary Figure 71. Estimated mutation rates of major centromere haplogroups from chromosomes 19 and 20. a-l)** Plots showing the estimated mutation rate across the **a-e)** chromosome 19 and **f-l)** chromosome 20 centromeric regions, separated by haplogroup. Individual data points from 10-kbp pairwise sequence alignments are shown. Mean mutation rates across the active α-satellite HOR array and flanking non-satellite sequences are shown as dotted red and black lines, respectively.

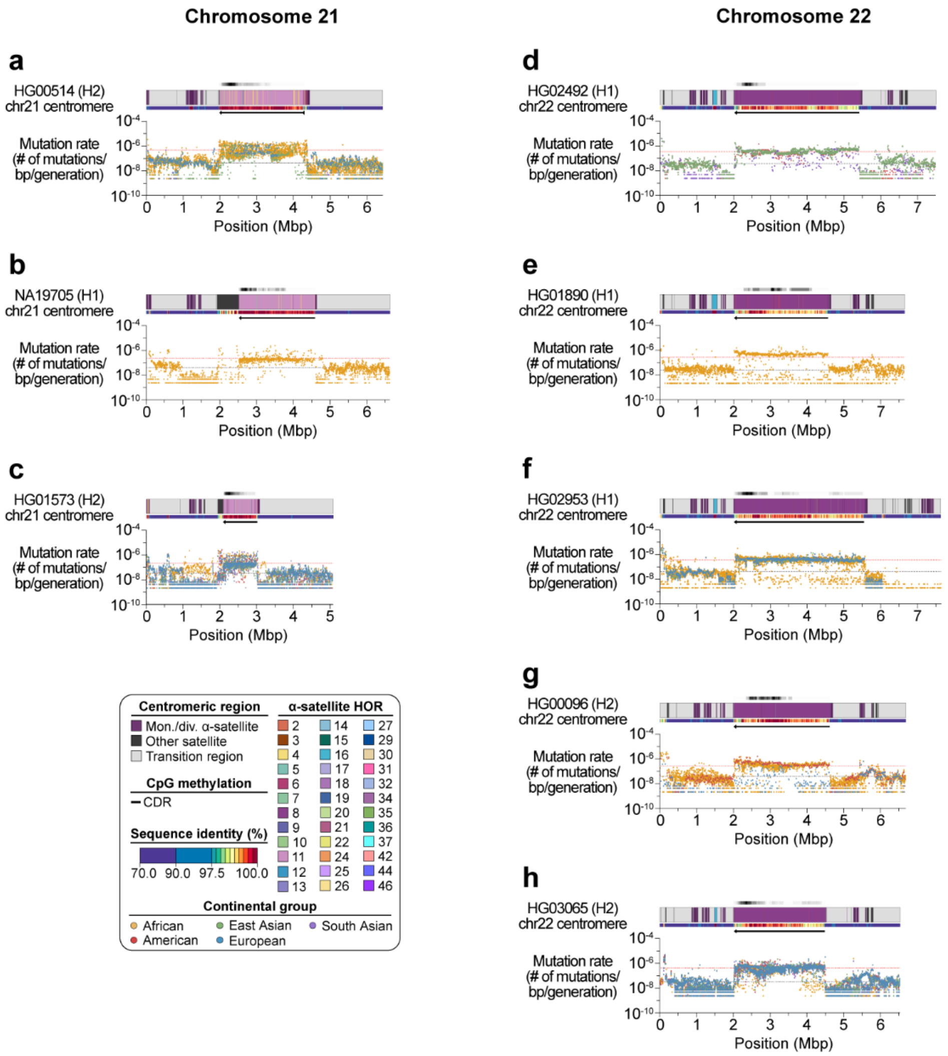


**Supplementary Figure 72. Estimated mutation rates of major centromere haplogroups from chromosomes 21 and 22. a-h)** Plots showing the estimated mutation rate across the **a-c)** chromosome 21 and **d-h)** chromosome 22 centromeric regions, separated by haplogroup. Individual data points from 10-kbp pairwise sequence alignments are shown. Mean mutation rates across the active α-satellite HOR array and flanking non-satellite sequences are shown as dotted red and black lines, respectively.

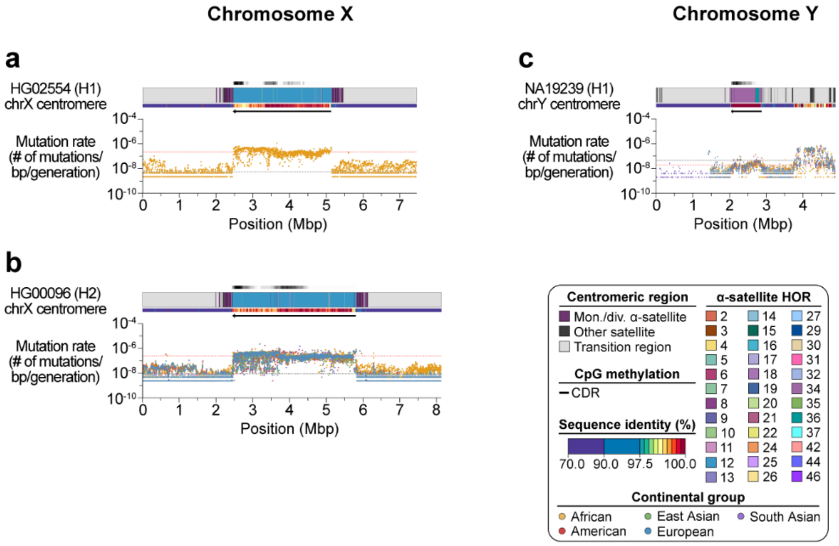


**Supplementary Figure 73. Estimated mutation rates of major centromere haplogroups from chromosomes X and Y. a-c)** Plots showing the estimated mutation rate across the **a,b)** chromosome X and **c)** chromosome Y centromeric regions, separated by haplogroup. Individual data points from 10-kbp pairwise sequence alignments are shown. Mean mutation rates across the active α-satellite HOR array and flanking non-satellite sequences are shown as dotted red and black lines, respectively.

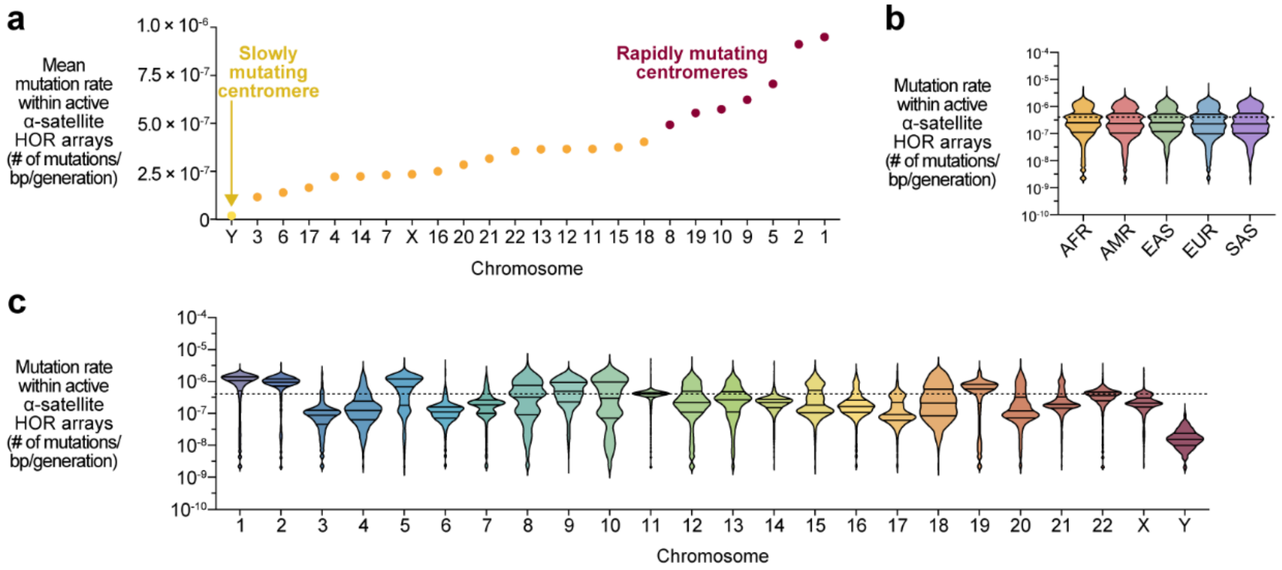


**Supplementary Figure 74. Estimated mutation rates of active α-satellite HOR arrays across chromosomes and continental groups. a)** Mean mutation rate of active α-satellite HOR arrays across chromosomes. There are seven chromosomes with rapidly mutating centromeres (red): chromosomes 1, 2, 5, 8, 9, 10, and 19. In contrast, chromosome Y is slowly mutating (yellow). **b,c)** Estimated mutation rate of active α-satellite HOR arrays, split by **b)** continental group and **c)** chromosome. Mutation rates $<10^{-10}$ mutations/bp/generation are not shown.

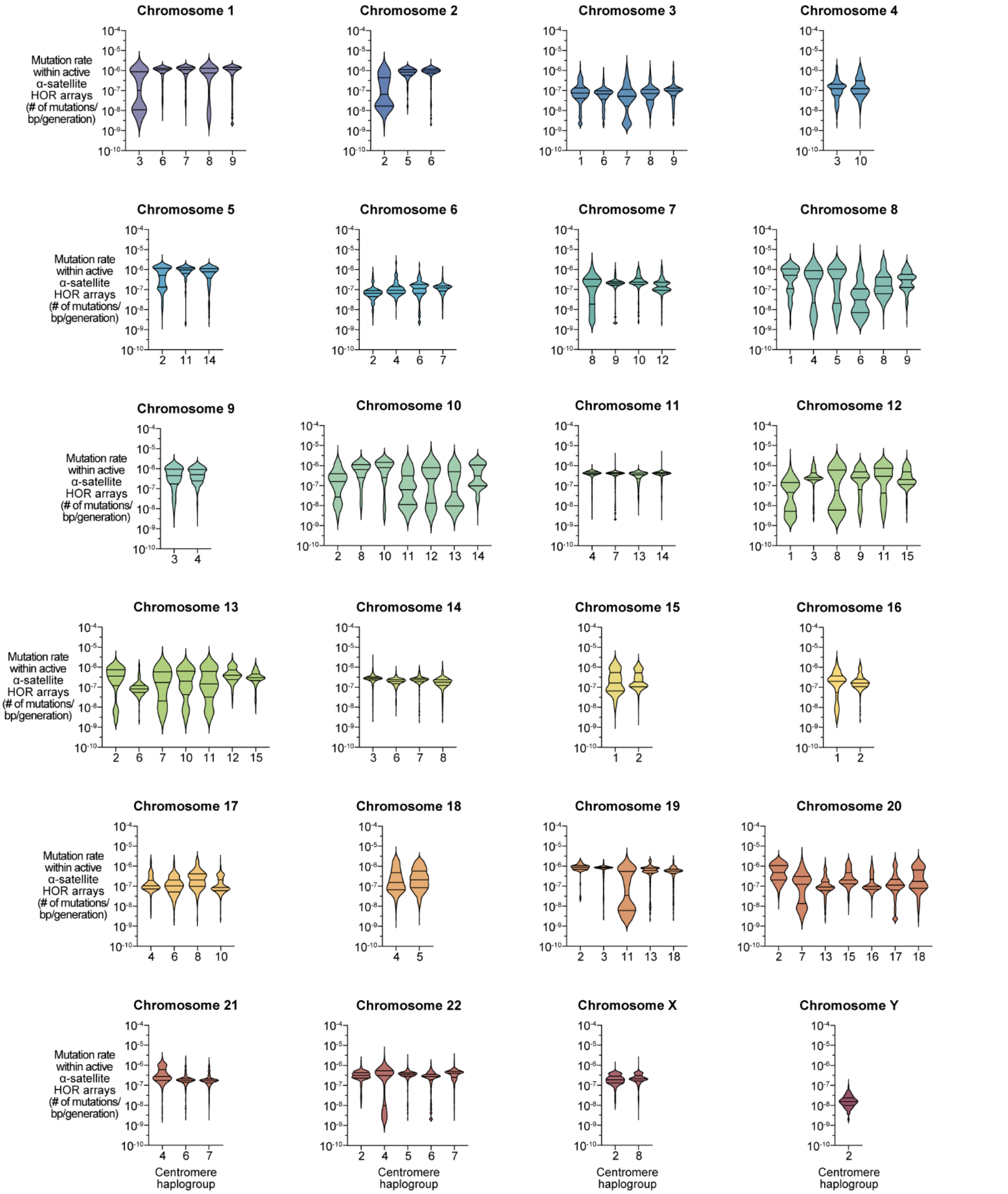


**Supplementary Figure 75. Estimated mutation rates of active α-satellite HOR arrays across centromere haplogroups.** Estimated mutation rates of α-satellite HOR arrays, split by major centromere haplogroups (defined as having ≥4 centromere haplotypes). The mutation rate across centromeric regions for each haplogroup are shown in **Supplementary Figs. 63-72**. Mutation rates $<10^{-10}$ mutations/bp/generation are not shown.

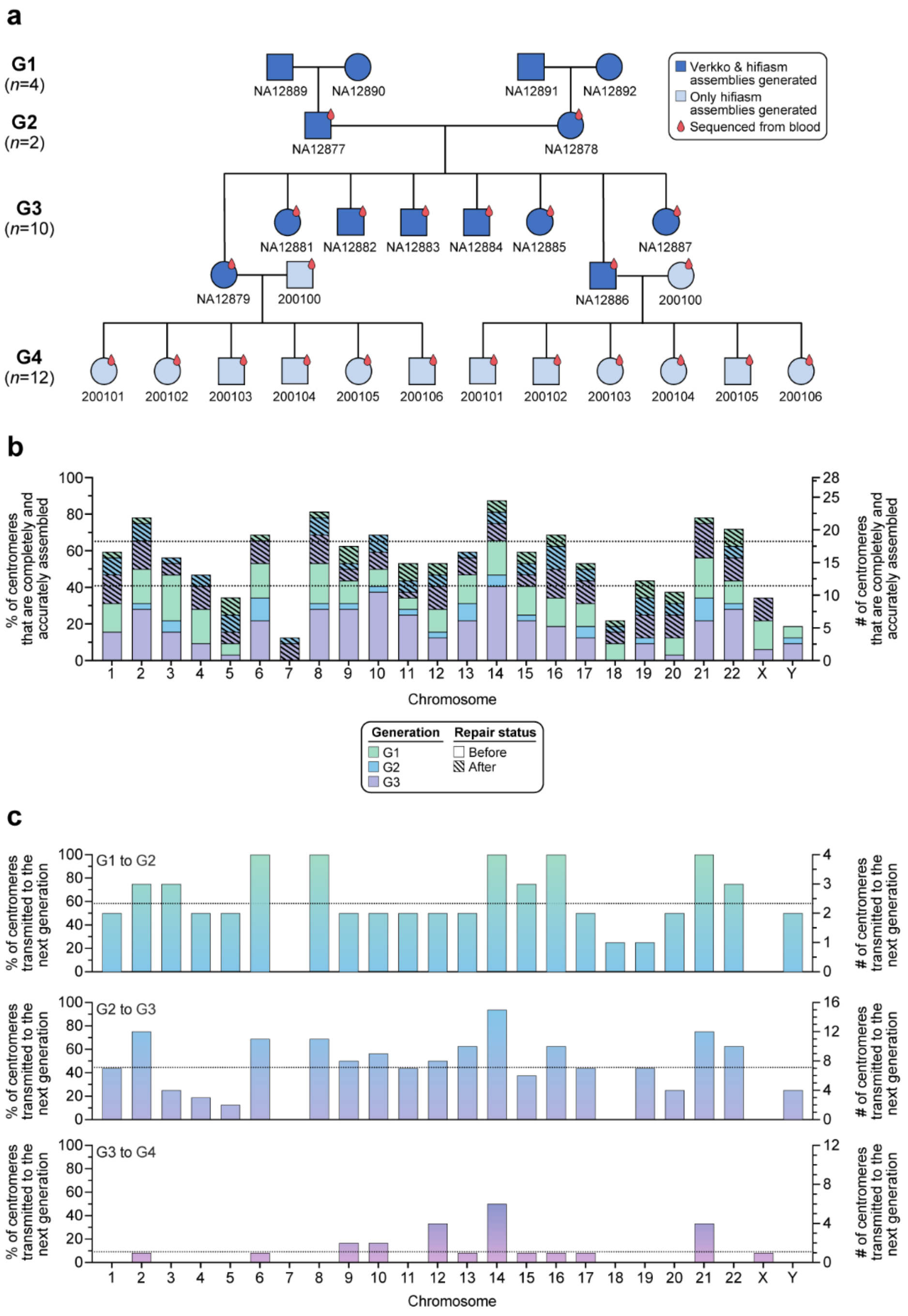


**Supplementary Figure 76. A 4-generation, 28-member pedigree enables accurate *de novo* mutation rate estimation in centromeres. a)** Pedigree of the 4-generation family (CEPH 1463)[11] and the genome assemblies generated for each member. **b)** Percentage and number of completely and accurately assembled centromeres before and after repair with AssemblyRepairer for generations 1-3 (G1-G3). Before repair, 288 centromeres (or 12.5 centromeres per chromosome) were accurately resolved on average; after repair, 483 centromeres (or 21 centromeres per chromosome) were accurately resolved on average, a 67.7% increase. **c)** Percentage and number of completely and accurately assembled centromeres transmitted to the next generation.

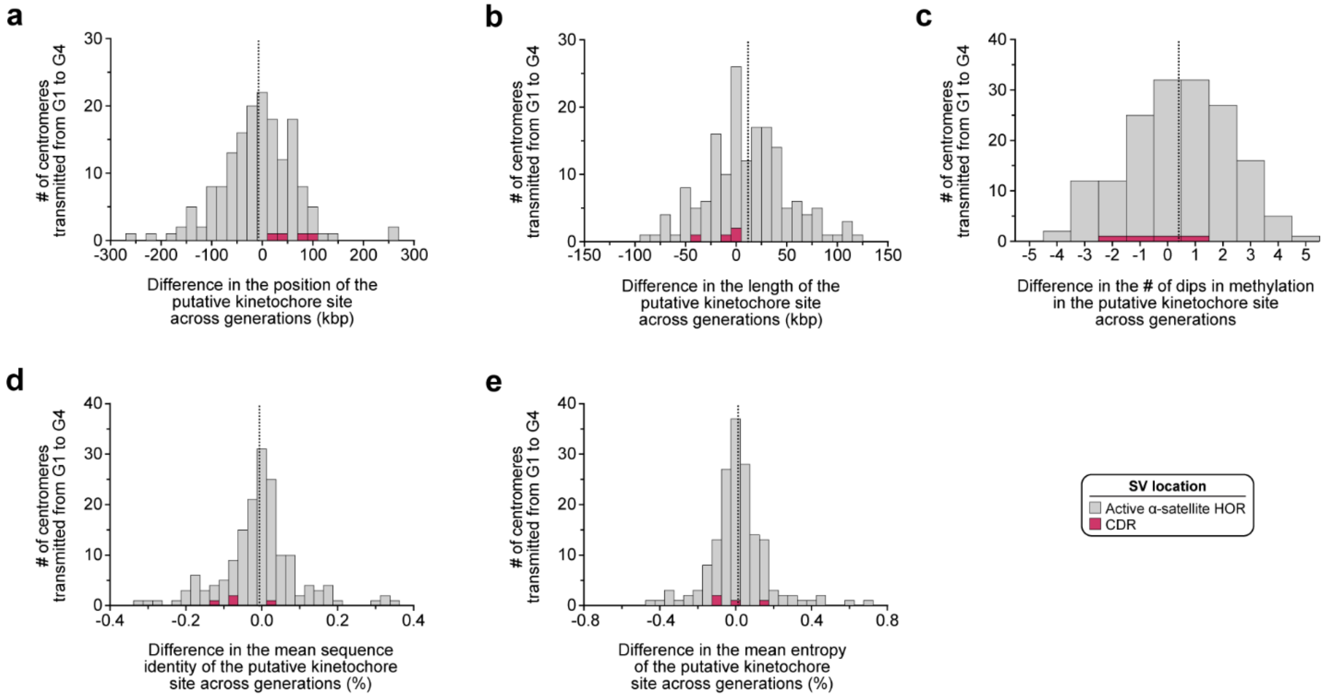


**Supplementary Figure 77. Differences in the position, length, distribution, and sequence composition of the putative kinetochore site across generations. a-e)** Changes in the **a)** position, **b)** length, **c)** number of dips in methylation, **d)** mean sequence identity, and **e)** mean entropy of the putative kinetochore site, marked by the CDR, from G1 to G4. From one generation to the next, the putative kinetochore site shifts in position, length, and sequence composition, reducing the mean sequence identity while maintaining entropy.

**SUPPLEMENTARY TABLES**

**1. Summary of the 320 diverse human genomes analyzed in this study.**

**2. Sequence validation of 2,110 centromeres from 65 diverse human genomes.**

**3. Estimated QV of 2,110 centromeres from 65 diverse human genomes.**

**4. Quantification of novel active α-satellite HOR variants in 2,110 centromeres from 65 diverse human genomes.**

**5. Validation of novel active α-satellite HOR variants across HGSVC and HPRC genome assemblies.**

**6. Length of active α-satellite HOR arrays in 2,110 completely and accurately assembled centromeres.**

**7. Frequency of active α-satellite HOR variants among the CHM13, CHM1, and 2,110 centromeres.**

**8. Frequency of active α-satellite HOR variants among the CHM13, CHM1, and 2,110 centromeres, separated by genome.**

**9. Length of CDRs in 2,110 completely and accurately assembled centromeres.**

**10. Mobile element insertions among 2,110 centromeres.**

**11. Mean sequence identity and entropy of active α-satellite HOR arrays and CDRs among 2,110 centromeres.**

**12. Estimated mutation rates of centromeric regions among 2,110 centromeres.**

**13. Sequence validation of 483 centromeres from a 4-generation, 28-member pedigree.**

**14. Estimated QV of 483 centromeres from a 4-generation, 28-member pedigree.**

**15. 244 transmitted centromeres across 4 generations of a 28-member pedigree.**

**16. *De novo* SNVs among centromeres transmitted across generations.**

**17. Estimated *de novo* mutation rates of centromeric regions transmitted across generations.**

**18. *De novo* SVs among centromeres transmitted across generations.**

**19. Estimated *de novo* SV mutation rates of centromeric regions transmitted across generations.**

**20. Variation in the position, length, distribution, and sequence composition of the putative kinetochore site across generations.**

**21. Datasets generated and/or used in this study.**
